# Discovery of Alkenyl Oxindole as a Novel PROTAC Moiety for Targeted Protein Degradation via CRL4^DCAF11^ Recruitment

**DOI:** 10.1101/2024.02.15.580430

**Authors:** Ying Wang, Tianzi Wei, Man Zhao, Aima Huang, Fan Sun, Lu Chen, Risheng Lin, Yubao Xie, Ming Zhang, Shiyu Xu, Zhihui Sun, Liang Hong, Rui Wang, Ruilin Tian, Guofeng Li

**Affiliations:** School of Pharmacy, Shenzhen University Medical School, Shenzhen University, Shenzhen 518055, China; Key University Laboratory of Metabolism and Health of Guangdong, Department of Medical Neuroscience, School of Medicine, Southern University of Science and Technology, Shenzhen 518055, China; Guangdong Key Laboratory of Chiral Molecule and Drug Discovery, School of Pharmaceutical Sciences, Sun Yat-sen University, Guangzhou 510006, China; Institute of Materia Medica and Research Unit of Peptide Science, Chinese Academy of Medical Sciences & Peking Union Medical College, Beijing 100050, China

**Keywords:** PROTAC, DCAF11, Alkenyl oxindole, BRD4, Antitumor

## Abstract

Alkenyl oxindoles have been characterized as autophagosome-tethering compounds (ATTECs), which can target mutant huntingtin protein (mHTT) for lysosomal degradation. In order to expand the application of alkenyl oxindoles for targeted protein degradation, we designed and synthesized a series of hetero-bifunctional compounds by conjugating different alkenyl oxindoles with the BRD4 inhibitor JQ1. Through structure-activity relationship study, we successfully developed JQ1-alkenyl oxindole conjugates that potently degrade BRD4. Unexpectedly, we found that these molecules degrade BRD4 through the ubiquitin-proteasome system, rather than the autophagy-lysosomal pathway. Using pooled CRISPR interference (CRISPRi) screening, we revealed that JQ1-alkenyl oxindole conjugates recruit the E3 ubiquitin ligase complex CRL4^DCAF11^ for substrate degradation. Furthermore, we validated the most potent hetero-bifunctional molecule HL435 as a promising drug-like lead compound to exert antitumor activity both *in vitro* and *in vivo*. Our research provides new employable PROTAC moieties for targeted protein degradation, providing new possibilities for drug discovery.

## 1. Introduction

Targeted protein degradation (TPD) has emerged as a promising approach for drug discovery. It uses multispecific small molecules to selectively recognize target proteins, facilitating their degradation via cell’s intrinsic protein degradation pathways^1^. Compared with traditional inhibitors, TPD drugs possess the unique ability to not only inhibit protein activity but also facilitate the degradation of target proteins. This dual functionality empowers TPD drugs to elicit stronger therapeutic effects and holds promise for targeting proteins that were previously deemed “undruggable”^2–4^. Currently, TPD strategies primarily utilize two major degradation pathways: the ubiquitin-proteasome system and the lysosomal degradation pathway^5–7^. According to mechanism of action, the major TPD strategies include proteolysis targeting chimeras (PROTACs), lysosome targeting chimeras (LYTACs) and autophagy targeting chimeras (AUTACs)^8–11^. Among them, PROTAC technology is the most extensively studied and has achieved significant breakthroughs. It has been successfully applied to degrade more than 100 target proteins, including those previously considered “undruggable”^12^. Moreover, more than 20 PROTACs are currently undergoing clinical trials since 2019^13–16^, indicating that PROTAC technology is a promising therapeutic strategy.

PROTACs are heterobifunctional molecules consisting of two ligand domains joined by a chemical linker. One ligand domain binds to the protein target of interest, and the other ligand recruits an E3 ubiquitin ligase. By engaging both the target protein and E3 ligase simultaneously, PROTACs facilitate the polyubiquitination and proteasomal degradation of the target protein^17^. While over 600 E3 ligases have been identified in the human genome^18^, only a small fraction (< 3%) have been successfully recruited by PROTACs^19^. The most commonly utilized E3 ligases include CRBN, VHL, MDM2 and IAPs. More recently, KEAP1^20, 21^, RNF114^22, 23^, DCAF15^24^ and DCAF16^25^ have expanded the toolbox of accessible E3 domains. However, the vast majority (> 90%) of reported PROTAC molecules continue to rely primarily on just two ligases: CRBN and VHL^12^. This narrow E3 diversity poses a major challenge, as the development and targeting potential of PROTAC degraders is constrained by the limited pool of recruited E3s. Therefore, broadening the range of E3 ligases that can be engaged by small molecule ligands could unlock new avenues to potentially degrade a wider range of protein targets, as well as circumvent the acquired drug resistance that caused by mutations in certain E3 ligases^26, 27^.

The alkenyl oxindole framework is commonly found in synthetic or natural compounds that exhibit a wide range of biological activities and have attracted research interests from pharmacologists and chemists^28^. Many alkenyl oxindoles have been developed as lead compounds or marketed drugs against tumors, such as sunitinib^29–31^. Recently, two alkenyl oxindoles (**10O5** and **AN1**) were found to act as molecular glues that tether mHTT to LC3, leading to the autophagy-lysosomal degradation of mHTT^32^. This prompted us to test whether this strategy can be expanded to degrade other substrates. To this end, we synthesized a series of heterobifunctional molecules by linking JQ1 with different alkenyl oxindoles, followed by assessing their targeted degradation activity. This led to the identification of HL435, a highly potent alkenyl oxindole-based BRD4 degrader. However, when we investigated the protein degradation mechanism of HL435, we found that it degraded BRD4 through the ubiquitin-proteasome system rather than the autophagy-lysosomal pathway. Based on this unexpected finding, we hypothesized that alkenyl oxindoles may act as novel E3 ligase ligands. To verify our hypothesis, we performed a pooled CRISPR interference (CRISPRi) screen, from which we revealed that the E3 ligase complex CRL4^DCAF11^ is in charge of HL435-induced proteasomal degradation of BRD4. We further validated the anti-tumor efficacy of HL435 both *in vitro* and *in vivo*. Overall, we discovered that alkenyl oxindoles can act as recruitment moiety for CRL4^DCAF11^ and developed alkenyl oxindole-based PROTAC molecules with high degradation efficiency and anti-tumor effects, expanding the toolbox of E3 ligases available for PROTAC drug development.

## 2. Results

### 2.1. Compounds development and structure-activity relationship studies on alkenyl oxindole-based hetero-bifunctional degraders

To explore the potential of alkenyl oxindole for target protein degradation, we designed and synthesized a series of hetero-bifunctional molecules by connecting JQ1 with different alkenyl oxindoles using various linkers, followed by examining their ability to degrade BRD4 (Table 1 and Figure S1). Firstly, JQ1 and the reported alkenyl oxindole (**10O5**) were connected directly with saturated or unsaturated alkane chains of different lengths to afford compounds **H1-H4**. However, they had little ability to degrade BRD4. When PEG linker was used to replace the alkane chain (**H5**), the degradation of BRD4 was observed at a concentration of 1.0 μM. Furthermore, the direct connection of linker and **10O5** through amide bond (**H6**) significantly improved the degradation ability. Therefore, we conducted a subsequent structural-activity study on the alkenyl oxindole moiety using the linker of compound **H6**. Subsequently, we examined the degradation activity of JQ1-alkenyl oxindole conjugates constructed from different alkenyl oxindole derivatives, including the addition of electron-poor substituents on the phenyl group or the benzo moiety of oxindole core (Table 1, **H7**- **H28**). The results showed that enhanced degradation activity can be achieved by substituting the alkenyl oxindole with trifluoromethyl group (Table 1, R = 6-CF_3_). These modifications led to the development of HL435 (**H27**), an excellent BRD4 degrader with a degradation efficiency > 99% at 1.0 μM.

**Table 1:**
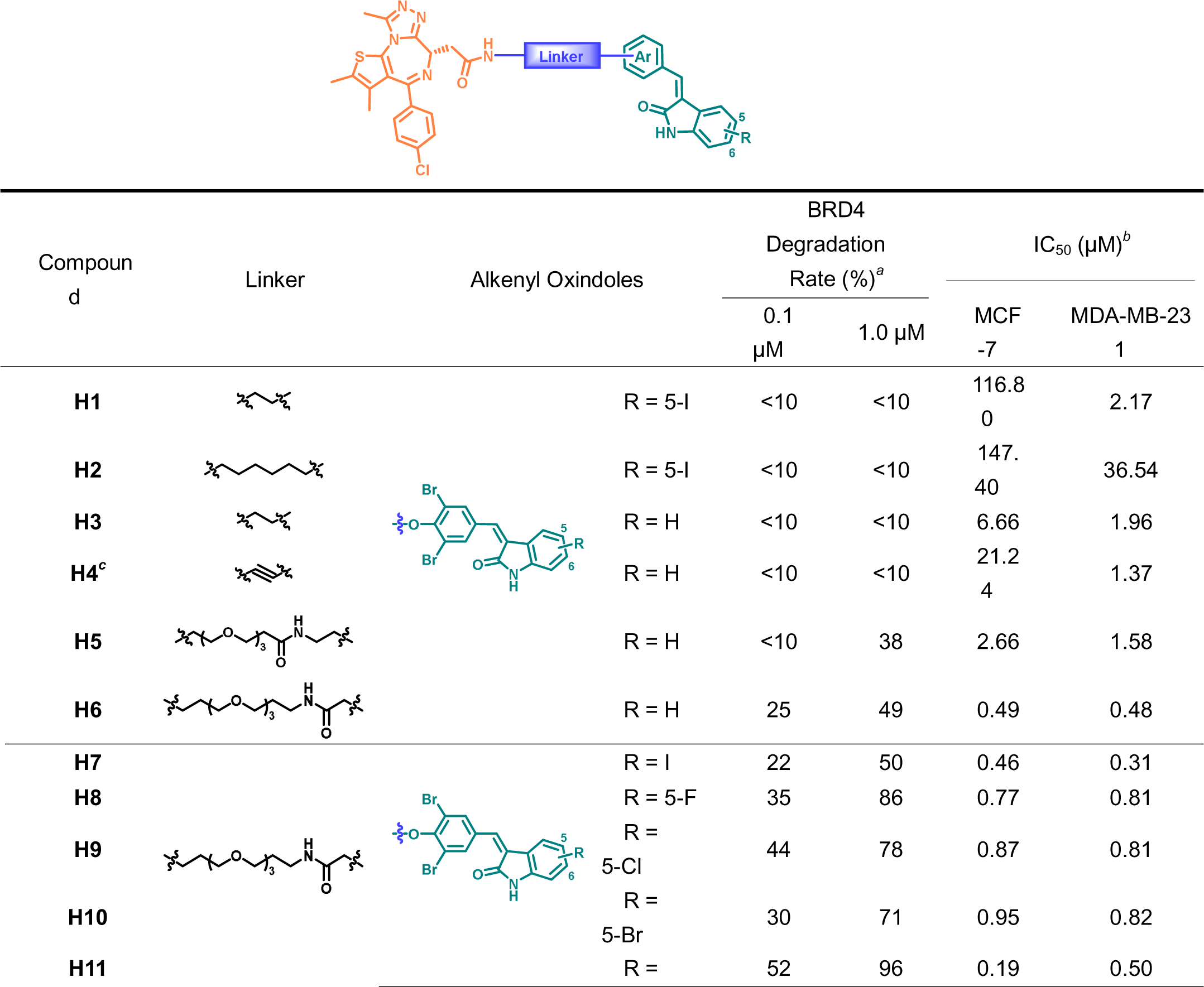
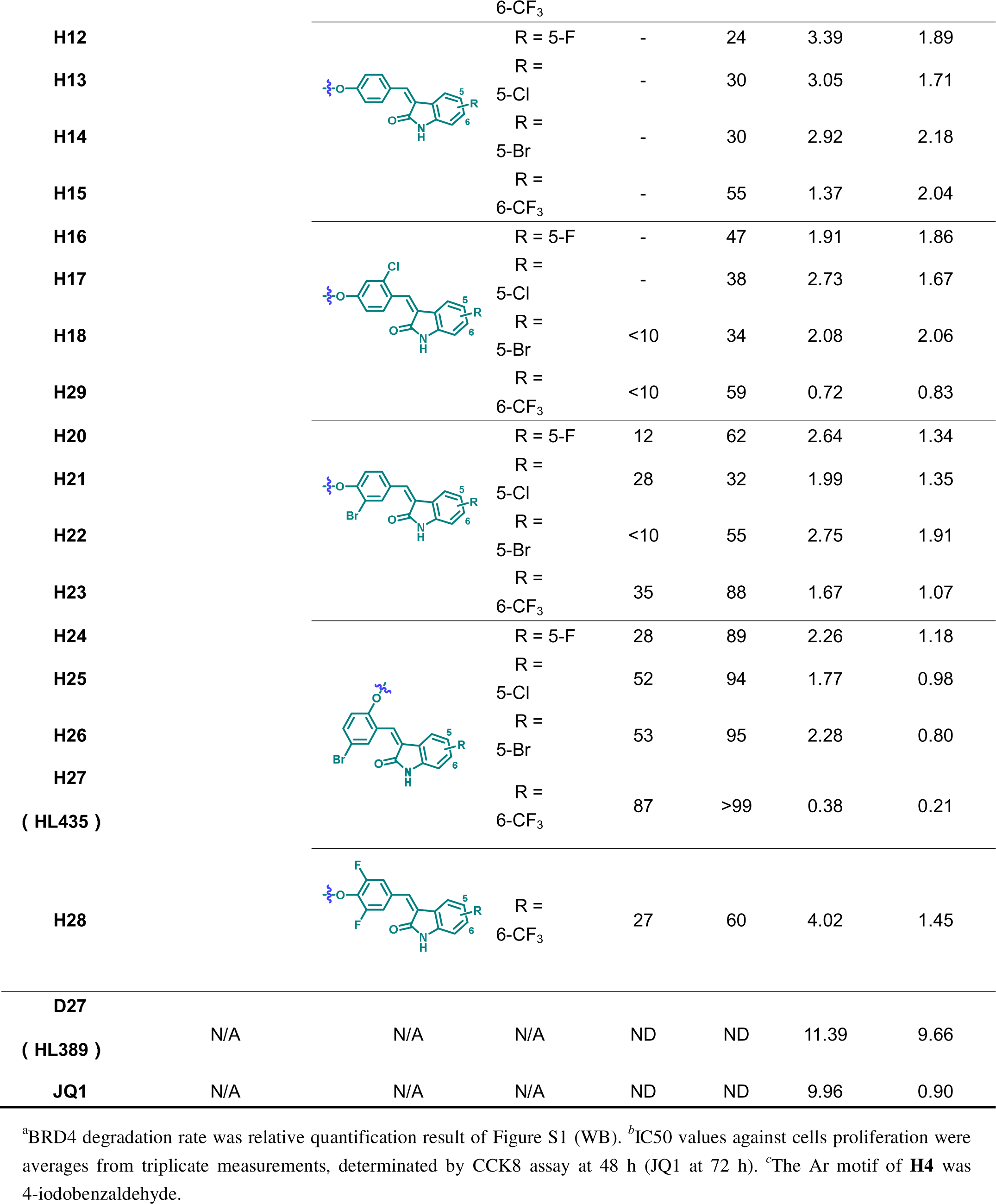
Target Degradation Efficiency and Antiproliferation Activities of Compounds.

### 2.2. HL435 potently degrades BRD4 through the ubiquitin-proteasome pathway

To evaluate the efficacy of HL435 (Figure 1A) in depleting BRD4, we conducted a concentration-dependent study in human breast cancer cells (Figure 1B and S3A). The maximum degradation efficiency (D_max_) of HL435 was > 99%, with DC_50_ values of 11.9 and 21.9 nM in MDA-MB-231 and MCF-7 cells, respectively (Figure 1C). Kinetics study of BRD4 degradation showed that degradation was observed just 1 h after HL435 treatment (Figure 1D and S3B), with a half-life of 1.38 and 1.31 h in MDA-MB-231 and MCF-7 cells (Figure 1E), respectively. Meanwhile, HL435 was demonstrated to efficiently degrade BRD4 in a concentration-dependent manner in multiple cell lines (Figure S2). Although the efficacies varied slightly among different cell lines, the D_max_ all > 99%.

**Figure 1.**
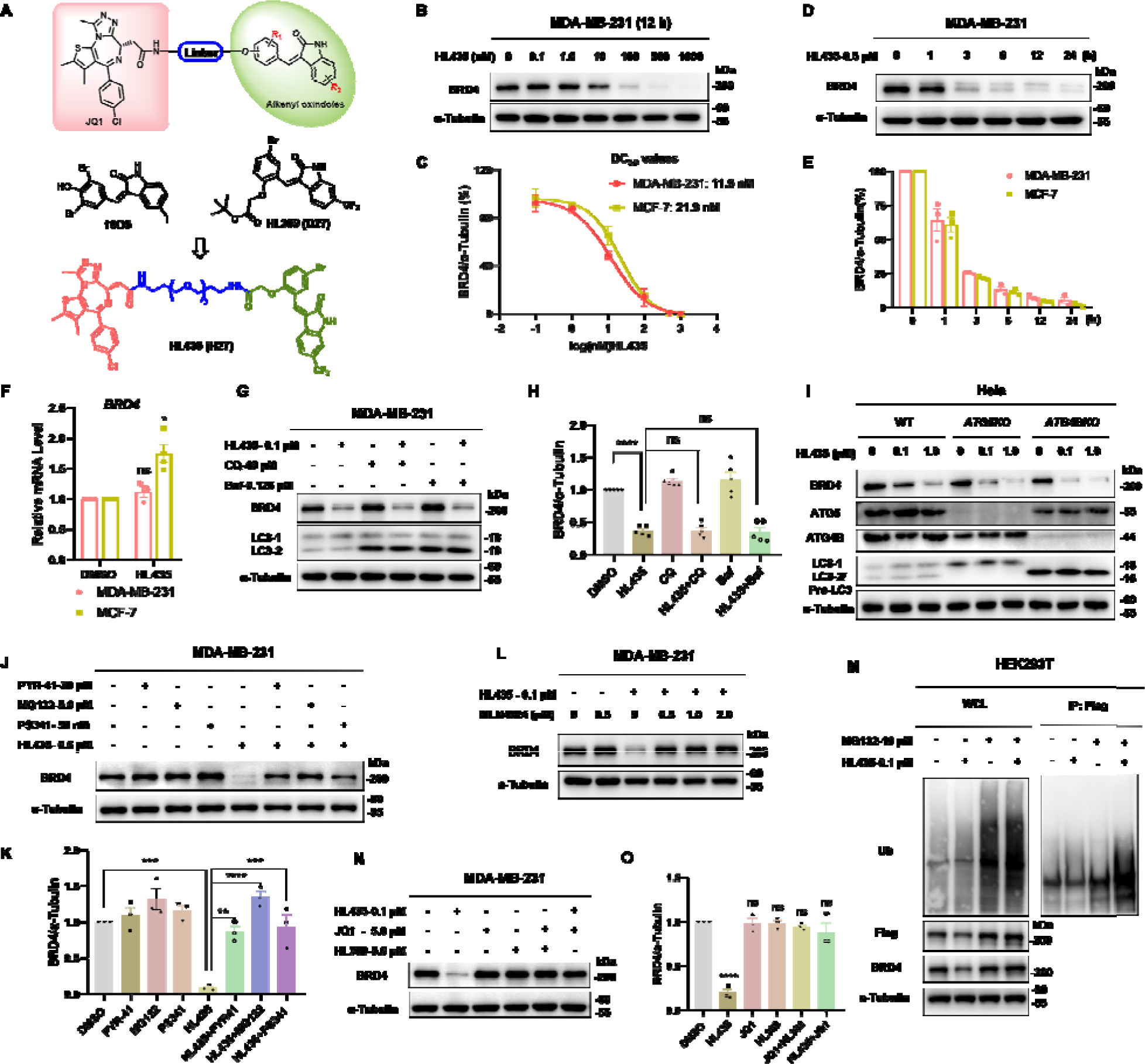
HL435 potently degrades BRD4 through the ubiquitin-proteasome pathway. **A**, Design and development of HL435 as a potent BRD4 degrader. **B**, Representative WB results, Cells were treated with gradient concentrations of HL435 for 12 h. **C**, The DC_50_ values of HL435 to degrade BRD4 in MDA-MB-231 and MCF-7 cells, from relative quantitative analysis. **D**, Representative WB results, MDA-MB-231cells were treated with HL435 at 0.5 μM for gradient time. **E**, Relative quantitative analysis for the time-dependent degradation of BRD4. **F**, The relative mRNA levels of *BRD4*, cells were treated with HL435 at 1.0 μM for 12 h, GAPDH used as control. **G**, Representative WB results, cells were co-treated with HL435 and chloroquine (CQ) or bafilomycin (Baf) for 6 h, CQ or Baf was pre-treated for 2 h. **H**, Relative quantitative analysis of BRD4 from **G**. **I**, Representative WB results (n=3). *WT*-Hela, *ATG5KO*-Hela or *ATG4BKO*-Hela cells were treated with indicated concentration of HL435 for 6 h. **J**, WB results for BRD4 degradation, cells were pre-treated with PYR-41, MG132 or PS-341 for 2 h, followed by HL435 treatment for 6 h. **K**Relative quantitative analysis of BRD4 from **J**. **L**, WB results, HL435 and MLN4924 were co-treated at indicated concentration for 6 h, MLN4924 was pre-treated for 2 h. followed by treatment with MG132 for 8 h and HL435 for 6 h. Cell lysates were Immunoprecipitated with anti-Flag magnetic beads and immunoblotted for ubiquitination level of BRD4. **N**, WB, JQ1, HL389 or HL435 was treated for 6 h. **O**, Relative quantitative analysis BRD4 from **N**. Data were presented as mean ± SEM. Statistical significance was determined by One-way analysis of variance (ANOVA) or Mann Whitney test (**F**). *p < 0.05, **P < 0.01, ***P < 0.001, ****P < 0.0001; ns, no statistical significance.

To validate the mechanism of BRD4 depletion induced by HL435, we first assessed the mRNA levels of *BRD4*. The mRNA level of *BRD4* in the HL435 treated group was not lower than that of control group in breast cancer cells, indicating that HL435 did not affect BRD4 at the transcriptional level but rather at the protein level (Figure 1F). Alkenyl oxindoles were found to bind both LC3 and mHTT for inducing autophagy degradation of mHTT^32^, so we explored whether autophagy-lysosomal inhibitors could rescue the degradation of BRD4 induced by JQ1- alkenyl oxindole-conjugated compounds. Surprisingly, both CQ and bafilomycin failed to rescue the degradation of BRD4 induced by HL435, **H1**, **H6** or **H7** (**10f**) in different cell lines (Figure 1G, 1H, S4B, S4D). We next treated WT-, *ATG5KO*-, and *ATG4BKO*-Hela cells with HL435 and found that the absence of LC3 and autophagosomes did not affect the efficiency of HL435 in degrading BRD4 (Figure 1I). Similar results were obtained for **H1** (Figure S4C). These results suggested that the degradation of BRD4 by JQ1- alkenyl oxindole-conjugated compounds was independent of the autophagy-lysosomal pathway, thus, we suspected that it was perhaps mediated by ubiquitin-proteasome system. To validate this hypothesis, E1 ubiquitin-activating enzyme inhibitor PYR-41 or proteasome inhibitors MG132 or PS-341 was employed to co-treatment with our compounds. As expected, pre-treatment with PYR-41, MG132 and PS-341 all successfully rescued the degradation of BRD4 induced by HL435, **H6** or **H7** (**10f**) (Figure 1J, 1K, S4D, S4E). Pre-treatment with NEDD8 activating E1 enzyme (NAE1) inhibitor MLN4924 also block the degradation of BRD4 by HL435 (Figure 1L), indicating that the degradation required the activation of Cullin-RING E3 ligase (CRL). When the degradation process was blocked by proteasome inhibitor MG132, HL435 increased the ubiquitination level of BRD4 (Figure 1M). Finally, we validated that both JQ1 and structurally modified alkenyl oxindole HL389, whether used alone or in combination, cannot deplete BRD4 in MDA-MB-231 cells. Moreover, an excess of JQ1 could competitively block the degradation of BRD4 induced by HL435 (Figure 1N, 1O). All these results suggested that JQ1-alkenyl oxindole-conjugated compounds including HL435 degrade BRD4 through the ubiquitin-proteasome pathway rather than the autophagy-lysosomal pathway, and the degradation process depends on compounds simultaneously interacting with the substrate protein and the ubiquitin-proteasome system.

### 2.3. A focused CRISPRi screen identified CRL4^DCAF11^ complex potentially responsible for HL435-induced proteasomal degradation activity

To identify the E3 ligase mediating HL435-induced degradation of BRD4, we determined to conduct a pooled CRISPR interference (CRISPRi) screen. We first constructed a dual-fluorescence reporter, containing the BRD4 bromodomain 1 (BD1) fused to mScarlet, followed by a P2A self-cleaving fragment and an enhanced green fluorescent protein (EGFP) for normalization (Figure 2A). This reporter was stably transduced into HEK293T cells constitutively expressing the CRISPRi machinery (dCas9-BFP-KRAB) from the CLYBL safe harbor locus^33^. Upon treatment with HL435, the relative BD1 intensity was significantly reduced, which could be fully restored by MG132 (Figure 2B-D), confirming the sensitivity of the reporter. Next, we designed a focused sgRNA library targeting all known human E1, E2, and E3 enzymes, consisting of 5071 sgRNAs against 993 genes with 5 sgRNAs per gene, and more than 100 non-targeting control sgRNAs. Using this library, we performed a fluorescence activated cell sorting (FACS)-based CRISPRi screen in the BD1 reporter cells based on relative BD1-mScarlet signal (BD1-mScarlet intensity normalized to EGFP intensity). In the HL435-treated group, knockdown of any components mediating HL435-induced degradation activity would result in an increased relative BD1-mScarlet signal (Figure 2E), which would not increase in the DMSO-treated group. As shown in Figure 2F and 2H, components of the CRL4^DCAF11^ complex, including the E3 ligase scaffold Cullin-4B (*CUL4B*), the RING-finger protein RING-box1 (*RBX1*), the adaptor Damage-specific DNA binding protein 1(*DDB1*) and the substrate receptor DDB1 and CUL4 associated factor 11 (*DCAF11*) were among the top positive hits, whose knockdown increased BD1-mScarlet signal in the HL435-treated group. In addition, *NAE1* and ubiquitin like modifier activating enzyme 3 (*UBA3*), which are responsible for CRL neddylation and activation, were also strong hits in the screen (Figure 2F). Importantly, these genes showed no phenotype or only weak phenotype in the DMSO group. These data indicated that the CRL4^DCAF11^ complex is specifically involved in HL435-induced substrate degradation (Figure 2F, G and Supplementary Table 1).

**Figure 2.**
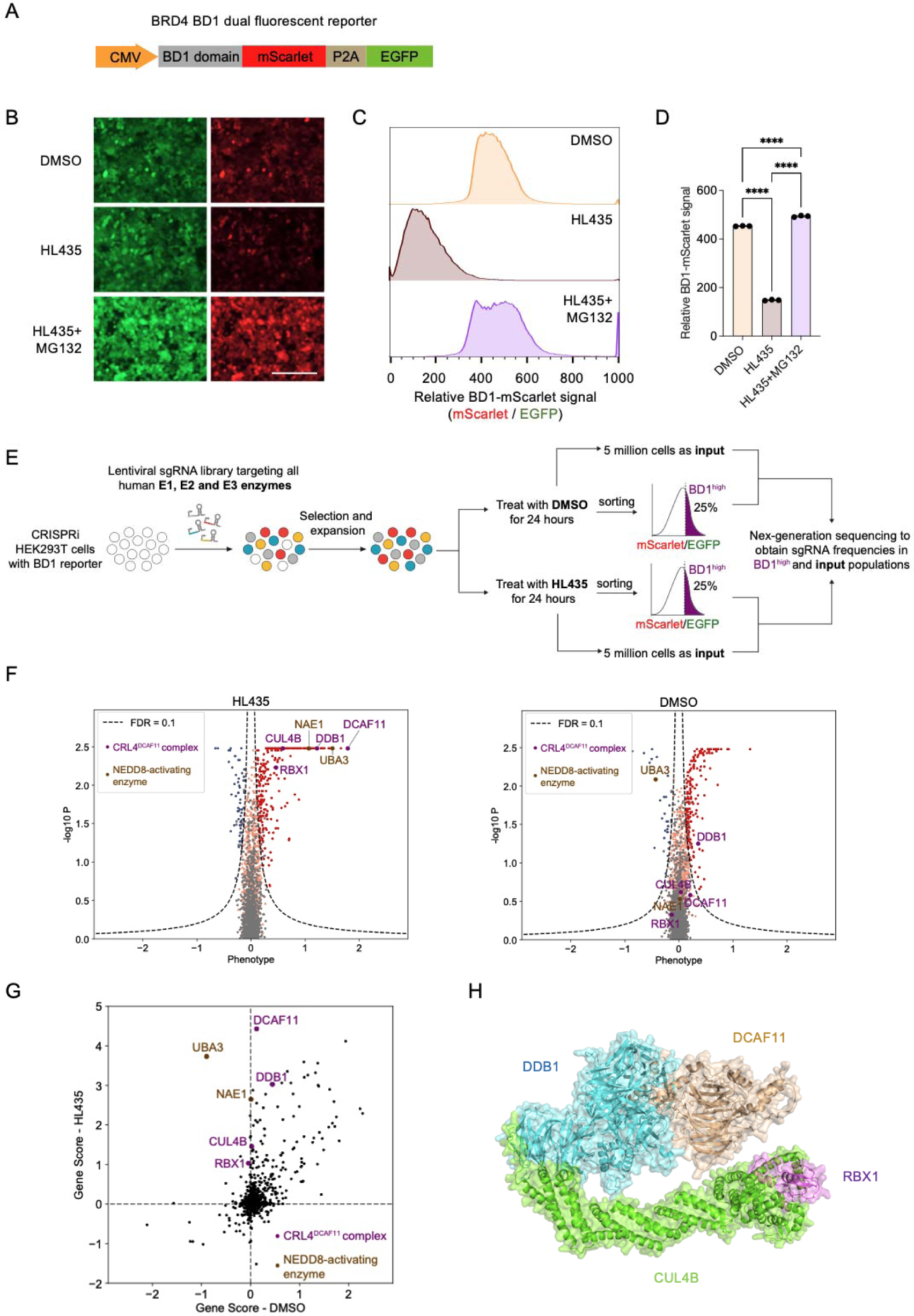
CRISPRi screen identified the CRL4^DCAF11^ complex as a potential target mediating HL435-induced proteasomal degradation of BRD4. **A,** Design of BRD4 BD1 dual fluorescent reporter. BRD4 BD1 domain was fused with mScarlet, followed by a P2A self-cleaving fragment and an EGFP. **B,** Validation of BD1 reporter response to HL435 treatment. Representative fluorescent microscope fields for BD1 reporter levels in HEK29T cells treated with DMSO, HL435 (50 nM), treatment. Relative BD1-mScarlet signal in HEK293T cells was determined by the ratio of mScarlet and EGFP as measured by flow cytometry. The cells were treated with DMSO, HL435 (50 nM), or HL435 (50 nM) and MG132 (5 μM) for 24 hours. **D,** Quantification of the relative BD1-mScarlet signal in the indicated groups was shown in the bar graph (mean ± s.d., n = 3 biological replicates). **E,** CRISPRi screen strategy. CRISPRi HEK293T cells harboring BD1 reporter were transduced with an sgRNA library targeting all human E1, E2, and E3 enzymes. Cells were treated with DMSO or HL435 (50 nM, 24 hours) and 5 million cells were taken as “input”. Cells with top 25% relative BD1-mScarlet signal (BD1^high^) were sorted via FACS. The frequencies of BD1^high^ and input cells expressing each sgRNA were determined by next-generation sequencing, and were compared to determine sgRNAs enriched or depleted in the BD1^high^ population. The screens were performed in duplicates. **F,** Screening results analyzed by the MAGeCK-iNC pipeline were shown for HL435-treated and DMSO-treated groups. A positive phenotype indicates the corresponding sgRNA was enriched in the BD1^high^ population and vice versa. Dots in red, blue, grey and orange represent positive hits, negative hits, negative control and other genes, respectively. Genes encoding components of the CRL4^DCAF11^ complex and the NEDD8-activating enzyme were highlighted. **G,** Scatter plot comparing gene scores for the screens under HL435 and DMSO treatment. Genes encoding components of the CRL4^DCAF11^ complex and the NEDD8-activating enzyme were highlighted. **H,** Predicted structure of CRL4^DCAF11^ complex using Alphafold. Statistical significance was determined by One-way ANOVA. ****P < 0.0001.

To validate the requirement of CRL4^DCAF11^ complex for the degradation activity of HL435, we individually cloned two separate sgRNAs targeting each of the following genes: *DCAF11*, *DDB1*, *CUL4B*, *RBX1, NAE1* and *UBA3,* followed by evaluating their effects on HL435-induced BD1 reporter degradation (Figure 3A). As expected, knocking down any of these genes blocked the BD1 reporter degradation upon HL435 treatment (Figure 3B-D). Additionally, we confirmed the requirement of DCAF11 in HL435-induced degradation of endogenous BRD4 (Figure 3E and F). Furthermore, we detected interaction between BRD4 BD1 and DCAF11 only in the presence of HL435, suggesting the formation of a ternary complex between BRD4, HL435 and DCAF11(Figure 3G). Taken together, these data indicate that the CRL4^DCAF11^ complex is responsible for HL435-induced proteasomal degradation of BRD4.

**Figure 3.**
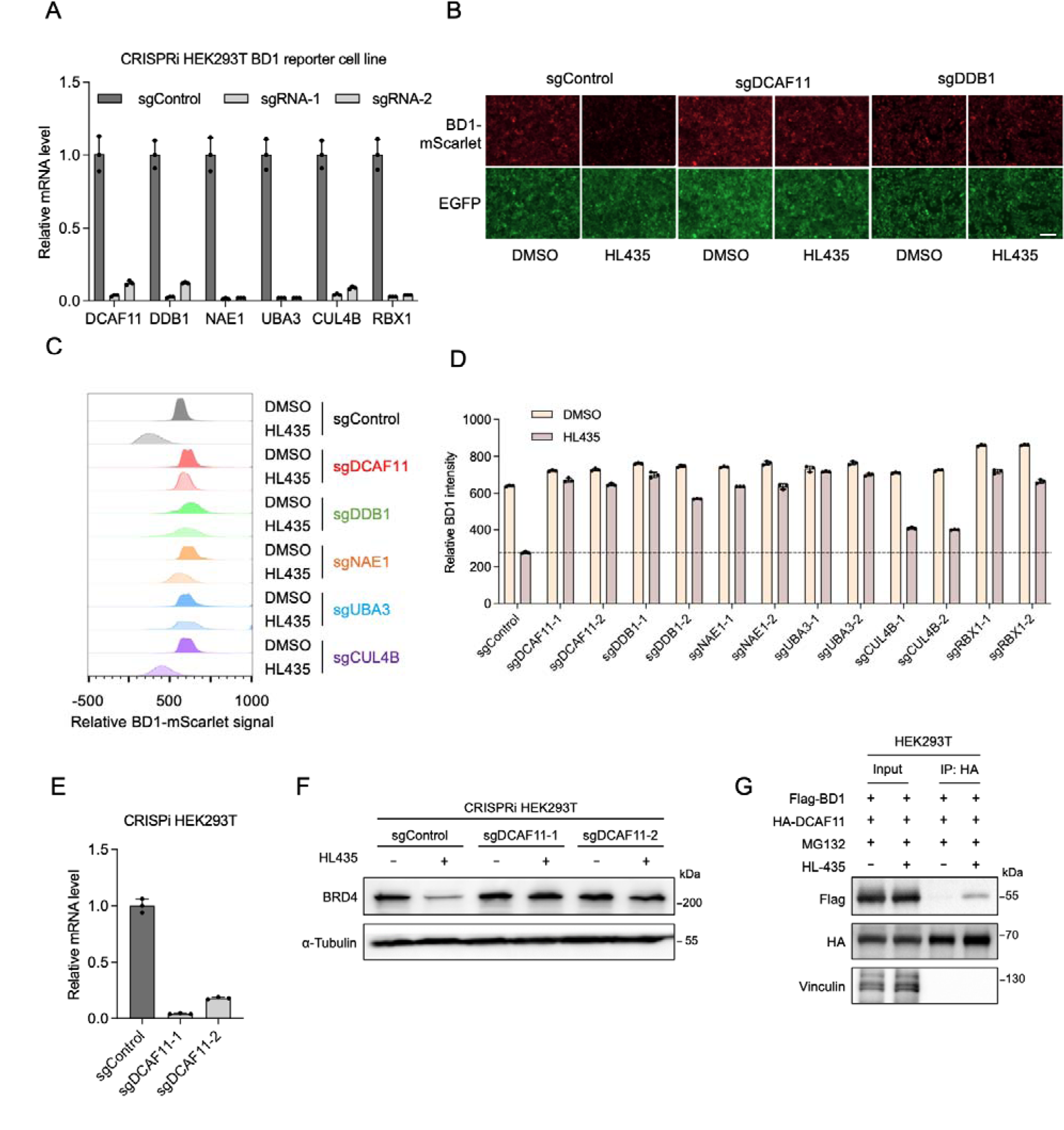
HL435 recruits CRL4^DCAF11^ complex to induce proteasomal degradation of BRD4. **A**, Validation of knockdown efficiency of different hit genes in CRISPRi HEK293T reporter cells by RT-qPCR (mean ± s.d., n = 3 technical replicates). **B**, Representative fluorescent microscope fields for BD1 reporter levels in the reporter cells expressing sgRNAs targeting individual hit genes after DMSO or HL435 treatment (50 nM, 24 hours). Bar = 200 μm. **C**, Relative BD1-mScarlet signal was quantified by flow cytometry for hit gene knockdown in BD1 reporter cells after DMSO or HL435 treatment (50 nM, 24 hours). **D**, Quantification of the relative BD1 intensity in the indicated groups was shown in the bar graph (mean ± s.d., n = 3 biological replicates). **E**, Validation of knockdown efficiency of two sgRNAs targeting *DCAF11* in CRISPRi HEK293T reporter cells by RT-qPCR (mean ± s.d., n = 3 technical replicates). **F**, Western blot showing endogenous protein levels of (50 nM, 24 hours). **G**, Co-immunoprecipitation analysis of HA-DCAF11 with Flag-BD1 in the absence or presence of HL435 (5 μM, 6 hours) in HEK293T cells.

### 2.4. HL435 potently inhibits proliferation and induced apoptosis of tumor cells *in vitro*

BRD4 has been identified as a potential therapeutic target for tumors owing to its contribution to tumor pathogenesis^34, 35^. As shown in Table 1, most of JQ1- alkenyl oxindole-conjugated compounds exhibited excellent anti-proliferation abilities in breast cancer cell lines MCF-7 and MDA-MB-231, with HL435 performed the best overall. The anti-proliferation activities of different compounds were positively correlated with their BRD4 degradation abilities, suggesting that BRD4 degradation contributed substantially to the anti-proliferation activity of breast cancer. Notably, the half-maximal inhibitory concentrations (IC_50_) of HL435 against 22RV1 (prostate cancer) was as low as 8.7 nM (Figure 4A), significantly superior to JQ1 (IC_50_ =157 nM). To further explore the biological effects of HL435 on -breast cancer, we performed flow cytometric analysis and Western blotting to assess the influence on cell cycle and apoptosis in MCF-7 and MDA-MB-231 cells. HL435 acted similarly to JQ1 at low concentration, blocking the cell cycle at G0/G1 phase, while higher concentrations of HL435 arrested cell cycle at G2/M phase (Figure 4B, 4C, S5A and S5B). Consistent with the results of flow cytometric analysis, immunoblotting results showed that HL435 treatment up-regulated P53 and P21 levels and down-regulated the levels of Cyclin D1 and Cyclin B1 (Figure 4E, S5C), which contributed to block the G1/S and G2/M transitions. The ability of apoptosis induction by HL435 in breast cancer cells was > 20-fold more potent than JQ1 (Figure 4D, S6A). Treatment with HL435 at 1.0µM for 36 h in MDA-MB-231 cells led to an apoptotic rate of 55.9 ± 1.9%, and the levels of cleaved caspase-9 and PARP1 were profoundly increased accordingly (Figure 4F, S6B). Oncogenes c-Myc is the key downstream signal used to evaluate the function of BRD4^36^. As displayed in Figure 4E, S5C and S7A, both mRNA and protein expression levels of c-Myc were significantly downregulated in breast cancer cells treated with HL435. These data indicated that HL435 not only exhibits excellent anti-proliferative capacity against multiple tumor cell lines, but also effectively arrests the cell cycle and induces apoptosis in breast cancer cells, validating the stronger therapeutic efficacy of degraders compared to inhibitors.

**Figure 4.**
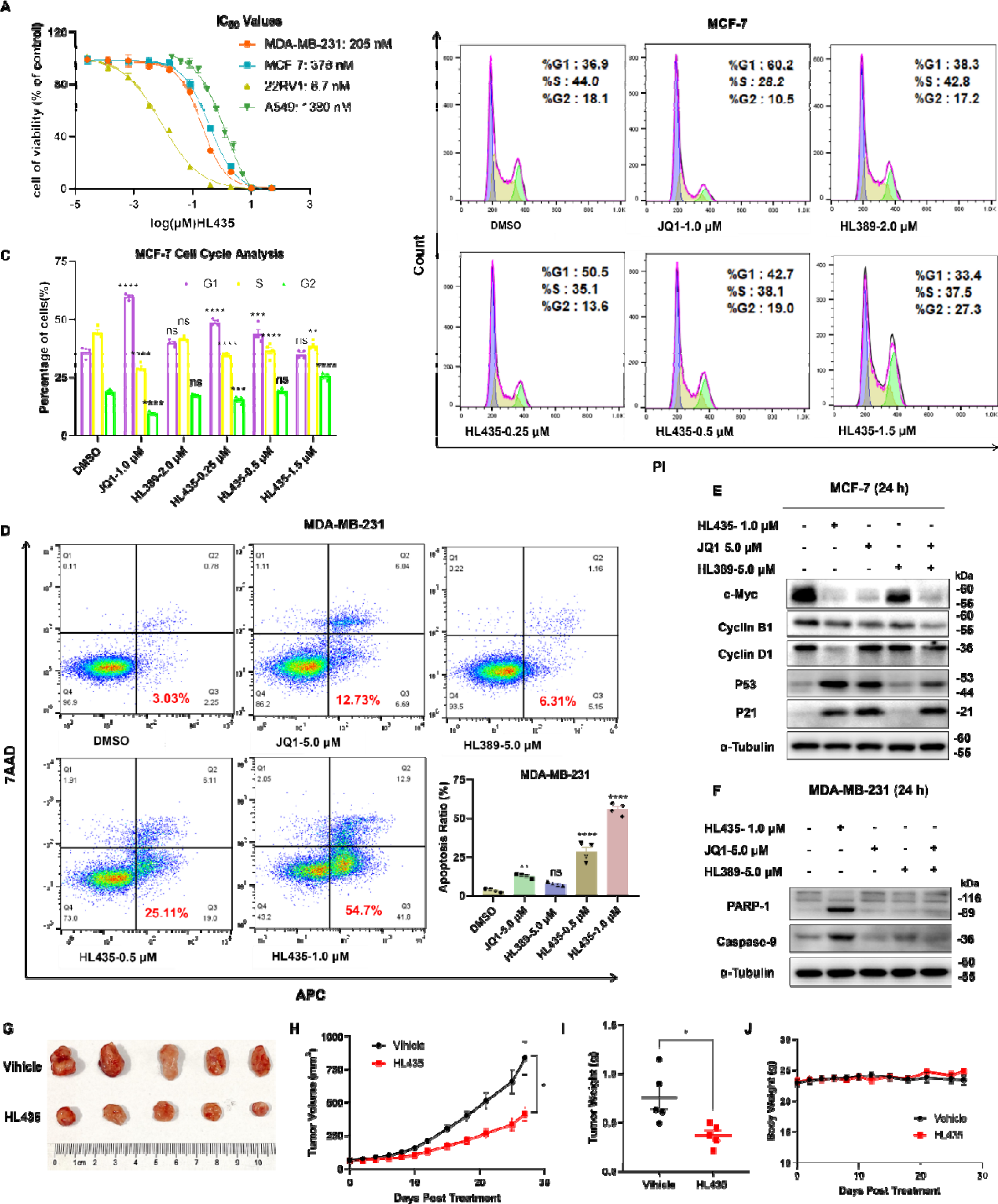
HL435 exhibited excellent anti-tumor efficacy in vitro and in vivo. **A**, IC_50_ values of HL435 against multiple tumor cell lines, determinated by the CCK8 assay after 48 h treatment. **B**, Representative flow cytometry analysis results of analysis of cell cycle for **B**. **D**, Representative flow cytometry analysis results and quantitative statistical analysis of apoptosis, MDA-MB-231 cells were treated with indicated compounds for 36 h before stained with an 7AAD/APC Apoptosis Detection kit. **E**, Representative Western blotting results of c-Myc and cycle relevant proteins in MCF-7 cells (n=3). **F**, Representative Western blotting results of apoptosis relevant proteins in MDA-MB-231 cells (n=3). **G**, Picture of stripped xenograft tumors at the end of experiment (day 45). NOD-SICD mice bearing the MDA-MB-231 xenograft were daily administered with vehicle (10% DMSO+90% corn oil, i.p.) or HL435 (20 mg/kg, i.p.) 6 days per week for 27 days. **H**, Growth curve of xenograft tumors after treatment. **I**, weights of stripped xenograft tumors at day 45. **J**, Body weight curves during treatment. Data was presented as mean ± SEM (n≥3). One-way ANOVA or Unpaired t test was employed to determine statistical significance. **P < 0.01, ***P < 0.001, ****P < 0.0001; ns, no statistical significance.

### 2.5. HL435 suppresses tumor growth *in vivo*

To evaluate the anti-tumor ability of HL435 *in vivo*, a mouse xenograft tumor model of MDA-MB-231 cells was employed. Mice bearing xenograft tumor were treated daily with HL435 (20 mg/kg) or vehicle (10% DMSO+90% corn oil) 6 days per week. After 27 days of treatment, the HL435 treatment group attenuated tumor progression, with a tumor growth inhibition rate (TGI) of 54.34% (Figure 4H) and a 51.12% reduction in tumor weight (Figure 4I) compared to vehicle group. Meanwhile, the body weight of HL435-treated group was comparable to that of the vehicle group, and no obvious toxicity or adverse effects were observed throughout the experiment period (Figure 4J), indicating that HL435 was tolerated well. These data further validated the anti-tumor efficacy of HL435 *in vivo*, providing a promising drug-like lead compound for anticancer drug development.

## 3. Discussion and Conclusions

Alkenyl oxindoles have been characterized as molecular glues that tether mHTT to LC3, enabling the lysosomal degradation of mHTT^32^. We initially sought to expand the versatility of this approach by conjugating alkenyl oxindoles to other substrate binding moieties, generating bifunctional molecules that may facilitate the degradation of various substrate proteins through the autophagy-lysosomal pathway. As a proof-of-principle, we generated a series of JQ1-alkenyl oxindole conjugates, from which we indeed identified molecules that potently degrade the target BRD4. However, we discovered that BRD4 degradation induced by JQ1-alkenyl oxindole conjugates does not occur via the autophagy-lysosomal pathway, but through the ubiquitin-proteasome pathway. We speculated that the oxidized indole structure may recruit E3 ubiquitin ligase for its degradation activity. To determine the responsible E3 ubiquitin ligase, we conducted a pooled CRISPRi screen, from which we identified the CRL4^DCAF11^ complex as a potential target for mediating the degradation activity of JQ1-alkenyl oxindole conjugates. We showed that alkenyl oxindoles can recruit DCAF11, thus acting as a novel PROTAC moiety for targeted protein degradation. Previously, the Cravatt^37^ and Gray^38^ groups respectively revealed that DCAF11 serves as an E3 ligase that can support protein degradation triggered by electrophilic PROTACs. Very recently, while we were preparing this manuscript for reviewing, similar findings were reported by Waldmann and Winter et al^39^. As alkenyl oxindole possesses Michael acceptor properties, they confirmed that it recruits DCAF11 through a covalent modification approach, potentially engaging with cysteine residues.

Previous structure-activity studies on PROTACs have mainly focused on the effects of different linker lengths on degradation activity, with few studies on the structure-activity of the E3 ligase ligand part. In this study, we found that the ability to degrade BRD4 were significantly improved through structural optimization of E3 ligase ligands, and an excellent BRD4 degrader HL435 was identified, whose JQ1 moiety was conjugated with the trifluoromethyl-substituted alkenyl oxindole via PEG chain. The D_max_ of HL435 > 99%, with DC_50_ values of 11.9 nM in MDA-MB-231 cells. To explore the druggability potential of hetero-bifunctional compounds conjugated with alkenyl oxindoles, we evaluated their anti-tumor abilities both *in vitro* and *in vivo*. Most of hetero-bifunctional compounds exhibited more excellent anti-proliferation abilities against breast cancer cells than JQ1 or alkenyl oxindoles, supporting the more excellent therapeutic potential of degraders compared to inhibitors. Consistent with the degradation efficiency of BRD4, HL435 showed the best anti-proliferative activity overall, with an IC_50_ as low as 8.7 nM against prostate cancer cells 22RV1. In addition to outstanding antiproliferative abilities against multiple tumor cells, HL435 can effectively arrest the cell cycle and induce apoptosis in breast cancer cells by blocking BRD4 downstream signaling pathway in a concentration-dependent manner. Finally, the anti-tumor efficacy of HL435 *in vivo* was validated in a mouse TNBC xenograft model, with good tolerability. These data suggested that HL435, a compound composed of structure modified alkenyl oxindole and BRD4 inhibitor JQ1, was a promising drug-like lead compound for anticancer drug development.

Although there are more than 600 E3 ubiquitin ligases in human cells, the ligand molecules currently available to recruit E3 ubiquitin ligases only cover less than 3%^18, 19^. In addition, with the emergence of E3 ubiquitin ligase resistance, PROTACs based on the same E3 ubiquitin ligase ligand may be ineffective^40–42^. Therefore, the development of new E3 ubiquitin ligase ligands can not only solve the limitations of existing ligands, but also be a major way to expand the scope of PROTACs therapeutic targets and provide better treatment opportunities^26^. Moreover, as DCAF11 is localized in the nucleus, the identification of ligands capable of recruiting DCAF11 provides new possibilities for targeting the degradation of nuclear proteins.

In summary, we discovered alkenyl oxindole as a novel PROTAC moiety for targeted protein degradation via CRL4^DCAF11^ recruitment. We also developed JQ1-alkenyl oxindole-conjugated bifunctional molecules with high BRD4 degradation efficiencies in multiple cell lines and proved their anti-cancer effect both *in vitro* and *in vivo*. Our study expands the E3 toolbox available for PROTACs, which will potentially broaden the spectrum of degradable proteins and improve the efficiency of target degradation, providing new possibilities for drug discovery.

## 4. Experimental Section

### 4.1. Cell lines culture and plasmid transfection

HCT116, MCF-7, 22RV1, A549, K562, THP-1, Hela and HEK293T cell lines were previously obtained from ATCC and cryopreserved in our laboratory. MDA-MB-231 was newly purchased from Procell (Wuhan, China). *ATG4B*KO-Hela, *ATG5*KO-Hela, *ATG4B*KO-HCT116 were kindly gifts from Professor Li (Sun Yat-sen University). The CRISPRi HEK293T cell line was established by integrating dCas9-BFP-KRAB cassette into the CLYBL safe harbor locus via homology-directed repair (HDR) as described previously^33^.

All cell lines were mycoplasma-free. HA-Ub and Flag-BRD4 plasmids were purchased from Miaoling Biology (Wuhan, China). Cell lines were cultured in RPMI-1640 medium (Gibco, Thermo Fisher Scientific, Waltham, MA, USA) or DMEM (Gibco) supplemented with 1% penicillin-streptomycin (Gibco) and 10% fetal bovine serum (FBS, sigma), the cultures were maintained in a CO_2_ incubator at 37 °C with 5% (v/v) CO_2_. For transient transfection, HEK293T cells were seeded into 6-well plates and cultured to about 50% density, then co-transfected with HA-Ub and Flag-BRD4 plasmids for 32 h by Hieff Trans^TM^ Liposomal Transfection Reagent (Yeasen, China).

### 4.2. DNA constructs

The coding sequence of BRD4 BD1 domain (amino acid N44-E168) was amplified from full-length of BRD4 cDNA in a pcDNA3.1-BRD4-3Flag with the forward primer (5’-ATGACGATGACAAGACTAGTaaccccccgcccccagagacctcca-3’) and reverse primer (5’-ccgccttcttctgtgggtagctcatttatt-3’) and it was fused with mScarlet-P2A-EGFP from pRT117 vector into the pLVX-3Flag-Hygro vector with the forward primer (5’-ctacccacagaagaaGGCGGTGGCTCGGTGAGCAA-3’) and reverse primer (5’-AGGGGCGGGATCCGCGGCCGCttactagtcggttcaactctaggtg-3’) by Hieff Clone^®^ Universal One Step Cloning Kit (YEASEN, 10922ES20). The coding sequence of DCAF11 (Youbio, L11006) was inserted into a pcDNA3.1-3HA vector with the forward primer (5’- TACCTGACTACGCTGGTACCatgggatcgcggaacagcagcag-3’) and reverse primer (5’- GATATCTGCAGAATTCctactggggtgaggaaaaggg-3’) by Hieff Clone^®^ Universal One Step Cloning Kit (YEASEN, 10922ES20).

### 4.3. CRISPRi screen

The overall CRIPSRi screen process was performed as described previously^33, 43, 44^. In brief, HEK293T cells harboring the BRD4 BD1 domain dual fluorescence reporter were infected with the sgRNA library targeting all human E1, E2, and E3 enzymes as described previously. The MOI value was controlled under 0.3 when library was transduced to the cells. The cells were selected by 2 μg/mL puromycin for 2 days. After expansion and puromycin selection, the cells were treated with DMSO and HL435 respectively. 24 hours later, 5 million cells were taken as “input”, and the remaining cells were subsequently collected for FACS, where the cells were sorted into the top 25% based on the ratio of mScarlet and EGFP signal. For each sample, cells corresponding to at least 4,000-fold over the library coverage were sorted per replicate. Sorted populations were collected and genomic DNA was isolated using DNAiso Reagent (Takara, 9770A). sgRNA cassettes were amplified by PCR and subjected to Next-generation sequencing (NGS) by NovaSeq 6000 PE150. Sequencing results were analyzed using MAGeCK-iNC as previously described^33^.

### 4.4. Cell viability assay

Cells were seeded in 96-well plates at a density of 2000-4000 cells per well. After overnight incubation, compounds were administered at the gradient concentrations for 2-3 days. Then, remove the old medium and add 100 μL fresh medium with 10% CCK8 reagent (Bimake; Selleck Chemicals; cat. no. B34304) for each well. And the plate was incubated in a cell incubator at 37 °C for 1-3 h. The optical density (OD) value at 450 nm, which stands for the vitality of the cells, was detected with a BioTek Synergy H1 microplate reader. IC_50_ values of compounds against cell lines were calculated from triplicate measurements.

### 4.5. Co-Immunoprecipitation and Western blot analysis

Cells were lysed in RIPA buffer (Beyotime, Haimen, China) supplemented with phosphatase inhibitors (Bimake; Selleck Chemicals, Houston, TX, USA) or protease inhibitor cocktail (Roche, Basel, Switzerland). The protein in each sample was quantified by a BCA protein assay (ThermoFisher, Rockford, I L), and boiled with 5×loading buffer (LB) for 5 min. For immunoprecipitation, 1 mg protein in each sample was incubated with anti-Flag (P2115, Beyotime, China) or anti-HA magnetic beads (P2121, Beyotime, China) overnight at 4°C. After washed away non-specifically bound proteins, anti-Flag magnetic beads were boiled with 1× LB for transsexual washout. 8%, 10% and 12% SDS-PAGE gels or 3% Tris-Acetate Polyacrylamide Gradient Gels were used to separate protein samples, and then transfered to a PVDF membrane (Millipore; Merck KGaA). 5% skim milk was used to block membranes at RT for 1 hour. Primary antibodies were blotted at 4°C overnight. The next day, the membranes were slowly flipped in secondary antibodies conjugated with horseradish peroxidase for 1 hour at RT. Images were captured by Tanon 5200 (Shanghai, China). Image J was used to quantify the intensities of bands. The antibodies used in this paper were as below: Anti-[-Tubulin (T6047), anti-LC3B (L7543) and anti-β-Actin were purchased from Sigma (St. Louis, MO, USA); Anti-BRD4 (13440), anti-PARP1 (9542), anti-Caspase 9 (9505), anti-HA (3724) and anti-Cyclin D1(2922) were purchased from Cell Signaling Technology (Danvers, MA); Anti-Ubiquitin (sc-8017) was from Santa Cruz (Dallas, TX, USA). Anti-ATG4B (M134), anti-ATG5 (M153) and anti-Flag were from MBL (Tokyo, Japan); Anti-Cyclin B1(55004-1-AP), anti-p53 (10442-1-AP), anti-p21 (10355-1-AP), anti-c-Myc (10828-1-AP), and anti-GAPDH (60004-1-Ig), anti-Vinculin (66305-1- Ig), Goat anti-Mouse IgG (H+L) (SA00001-1) and Goat anti-Rabbit IgG (H+L) (SA00001-2) were purchased from Proteintech.

### 4.6. Quantitative real-time polymerase chain reaction (qRT-PCR) analysis

MDA-MB-231 or MCF-7 cells were plated in 12-well plates and treated with compounds for 12 h after overnight incubation. Total RNA was extracted using TRIzol (Invitrogen). A High-Capacity cDNA Reverse Transcription kit (Thermo Fisher Scientific) was used to create cDNA from purified RNA. The real-time PCR was conducted on a real-time fluorescence quantitative PCR equipment (light-Cycler480II, Roche) according to the protocol of SYBR Green qPCR Mix (Dongsheng Biotech, China). Results analyses were performed from three or four biological replicates, each data in biological replicate was triplicate. The expression level of genes was calculated with the 2^-ΔΔCt^ technique, and GAPDH was used as an internal reference. The expression level of each gene in the DMSO group was normalized to 1, and that of treatment groups were presented as fold-change relative to the DMSO group.

The primer sequences used were as follows:

GAPDH-F: GAGTCAACGGATTTGGTCGT, GAPDH-R: GACAAGCTTCCCGTTCTCAG; BRD4-F: CTCCGCAGACATGCTAGTGA, BRD4-R: GTAGGATGACTGGGCCTCTG; c-MYC-F: CACCGAGTCGTAGTCGAGGT, c-MYC-R: GCTGCTTAGACGCTGGATTT; P21-F: TGTCCGTCAGAACCCATGC, P21-R: AAAGTCGAAGTTCCATCGCTC; DCAF11-F: CAATGATCTGGGCTTCACTGAT, DCAF11-R: TCTTGGCAAGCAGACATGAAT; DDB1-F: ATGTCGTACAACTACGTGGTAAC, DDB1-R: CGAAGTAAAGTGTCCGGTCAC; NAE1-F: ACCTGTTCGAGGCACAATTCC, NAE1-R: TCTTTGCTTTTTCACGGTAAACG; UBA3-F: CGATCTGGACCCTTCACACAC, UBA3-R: GCCAGCTCCAATGACTAGAAC; CUL4B-F: ACTCCTCCTTTACAACCCAGG, CUL4B-R: TCTTCGCATCAAACCCTACAAAC; RBX1-F: TTGTGGTTGATAACTGTGCCAT, RBX1-R: GACGCCTGGTTAGCTTGACAT; Tubulin-F: ACCTTAACCGCCTTATTAGCCA, Tubulin-R: ACATTCAGGGCTCCATCAAATC.

### 4.7. Cell cycle assay

The influence of compounds on the cell cycle was detected by cell flow cytometry following instructions of the Cell Cycle Analysis Kit (C1052, Beyotime). In brief, seed cells into a 6-well plate at an appropriate density. After overnight incubation, compounds were administered at the respective concentration for 24 h. Pre-cooled PBS was used to wash cells before and after centrifugation, followed by overnight fixation in 70% ethanol at 4 °C. On the next day, ethanol was removed by centrifugation, then cells were dealt with RNase for 30 min at 37 °C. Subsequently, they were stained with propidium iodide (PI) at room temperature for an additional 30 min. If stored at 4 °C, the stained cells can be detected on a flow cytometer (BD FACSCalibur, BD Biosciences, USA) within 24 h and analyzed for cell cycle distribution using FlowJo software.

### 4.8. Cell apoptosis assay

The effect of compounds on inducing apoptosis was detected using cell flow cytometry according to the instructions of Annexin V APC/7-AAD apoptosis kit (AP105-100, liankebio, China). In brief, seed cells into a 6-well plate at an appropriate density. On the next day, compounds were administered at the respective concentrations for 36 hours. 500 μL 1X binding buffer was used to resuspend the harvested cells, and then add Annexin V-APC (5 μL) and 7-AAD (10 μL). Gently mix the solution and incubate it in the dark at RT for 5 minutes. Finally, a flow cytometer (BD FACSCalibur, BD Biosciences, USA) was employed to detect the cells as soon as possible.

### 4.9. Animal experiments

Animal experiments were performed at the Experimental Animal Center of Sun Yat-sen University (East Campus), and female NOD-SCID mice were purchased from Guangdong Yaokang Biotechnology Co., LTD. Inject subcutaneously 5 million MDA-MB-231 cells on the right dorsal side of each mouse at the age of 6-7 weeks. When tumors size reached 60-70 mm^3^ about half a month later, mice were randomly divided into 2 groups. Mice were daily injected with vehicle (10% DMSO+90% corn oil, i.p.) or HL435 (20 mg/kg, i.p.) 6 days per week for 27 days. Volume of xenograft tumor and body weight of each mouse were measured every 2-4 days. Volume of xenograft tumor = length x width^2^/2. sacrifice all of mice at day 27 post treatment and xenograft tumors were excised for weight measurement. TGI (%) = [1 - (TV_Treatment/Dx_ - TV_Treatment/D1_)/ (TV_Vehicle/Dx_– TV_Vehicle/D1_)] × 100%, X = days post treament. The animal experiments were conducted strictly according to animal ethics guidelines and the protocol (No. SYSU-IACUC-2023-000327), approved by the Institutional Animal Care and Use Committee (IACUC) of Sun Yat-sen University Cancer Center.

### 4.10. Chemistry

Unless otherwise stated, all solvents and the compounds without provided synthesis routes were commercially purchased. All solvents were purified and dried according to standard methods before use. The spectra of ^1^H nuclear magnetic resonance (NMR) was recorded on a Varian instrument (500 MHz or 400 MHz), and the tetramethylsilane signal or residual protio solvent signals was used as the internal standard. ^13^C NMR was recorded on a Varian instrument (125 MHz or 100 MHz). Data for ^1^H NMR were recorded as follows: chemical shift (δ, ppm), multiplicity (s = singlet, d = doublet, t = triplet, m = multiplet, q = quartet or unresolved, coupling constant (s) in Hz, integration). Data for ^13^C NMR were reported in terms of chemical shift (δ, ppm). The progress of the reaction was monitored by thin-layer chromatography (TLC) on glass plates coated with a fluorescent indicator (GF254). Flash column chromatography was performed on silica gel (200-300 mesh). The ESI ionization sources were employed to obtain high resolution mass spectra (HRMS). The purity of final key products was confirmed by a Waters e2695 HPLC system equipped with an XBridge C18 (5 um, 4.6 x 250 mm) and eluted with methanol/water (97.5: 2.5) at a flow rate of 1.0 mL/min. The yields indicated were from single step reactions. All compounds used in biological tests have been further purified by preparative liquid chromatography, and all of them showed > 95% purity using the HPLC methods described above.

## Supporting information

Supplementary results and synthetic methods of compounds H1 - H28

NMR Spectra of compounds H1-H28 and D27

Supplementary Table1 for CRISPRi screen

## Acknowledgements

The authors are grateful for financial support from the National Natural Science Foundation of China (22271317 to L.H., 22101306 to Ming Z., 32100766 and 82171416 to R.T.), the Medical Innovation and Development Project of Lanzhou University (lzuyxcx-2022-156 to R.W.), the CAMS Innovation Fund for Medical Sciences (CIFMS) (2019-I2M-5-074, 2021-I2M-1-026, 2021-I2M-3-001 and 2022-I2M-2-002 to R.W.), Guangdong Basic and Applied Basic Research Foundation (2023B1515020075 to R.T.), the Science, Technology and Innovation Commission of Shenzhen Municipality (RCBS20210609103800006, JCYJ20220530112602006 and RCYX20221008092845052 to R.T.), the Lingang Laboratory Grant (LG-QS-202203-11 to R.T.), and the China Postdoctoral Science Foundation (2023M731523 to T.W.).

## Conflict of interest

All authors declare no conflict of interest.

## Author contributions

G.L., R.T., L.H., and R.W conceived and designed the project. Y.W., T.W., Man Z., A.H., F.S., L.C., R.L., Y.X., Ming Z., S.X. and Z.S. performed the experimental work. Y.W., T.W. and Man Z. analyzed the results and wrote the manuscript. Contributions to the experimental work include: compounds development and structure-activity relationship studies, A.H., Y.W., F.S., L.C., Man Z., Ming Z., S.X. and Z.S.; degradation mechanism verification and anti-tumor efficacy research in vivo and in vitro, Y.W., Man Z. and Y.X.; Identification of recruiting E3 ligase by pooled CRISPRi screening and validation, T.W. and R.L. All authors edited and approved the manuscript.

## Data Availability Statement

Data supporting the findings of this study is available in the supplementary information of this article.

## Notes

### Competing Interest Statement

The authors have declared no competing interest.

### Summary of Updates

we inserted “Very recently, while we were preparing this manuscript for reviewing, similar findings were reported by Waldmann and Winter et al (39). As alkenyl oxindole possesses Michael acceptor properties, they confirmed that it recruits DCAF11 through a covalent modification approach, potentially engaging with cysteine residues.” in the discussion section.

