## Supplementary results and synthetic methods of compounds H1 - H28 for "Discovery of Alkenyl Oxindole as a Novel PROTAC Moiety for Targeted Protein Degradation via CRL4^DCAF11^ Recruitment"

### Contents

**Figure S1:** Western blotting results of our developed compounds to degrade BRD4……………………......3

**Figure S2:** Western blotting results of BRD4 degradation by HL435 in multiple cells…………….........4

**Figure S3:** Concentration-dependent and kinetics studies of HL435-inducecd BRD4 degradation in MCF-7 cells.........................................................................................................................................4

**Figure S4:** The mechanism of BRD4 degradation by compounds conjugated alkenyl oxindoles…………...5

**Figure S5:** Effects of HL435 on cell cycle in MDA-MB-231 cells ……………………………………....6

**Figure S6:** Effects of HL435 on cell apoptosis in MCF-7 cells ……………………………………..….7

**Figure S7:** Effects of HL435 on the transcript levels of *c-MYC* and *P21* in BC cells……………………….8

The synthetic methods of compounds **H1 - H28**…………………………………………………………..9

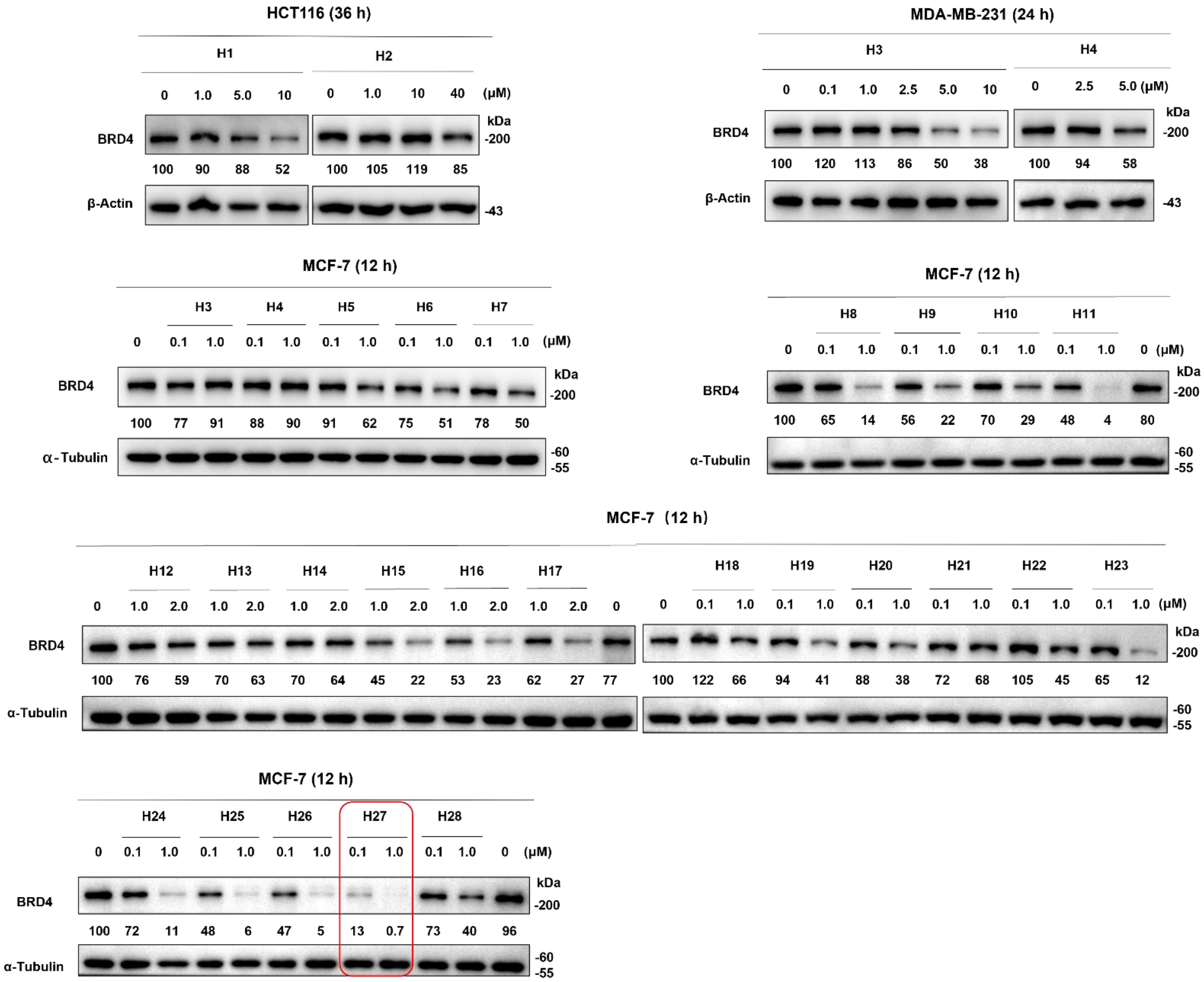


**Figure S1.** All compounds we developed could degrade BRD4 in a concentration-dependent manner. Western blotting results for BRD4 degradation. Image J was employed for relative quantitative analysis.

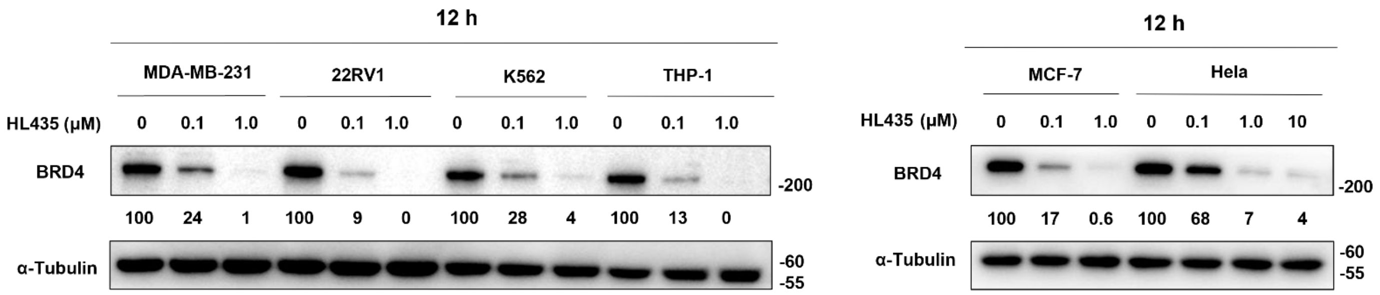


**Figure S2.** HL435 potently degraded BRD4 in multiple cell lines. Western blotting results for BRD4 degradation. Image J was employed for relative quantitative analysis.

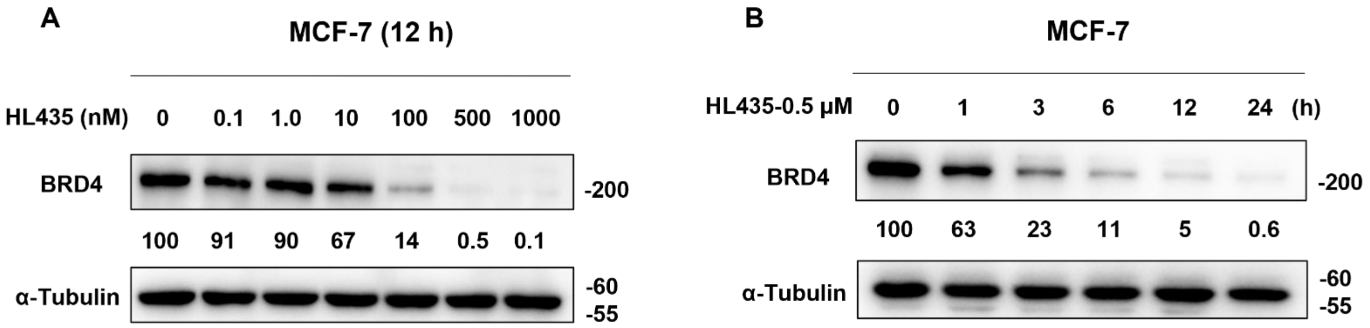


**Figure S3.** HL435 degraded BRD4 in concentration- and time-dependent manner in MCF-7 cells. (**A**) Representative Western blotting results for concentration-dependent studies of BRD4 degradation. (**B**) Representative Western blotting results for kinetics studies of BRD4 degradation. Image J was employed for relative quantitative analysis.

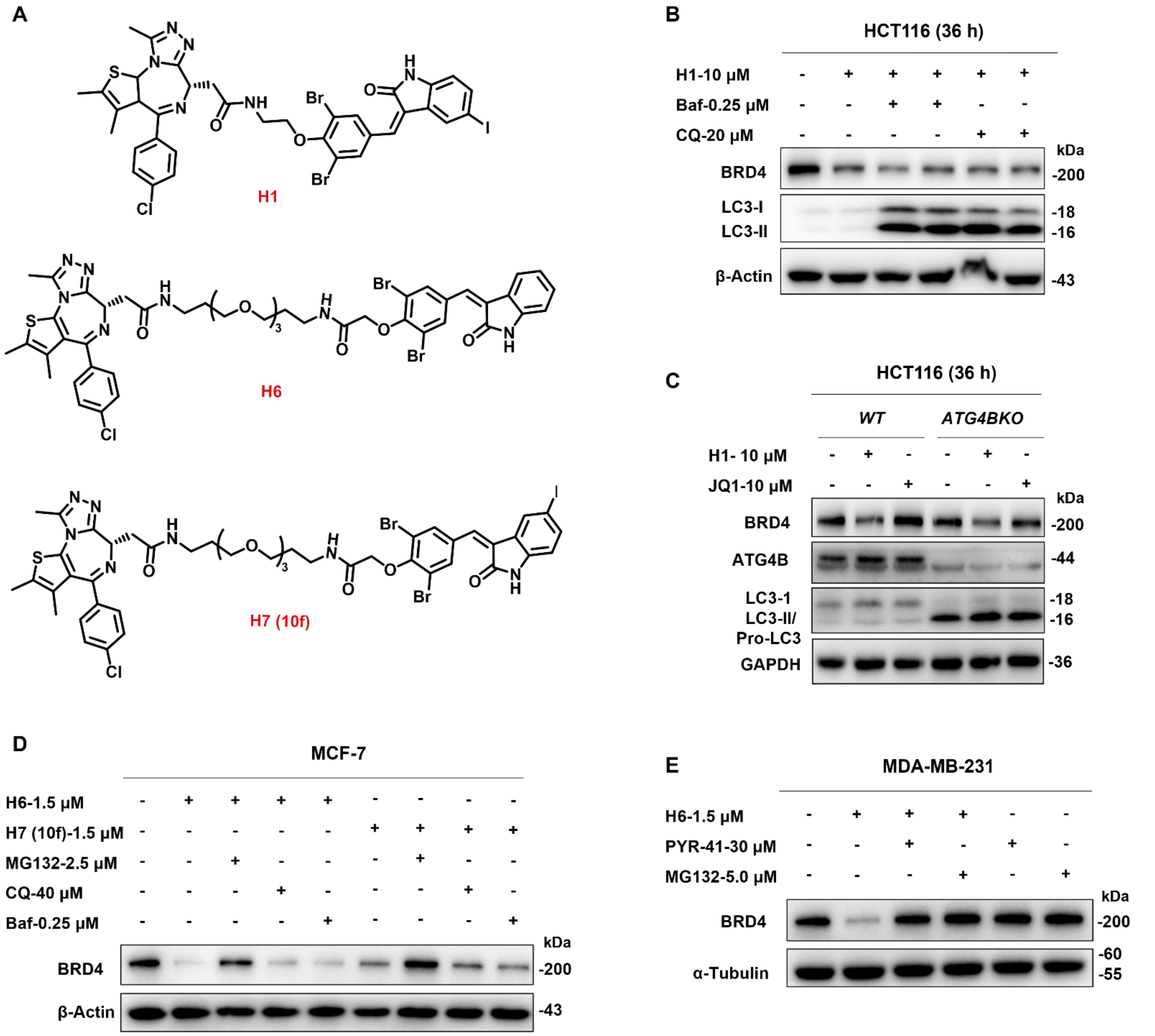


**Figure S4.** Compounds conjugated alkenyl oxindoles degraded BRD4 via ubiquitin proteasome pathway. (**A**) Structure of compounds conjugated alkenyl oxindoles. (**B**) Autophagy-lysosome inhibitor CQ or Baf cannot rescue the degradation of BRD4 by **H1**. Representative Immunoblotting results, cells were co-treated with **H1** and CQ or Baf for 36 h. (**C**) **H1** degraded BRD4 independent of LC3 and autophagosomes. Representative Immunoblotting results, WT- or *ATG4BKO*- HCT116 cells were treated with **H1** or JQ1 for 36 h. (**D**) Representative Immunoblotting results, cells were pre-treated with MG132, CQ or Baf for 2 h, followed by **H6** or **H7 (10f)** treatment for 6 h. (**E**) Representative Immunoblotting results, cells were pre-treated with PYR-41 or MG132 for 2 h, followed by **H6** treatment for 6 h.

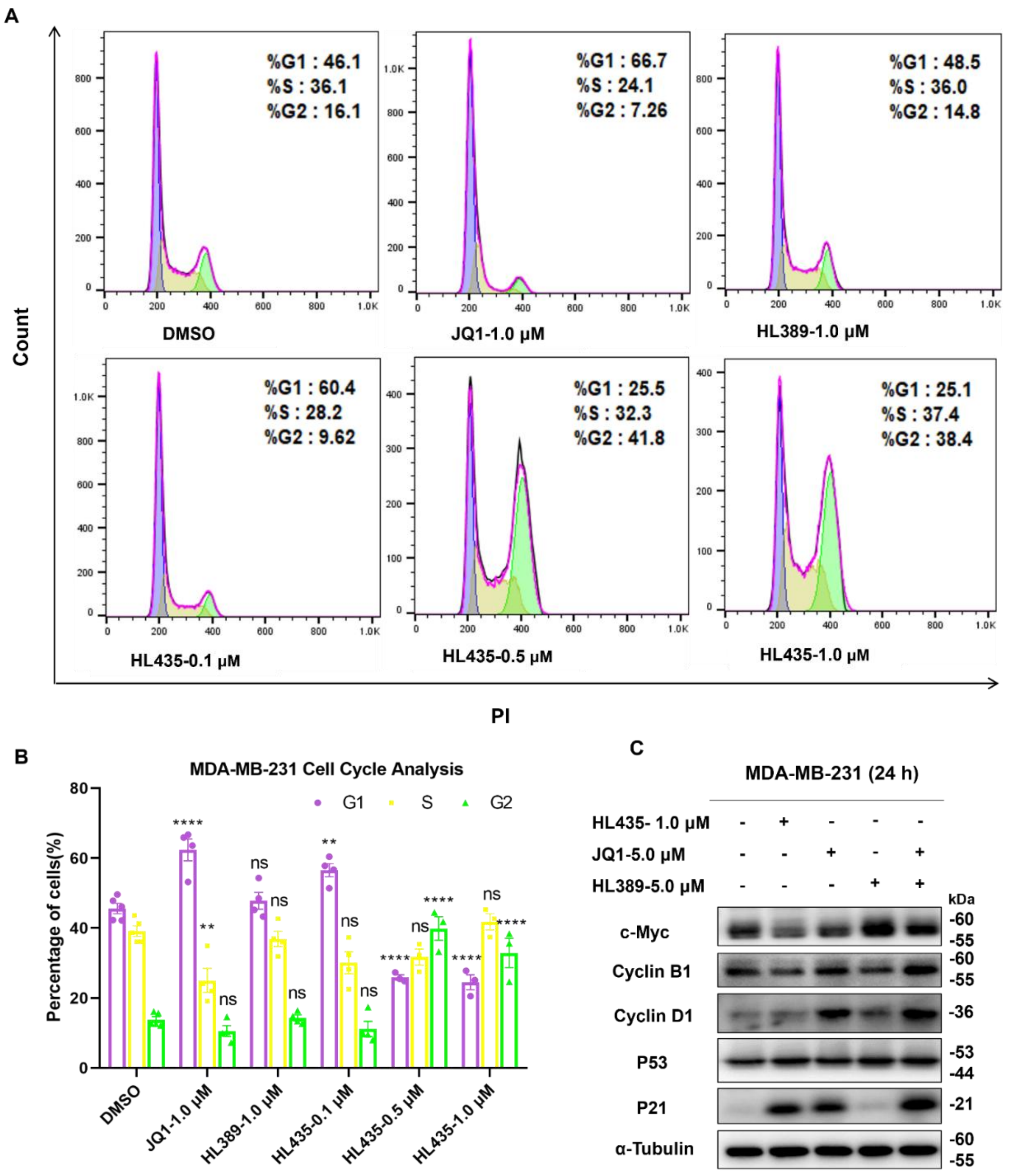


**Figure S5.** HL435 arrested cell cycle in MDA-MB-231 cells. (**A**) Representative flow cytometry analysis results of the cell cycle. MDA-MB-231 cells were treated with indicated compounds for 24 h before stained with PI. (**B**) Quantitative statistical analysis of cell cycle for **A**. (**C**) Representative Western blotting results of cycle relevant proteins in MDA-MB-231 cells. Data was presented as mean ± SEM (n=3). Statistical significance was determined by One-way ANOVA . **P < 0.01, ***P < 0.001, ****P < 0.0001; ns, no statistical significance.

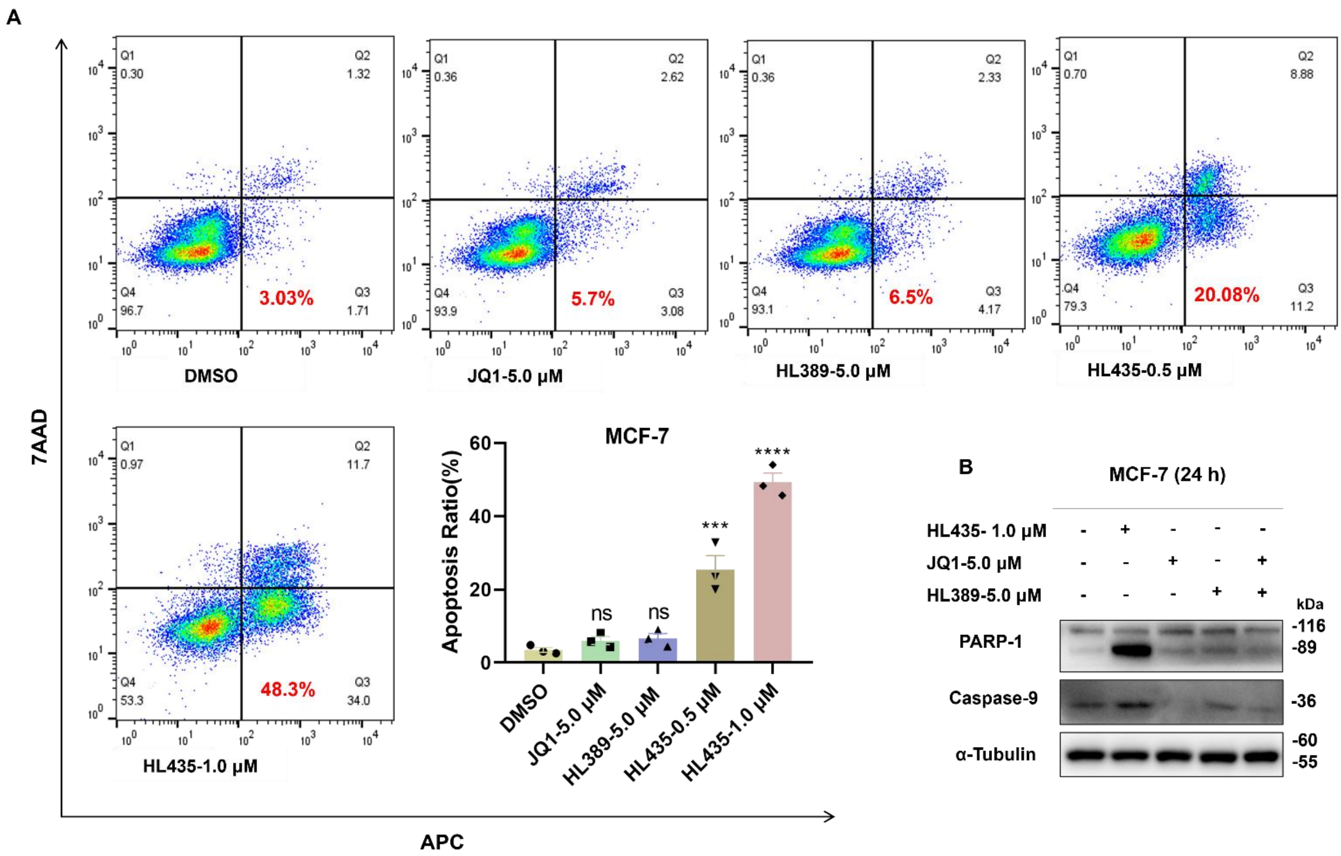


**Figure S6.** HL435 induced cell apoptosis in MCF-7 cells. (**A**) Representative flow cytometry analysis results and quantitative statistical analysis of apoptosis, MCF-7 cells were treated with indicated compounds for 36 h before stained with an 7AAD/APC Apoptosis Detection kit. (**B**) Representative Western blotting results of apoptosis relevant proteins. Data was presented as mean ± SEM (n=3). Statistical significance was determined by One-way ANOVA . ***P < 0.001, ****P < 0.0001; ns, no statistical significance.

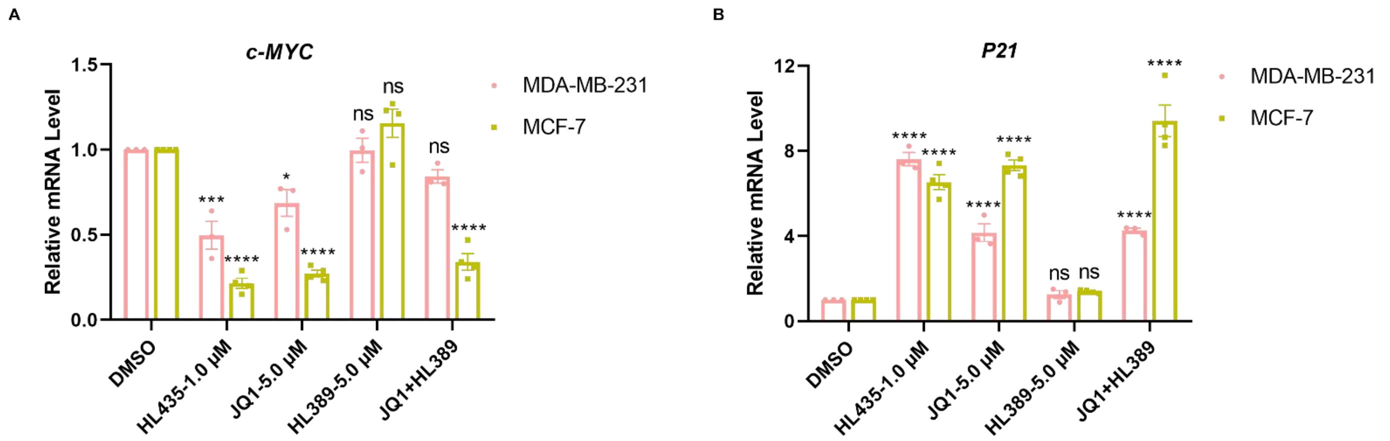


**Figure S7.** Effects of HL435 on the transcript levels of *c-MYC* and *P21* in BC cells. (**A**) qRT-PCR analysis of relative mRNA level of *c-MYC* in BC cells, treated with indicated compounds for 12 h. (**B**) qRT-PCR analysis of relative mRNA level of *P21*, BC cells were treated with indicated compounds for 12 h. Data was presented as mean ± SEM. Statistical significance was determined by One-way ANOVA . *P < 0.05, ***P < 0.001, ****P < 0.0001; ns, no statistical significance.

**The synthetic methods of compounds H1 - H28**

**(*S, Z*)-2-(4-(4-chlorophenyl)-2,3,9-trimethyl-6*H*-thieno[3,2-*f*][1,2,4]triazolo[4,3-*a*][1,4]diazepin-6-yl)-*N*-(2-(2,6-dibromo-4-((5-iodo-2-oxoindolin-3-ylidene)methyl)phenoxy)ethyl)acetamide (H1)**

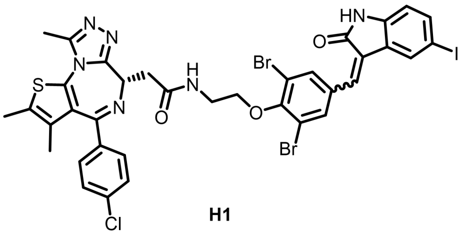

**H1**

The tert-butyl(2-bromoethyl)carbamate (2.91g, 13mmol) was added into a round-bottom flask filled with 3,5-dibromo-4-hydroxybenzaldehyde (**A1**; 2.80 g, 10 mmol), cesium carbonate (4.89g, 15mmol), potassium iodide (0.42g, 2.5 mmol), acetonitrile (70 mL) and magnetic stirrer. The mixture refluxed at 80℃ under argon atmosphere until the 3,5-dibromo-4-hydroxybenzaldehyde was consumed completely (monitored by TLC). Then, the mixture was extracted with methylene chloride (3 x 50 mL), and the combined organic layers were washed with saturated NaCl solution, and dried with anhydrous $Na_2SO_4$. Subsequently, the solvent was evaporated under vacuum and the residue was purified by flash column chromatography on silica gel to obtain the product **B1** (white solid 0.79 g, 18.8% yield). The 5-iodolindolin-2-one (**C1**, 0.26 g, 1.0 mmol) was added into the solution of **B1** (0.46 g, 1.1mmol), piperidine (15 uL) and $CH_3OH$ (10 mL), followed by refluxing at 65 ℃ for 13h. Then, the reaction product was purified by flash column chromatography to obtain the product **D1** (yellow solid 0.72 g, 77.32% yield).

5 mL HCl was added in 0.50 g **D1** (0.75 mmol) dissolved in 20 ml ethyl acetate and reacted at RT for 6 h. The mixture was extracted with ethyl acetate and ammonia, and the organic layers was purified to obtain the product **E1** (orange solid 0.41 g, 96% yield). Next, the mixture of JQ1 (carboxylic acid form, 0.4 g, 1.0 mmol), **E1** (0.41 g, 1.2 mmol), HBTU (0.56 g, 2.0 mmol) and DIPEA (0.6 mL, 4.0 mmol) were allowed to react in anhydrous DMF (3 mL) at room temperature for 3 h. After completion of the reaction, the reaction solution was extracted with ethyl acetate and the combine organic layers were dehydrated with anhydrous $Na_2SO_4$ and the solvent evaporated. The product was purified by flash column chromatography to afford **H1** (yellow solid 0.50 g, 53.0% yield, 95.2% purity). **$^1$H NMR** (400 MHz, DMSO-$d_6$) $\delta$ 10.79 (d, $J$ = 19.8 Hz, 1H), 8.79 (s, 1H), 8.53 (d, $J$ = 5.9 Hz, 1H), 8.02 (d, $J$ = 10.3 Hz, 1H), 7.84 (s, 1H), 7.70 (s, 1H), 7.62 – 7.50 (m, 2H), 7.44 (s, 4H), 6.71 (dd, $J$ = 19.8, 8.1 Hz, 1H), 4.56 (t, $J$ = 6.4 Hz, 1H), 4.10 (d, $J$ = 5.0 Hz, 2H), 3.75 – 3.54 (m, 4H), 2.61 (d, $J$ = 10.8 Hz, 3H), 2.40 (s, 3H), 1.62 (s, 3H). **$^{13}$C NMR** (101 MHz, DMSO-$d_6$) $\delta$ 170.43, 166.97, 163.52, 155.57, 154.18, 150.32, 143.21, 140.97, 139.04, 137.98, 137.20, 136.67, 135.69, 135.00, 134.17, 133.97, 133.69, 133.36, 132.70, 131.20, 130.65, 130.34, 130.06, 128.88, 128.49, 127.57,

127.30, 123.54, 118.41, 117.75, 113.19, 112.46, 84.68, 72.45, 54.22, 38.03, 14.53, 13.16, 11.77. **ESI-MS**: $m/z$ $[M + H]^+$ calcd for $C_{36}H_{29}O_3N_6ClBr_2IS^+$, 946.9088; found, 946.9087.

**Scheme I**

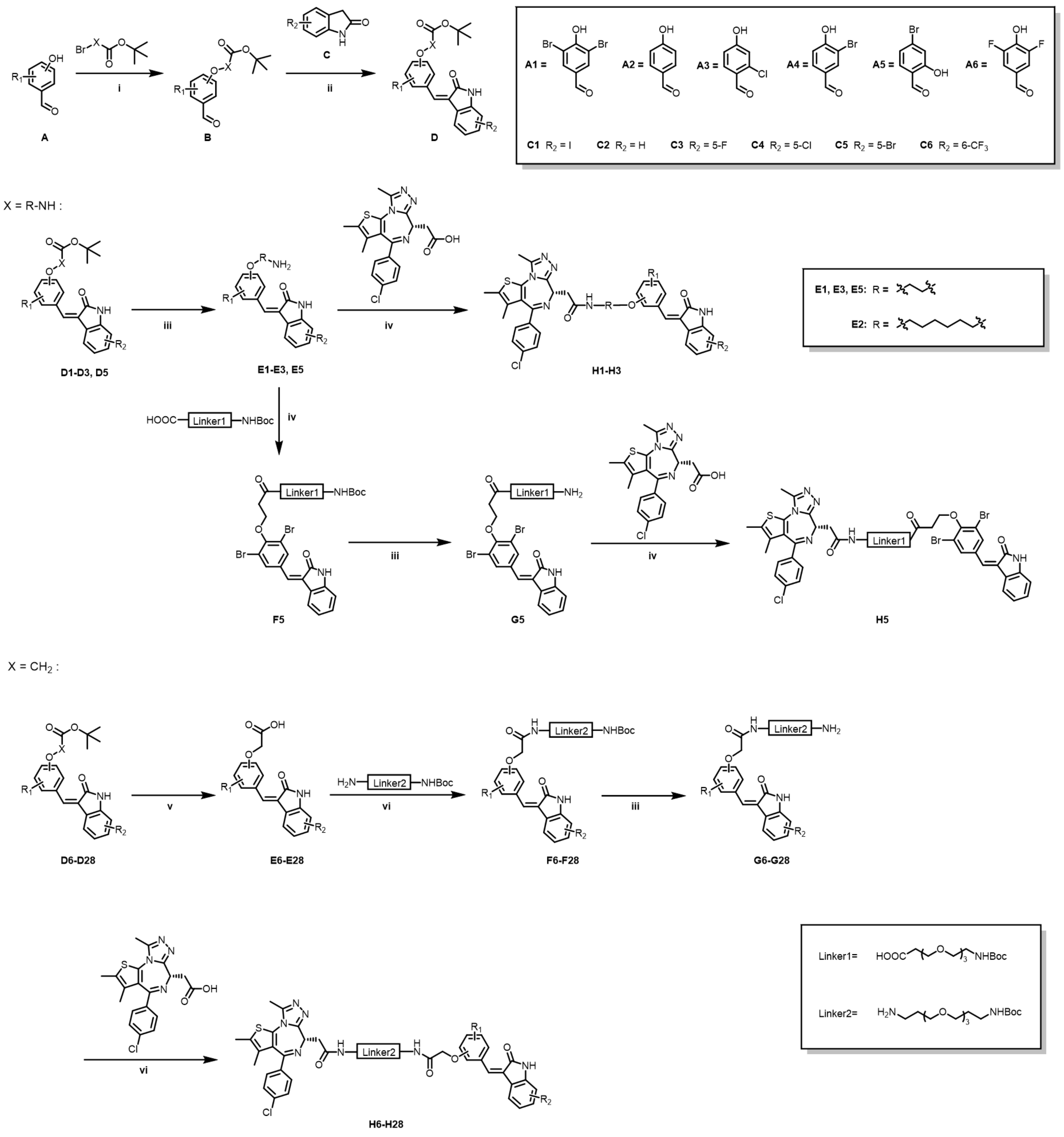


Scheme I: i) $Cs_2CO_3$, KI, Acetonitrile, 80℃ reflux; ii) Piperidine, $CH_3OH$, 65℃ reflux; iii) HCl, EA, rt; iv) HBTU, DIPEA, DMF, rt; v) TFA, DCM, rt; vi) HOBT, EDCI, DIPEA, DMF, rt.

**Scheme II**

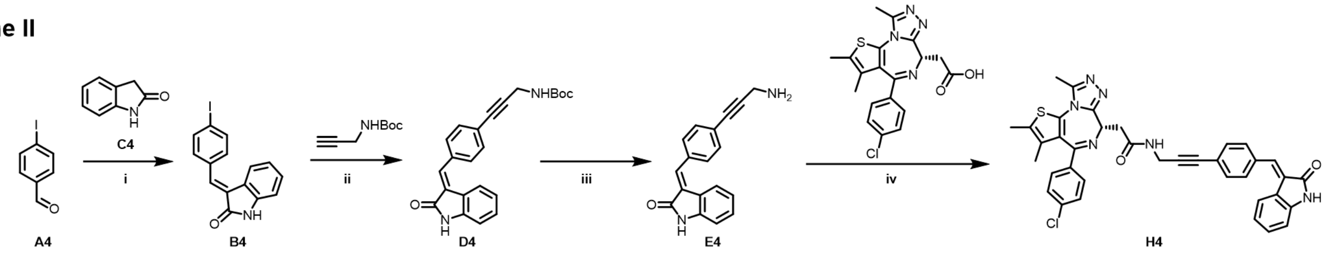


Scheme II: i) Piperidine, $CH_3OH$, 65℃ reflux; ii) $PdCl_2(Pph)_3$, CuI, $Et_3N$, 50℃, Argon; iii) HCl, EA, rt; iv) HBTU, DIPEA, DMF, rt.

**(*S, Z*)-2-(4-(4-chlorophenyl)-2,3,9-trimethyl-6*H*-thieno[3,2-*f*][1,2,4]triazolo[4,3-*a*][1,4]diazepin-6-yl)-*N*-(6-(2,6-dibromo-4-((5-iodo-2-oxoindolin-3-ylidene)methyl)phenoxy)hexyl)acetamide (H2)**

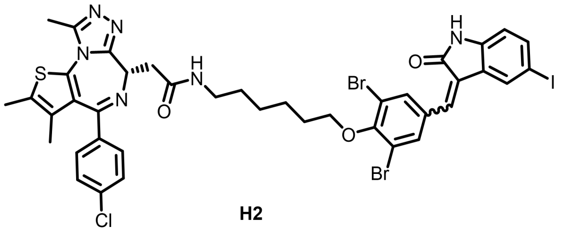

H2

The title compound **H2** (yellow solid, 35.2% yield) was synthesized according to the procedures for the preparation of **H1** from **A1** (3,5-dibromo-4-hydroxybenzaldehyde), **C1** (5-iodoindolin-2-one) and tert-butyl (6-bromohexyl) carbamate. **$^1$H NMR** (400 MHz, DMSO-$d_6$) $\delta$ 10.83 (s, 1H), 8.79 (s, 1H), 8.22 (s, 1H), 8.06 (s, 1H), 8.00 (s, 1H), 7.85 (s, 1H), 7.58 (t, $J$ = 8.7 Hz, 2H), 7.47 (dd, $J$ = 18.1, 8.4 Hz, 5H), 6.75 (dd, $J$ = 20.6, 8.2 Hz, 1H), 4.54 (s, 1H), 4.04 (dd, $J$ = 11.5, 5.8 Hz, 3H), 3.37 – 2.98 (m, 5H), 2.61 (s, 3H), 2.42 (s, 3H), 1.84 (d, $J$ = 6.8 Hz, 3H), 1.64 (s, 3H), 1.54 (d, $J$ = 6.7 Hz, 5H), 1.44 (s, 3H). **$^{13}$C NMR** (101 MHz, DMSO-$d_6$) $\delta$ 169.93, 166.98, 163.51, 155.59, 154.56, 154.04, 150.32, 149.29, 148.86, 143.19, 140.93, 137.19, 136.66, 135.74, 135.08, 134.24, 133.94, 133.44, 133.14, 132.69, 131.22, 130.56, 130.30, 128.91, 127.58, 127.17, 117.81, 112.46, 99.99, 84.67, 73.99, 54.39, 38.92, 38.16, 31.61, 29.96, 29.44, 26.64, 25.63, 14.53, 13.14, 11.74. **ESI-MS**: *m/z* $[M + H]^+$ calcd for $C_{40}H_{37}O_3N_6ClBr_2IS^+$, 1002.9722; found, 1002.9720; purity: 97%.

**(*S, Z*)-2-(4-(4-chlorophenyl)-2,3,9-trimethyl-6*H*-thieno[3,2-*f*][1,2,4]triazolo[4,3-*a*][1,4]diazepin-6-yl)-*N*-(2-(2,6-dibromo-4-((2-oxoindolin-3-ylidene)methyl)phenoxy)ethyl)acetamide (H3)**

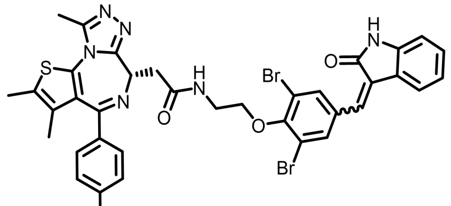

H3

The title compound **H3** (yellow solid, 54.7% yield) was synthesized according to the procedures for the preparation of **H1** from **A1** (3,5-dibromo-4-hydroxybenzaldehyde), **C2** (indolin-2-one) and tert-butyl(2-bromoethyl) carbamate. **$^1$H NMR** (400 MHz, DMSO-$d_6$) $\delta$ 10.68 (d, $J$ = 24.4 Hz, 1H), 8.80 (s, 1H), 8.53 (s, 1H), 8.00 (s, 1H), 7.84 – 7.60 (m, 2H), 7.49 (d, $J$ = 28.7 Hz, 5H), 7.24 (s, 1H), 7.15 – 6.77 (m, 2H), 4.56 (s, 1H), 4.09 (s, 2H), 3.60 (s, 3H), 2.94 (d, $J$ = 17.6 Hz, 1H), 2.60 (s, 3H), 2.41 (s, 3H), 1.63 (s, 3H). **$^{13}$C NMR** (101 MHz, DMSO-$d_6$) $\delta$ 170.48, 167.55, 167.23, 163.56, 155.57, 153.84, 153.46, 150.37, 137.19, 136.95, 136.39, 135.69, 133.80, 133.32, 132.67, 131.86, 131.25, 130.65, 130.34, 130.07, 128.87, 124.91, 122.71, 121.82, 121.63, 121.47, 120.90, 120.76, 120.57, 118.39, 117.66, 114.71, 110.11, 100.00, 72.37, 54.21, 14.51, 13.14, 11.74. **ESI-MS**: $m/z$ $[M + H]^+$ calcd for $C_{36}H_{30}O_3N_6ClBr_2S^+$, 821.0131; found, 821.0116; purity: > 99%.

**(*S, E*)-2-(4-(4-chlorophenyl)-2,3,9-trimethyl-6*H*-thieno[3,2-*f*][1,2,4]triazolo[4,3-*a*][1,4]diazepin-6-yl)-*N*-(3-(4-((2-oxoindolin-3-ylidene)methyl)phenyl)prop-2-yn-1-yl)acetamide (H4)**

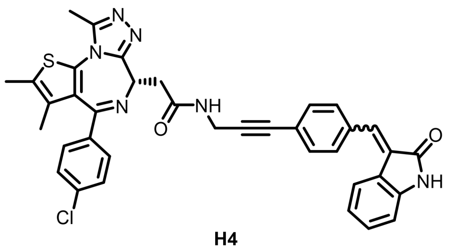

**H4**

The indolin-2-one (0.21 g, 1.6 mmol) was added into a round-bottom flask filled with 4-iodobenzaldehyde (**A4**; 0.46 g, 2 mmol), pyridine (34.06 mg, 0.4 mmol), $CH_3OH$ (8 mL) and magnetic stirrer. The mixture refluxed at 65 ℃ until the indolin-2-one was consumed completely. After solvent evaporated, the residue was purified by flash column chromatography to afford **B4** (0.49 g, 87.5% yield). Then, the mixture of **B4** (0.49 g, 1.4 mmol), tert-butyl prop-2-yn-1-ylcarbamate (0.33 g, 2.1 mmol), $PdCl_2(Pph)_3$ (24.74 mg, 0.036 mmol), CuI (13.42 mg, 0.007 mmol) and $Et_3N$ (3 mL) were allowed to react at 50℃ under argon atmosphere to afford the product **C4** (0.48 g, 91.6% yield). After that, 5 mL HCl was added into 0.48 g **C4** (1.28 mmol) dissolved in 20 ml ethyl acetate and reacted at rt for 6 h. The mixture was extracted with ethyl acetate and ammonia, and the organic layers was purified to obtain the product **D4** (0.31 g, 88.9% yield). Next, the mixture of JQ1 (carboxylic acid form, 0.4 g, 1.0 mmol), **D4** (0.31 g, 1.14 mmol), HBTU (0.56 g, 2.0 mmol) and DIPEA (0.6 mL, 4.0 mmol) were allowed to react in anhydrous DMF (3 mL) at room temperature for 3 h. After completion of the reaction, the reaction solution was extracted with ethyl acetate and the combine organic layers were dehydrated with anhydrous $Na_2SO_4$ and the solvent evaporated. The product was purified by flash column chromatography to afford **H4** (yellow solid 0.32 g, 48.3% yield).

**$^1$H NMR** (400 MHz, DMSO-*d*$_6$) $\delta$ 10.66 (d, J = 12.4 Hz, 1H), 8.85 (s, 1H), 8.42 (d, J = 8.1 Hz, 1H), 7.73 (d, J = 7.9 Hz, 2H), 7.62 (s, 1H), 7.58 (d, J = 8.0 Hz, 1H), 7.54 – 7.47 (m, 1H), 7.43 (d, J = 8.3 Hz, 2H), 7.34 (d, J = 8.3 Hz, 2H), 7.24 (dd, J = 14.2, 7.0 Hz, 1H), 6.89 (d, J = 7.8 Hz, 1H), 6.84 (t, J = 7.3 Hz, 1H), 4.63 – 4.42 (m, 1H), 4.26 (ddd, J = 22.2, 17.7, 5.2 Hz, 2H), 3.28 (d, J = 5.3 Hz, 2H), 2.61 (s, 3H), 2.41 (s, 3H), 1.61 (s, 3H). **$^{13}$C NMR** (101 MHz, DMSO-*d*$_6$) $\delta$ 170.05, 169.02, 167.53, 163.67, 155.45, 150.46, 143.55, 141.39, 137.12, 135.95, 135.70, 135.17, 135.07, 134.53, 132.73, 132.53, 132.24, 131.65, 131.32, 130.89, 130.64, 130.32, 130.12, 130.03, 128.83, 128.61, 125.25, 123.90, 123.00, 121.67, 121.15, 120.48, 110.71, 109.91, 89.62, 81.76, 54.30, 37.88, 31.99, 29.88, 29.18, 14.49, 13.15, 11.76. **ESI-MS**: *m*/*z* [M + H]$^+$ calcd for $C_{37}H_{29}O_2N_6ClS^+$, 657.1834; found, 657.1835; purity: > 99%.

**(*S*,*Z*)-3-(2-(2-(2-(2-(4-(4-chlorophenyl)-2,3,9-trimethyl-6*H*-thieno[3,2-*f*][1,2,4]triazolo[4,3-*a*][1,4] diazepin-6-yl)acetamido)ethoxy)ethoxy)ethoxy)-*N*-(2-(2,6-dibromo-4-((2-oxoindolin-3-ylidene)methyl) phenoxy)ethyl)propenamide (H5)**

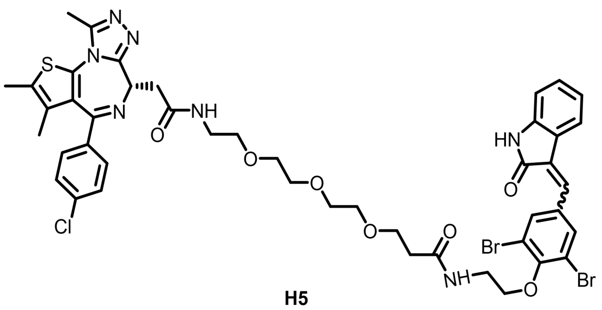

H5

The compound **E5** was synthesized through the same synthetic method of **E1**. The mixture of **E5** (0.44 g, 1.0 mmol), 2,2-dimethyl-4-oxo-3,8,11,14-tetraoxa-5-azaheptadecan-17-oic acid (Linker1, 0.39 g, 1.2 mmol), HBTU (0.56 g, 2.0 mmol) and DIPEA (0.6 mL, 4.0 mmol) were reacted in anhydrous DMF (2 mL) at room temperature for 3 h. After completion of the reaction, the reaction solution was extracted with ethyl acetate and the combine organic layers were dehydrated with anhydrous $Na_2SO_4$ and the solvent evaporated. The product was purified by flash column chromatography to afford **F5** (0.31 g, 41.2% yield) and was dissolved in 10 mL ethyl acetate follow by adding 4 mL HCl to react at RT for 6 h. The mixture was extracted with ethyl acetate and ammonia, and the organic layers was purified to obtain the product **G5** (0.23 g, 86.3% yield). Then, JQ1 (carboxylic acid form, 0.12 g, 0.3 mmol), **G5** (0.23 g, 0.36 mmol), HBTU (0.19 g, 0.67 mmol) and DIPEA (0.2 mL, 1.33 mmol) were dissolved in 2 mL anhydrous DMF and reacted at room temperature for 3 h. After completion of the reaction, the reaction solution was extracted with ethyl acetate and the combine organic layers were dehydrated with anhydrous $Na_2SO_4$ and the solvent evaporated. The product was purified by flash column chromatography to afford **H5** (yellow solid 0.12 g, 37.5% yield). **$^1$H NMR** (400 MHz, DMSO-*d*$_6$) $\delta$ 10.68 (d, *J* = 24.5 Hz, 1H), 8.75 (d, *J* = 24.8 Hz, 1H), 8.28 (s, 1H), 8.16 (s,

1H), 7.95 (d, $J$ = 24.9 Hz, 1H), 7.80 – 7.61 (m, 2H), 7.45 (d, $J$ = 13.8 Hz, 5H), 7.32 – 6.74 (m, 1H), 4.52 (s, 1H), 4.01 (d, $J$ = 17.3 Hz, 4H), 3.54 (dd, $J$ = 36.8, 26.6 Hz, 16H), 3.28 (s, 4H), 2.59 (s, 3H), 2.39 (t, $J$ = 21.2 Hz, 6H), 1.61 (s, 3H). **$^{13}$C NMR** (101 MHz, DMSO-$d_6$) $\delta$ 170.94, 170.23, 163.51, 155.58, 137.22, 136.38, 135.70, 133.79, 132.71, 131.86, 131.20, 130.61, 130.31, 130.06, 128.91, 121.77, 118.34, 117.62, 72.41, 70.23, 70.09, 69.66, 67.19, 64.25, 54.29, 37.98, 36.54, 14.50, 13.13, 11.74. **ESI-MS**: $m/z$ [M + H]$^+$ calcd for $C_{45}H_{47}O_7N_7ClBr_2S^+$, 1024.1078; found, 1024.1074; purity: > 99%.

**(*S, Z*)-2-(4-(4-chlorophenyl)-2,3,9-trimethyl-6*H*-thieno[3,2-*f*][1,2,4]triazolo[4,3-*a*][1,4]diazepin-6-yl)-*N*-(1-(2,6-dibromo-4-((2-oxoindolin-3-ylidene)methyl)phenoxy)-2-oxo-7,10,13-trioxa-3-azahexadecan-16-yl)acetamide (H6)**

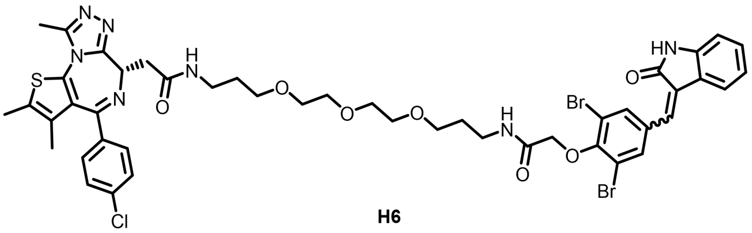

H6

The compound **D6** was synthesized through the same synthetic method of **D1**. Trifluoroacetic acid (TFA, 3 mL) was added to 3 ml of methylene chloride dissolved **D6** (0.51 g, 1.0 mmol) and stirred at room temperature. After full conversion of **D6**, the reaction mixture was concentrated and the product was purified by column chromatography to afford **E6** (0.42g, 93.4% yield). Then, the mixture of **E6** (0.42g, 0.93 mmol), tert-butyl (3-(2-(2-(3-aminopropoxy)ethoxy)ethoxy)propyl) carbamate (Linker2, 0.36 g, 1.13 mmol), EDCI (0.27 g, 1.40 mmol), HOBT (0.40 g, 1.86 mmol) and DIPEA (0.45 mL, 2.80 mmol) in 10 mL of DMF was stirred at room temperature for 3 h. After completion of the reaction, the mixture was extracted with ethyl acetate and the combined organic layers were dried over $Na_2SO_4$. After the solvent was evaporated, the **F6** (0.29 g, 41.2% yield) was purified by flash column chromatography. The **F6** (0.29 g, 0.38 mmol) was dissolved in 10 mL ethyl acetate follow by adding 4 mL HCl to react at RT for 6 h. The mixture was extracted with ethyl acetate and ammonia, and the organic layers was purified to obtain the product **G6** (0.19 g, 77.6% yield). Then, the mixture of JQ1 (carboxylic acid form, 0.10 g, 0.25 mmol), **G6** (0.19 g, 0.30 mmol), EDCI (0.23 g, 1.17 mmol), HOBT (0.34 g, 1.57 mmol) and DIPEA (0.38 mL, 2.34 mmol) in 10 mL of DMF was stirred at room temperature for 3 h. After completion of the reaction, the mixture was extracted with ethyl acetate and the combined organic layers were dried over $Na_2SO_4$. After the solvent was evaporated, the **F6** (0.10 g, 38.9% yield) was purified by flash column chromatography. **$^1$H NMR** (400 MHz, DMSO-$d_6$) $\delta$ 10.69 (d, $J$ = 26.1 Hz, 1H), 8.80 (s, 1H), 8.23 – 8.09 (m, 2H), 7.99 (s, 1H), 7.45 (td, $J$ = 20.1, 11.9 Hz, 6H), 7.25 (dd, $J$ = 13.8, 7.0 Hz, 1H), 6.87 (dd, $J$ = 17.5, 8.9 Hz, 2H), 4.51 (d, $J$ = 6.9 Hz, 1H), 4.45 (d, $J$ = 10.9 Hz, 2H), 3.56 – 3.40 (m, 13H), 3.31 – 3.07 (m, 8H), 2.59 (s, 3H), 2.40 (s, 3H), 1.78 – 1.65 (m,

4H), 1.62 (s, 3H). **$^{13}$C NMR** (101 MHz, DMSO-$d_6$) $\delta$ 169.96, 168.65, 168.57, 167.51, 166.69, 163.50, 155.57, 152.88, 152.53, 150.27, 143.82, 141.57, 137.22, 136.37, 135.71, 134.58, 133.94, 133.77, 133.12, 132.71, 132.46, 131.25, 131.17, 130.57, 130.31, 130.04, 128.93, 124.89, 121.79, 120.88, 120.59, 118.16, 117.41, 110.10, 71.45, 70.25, 70.10, 70.01, 68.89, 68.53, 54.36, 38.14, 36.59, 36.28, 29.92, 29.62, 14.50, 13.13, 11.75. **ESI-MS**: *m*/*z* [M + H]$^+$ calcd for $C_{46}H_{49}O_7N_7ClBr_2S^+$, 1038.1445; found, 1038.1439; purity: > 99%.

**(*S, Z*)-2-(4-(4-chlorophenyl)-2,3,9-trimethyl-6*H*-thieno[3,2-*f*][1,2,4]triazolo[4,3-*a*][1,4]diazepin-6-yl)-*N*-(1-(2,6-dibromo-4-((5-iodo-2-oxoindolin-3-ylidene)methyl)phenoxy)-2-oxo-7,10,13-trioxa-3-azahexadecan-16-yl)acetamide (H7)**

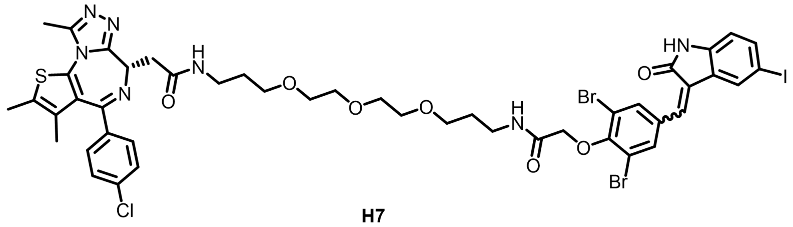


H7

The title compound **H7** (yellow solid, 31.2% yield) was synthesized according to the procedures for the preparation of **H6** from **A1**(3,5-dibromo-4-hydroxybenzaldehyde), **C1** (5-iodoindolin-2-one), tert-butyl 2-bromoacetate and Linker2. **$^1$H NMR** (400 MHz, DMSO-$d_6$) $\delta$ 10.81 (d, $J$ = 23.8 Hz, 1H), 8.78 (s, 1H), 8.27 – 8.10 (m, 2H), 8.03 (d, $J$ = 6.9 Hz, 1H), 7.85 (s, 1H), 7.61 – 7.37 (m, 6H), 6.72 (dd, $J$ = 20.7, 8.0 Hz, 1H), 4.51 (t, $J$ = 6.8 Hz, 1H), 4.45 (d, $J$ = 9.6 Hz, 2H), 3.47 (dd, $J$ = 16.7, 11.5 Hz, 15H), 3.31 – 3.09 (m, 6H), 2.59 (s, 3H), 2.39 (s, 3H), 1.78 – 1.65 (m, 4H), 1.60 (s, 3H). **$^{13}$C NMR** (101 MHz, DMSO-$d_6$) $\delta$ 171.70, 169.96, 168.02, 166.95, 166.65, 163.50, 155.56, 153.23, 152.75, 150.27, 143.26, 141.02, 139.88, 139.12, 137.22, 136.62, 135.70, 134.82, 134.12, 133.95, 133.73, 132.71, 131.16, 130.57, 130.30, 130.02, 128.93, 128.67, 127.55, 123.50, 118.19, 117.50, 113.21, 112.48, 84.67, 84.36, 71.45, 70.25, 70.10, 70.01, 68.89, 68.53, 54.36, 38.13, 36.59, 36.28, 29.92, 29.62, 14.51, 13.14, 11.76. **ESI-MS**: *m*/*z* [M + H]$^+$ calcd for $C_{46}H_{48}O_7N_7ClBr_2IS^+$, 1164.0401; found, 1164.0399; purity: > 99%.

**(*S, Z*)-2-(4-(4-chlorophenyl)-2,3,9-trimethyl-6*H*-thieno[3,2-*f*][1,2,4]triazolo[4,3-*a*][1,4]diazepin-6-yl)-*N*-(1-(2,6-dibromo-4-((5-fluoro-2-oxoindolin-3-ylidene)methyl)phenoxy)-2-oxo-7,10,13-trioxa-3-azahexadecan-16-yl)acetamide (H8)**

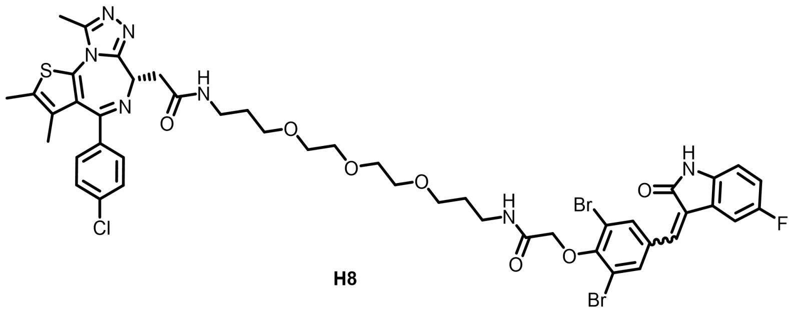

**H8**

The title compound **H8** (yellow solid, 24.6% yield) was synthesized according to the procedures for the preparation of **H6** from **A1** (3,5-dibromo-4-hydroxybenzaldehyde), **C3** (5-fluoroindolin-2-one), tert-butyl 2-bromoacetate and Linker2. **$^{1}$H NMR** (400 MHz, DMSO-*d*$_6$) $\delta$ 10.05 (s, 1/3H), 9.81 (s, 1/3H), 8.82 (s, 2/3H), 7.96 (s, 1H), 7.74 (d, *J* = 12.2 Hz, 5/3H), 7.60 (s, 1H), 7.50 (d, *J* = 8.1 Hz, 3H), 7.41 (d, *J* = 7.9 Hz, 2H), 7.21 (d, *J* = 8.9 Hz, 2/3H), 7.11 – 6.86 (m, 2H), 4.65 (s, 1H), 4.53 (d, *J* = 11.5 Hz, 2H), 3.61 (s, 6H), 3.55 (s, 6H), 3.48 (d, *J* = 6.0 Hz, 3H), 3.36 (d, *J* = 7.8 Hz, 4H), 2.62 (s, 3H), 2.44 (s, 3H), 1.87 (s, 2H), 1.79 (s, 2H), 1.69 (d, *J* = 4.5 Hz, 3H). **$^{13}$C NMR** (101 MHz, DMSO-*d*$_6$) $\delta$ 169.90, 168.60, 167.50, 166.68, 166.62, 163.47, 156.64, 155.56, 152.77, 150.25, 137.21, 136.57, 135.70, 134.82, 134.14, 133.80, 133.65, 132.73, 131.14, 130.57, 130.29, 130.02, 129.49, 128.93, 118.28, 117.51, 71.44, 70.25, 70.10, 70.01, 68.87, 68.52, 54.36, 38.12, 36.56, 36.25, 29.93, 29.64, 14.51, 13.13, 11.76. **ESI-MS**: *m*/*z* [M + H]$^+$ calcd for $C_{46}H_{48}O_7N_7ClBr_2FS^+$, 1056.1349; found, 1056.1354; purity: > 99%.

**(*S, Z*)-2-(4-(4-chlorophenyl)-2,3,9-trimethyl-6*H*-thieno[3,2-*f*][1,2,4]triazolo[4,3-*a*][1,4]diazepin-6-yl)-*N*-(1-(2,6-dibromo-4-((5-chloro-2-oxoindolin-3-ylidene)methyl)phenoxy)-2-oxo-7,10,13-trioxa-3-azahexadecan-16-yl)acetamide (H9)**

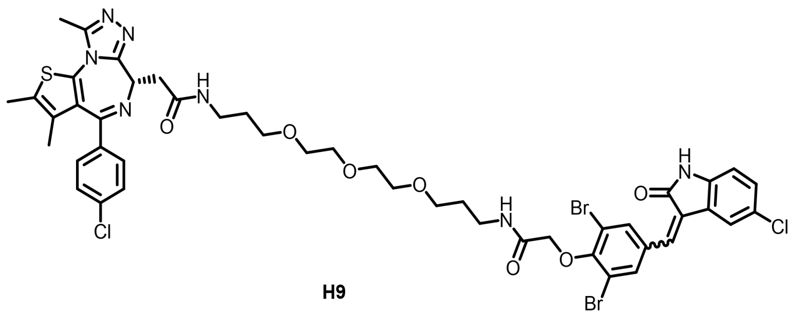

**H9**

The title compound **H9** (yellow solid, 33.2% yield) was synthesized according to the procedures for the preparation of **H6** from **A1** (3,5-dibromo-4-hydroxybenzaldehyde), **C4** (5-chloroindolin-2-one), tert-butyl 2-bromoacetate and Linker2. **$^{1}$H NMR** (400 MHz, DMSO-*d*$_6$) $\delta$ 10.13 (s, 1/3H), 9.88 (s, 1/3H), 8.82 (s, 1H), 7.98 (s, 1H), 7.80 (s, 2/3H), 7.72 (s, 5/3H), 7.59 (d, *J* = 10.1 Hz, 1H), 7.49 (t, *J* = 9.2 Hz, 3H), 7.41 (d, *J* = 8.1 Hz, 2H), 7.26 (dd, *J* = 16.5, 8.2 Hz, 1H), 7.08 – 6.86 (m, 1H), 4.66 (d, *J* = 6.5 Hz, 1H), 4.53 (d, *J* = 11.2 Hz, 2H), 3.61 (s, 6H), 3.56 (d, *J* = 8.3 Hz, 6H), 3.48 (d, *J* = 5.9 Hz, 3H), 3.42 – 3.25 (m, 4H), 2.62 (s, 3H), 2.44 (s, 3H), 1.87 (d, *J* = 3.5 Hz, 2H), 1.79 (d, *J* = 4.1 Hz, 2H), 1.69 (d, *J* = 6.9 Hz, 3H). **$^{13}$C NMR** (101

MHz, DMSO-$d_6$) $\delta$ 169.94, 167.24, 166.62, 163.48, 159.78, 155.55, 153.23, 152.14, 150.23, 149.91, 140.21, 138.59, 136.62, 135.70, 135.03, 133.90, 133.66, 133.21, 132.72, 131.13, 130.56, 130.29, 130.00, 128.92, 127.86, 126.07, 125.56, 120.59, 118.25, 117.51, 113.97, 111.48, 108.58, 100.00, 71.43, 70.25, 70.11, 70.01, 68.88, 68.53, 54.36, 38.13, 36.57, 36.27, 29.93, 29.64, 14.50, 13.12, 11.76. **ESI-MS**: *m*/*z* $[M + H]^+$ calcd for $C_{46}H_{48}O_7N_7Cl_2Br_2S^+$, 1072.1054; found, 1072.1058; purity: > 99%.

**(*S, Z*)-2-(4-(4-chlorophenyl)-2,3,9-trimethyl-6*H*-thieno[3,2-*f*][1,2,4]triazolo[4,3-*a*][1,4]diazepin-6-yl)-*N*-(1-(2,6-dibromo-4-((5-bromo-2-oxoindolin-3-ylidene)methyl)phenoxy)-2-oxo-7,10,13-trioxa-3-azahexa decan-16-yl)acetamide (H10)**

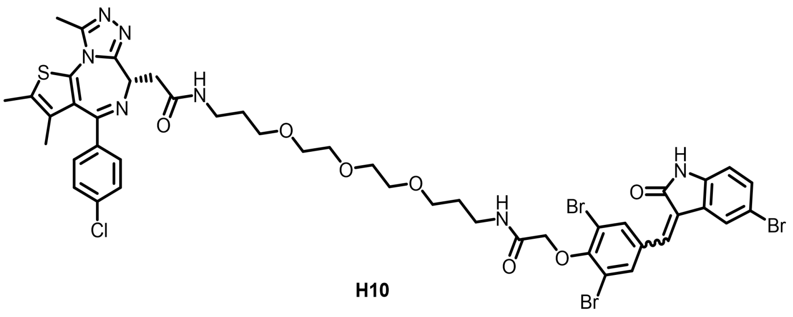

**H10**

The title compound **H10** (yellow solid, 56.8% yield) was synthesized according to the procedures for the preparation of **H6** from **A1** (3,5-dibromo-4-hydroxybenzaldehyde), **C5** (5- bromoindolin-2-one), tert-butyl 2-bromoacetate and Linker2. $^1$**H NMR** (400 MHz, DMSO-$d_6$) $\delta$ 10.13 (s, 1/3H), 9.88 (s, 1/3H), 8.82 (s, 1H), 7.98 (s, 1H), 7.80 (s, 2/3H), 7.72 (s, 5/3H), 7.59 (d, *J* = 10.1 Hz, 1H), 7.49 (t, *J* = 9.2 Hz, 3H), 7.41 (d, *J* = 8.1 Hz, 2H), 7.26 (dd, *J* = 16.5, 8.2 Hz, 1H), 7.08 – 6.86 (m, 1H), 4.66 (d, *J* = 6.5 Hz, 1H), 4.53 (d, *J* = 11.2 Hz, 2H), 3.61 (s, 6H), 3.56 (d, *J* = 8.3 Hz, 6H), 3.48 (d, *J* = 5.9 Hz, 3H), 3.42 – 3.25 (m, 4H), 2.62 (s, 3H), 2.44 (s, 3H), 1.87 (d, *J* = 3.5 Hz, 2H), 1.79 (d, *J* = 4.1 Hz, 2H), 1.69 (d, *J* = 6.9 Hz, 3H). $^{13}$**C NMR** (101 MHz, DMSO-$d_6$) $\delta$ 169.93, 169.77, 168.20, 167.12, 166.61, 163.48, 155.55, 153.26, 150.24, 142.89, 140.58, 138.34, 137.21, 136.65, 135.70, 135.07, 134.29, 134.07, 133.93, 133.39, 132.73, 132.24, 131.13, 130.57, 130.29, 130.00, 128.93, 127.72, 127.22, 126.03, 125.32, 123.37, 123.03, 118.25, 117.81, 117.52, 113.72, 113.17, 112.71, 111.99, 71.44, 70.26, 70.11, 70.02, 68.88, 68.53, 54.36, 38.14, 36.57, 36.27, 29.94, 29.64, 14.52, 13.14, 11.77. **ESI-MS**: *m*/*z* $[M + H]^+$ calcd for $C_{46}H_{48}O_7N_7ClBr_3S^+$, 1118.0528; found, 1118.0527; purity: > 99%.

**(*S, Z*)-2-(4-(4-chlorophenyl)-2,3,9-trimethyl-6*H*-thieno[3,2-*f*][1,2,4]triazolo[4,3-*a*][1,4]diazepin-6-yl)-*N*-(1-(2,6-dibromo-4-((2-oxo-6-(trifluoromethyl)indolin-3-ylidene)methyl)phenoxy)-2-oxo-7,10,13-trioxa-3-azahexadecan-16-yl)acetamide (H11)**

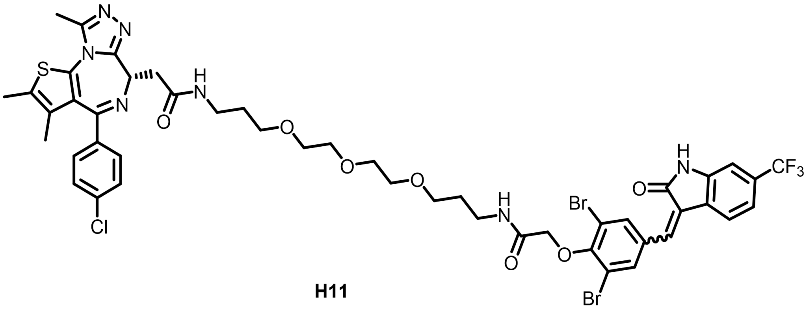

**H11**

The title compound **H11** (yellow solid, 39.7% yield) was synthesized according to the procedures for the preparation of **H6** from **A1** (3,5-dibromo-4-hydroxybenzaldehyde), **C6** (6-(trifluoromethyl)indolin-2-one), tert-butyl 2-bromoacetate and Linker2. **$^1$H NMR** (400 MHz, DMSO-*d*6) *δ* 10.99 (d, *J* = 25.6 Hz, 1H), 8.82 (s, 1H), 8.18 (d, *J* = 16.4 Hz, 2H), 8.00 (d, *J* = 24.9 Hz, 1H), 7.87 (d, *J* = 7.8 Hz, 2/3H), 7.59 – 7.32 (m, 5H), 7.25 (d, *J* = 7.5 Hz, 1/3H), 7.08 (d, *J* = 20.1 Hz, 1H), 4.63 – 4.37 (m, 3H), 3.69 – 3.40 (m, 13H), 3.32 – 3.08 (m, 6H), 2.59 (s, 3H), 2.40 (s, 3H), 1.72 (dd, *J* = 18.4, 6.2 Hz, 4H), 1.61 (s, 3H). **$^{13}$C NMR** (101 MHz, DMSO-*d*6) *δ* 169.93, 167.19, 166.60, 163.48, 155.55, 153.49, 150.24, 141.79, 137.21, 136.84, 136.48, 135.70, 133.93, 133.47, 132.73, 131.12, 130.57, 130.28, 130.00, 128.92, 127.42, 121.11, 118.64, 118.33, 117.54, 106.25, 71.45, 70.25, 70.10, 70.01, 68.87, 68.52, 54.36, 38.12, 36.56, 36.26, 29.93, 29.63, 14.49, 13.12, 11.75. **ESI-MS**: *m/z* [M + H]$^+$ calcd for $C_{47}H_{48}O_7N_7ClBr_2F_3S^+$, 1106.1328; found, 1106.1326; purity: > 99%.

**(*S, Z*)-2-(4-(4-chlorophenyl)-2,3,9-trimethyl-6*H*-thieno[3,2-*f*][1,2,4]triazolo[4,3-*a*][1,4]diazepin-6-yl)-*N*-(1-(4-((5-fluoro-2-oxoindolin-3-ylidene)methyl)phenoxy)-2-oxo-7,10,13-trioxa-3-azahexadecan-16-yl)acetamide (H12)**

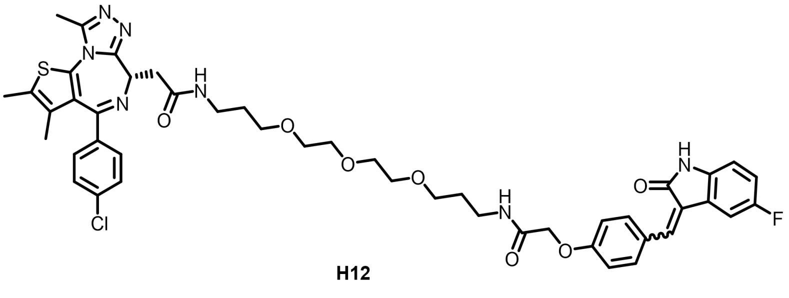

**H12**

The title compound **H12** (yellow solid, 29.4% yield) was synthesized according to the procedures for the preparation of **H6** from **A2** (4-hydroxybenzaldehyde), **C3** (5-fluoroindolin-2-one), tert-butyl 2-bromoacetate and Linker2. **$^1$H NMR** (400 MHz, DMSO-*d*6) *δ* 10.60 (s, 1H), 8.48 (d, *J* = 8.3 Hz, 1/3H), 8.16 (d, *J* = 22.8 Hz, 2H), 7.83 (s, 1/3H), 7.71 (d, *J* = 8.1 Hz, 4/3H), 7.65 (s, 2/3H), 7.61 (d, *J* = 8.5 Hz, 1/3H), 7.48 (d, *J* = 8.2 Hz, 2H), 7.42 (d, *J* = 8.0 Hz, 2H), 7.34 (d, *J* = 9.4 Hz, 2/3H), 7.17 – 6.95 (m, 3H), 6.90 – 6.75 (m, 1H), 4.58 (s, 2H), 4.52 (s, 1H), 3.59 – 3.38 (m, 13H), 3.29 – 3.10 (m, 6H), 2.59 (s, 3H), 2.40 (s, 3H), 1.73 – 1.64 (m, 4H), 1.61 (s, 3H). **$^{13}$C NMR** (101 MHz, DMSO-*d*6) *δ* 169.93, 169.31, 167.86,

167.62, 167.56, 163.49, 160.30, 159.67, 156.46, 155.57, 150.26, 139.50, 138.82, 137.99, 137.22, 136.97, 135.70, 135.03, 132.73, 131.95, 131.14, 130.57, 130.29, 130.02, 128.93, 127.81, 127.29, 126.09, 124.51, 122.49, 116.57, 116.34, 115.58, 114.99, 110.41, 109.70, 70.22, 70.02, 68.64, 68.51, 67.47, 54.36, 38.12, 36.34, 36.26, 29.92, 29.73, 14.50, 13.12, 11.75. **ESI-MS**: *m/z* $[M + H]^+$ calcd for $C_{46}H_{50}O_7N_7ClFS^+$, 898.3160; found, 898.3161; purity: > 99%.

**(*S, Z*)-2-(4-((5-chloro-2-oxoindolin-3-ylidene)methyl)phenoxy)-*N*-(1-(4-(4-chlorophenyl)-2,3,9-trimethyl-6*H*-thieno[3,2-*f*][1,2,4]triazolo[4,3-*a*][1,4]diazepin-6-yl)-2-oxo-7,10,13-trioxa-3-azahexadecan-16-yl)acetamide (H13)**

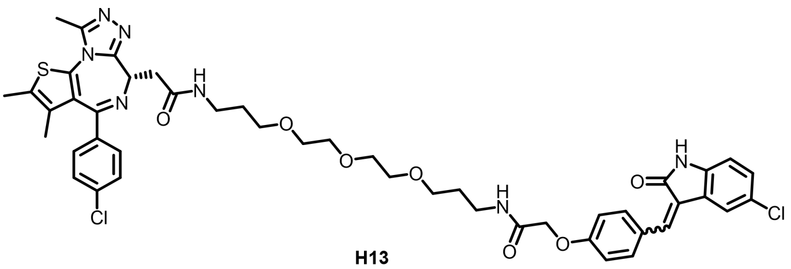


H13

The title compound **H13** (yellow solid, 38.5% yield) was synthesized according to the procedures for the preparation of **H6** from **A2** (4-hydroxybenzaldehyde), **C4** (5-chloroindolin-2-one), tert-butyl 2-bromoacetate and Linker2. **$^1$H NMR** (400 MHz, DMSO-$d_6$) $\delta$ 10.72 (s, 1H), 8.49 (d, $J$ = 8.1 Hz, 1H), 8.17 (d, $J$ = 19.6 Hz, 2H), 7.85 (d, $J$ = 31.6 Hz, 1H), 7.76 – 7.61 (m, 2H), 7.55 (s, 1H), 7.45 (dd, $J$ = 24.1, 7.7 Hz, 4H), 7.24 (dd, $J$ = 27.2, 8.0 Hz, 1H), 7.10 (dd, $J$ = 26.6, 8.0 Hz, 2H), 6.86 (dd, $J$ = 28.2, 8.3 Hz, 1H), 4.55 (d, $J$ = 25.2 Hz, 3H), 3.65 – 3.38 (m, 12H), 3.21 (d, $J$ = 6.8 Hz, 7H), 2.59 (s, 3H), 2.40 (s, 3H), 1.68 (s, 4H), 1.61 (s, 3H). **$^{13}$C NMR** (101 MHz, DMSO-$d_6$) $\delta$ 169.99, 168.94, 167.61, 163.49, 160.34, 159.72, 155.57, 150.80, 150.25, 141.93, 139.38, 138.22, 137.21, 135.70, 135.10, 132.72, 131.96, 131.14, 130.57, 130.29, 130.01, 129.71, 128.92, 127.28, 125.51, 125.37, 123.73, 123.23, 121.95, 119.83, 115.55, 115.01, 111.87, 111.01, 70.22, 70.02, 68.66, 68.51, 67.48, 54.36, 38.13, 36.35, 29.92, 29.73, 14.50, 13.12, 11.75. **ESI-MS**: *m/z* $[M + H]^+$ calcd for $C_{46}H_{50}O_7N_7Cl_2S^+$, 914.2864; found, 914.2867; purity: > 99%.

**(*S, Z*)-2-(4-((5-bromo-2-oxoindolin-3-ylidene)methyl)phenoxy)-*N*-(1-(4-(4-chlorophenyl)-2,3,9-trimethyl-6*H*-thieno[3,2-*f*][1,2,4]triazolo[4,3-*a*][1,4]diazepin-6-yl)-2-oxo-7,10,13-trioxa-3-azahexadecan-16-yl)acetamide (H14)**

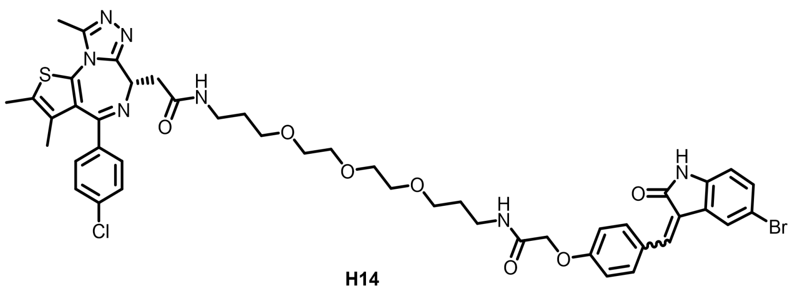

**H14**

The title compound **H14** (yellow solid, 50.0% yield) was synthesized according to the procedures for the preparation of **H6** from **A2** (4-hydroxybenzaldehyde), **C5** (5-bromoindolin-2-one), tert-butyl 2-bromoacetate and Linker2. **$^{1}$H NMR** (400 MHz, DMSO-*d*$_6$) *δ* 10.73 (s, 1H), 8.49 (s, 1H), 8.17 (d, *J* = 18.1 Hz, 2H), 7.91 (d, *J* = 16.4 Hz, 1H), 7.77 – 7.60 (m, 3H), 7.56 – 7.37 (m, 5H), 7.10 (d, *J* = 26.8 Hz, 2H), 6.91 – 6.72 (m, 1H), 4.59 (s, 2H), 4.53 (s, 1H), 3.57 – 3.39 (m, 13H), 3.22 (d, *J* = 5.1 Hz, 6H), 2.60 (d, *J* = 3.8 Hz, 3H), 2.40 (d, *J* = 3.2 Hz, 3H), 1.69 (s, 4H), 1.62 (s, 3H). **$^{13}$C NMR** (101 MHz, DMSO-*d*$_6$) *δ* 169.93, 167.61, 163.49, 160.36, 159.75, 155.56, 150.24, 142.28, 139.09, 138.22, 137.22, 135.71, 135.12, 132.72, 132.49, 131.96, 131.13, 130.57, 130.30, 130.03, 128.93, 127.27, 125.41, 124.66, 122.57, 115.55, 115.01, 113.08, 112.37, 111.54, 70.24, 70.03, 68.66, 54.36, 38.12, 36.35, 29.94, 29.74, 14.51, 13.12, 11.75. **ESI-MS**: *m*/*z* [M + H]$^+$ calcd for $C_{46}H_{50}O_7N_7ClBrS^+$, 960.2340; found, 960.2340; purity: > 99%.

**(*S, Z*)-2-(4-(4-chlorophenyl)-2,3,9-trimethyl-6*H*-thieno[3,2-*f*][1,2,4]triazolo[4,3-*a*][1,4]diazepin-6-yl)-*N*-(2-oxo-1-(4-((2-oxo-6-(trifluoromethyl)indolin-3-ylidene)methyl)phenoxy)-7,10,13-trioxa-3-azahexadecan-16-yl)acetamide (H15)**

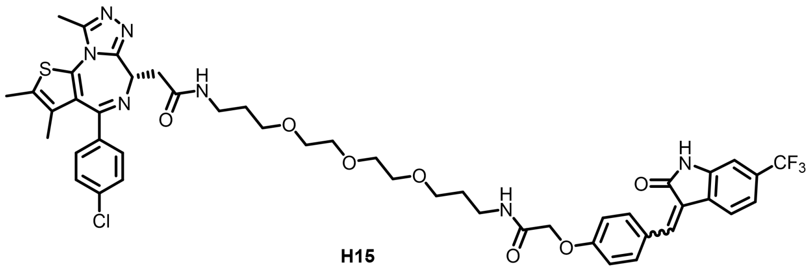

**H15**

The title compound **H15** (yellow solid, 36.7% yield) was synthesized according to the procedures for the preparation of **H6** from **A2** (4-hydroxybenzaldehyde), **C6** (6-(trifluoromethyl)indolin-2-one), tert-butyl 2-bromoacetate and Linker2. **$^{1}$H NMR** (400 MHz, DMSO-*d*$_6$) *δ* 10.88 (d, *J* = 10.9 Hz, 1H), 8.54 (d, *J* = 8.7 Hz, 1H), 8.17 (dd, *J* = 14.9, 8.4 Hz, 2H), 8.01 – 7.85 (m, 1H), 7.82 (d, *J* = 8.0 Hz, 1H), 7.76 (d, *J* = 3.1 Hz, 2H), 7.45 (dd, *J* = 23.6, 8.3 Hz, 4H), 7.34 (d, *J* = 7.6 Hz, 1H), 7.23 (d, *J* = 7.8 Hz, 1H), 7.09 (dd, *J* = 23.4, 8.8 Hz, 3H), 4.59 (s, 2H), 4.55 – 4.47 (m, 1H), 3.57 – 3.40 (m, 13H), 3.23 (dd, *J* = 12.9, 6.8 Hz, 6H), 2.59 (s, 3H), 2.40 (s, 3H), 1.73 – 1.65 (m, 4H), 1.61 (s, 3H). **$^{13}$C NMR** (101 MHz, DMSO-*d*$_6$) *δ* 169.92, 169.04, 168.35, 167.60, 167.55, 163.51, 159.97, 155.59, 150.24, 143.52, 139.78, 137.24, 135.73, 135.42, 132.76, 132.31, 131.15, 130.58, 130.31, 130.00, 128.92, 127.21, 125.51, 125.36, 124.88, 122.94, 118.37, 115.60,

115.05, 106.51, 70.22, 70.04, 68.66, 67.45, 54.35, 38.11, 36.34, 29.91, 29.74, 14.48, 13.10, 11.73. **ESI-MS**: *m/z* $[M + H]^+$ calcd for $C_{47}H_{50}O_7N_7ClF_3S^+$, 948.3128; found, 948.3132; purity: > 99%.

**(*S, Z*)-2-(3-chloro-4-((5-fluoro-2-oxoindolin-3-ylidene)methyl)phenoxy)-*N*-(1-(4-(4-chlorophenyl)-2,3,9-trimethyl-6*H*-thieno[3,2-*f*][1,2,4]triazolo[4,3-*a*][1,4]diazepin-6-yl)-2-oxo-7,10,13-trioxa-3-azahexadecan-16-yl)acetamide (H16)**

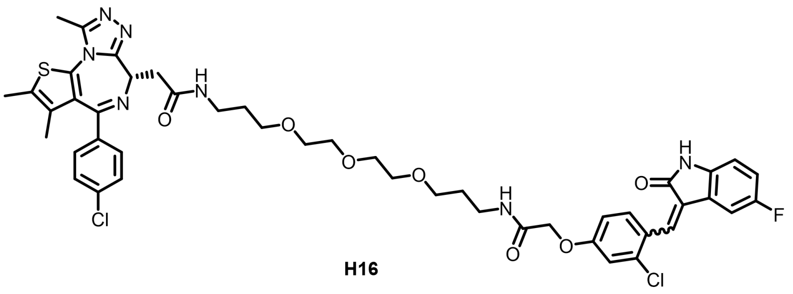

**H16**

The title compound **H16** (yellow solid, 41.2% yield) was synthesized according to the procedures for the preparation of **H6** from **A3** (2-chloro-4-hydroxybenzaldehyde), **C3** (5-fluoroindolin-2-one), tert-butyl 2-bromoacetate and Linker2. **$^1$H NMR** (400 MHz, DMSO-$d_6$) $\delta$ 10.63 (d, $J$ = 40.6 Hz, 1H), 8.17 (d, $J$ = 5.4 Hz, 2H), 7.88 – 7.69 (m, 1H), 7.61 (s, 1H), 7.45 (dd, $J$ = 23.6, 7.0 Hz, 4H), 7.27 (s, 1H), 7.18 – 6.93 (m, 3H), 6.88 (s, 1H), 4.62 (s, 2H), 4.51 (s, 1H), 3.46 (dd, $J$ = 24.5, 10.0 Hz, 13H), 3.21 (s, 6H), 2.59 (s, 3H), 2.40 (s, 3H), 1.67 (d, $J$ = 5.9 Hz, 4H), 1.61 (s, 3H). **$^{13}$C NMR** (101 MHz, DMSO-$d_6$) $\delta$ 169.92, 168.74, 167.33, 163.49, 160.15, 155.57, 150.26, 139.81, 137.21, 135.71, 134.84, 133.45, 132.72, 131.76, 131.15, 130.57, 130.29, 130.02, 128.93, 125.29, 117.25, 117.01, 116.64, 114.73, 70.22, 70.02, 68.59, 68.51, 67.68, 54.35, 38.11, 36.31, 36.26, 29.92, 29.73, 14.50, 13.12, 11.75. **ESI-MS**: *m/z* $[M + H]^+$ calcd for $C_{46}H_{49}O_7N_7Cl_2FS^+$, 932.2770; found, 932.2779; purity: > 99%.

**(*S, Z*)-2-(3-chloro-4-((5-chloro-2-oxoindolin-3-ylidene)methyl)phenoxy)-*N*-(1-(4-(4-chlorophenyl)-2,3,9-trimethyl-6*H*-thieno[3,2-*f*][1,2,4]triazolo[4,3-*a*][1,4]diazepin-6-yl)-2-oxo-7,10,13-trioxa-3-azahexadecan-16-yl)acetamide (H17)**

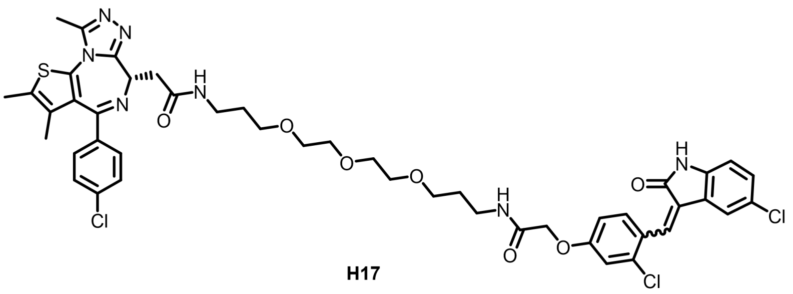

**H17**

The title compound **H17** (yellow solid, 38.9% yield) was synthesized according to the procedures for the preparation of **H6** from **A3** (2-chloro-4-hydroxybenzaldehyde), **C4** (5-chloroindolin-2-one), tert-butyl 2-bromoacetate and Linker2. **$^1$H NMR** (400 MHz, DMSO-$d_6$) $\delta$ 10.75 (d, $J$ = 42.0 Hz, 1H), 8.18 (d, $J$ = 6.5

Hz, 2H), 7.81 (dd, $J$ = 31.0, 19.9 Hz, 2H), 7.61 (s, 1H), 7.45 (dd, $J$ = 24.0, 8.3 Hz, 4H), 7.33 – 7.25 (m, 2H), 7.23 (s, 1H), 7.18 – 7.07 (m, 1H), 6.90 (d, $J$ = 8.3 Hz, 1H), 4.61 (d, $J$ = 11.0 Hz, 2H), 4.51 (t, $J$ = 6.9 Hz, 1H), 3.55 – 3.38 (m, 16H), 3.28 – 3.10 (m, 6H), 2.59 (s, 3H), 2.40 (s, 3H), 1.71 – 1.65 (m, 4H), 1.61 (s, 3H). **$^{13}$C NMR** (101 MHz, DMSO-$d_6$) $\delta$ 169.91, 168.46, 167.31, 163.48, 160.21, 155.57, 150.25, 142.24, 137.21, 135.70, 134.93, 133.66, 132.72, 131.79, 131.14, 130.57, 130.29, 130.02, 128.92, 127.88, 125.52, 125.30, 122.74, 122.34, 116.59, 114.74, 112.08, 70.23, 70.02, 68.60, 68.51, 67.69, 54.36, 38.11, 36.32, 36.25, 29.92, 29.74, 14.50, 13.12, 11.75. **ESI-MS**: $m/z$ $[M + H]^+$ calcd for $C_{46}H_{49}O_7N_7Cl_3S^+$, 950.2445; found, 950.2448; purity: > 99%.

**(*S*, *Z*)-2-(4-((5-bromo-2-oxoindolin-3-ylidene)methyl)-3-chlorophenoxy)-*N*-(1-(4-(4-chlorophenyl)-2,3,9-trimethyl-6*H*-thieno[3,2-*f*][1,2,4]triazolo[4,3-*a*][1,4]diazepin-6-yl)-2-oxo-7,10,13-trioxa-3-azahexadecan-16-yl)acetamide (H18)**

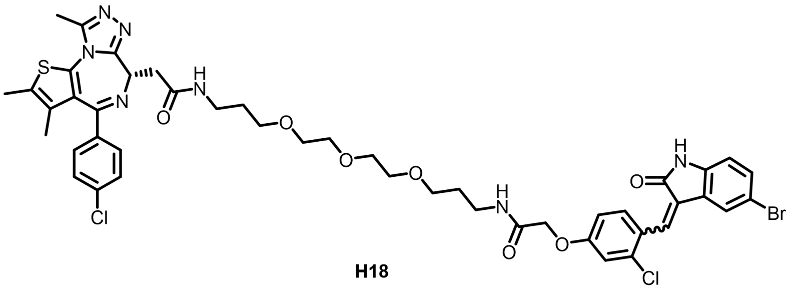


**H18**

The title compound **H18** (yellow solid, 33.9% yield) was synthesized according to the procedures for the preparation of **H6** from **A3** (2-chloro-4-hydroxybenzaldehyde), **C5** (5-bromoindolin-2-one), tert-butyl 2-bromoacetate and Linker2. **$^1$H NMR** (400 MHz, DMSO-$d_6$) $\delta$ 10.77 (d, $J$ = 42.2 Hz, 1H), 8.19 (d, $J$ = 5.4 Hz, 2H), 7.91 (d, $J$ = 17.1 Hz, 1H), 7.78 (d, $J$ = 8.5 Hz, 1H), 7.61 (s, 1H), 7.45 (dd, $J$ = 23.6, 8.1 Hz, 5H), 7.33 (d, $J$ = 37.8 Hz, 2H), 7.14 (d, $J$ = 8.7 Hz, 1H), 6.82 (dd, $J$ = 29.9, 8.1 Hz, 1H), 4.62 (d, $J$ = 10.9 Hz, 2H), 4.52 (t, $J$ = 6.7 Hz, 1H), 3.56 – 3.39 (m, 15H), 3.25 (dd, $J$ = 19.3, 10.7 Hz, 6H), 2.60 (s, 3H), 2.40 (s, 3H), 1.69 (s, 4H), 1.62 (s, 3H). **$^{13}$C NMR** (101 MHz, DMSO-$d_6$) $\delta$ 169.92, 168.33, 167.30, 163.45, 160.22, 155.53, 150.25, 142.60, 137.17, 135.66, 134.96, 133.64, 133.10, 132.72, 131.78, 131.14, 130.57, 130.25, 129.98, 128.92, 127.74, 125.30, 125.05, 124.71, 123.21, 116.57, 114.72, 113.16, 112.58, 70.23, 70.02, 68.61, 67.70, 54.35, 38.11, 36.30, 29.92, 29.72, 14.51, 13.13, 11.72. **ESI-MS**: $m/z$ $[M + H]^+$ calcd for $C_{46}H_{49}O_7N_7Cl_2BrS^+$, 994.1949; found, 994.1948; purity: > 99%.

**(*S*, *Z*)-2-(3-chloro-4-((2-oxo-6-(trifluoromethyl)indolin-3-ylidene)methyl)phenoxy)-*N*-(1-(4-(4-chlorophenyl)-2,3,9-trimethyl-6*H*-thieno[3,2-*f*][1,2,4]triazolo[4,3-*a*][1,4]diazepin-6-yl)-2-oxo-7,10,13-trioxa-3-azahexadecan-16-yl)acetamide (H19)**

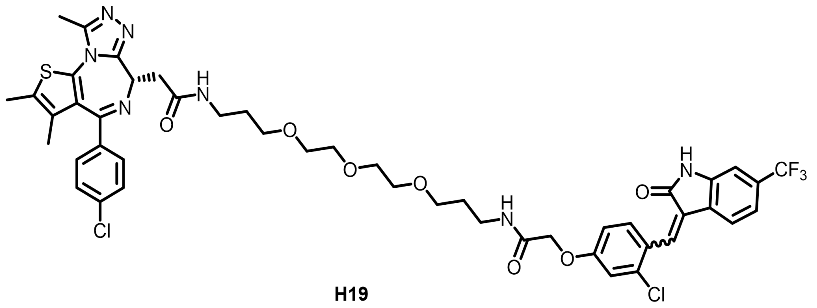

**H19**

The title compound **H19** (yellow solid, 43.6% yield) was synthesized according to the procedures for the preparation of **H6** from **A3** (2-chloro-4-hydroxybenzaldehyde), **C6** (6-(trifluoromethyl)indolin-2-one), tert-butyl 2-bromoacetate and Linker2. **$^1$H NMR** (400 MHz, DMSO-$d_6$) $\delta$ 10.93 (d, $J$ = 26.3 Hz, 1H), 8.19 (d, $J$ = 6.1 Hz, 2H), 7.80 (d, $J$ = 8.6 Hz, 1H), 7.72 (s, 1H), 7.45 (dd, $J$ = 23.8, 8.5 Hz, 5H), 7.33 (d, $J$ = 8.4 Hz, 1H), 7.28 (s, 1H), 7.20 (d, $J$ = 8.1 Hz, 1H), 7.14 – 6.96 (m, 2H), 4.63 (s, 2H), 4.52 (t, $J$ = 6.9 Hz, 1H), 3.46 (ddd, $J$ = 18.0, 12.0, 7.9 Hz, 16H), 3.21 (dt, $J$ = 23.5, 12.7 Hz, 6H), 2.60 (s, 3H), 2.40 (s, 3H), 1.69 (d, $J$ = 6.1 Hz, 4H), 1.62 (s, 3H). **$^{13}$C NMR** (101 MHz, DMSO-$d_6$) $\delta$ 169.93, 168.47, 167.28, 163.49, 160.35, 155.57, 150.26, 143.88, 137.21, 135.70, 135.11, 132.72, 131.96, 131.14, 130.56, 130.29, 130.02, 128.92, 127.42, 125.21, 124.90, 123.34, 118.58, 116.59, 114.86, 70.23, 70.02, 68.60, 68.51, 67.68, 54.35, 38.12, 36.31, 36.26, 29.91, 29.74, 14.49, 13.11, 11.74. **ESI-MS**: *m/z* $[M + H]^+$ calcd for $C_{47}H_{48}O_7N_7Cl_2F_3S^+$, 982.2719; found, 982.2745; purity: > 99%.

**(*S, Z*)-2-(2-bromo-4-((5-fluoro-2-oxoindolin-3-ylidene)methyl)phenoxy)-*N*-(1-(4-(4-chlorophenyl)-2,3,9-trimethyl-6*H*-thieno[3,2-*f*][1,2,4]triazolo[4,3-*a*][1,4]diazepin-6-yl)-2-oxo-7,10,13-trioxa-3-azahexadecan-16-yl)acetamide (H20)**

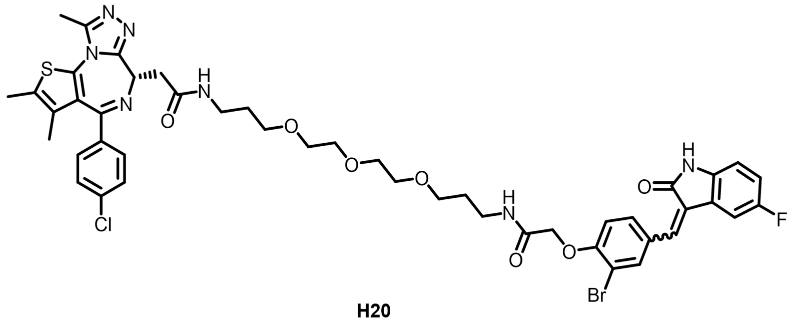

**H20**

The title compound **H20** (yellow solid, 52.3% yield) was synthesized according to the procedures for the preparation of **H6** from **A4** (3-bromo-4-hydroxybenzaldehyde), **C3** (5-fluoroindolin-2-one), tert-butyl 2-bromoacetate and Linker2. **$^1$H NMR** (400 MHz, DMSO-$d_6$) $\delta$ 10.64 (d, $J$ = 10.7 Hz, 1H), 8.19 (s, 1H), 7.96 (s, 2H), 7.87 – 7.69 (m, 1H), 7.60 (d, $J$ = 15.8 Hz, 1H), 7.45 (dd, $J$ = 24.4, 7.5 Hz, 4H), 7.25 (d, $J$ = 8.2 Hz, 1H), 7.19 – 6.95 (m, 2H), 6.84 (d, $J$ = 26.0 Hz, 1H), 4.71 (s, 2H), 4.52 (s, 1H), 3.46 (dd, $J$ = 25.3, 8.2 Hz, 13H), 3.23 (s, 6H), 2.59 (s, 3H), 2.40 (s, 3H), 1.68 (s, 4H), 1.62 (s, 3H). **$^{13}$C NMR** (101 MHz, DMSO-$d_6$) $\delta$ 169.91, 169.02, 167.78, 167.14, 167.06, 166.01, 163.48, 155.90, 155.57, 150.25, 139.75, 137.21, 136.32,

135.70, 134.66, 134.36, 132.73, 131.14, 130.73, 130.57, 130.28, 130.02, 128.92, 128.80, 122.23, 116.75, 114.35, 111.75, 70.23, 70.04, 70.01, 68.55, 68.51, 68.32, 55.39, 54.36, 38.11, 36.37, 36.25, 29.92, 29.65, 14.50, 13.12, 11.75. **ESI-MS**: *m*/*z* $[M + H]^+$ calcd for $C_{46}H_{49}O_7N_7ClBrFS^+$, 978.2244; found, 978.2243; purity: > 99%.

**(*S, Z*)-2-(2-bromo-4-((5-chloro-2-oxoindolin-3-ylidene)methyl)phenoxy)-*N*-(1-(4-(4-chlorophenyl)-2,3,9-trimethyl-6*H*-thieno[3,2-*f*][1,2,4]triazolo[4,3-*a*][1,4]diazepin-6-yl)-2-oxo-7,10,13-trioxa-3-azahexadecan-16-yl)acetamide (H21)**

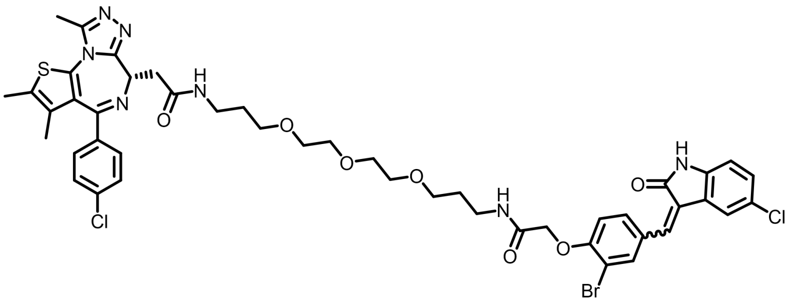

**H21**

The title compound **H21** (yellow solid, 43.6% yield) was synthesized according to the procedures for the preparation of **H6** from **A4** (3-bromo-4-hydroxybenzaldehyde), **C4** (5-chloroindolin-2-one), tert-butyl 2-bromoacetate and Linker2. **$^1$H NMR** (400 MHz, DMSO-*d*$_6$) $\delta$ 10.76 (d, *J* = 9.0 Hz, 1H), 8.19 (s, 1H), 8.06 – 7.91 (m, 2H), 7.84 (d, *J* = 30.1 Hz, 1H), 7.73 (d, *J* = 8.4 Hz, 1H), 7.63 (s, 1H), 7.45 (dd, *J* = 24.3, 8.1 Hz, 5H), 7.26 (dd, *J* = 25.1, 8.3 Hz, 1H), 7.13 (dd, *J* = 17.8, 8.5 Hz, 1H), 6.87 (dd, *J* = 25.0, 8.2 Hz, 1H), 4.72 (s, 2H), 4.52 (t, *J* = 6.8 Hz, 1H), 3.46 (dd, *J* = 25.5, 8.3 Hz, 13H), 3.31 – 3.06 (m, 6H), 2.59 (s, 3H), 2.40 (s, 3H), 1.68 (s, 4H), 1.61 (s, 3H). **$^{13}$C NMR** (101 MHz, DMSO-*d*$_6$) $\delta$ 169.91, 168.75, 167.53, 167.13, 163.48, 155.96, 155.57, 150.25, 142.16, 139.64, 137.41, 137.21, 137.09, 136.53, 135.70, 134.64, 134.45, 132.73, 131.13, 130.88, 130.57, 130.29, 130.01, 129.07, 128.92, 128.76, 126.66, 125.89, 125.41, 124.99, 122.96, 122.08, 114.30, 113.74, 112.00, 111.73, 111.20, 70.23, 70.05, 68.56, 68.52, 68.33, 55.39, 54.36, 38.12, 36.39, 36.26, 29.92, 29.66, 14.51, 13.13, 11.76. **ESI-MS**: *m*/*z* $[M + H]^+$ calcd for $C_{46}H_{49}O_7N_7Cl_2BrS^+$, 994.1949; found, 994.1952; purity: > 99%.

**(*S, Z*)-2-(2-bromo-4-((5-bromo-2-oxoindolin-3-ylidene)methyl)phenoxy)-*N*-(1-(4-(4-chlorophenyl)-2,3,9-trimethyl-6*H*-thieno[3,2-*f*][1,2,4]triazolo[4,3-*a*][1,4]diazepin-6-yl)-2-oxo-7,10,13-trioxa-3-azahexadecan-16-yl)acetamide (H22)**

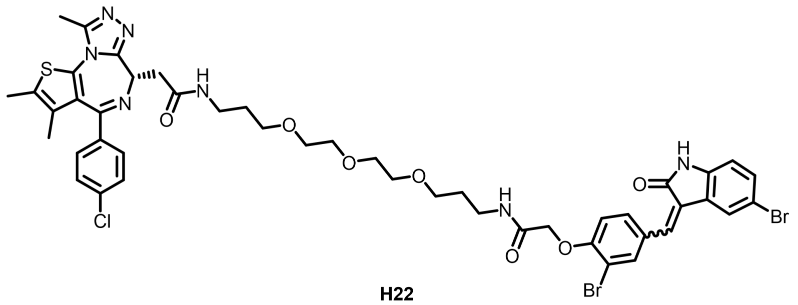

**H22**

The title compound **H22** (yellow solid, 69.3% yield) was synthesized according to the procedures for the preparation of **H6** from **A4** (3-bromo-4-hydroxybenzaldehyde), **C5** (5-bromoindolin-2-one), tert-butyl 2-bromoacetate and Linker2. **$^{1}$H NMR** (400 MHz, DMSO-$d_6$) $\delta$ 10.80 (d, $J$ = 3.3 Hz, 1H), 9.01 (s, 1H), 8.20 (s, 1H), 8.00 (d, $J$ = 8.4 Hz, 1H), 7.95 – 7.78 (m, 2H), 7.63 (d, $J$ = 7.1 Hz, 2H), 7.54 – 7.25 (m, 7H), 7.13 (dd, $J$ = 17.1, 8.6 Hz, 1H), 6.83 (dd, $J$ = 26.1, 8.2 Hz, 1H), 4.72 (s, 2H), 4.51 (d, $J$ = 7.0 Hz, 1H), 3.64 – 3.45 (m, 13H), 3.19 (dd, $J$ = 17.6, 10.1 Hz, 6H), 2.59 (s, 3H), 2.40 (s, 3H), 1.68 (s, 4H), 1.61 (s, 3H). **$^{13}$C NMR** (101 MHz, DMSO-$d_6$) $\delta$ 169.95, 167.40, 167.15, 167.06, 163.48, 156.51, 155.56, 150.24, 144.26, 143.30, 140.04, 137.42, 137.20, 137.09, 136.48, 135.71, 134.61, 134.49, 132.84, 132.71, 131.34, 131.13, 130.59, 130.27, 129.99, 129.08, 128.95, 127.81, 126.70, 126.66, 124.83, 124.45, 122.85, 119.31, 114.38, 114.28, 113.75, 112.57, 111.72, 110.46, 70.24, 70.03, 68.54, 68.30, 68.22, 68.20, 54.36, 53.86, 42.16, 38.12, 36.39, 36.18, 29.92, 29.65, 14.49, 13.12, 12.85, 11.75. **ESI-MS**: $m/z$ [M + H]$^+$ calcd for $C_{46}H_{49}O_7N_7ClBr_2S^+$, 1038.1444; found, 1038.1455; purity: > 99%.

**(*S*, *Z*)-2-(2-bromo-4-((2-oxo-6-(trifluoromethyl)indolin-3-ylidene)methyl)phenoxy)-*N*-(1-(4-(4-chloro-phenyl)-2,3,9-trimethyl-6*H*-thieno[3,2-*f*][1,2,4]triazolo[4,3-*a*][1,4]diazepin-6-yl)-2-oxo-7,10,13-trioxa-3 -azahexadecan-16-yl)acetamide (H23)**

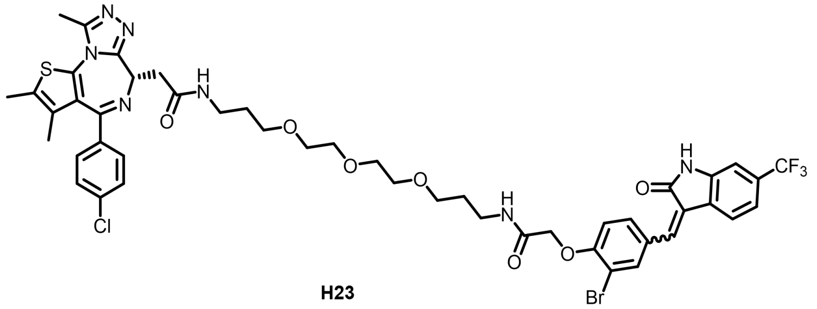

**H23**

The title compound **H23** (yellow solid, 36.7% yield) was synthesized according to the procedures for the preparation of **H6** from **A4** (3-bromo-4-hydroxybenzaldehyde), **C6** (6-(trifluoromethyl)indolin-2-one), tert-butyl 2-bromoacetate and Linker2. **$^{1}$H NMR** (400 MHz, DMSO-$d_6$) $\delta$ 10.92 (d, $J$ = 17.4 Hz, 1H), 8.19 (s, 1H), 7.99 (d, $J$ = 14.8 Hz, 2H), 7.83 (dd, $J$ = 45.1, 8.1 Hz, 1H), 7.71 (d, $J$ = 7.6 Hz, 1H), 7.45 (dd, $J$ = 24.5, 8.2 Hz, 4H), 7.30 (dd, $J$ = 48.2, 8.0 Hz, 1H), 7.11 (dd, $J$ = 24.6, 14.1 Hz, 2H), 4.72 (s, 2H), 4.52 (t, $J$ = 6.8 Hz, 1H), 3.47 (dd, $J$ = 20.0, 9.1 Hz, 12H), 3.31 – 3.07 (m, 6H), 2.59 (s, 3H), 2.40 (s, 3H), 1.69 (d, $J$ = 5.0 Hz,

4H), 1.61 (s, 3H). **$^{13}$C NMR** (101 MHz, DMSO-*d*$_6$) *δ* 169.93, 168.76, 167.48, 167.02, 163.48, 156.77, 156.15, 155.56, 150.25, 143.79, 141.18, 138.83, 138.10, 137.33, 137.21, 135.69, 134.91, 134.72, 132.73, 131.12, 131.00, 130.56, 130.28, 130.00, 129.45, 128.92, 128.69, 125.14, 124.51, 123.68, 123.33, 122.93, 120.52, 118.40, 114.35, 113.74, 111.83, 111.24, 105.98, 70.22, 70.05, 70.01, 68.55, 68.51, 68.23, 55.38, 54.36, 38.12, 36.37, 36.25, 29.92, 29.64, 14.48, 13.11, 11.75. **ESI-MS**: *m*/*z* $[M + H]^+$ calcd for $C_{47}H_{49}O_7N_7ClBrF_3S^+$, 1028.2212; found, 1028.2210; purity: > 99%.

**(*S*, *Z*)-2-(4-bromo-2-((5-fluoro-2-oxoindolin-3-ylidene)methyl)phenoxy)-*N*-(1-(4-(4-chlorophenyl)-2,3,9-trimethyl-6*H*-thieno[3,2-*f*][1,2,4]triazolo[4,3-*a*][1,4]diazepin-6-yl)-2-oxo-7,10,13-trioxa-3-azahexadecan-16-yl)acetamide (H24)**

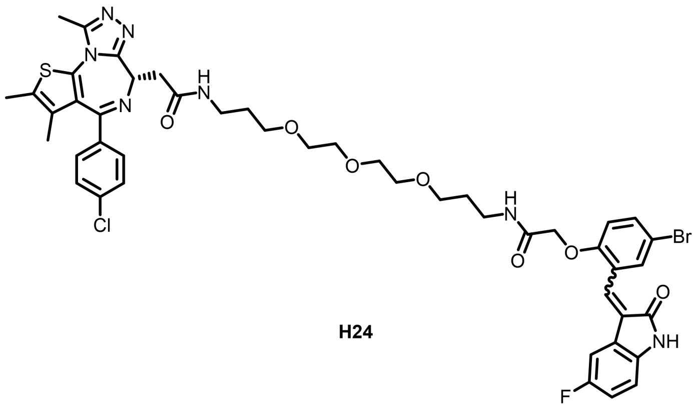

**H24**

The title compound **H24** (yellow solid, 38.9% yield) was synthesized according to the procedures for the preparation of **H6** from **A5** (5-bromo-2-hydroxybenzaldehyde), **C3** (5-fluoroindolin-2-one), tert-butyl 2-bromoacetate and Linker2. **$^1$H NMR** (400 MHz, DMSO-*d*$_6$) *δ* 10.66 (s, 1H), 8.19 (s, 1H), 8.00 (s, 1H), 7.80 (s, 1H), 7.75 (s, 1H), 7.65 (d, *J* = 8.6 Hz, 1H), 7.45 (dd, *J* = 24.0, 8.1 Hz, 4H), 7.10 (t, *J* = 8.3 Hz, 1H), 7.01 (d, *J* = 8.3 Hz, 2H), 6.95 – 6.79 (m, 1H), 4.62 (s, 2H), 4.52 (t, *J* = 6.8 Hz, 1H), 3.47 (dd, *J* = 18.2, 5.5 Hz, 13H), 3.28 – 3.11 (m, 6H), 2.60 (s, 3H), 2.40 (s, 3H), 1.71 – 1.57 (m, 7H). **$^{13}$C NMR** (101 MHz, DMSO-*d*$_6$) *δ* 169.90, 169.20, 168.74, 168.11, 167.41, 163.50, 156.01, 155.57, 150.30, 139.85, 137.20, 135.74, 134.31, 132.77, 132.48, 132.10, 131.18, 130.59, 130.28, 130.02, 128.92, 125.75, 117.19, 116.94, 115.52, 112.63, 112.58, 111.47, 109.91, 109.64, 70.21, 70.00, 68.49, 68.06, 54.36, 38.14, 36.24, 29.91, 29.67, 14.50, 13.17, 11.75. **ESI-MS**: *m*/*z* $[M + H]^+$ calcd for $C_{46}H_{49}O_7N_7ClBrFS^+$, 978.2246; found, 978.2248; purity: > 99%.

**(*S*, *Z*)-2-(4-bromo-2-((5-chloro-2-oxoindolin-3-ylidene)methyl)phenoxy)-*N*-(1-(4-(4-chlorophenyl)-2,3,9-trimethyl-6*H*-thieno[3,2-*f*][1,2,4]triazolo[4,3-*a*][1,4]diazepin-6-yl)-2-oxo-7,10,13-trioxa-3-azahexadecan-16-yl)acetamide (H25)**

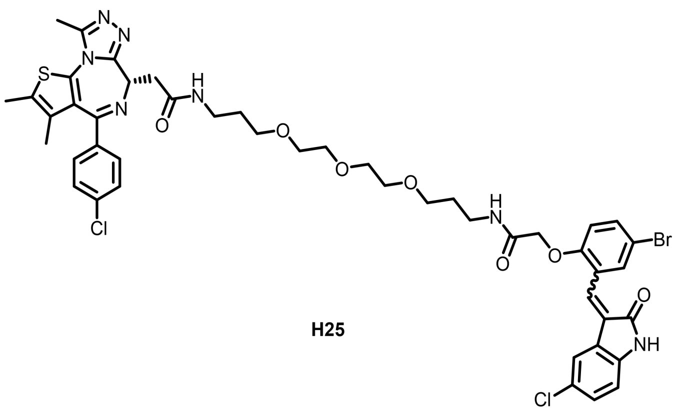

**H25**

The title compound **H25** (yellow solid, 58.4% yield) was synthesized according to the procedures for the preparation of **H6** from **A5** (5-bromo-2-hydroxybenzaldehyde), **C4** (5-chloroindolin-2-one), tert-butyl 2-bromoacetate and Linker2. **$^1$H NMR** (400 MHz, DMSO-$d_6$) $\delta$ 10.76 (d, $J$ = 9.7 Hz, 1H), 8.19 (s, 1H), 8.09 (d, $J$ = 16.8 Hz, 1H), 8.01 (s, 1H), 7.82 (s, 1H), 7.76 (s, 1H), 7.65 (d, $J$ = 8.6 Hz, 1H), 7.45 (dd, $J$ = 23.3, 7.9 Hz, 4H), 7.28 (d, $J$ = 14.2 Hz, 2H), 6.99 (dd, $J$ = 16.9, 8.9 Hz, 1H), 6.87 (dd, $J$ = 21.7, 8.1 Hz, 1H), 4.62 (s, 2H), 4.52 (t, $J$ = 6.3 Hz, 1H), 3.47 (dd, $J$ = 17.5, 5.7 Hz, 12H), 3.29 – 3.07 (m, 6H), 2.60 (s, 3H), 2.40 (s, 3H), 1.65 (d, $J$ = 23.6 Hz, 7H). **$^{13}$C NMR** (101 MHz, DMSO-$d_6$) $\delta$ 174.71, 169.95, 168.51, 167.36, 163.46, 156.09, 155.59, 150.29, 142.28, 137.19, 135.72, 134.38, 132.70, 132.23, 131.65, 131.13, 130.57, 130.27, 130.01, 128.90, 128.22, 125.66, 125.41, 122.38, 120.58, 115.48, 112.60, 112.01, 70.20, 70.00, 68.49, 36.24, 29.95, 29.67, 14.53, 13.16, 11.80. **ESI-MS**: $m/z$ [M + H]$^+$ calcd for $C_{46}H_{49}O_7N_7Cl_2BrS^+$, 994.1949; found, 994.1950; purity: > 99%.

**(*S, Z*)-2-(4-bromo-2-((5-bromo-2-oxoindolin-3-ylidene)methyl)phenoxy)-*N*-(1-(4-(4-chlorophenyl)-2,3,9-trimethyl-6*H*-thieno[3,2-*f*][1,2,4]triazolo[4,3-*a*][1,4]diazepin-6-yl)-2-oxo-7,10,13-trioxa-3-azahexadecan-16-yl)acetamide (H26)**

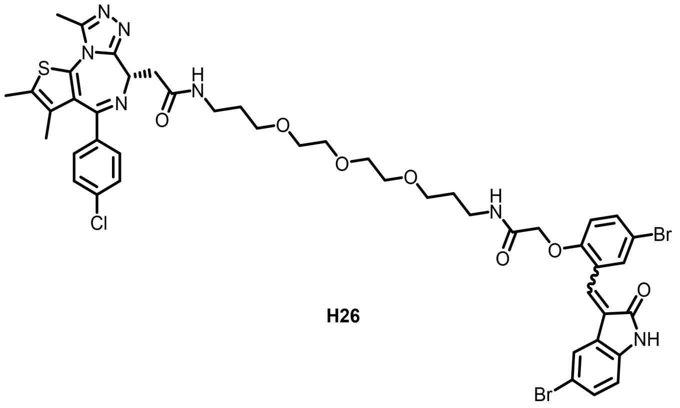

**H26**

The title compound **H26** (yellow solid, 46.9% yield) was synthesized according to the procedures for the preparation of **H6** from **A5** (5-bromo-2-hydroxybenzaldehyde), **C5** (5-bromoindolin-2-one), tert-butyl 2-bromoacetate and Linker2. **$^1$H NMR** (400 MHz, DMSO-$d_6$) $\delta$ 10.76 (d, $J$ = 9.1 Hz, 1H), 8.19 (s, 1H), 8.00 (s, 1H), 7.82 (s, 1H), 7.75 (s, 1H), 7.65 (d, $J$ = 8.8 Hz, 1H), 7.54 – 7.33 (m, 6H), 6.99 (dd, $J$ = 16.1, 8.9 Hz,

1H), 6.93 – 6.73 (m, 1H), 4.62 (s, 2H), 4.51 (t, $J$ = 6.7 Hz, 1H), 3.62 – 3.38 (m, 12H), 3.20 (ddd, $J$ = 21.9, 18.1, 10.6 Hz, 6H), 2.59 (s, 3H), 2.40 (s, 3H). **$^{13}$C NMR** (101 MHz, DMSO-$d_6$) $\delta$ 169.91, 168.34, 167.48, 167.33, 167.09, 163.47, 156.29, 156.06, 155.57, 150.25, 142.60, 137.20, 135.69, 134.39, 134.16, 132.99, 132.72, 132.62, 132.22, 131.64, 131.14, 130.57, 130.29, 130.02, 128.92, 128.10, 127.16, 126.98, 125.65, 125.19, 123.44, 115.47, 113.08, 112.56, 70.21, 70.01, 68.51, 68.04, 54.35, 38.11, 36.25, 29.92, 29.64, 14.51, 13.13, 11.76. **ESI-MS**: *m*/*z* [M + H]$^+$ calcd for $C_{46}H_{49}O_7N_7ClBr_2S^+$, 1038.1444; found, 1038.1451; purity: > 99%.

**(*S*, *Z*)-2-(4-bromo-2-((2-oxo-6-(trifluoromethyl)indolin-3-ylidene)methyl)phenoxy)-*N*-(1-(4-(4-chloro phenyl)-2,3,9-trimethyl-6*H*-thieno[3,2-*f*][1,2,4]triazolo[4,3-*a*][1,4]diazepin-6-yl)-2-oxo-7,10,13-trioxa-3 -azahexadecan-16-yl)acetamide (H27)**

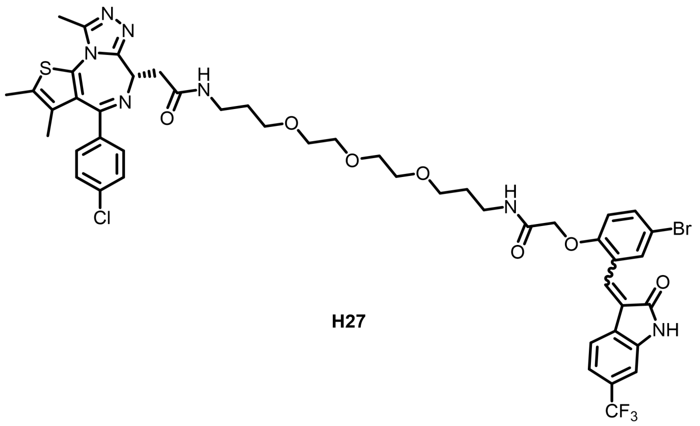


H27

The title compound **H27** (yellow solid, 29.8% yield) was synthesized according to the procedures for the preparation of **H6** from **A5** (5-bromo-2-hydroxybenzaldehyde), **C6** (6-(trifluoromethyl)indolin-2-one), tert-butyl 2-bromoacetate and Linker2. **$^1$H NMR** (400 MHz, DMSO-$d_6$) $\delta$ 10.91 (s, 1H), 8.18 (s, 1H), 7.98 (s, 1H), 7.83 (d, $J$ = 11.1 Hz, 2H), 7.74 – 7.56 (m, 1H), 7.56 – 7.32 (m, 5H), 7.25 (d, $J$ = 7.5 Hz, 1H), 7.14 – 6.92 (m, 2H), 4.63 (s, 2H), 4.51 (s, 1H), 3.47 (d, $J$ = 22.6 Hz, 13H), 3.27 – 3.07 (m, 6H), 2.59 (s, 3H), 2.40 (s, 3H), 1.65 (d, $J$ = 23.5 Hz, 7H). **$^{13}$C NMR** (101 MHz, DMSO-$d_6$) $\delta$ 169.92, 168.50, 167.34, 163.48, 161.74, 156.47, 156.06, 155.57, 153.36, 150.27, 144.75, 143.88, 137.89, 137.21, 135.70, 135.01, 134.57, 134.12, 133.50, 132.72, 132.28, 131.15, 130.57, 130.29, 130.02, 128.92, 127.86, 127.21, 126.64, 125.66, 125.14, 123.29, 120.83, 118.55, 115.48, 112.76, 106.66, 70.20, 70.00, 68.51, 68.03, 54.35, 38.11, 36.25, 29.91, 29.62, 14.50, 13.12, 11.75. **ESI-MS**: *m*/*z* [M + H]$^+$ calcd for $C_{47}H_{49}O_7N_7ClBrF_3S^+$, 1028.2212; found, 1028.2217; purity: > 99%.

**(*S*, *Z*)-2-(4-(4-chlorophenyl)-2,3,9-trimethyl-6*H*-thieno[3,2-*f*][1,2,4]triazolo[4,3-*a*][1,4]diazepin-6-yl)-*N*-(1-(2,6-difluoro-4-((2-oxo-6-(trifluoromethyl)indolin-3-ylidene)methyl)phenoxy)-2-oxo-7,10,13-trioxa-3 -azahexadecan-16-yl)acetamide (H28)**

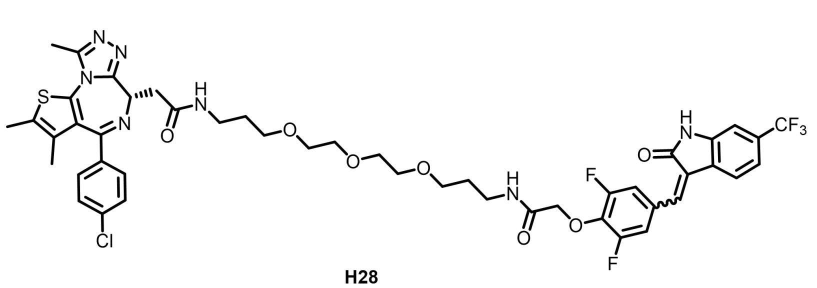

**H28**

The title compound **H28** (yellow solid, 22.3% yield) was synthesized according to the procedures for the preparation of **H6** from **A6** (3,5-difluoro-4-hydroxybenzaldehyde), **C6** (6-(trifluoromethyl)-2-one), tert-butyl 2-bromoacetate and Linker2. **$^1$H NMR** (400 MHz, DMSO-$d_6$) $\delta$ 10.98 (d, $J$ = 36.3 Hz, 1H), 8.35 (d, $J$ = 10.2 Hz, 1H), 8.20 (s, 1H), 8.12 (s, 1H), 7.95 (s, 1H), 7.87 (d, $J$ = 7.8 Hz, 1H), 7.69 (d, $J$ = 9.9 Hz, 1H), 7.56 (d, $J$ = 8.8 Hz, 1H), 7.48 (d, $J$ = 8.3 Hz, 2H), 7.40 (dd, $J$ = 15.1, 8.2 Hz, 3H), 7.08 (d, $J$ = 17.7 Hz, 1H), 4.70 (d, $J$ = 8.1 Hz, 2H), 4.51 (t, $J$ = 6.9 Hz, 1H), 3.46 (dd, $J$ = 26.4, 8.3 Hz, 12H), 3.30 – 3.04 (m, 6H), 2.59 (s, 3H), 2.40 (s, 3H), 1.66 (dd, $J$ = 18.7, 12.6 Hz, 7H). **$^{13}$C NMR** (101 MHz, DMSO-$d_6$) $\delta$ 169.92, 167.29, 163.48, 155.56, 152.62, 150.25, 144.05, 141.58, 137.75, 137.21, 135.70, 132.73, 131.13, 130.57, 130.28, 130.01, 128.92, 126.57, 120.90, 117.05, 116.80, 70.22, 70.01, 68.58, 68.51, 54.36, 38.11, 36.29, 29.92, 29.67, 14.48, 13.11, 11.74. **ESI-MS**: *m/z* [M + H]$^+$ calcd for $C_{47}H_{48}O_7N_7ClF_5S^+$, 984.2950; found, 984.2942; purity: > 99%.

**Tert-butyl (Z)-2-(4-bromo-2-((2-oxo-6-(trifluoromethyl)indolin-3-ylidene)methyl)phenoxy)acetate (D27)**

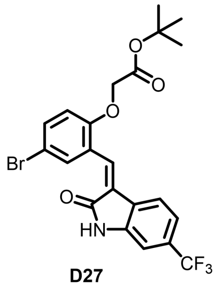

**D27**

The title compound **D27** (yellow solid, 56.8% yield) was synthesized according to the procedures for the preparation of **D1** from **A5** (5-bromo-2-hydroxybenzaldehyde), **C6** (6-(trifluoromethyl)indolin-2-one) and tert-butyl 2-bromoacetate. **$^1$H NMR** (400 MHz, DMSO-$d_6$) $\delta$ 10.92 (s, 1H), 7.82 (d, J = 2.2 Hz, 1H), 7.75 (s, 1H), 7.66 (dd, J = 8.9, 2.2 Hz, 1H), 7.46 (d, J = 8.0 Hz, 1H), 7.25 (d, J = 8.0 Hz, 1H), 7.12 – 7.06 (m, 2H), 4.81 (s, 2H), 1.38 (s, 9H). **$^{13}$C NMR** (101 MHz, DMSO-$d_6$) $\delta$ 168.49, 167.68, 155.81, 143.85, 134.58, 133.52, 132.27, 128.00, 125.52, 125.15, 123.36, 118.61, 115.56, 114.97, 112.84, 112.43, 106.74, 99.99, 82.21, 66.03, 28.07. **ESI-MS**: *m/z* [M + H]$^+$ calcd for $C_{22}H_{20}O_4NBrF_3^+$, 498.0522; found, 498.0529; purity: 97%.
