## Supplementary material for "Discovery of Alkenyl Oxindole as a Novel PROTAC Moiety for Targeted Protein Degradation via CRL4^DCAF11^ Recruitment": NMR Spectra of compounds H1-H28 and D27

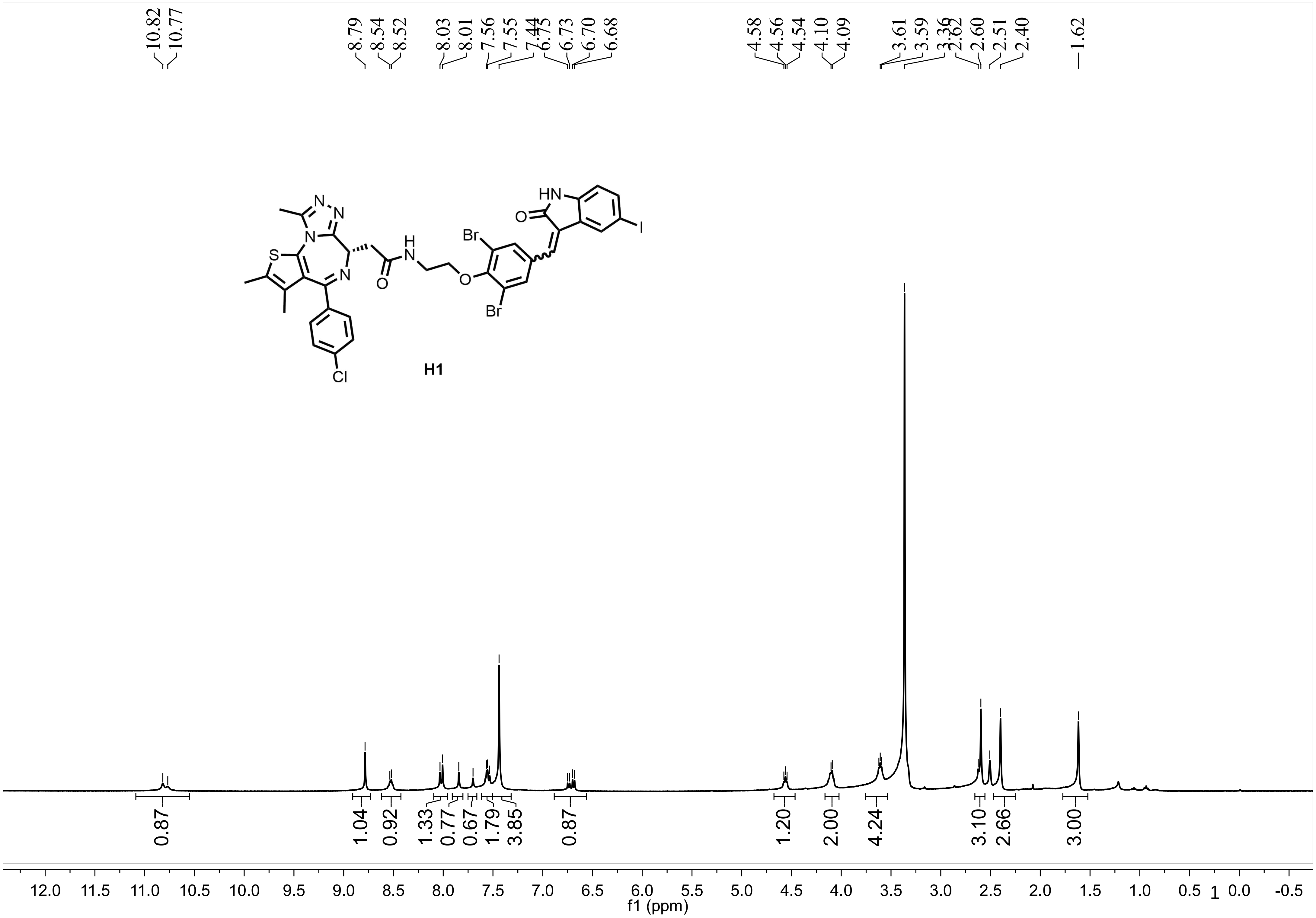

10.82
10.77
8.79
8.54
8.52
8.03
8.01
7.56
7.55
7.44
6.75
6.73
6.70
6.68
4.58
4.56
4.54
4.10
4.09
3.61
3.59
3.36
2.62
2.60
2.51
2.40
1.62
H1
0.87
1.04
0.92
1.33
0.77
0.67
1.79
3.85
0.87
1.20
2.00
4.24
3.10
2.66
3.00
f1 (ppm)


1

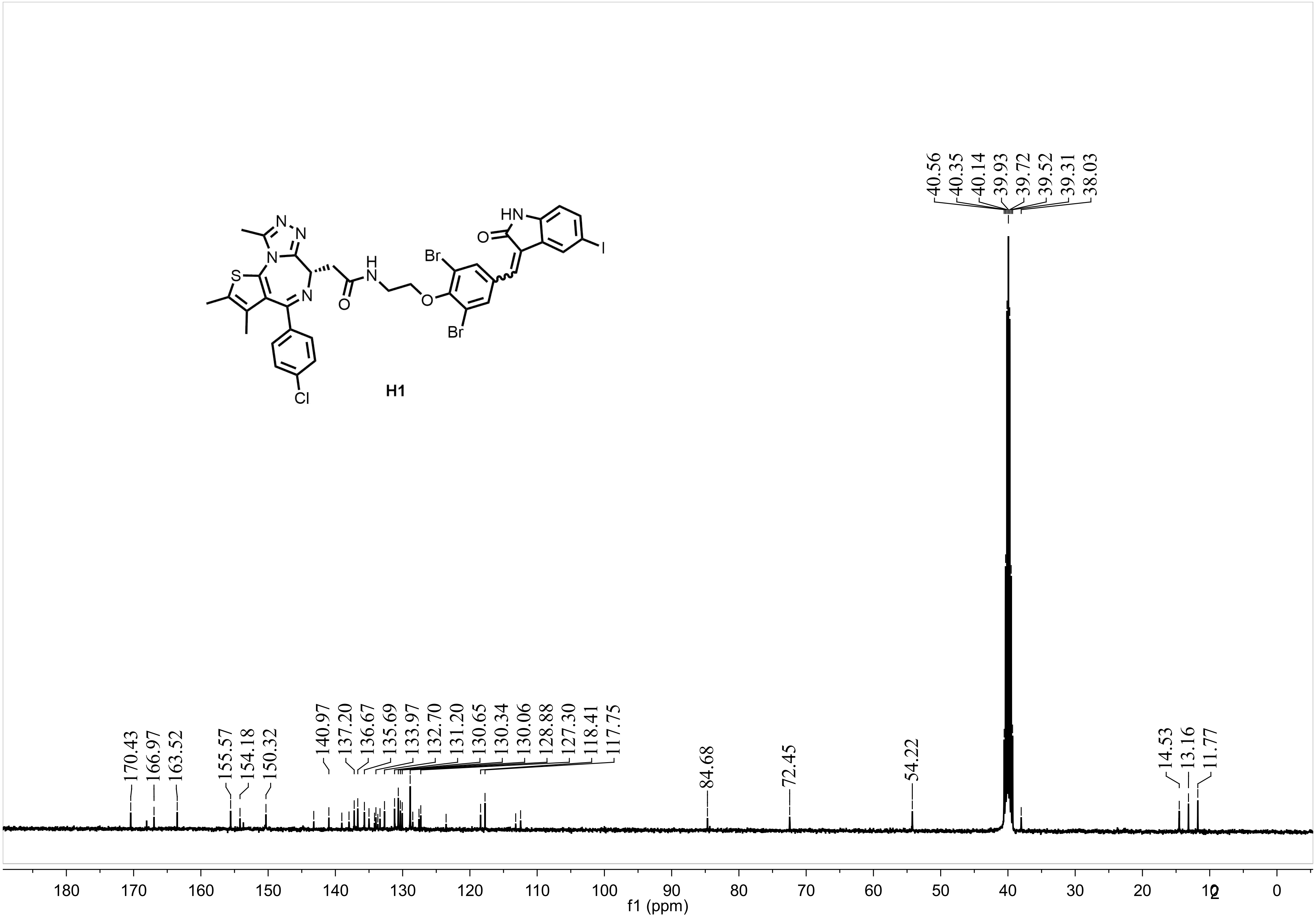
H1
Br
Br
HN
O
O
N
N
N
S
Cl
I
H
170.43
166.97
163.52
155.57
154.18
150.32
140.97
137.20
136.67
135.69
133.97
132.70
131.20
130.65
130.34
130.06
128.88
127.30
118.41
117.75
84.68
72.45
54.22
40.56
40.35
40.14
39.93
39.72
39.52
39.31
38.03
14.53
13.16
11.77
180
170
160
150
140
130
120
110
100
90
80
70
60
50
40
30
20
10
0
f1 (ppm)

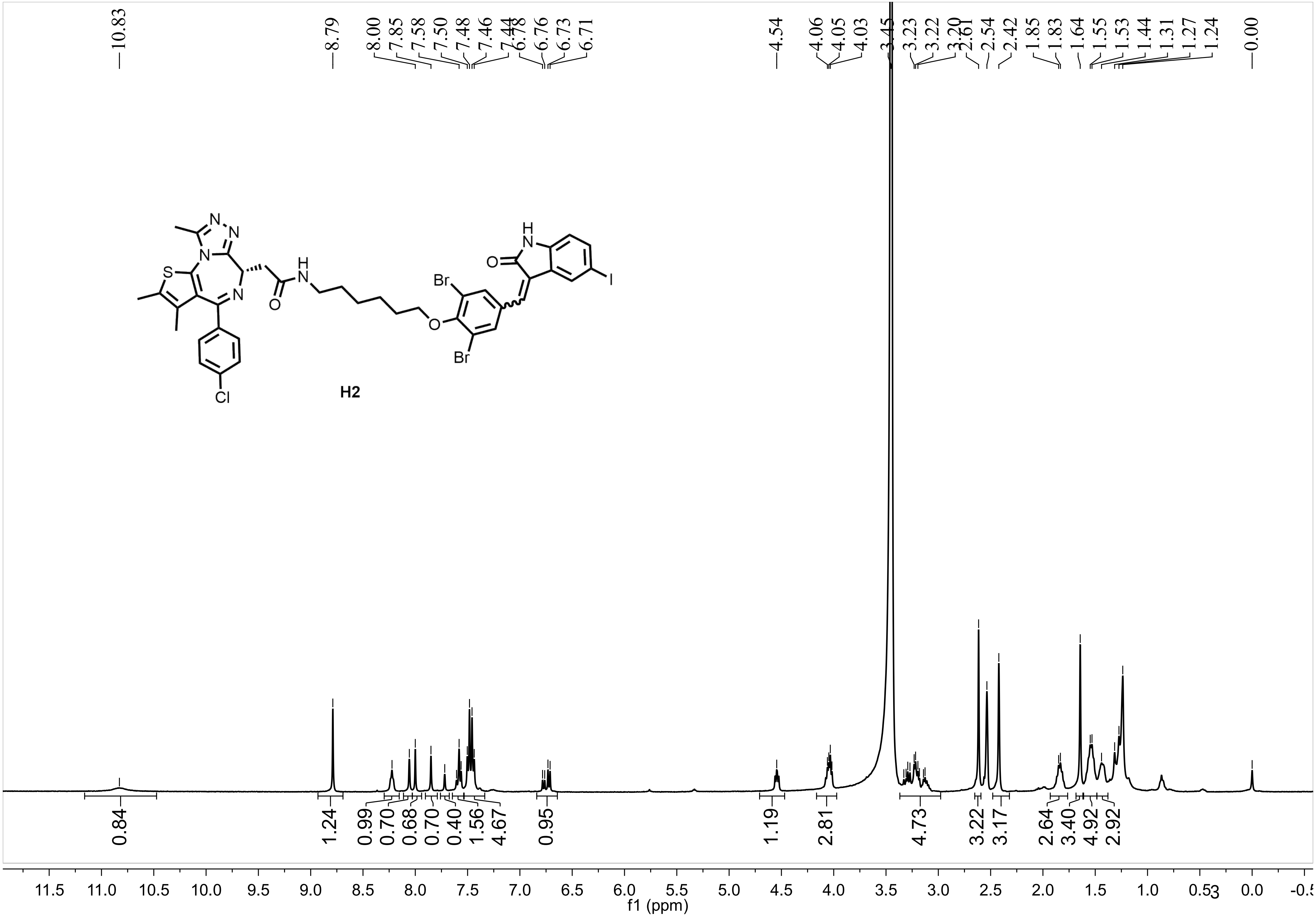
10.83
8.79
8.00
7.85
7.58
7.50
7.48
7.46
7.44
6.78
6.76
6.73
6.71
4.54
4.06
4.05
4.03
3.45
3.23
3.22
3.20
2.61
2.54
2.42
1.85
1.83
1.64
1.55
1.53
1.44
1.31
1.27
1.24
0.00
H2
0.84
1.24
0.99
0.70
0.68
0.70
0.40
1.56
4.67
0.95
1.19
2.81
4.73
3.22
3.17
2.64
3.40
4.92
2.92
11.5 11.0 10.5 10.0 9.5 9.0 8.5 8.0 7.5 7.0 6.5 6.0 5.5 5.0 4.5 4.0 3.5 3.0 2.5 2.0 1.5 1.0 0.5 0.0
f1 (ppm)

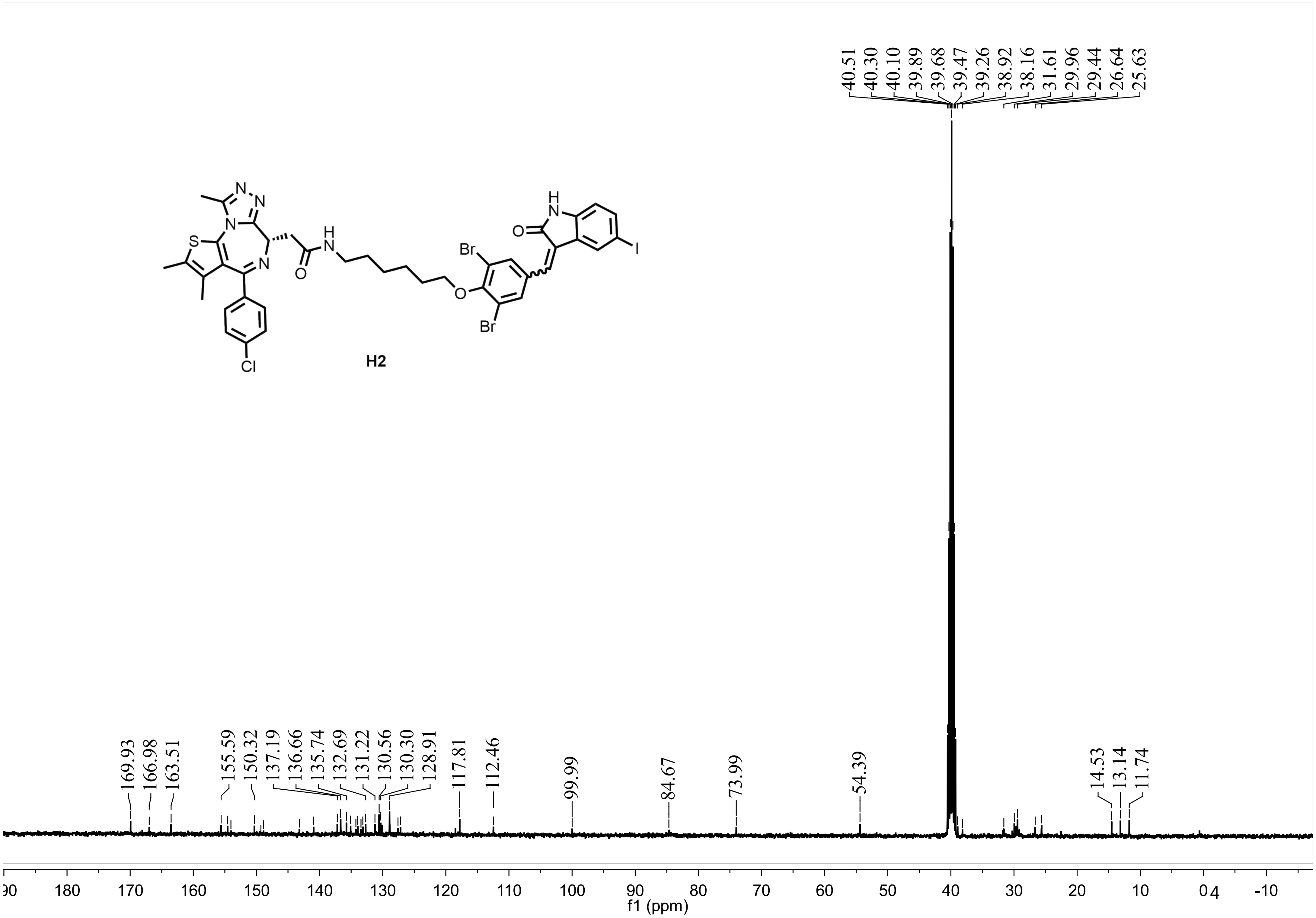
40.51
40.30
40.10
39.89
39.68
39.47
39.26
38.92
38.16
31.61
29.96
29.44
26.64
25.63
H2
Br
Br
O
N
N
N
S
N
H
O
Cl
I
169.93
166.98
163.51
155.59
150.32
137.19
136.66
135.74
132.69
131.22
130.56
130.30
128.91
117.81
112.46
99.99
84.67
73.99
54.39
14.53
13.14
11.74
180
170
160
150
140
130
120
110
100
90
80
70
60
50
40
30
20
10
0
-10
f1 (ppm)

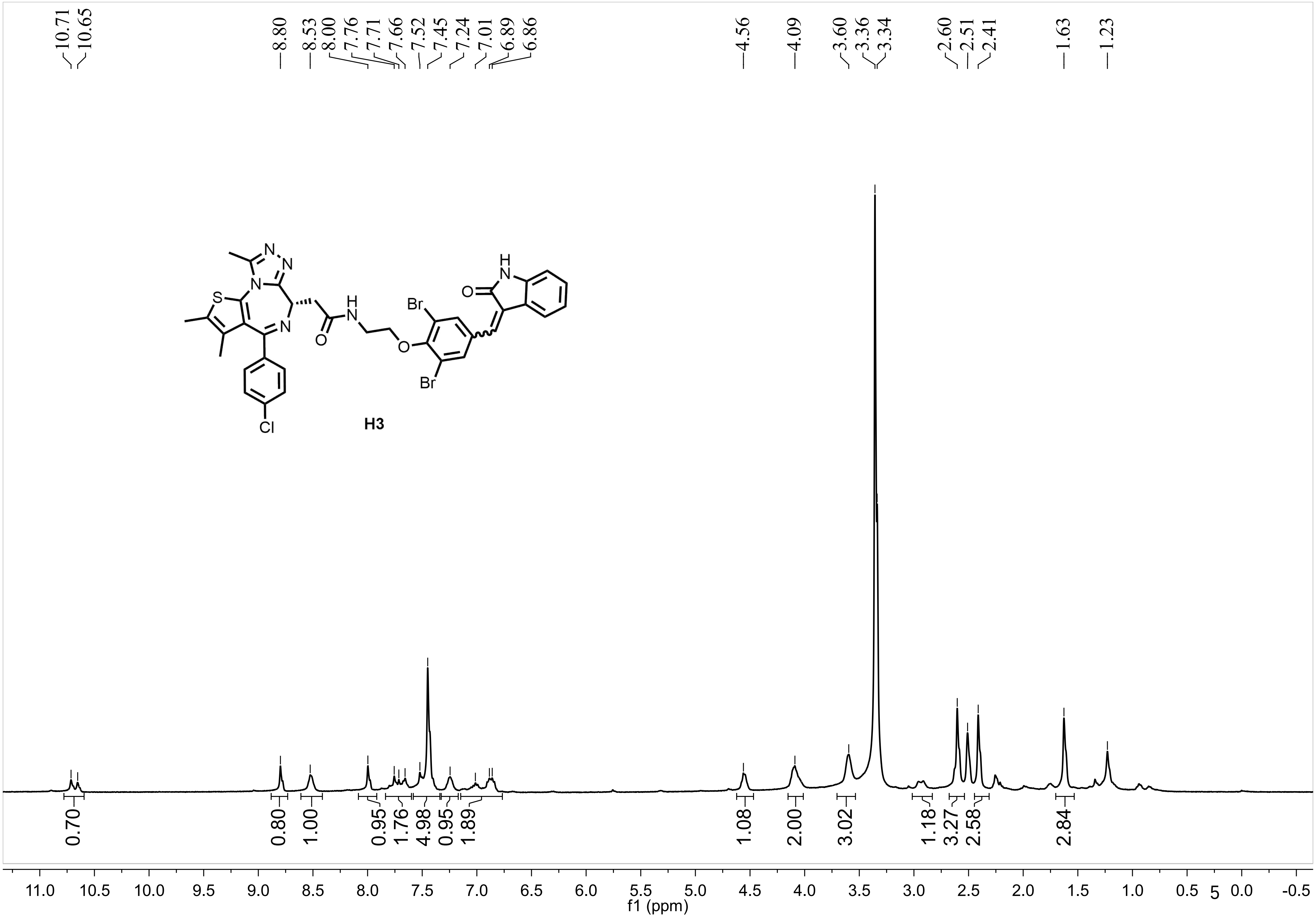
10.71
10.65
8.80
8.53
8.00
7.76
7.71
7.66
7.52
7.45
7.24
7.01
6.89
6.86
4.56
4.09
3.60
3.36
3.34
2.60
2.51
2.41
1.63
1.23
H3
0.70
0.80
1.00
0.95
1.76
4.98
0.95
1.89
1.08
2.00
3.02
1.18
3.27
2.58
2.84
11.0 10.5 10.0 9.5 9.0 8.5 8.0 7.5 7.0 6.5 6.0 5.5 5.0 4.5 4.0 3.5 3.0 2.5 2.0 1.5 1.0 0.5 0.0 -0.5
f1 (ppm)

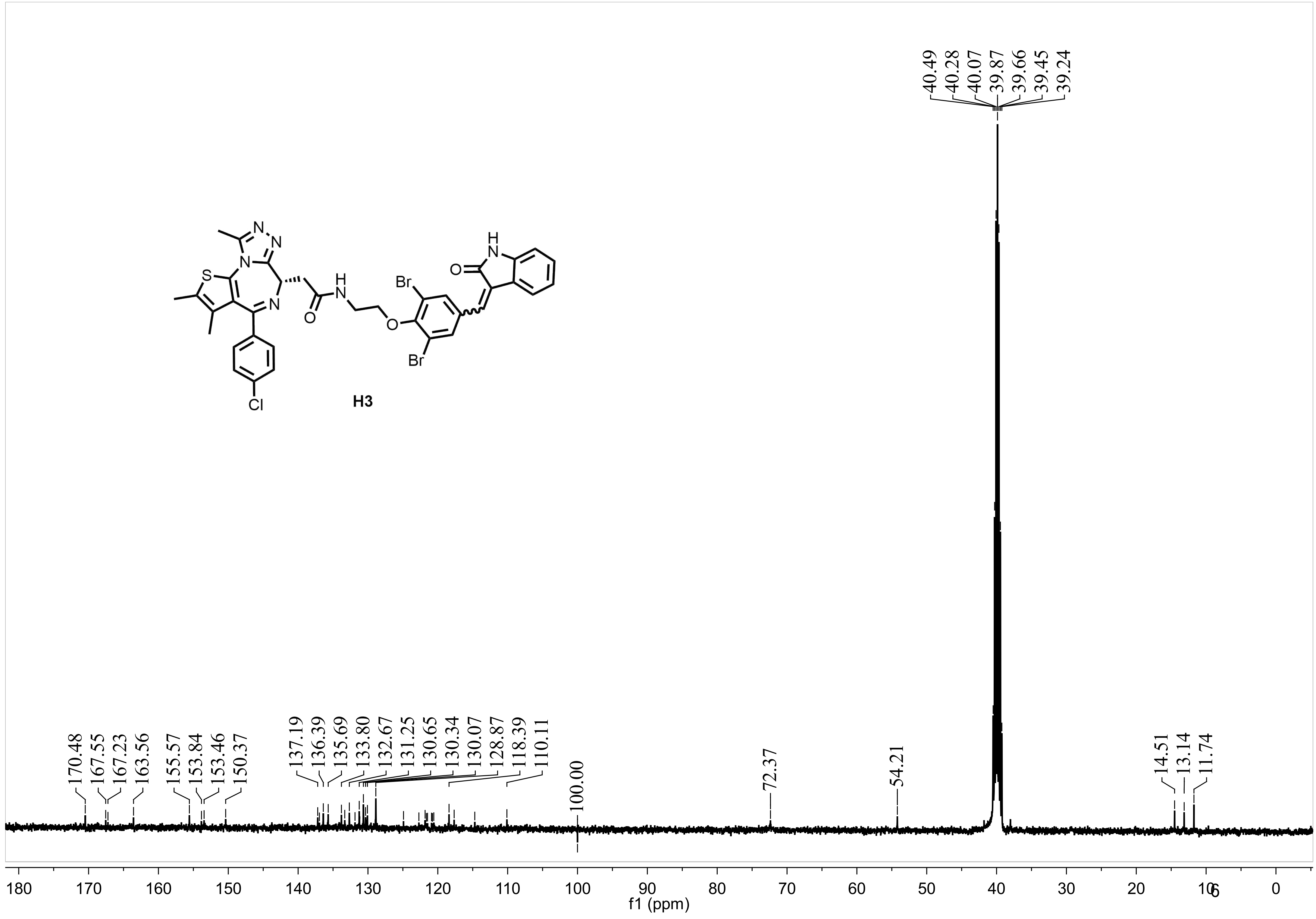
40.49
40.28
40.07
39.87
39.66
39.45
39.24
H3
170.48
167.55
167.23
163.56
155.57
153.84
153.46
150.37
137.19
136.39
135.69
133.80
132.67
131.25
130.65
130.34
130.07
128.87
118.39
110.11
100.00
72.37
54.21
14.51
13.14
11.74
180 170 160 150 140 130 120 110 100 90 80 70 60 50 40 30 20 10 0
f1 (ppm)

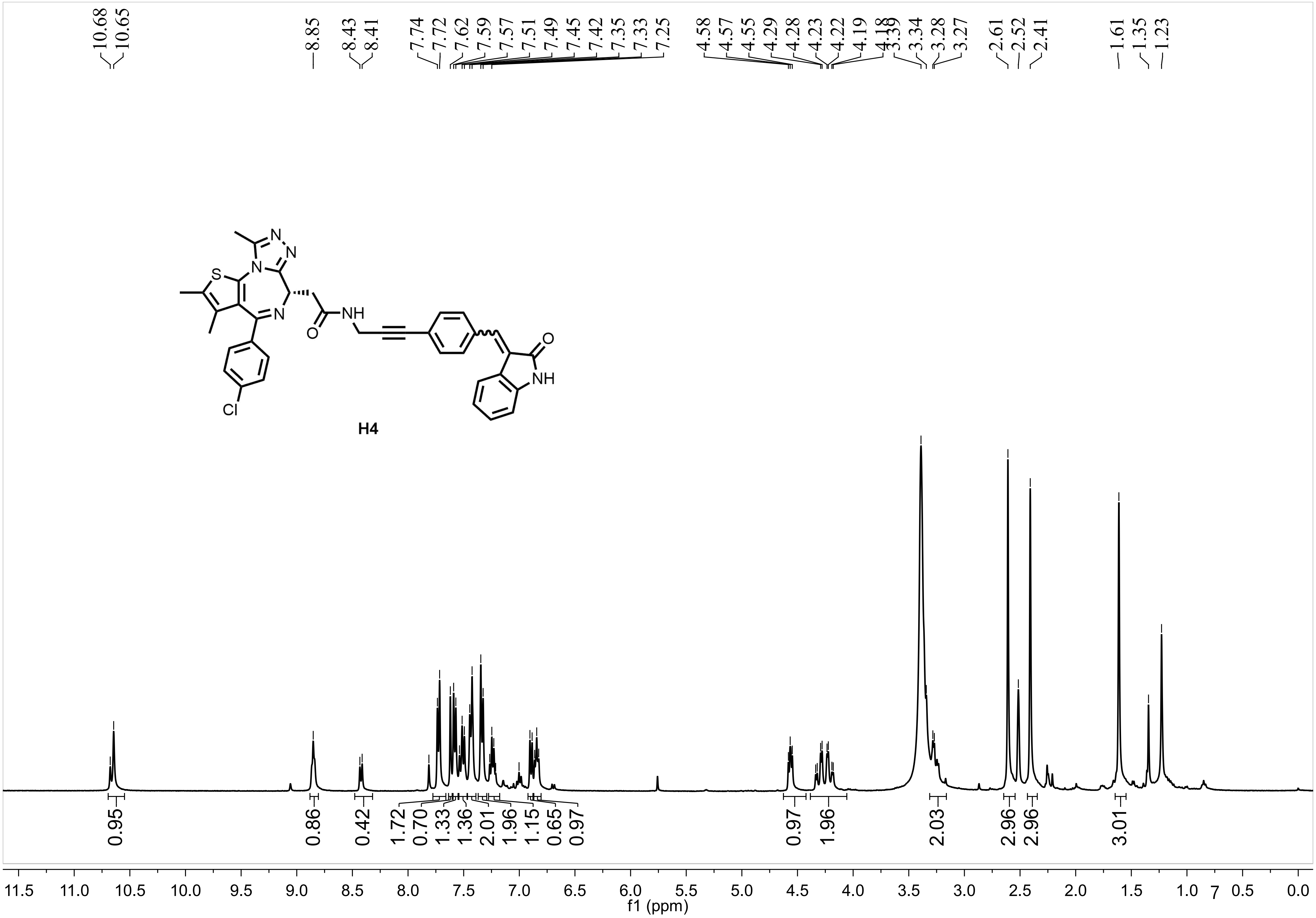
10.68
10.65
8.85
8.43
8.41
7.74
7.72
7.62
7.59
7.57
7.51
7.49
7.45
7.42
7.35
7.33
7.25
4.58
4.57
4.55
4.29
4.28
4.23
4.22
4.19
4.18
3.39
3.34
3.28
3.27
2.61
2.52
2.41
1.61
1.35
1.23
S
N
N
N
N
NH
O
Cl
O
NH
H4
0.95
0.86
0.42
1.72
0.70
1.33
1.36
2.01
1.96
1.15
0.65
0.97
0.97
1.96
2.03
2.96
2.96
3.01
11.5 11.0 10.5 10.0 9.5 9.0 8.5 8.0 7.5 7.0 6.5 6.0 5.5 5.0 4.5 4.0 3.5 3.0 2.5 2.0 1.5 1.0 0.5 0.0
f1 (ppm)

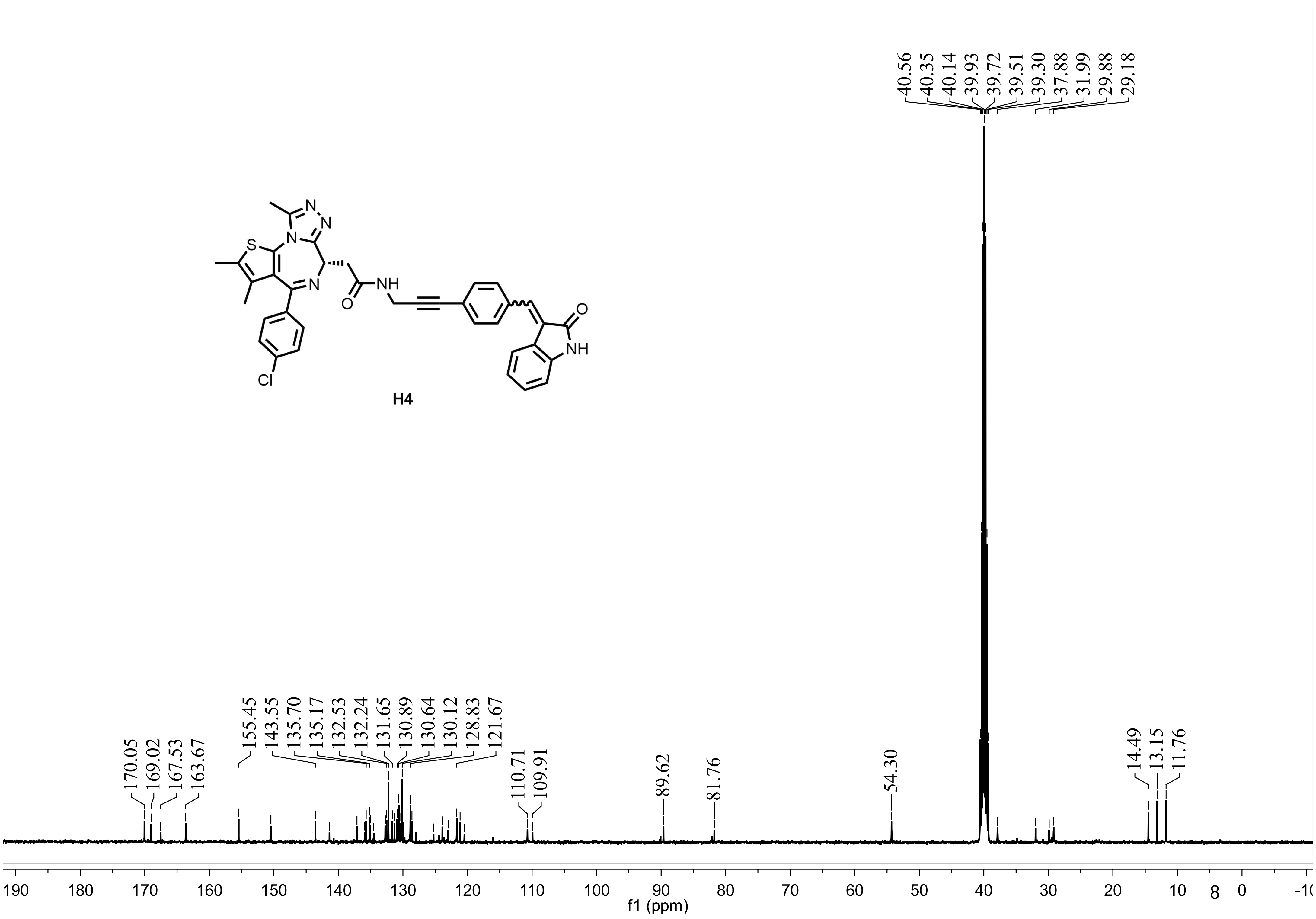
40.56
40.35
40.14
39.93
39.72
39.51
39.30
37.88
31.99
29.88
29.18
N
N
N
S
N
N
O
NH
O
NH
Cl
H4
170.05
169.02
167.53
163.67
155.45
143.55
135.70
135.17
132.53
132.24
131.65
130.89
130.64
130.12
128.83
121.67
110.71
109.91
89.62
81.76
54.30
14.49
13.15
11.76
190 180 170 160 150 140 130 120 110 100 90 80 70 60 50 40 30 20 10 0
f1 (ppm)

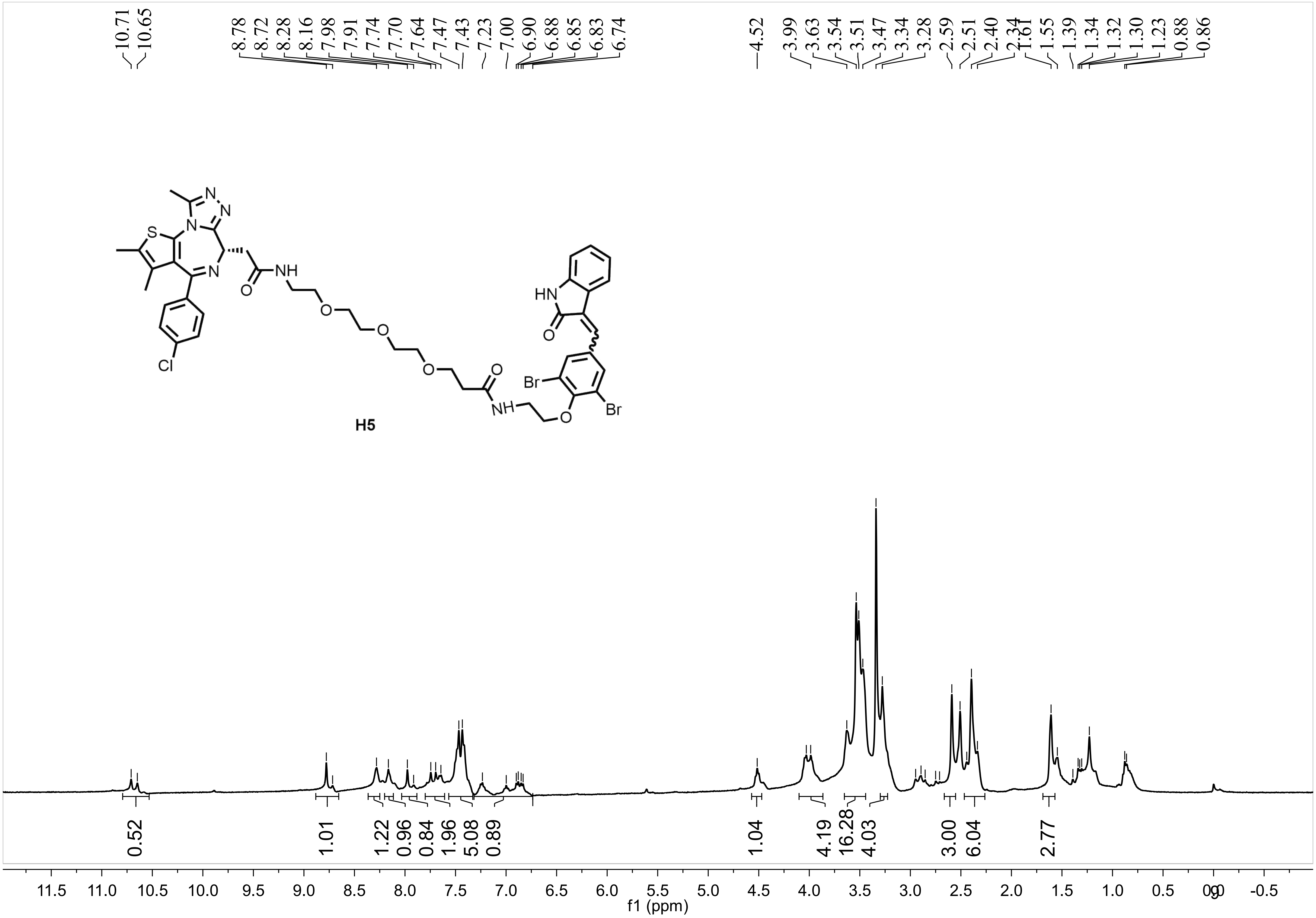
10.71
10.65
8.78
8.72
8.28
8.16
7.98
7.91
7.74
7.70
7.64
7.47
7.43
7.23
7.00
6.90
6.88
6.85
6.83
6.74
4.52
3.99
3.63
3.54
3.51
3.47
3.34
3.28
2.59
2.51
2.40
2.34
1.61
1.55
1.39
1.34
1.32
1.30
1.23
0.88
0.86
N
N
N
S
N
NH
O
O
O
O
O
Cl
HN
O
Br
Br
NH
O
H5
0.52
1.01
1.22
0.96
0.84
1.96
5.08
0.89
1.04
4.19
16.28
4.03
3.00
6.04
2.77
11.5
11.0
10.5
10.0
9.5
9.0
8.5
8.0
7.5
7.0
6.5
6.0
5.5
5.0
4.5
4.0
3.5
3.0
2.5
2.0
1.5
1.0
0.5
0.0
-0.5
f1 (ppm)

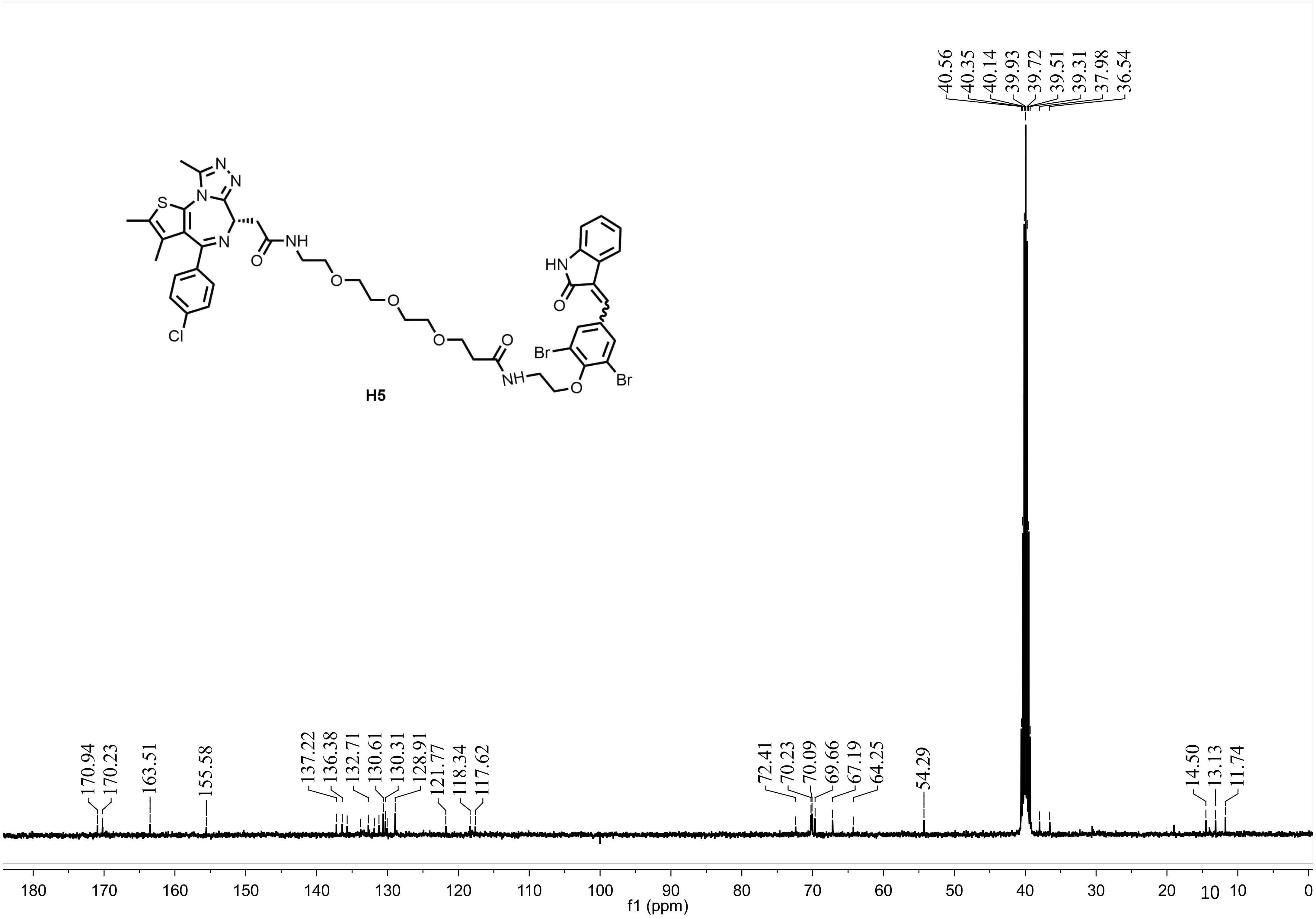

40.56
40.35
40.14
39.93
39.72
39.51
39.31
37.98
36.54
H5
HN
O
NH
Br
Cl
S
N
170.94
170.23
163.51
155.58
137.22
136.38
132.71
130.61
130.31
128.91
121.77
118.34
117.62
72.41
70.23
70.09
69.66
67.19
64.25
54.29
14.50
13.13
11.74
180 170 160 150 140 130 120 110 100 90 80 70 60 50 40 30 20 10 0
f1 (ppm)

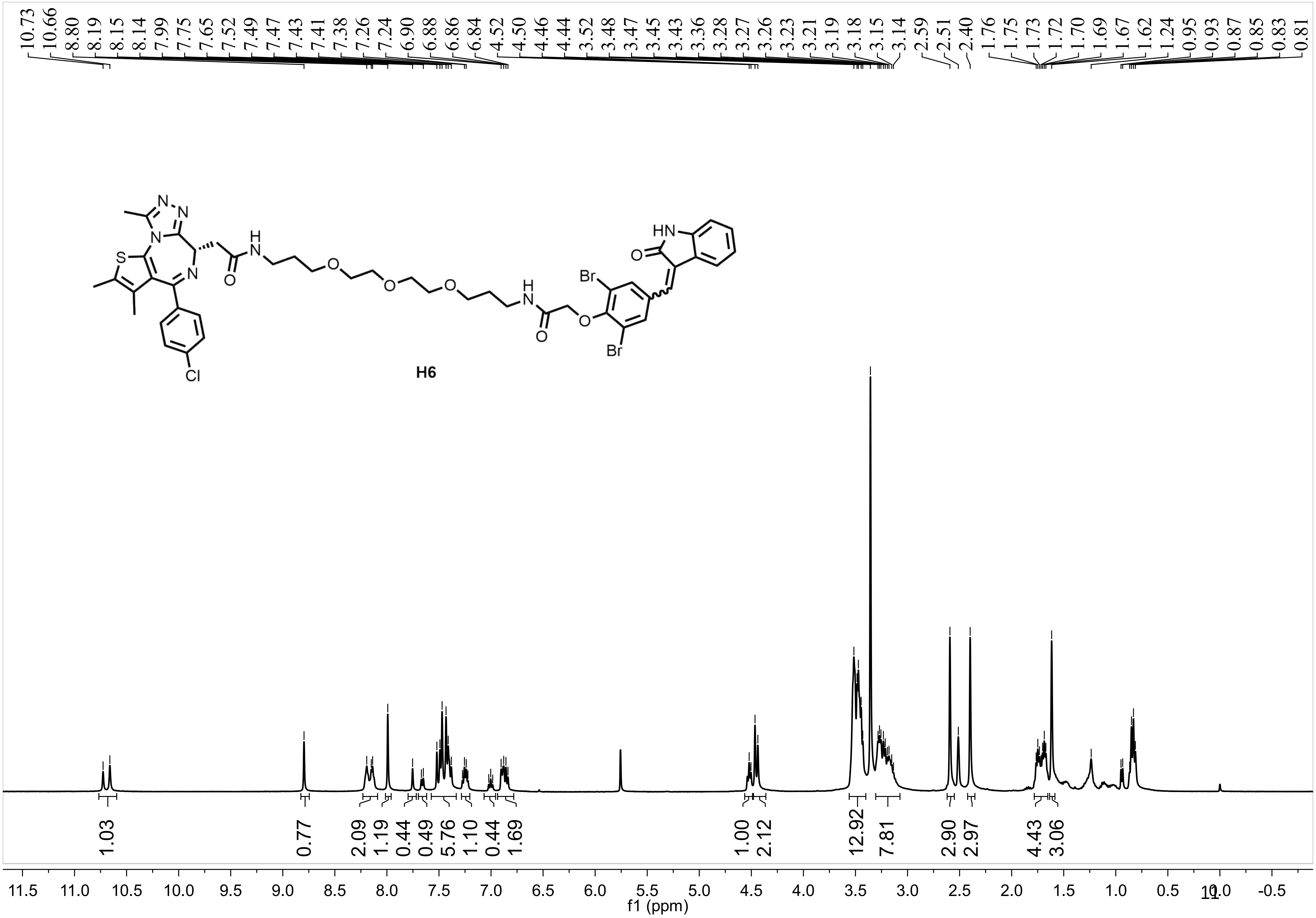
10.73 10.66 8.80 8.19 8.15 8.14 7.99 7.75 7.65 7.52 7.49 7.47 7.43 7.41 7.38 7.26 7.24 6.90 6.88 6.86 6.84 4.52 4.50 4.46 4.44 3.52 3.48 3.47 3.45 3.43 3.36 3.28 3.27 3.26 3.23 3.21 3.19 3.18 3.15 3.14 2.59 2.51 2.40 1.76 1.75 1.73 1.72 1.70 1.69 1.67 1.62 1.24 0.95 0.93 0.87 0.85 0.83 0.81
H6
1.03 0.77 2.09 1.19 0.44 0.49 5.76 1.10 0.44 1.69 1.00 2.12 12.92 7.81 2.90 2.97 4.43 3.06
11.5 11.0 10.5 10.0 9.5 9.0 8.5 8.0 7.5 7.0 6.5 6.0 5.5 5.0 4.5 4.0 3.5 3.0 2.5 2.0 1.5 1.0 0.5 0.0 -0.5
f1 (ppm)

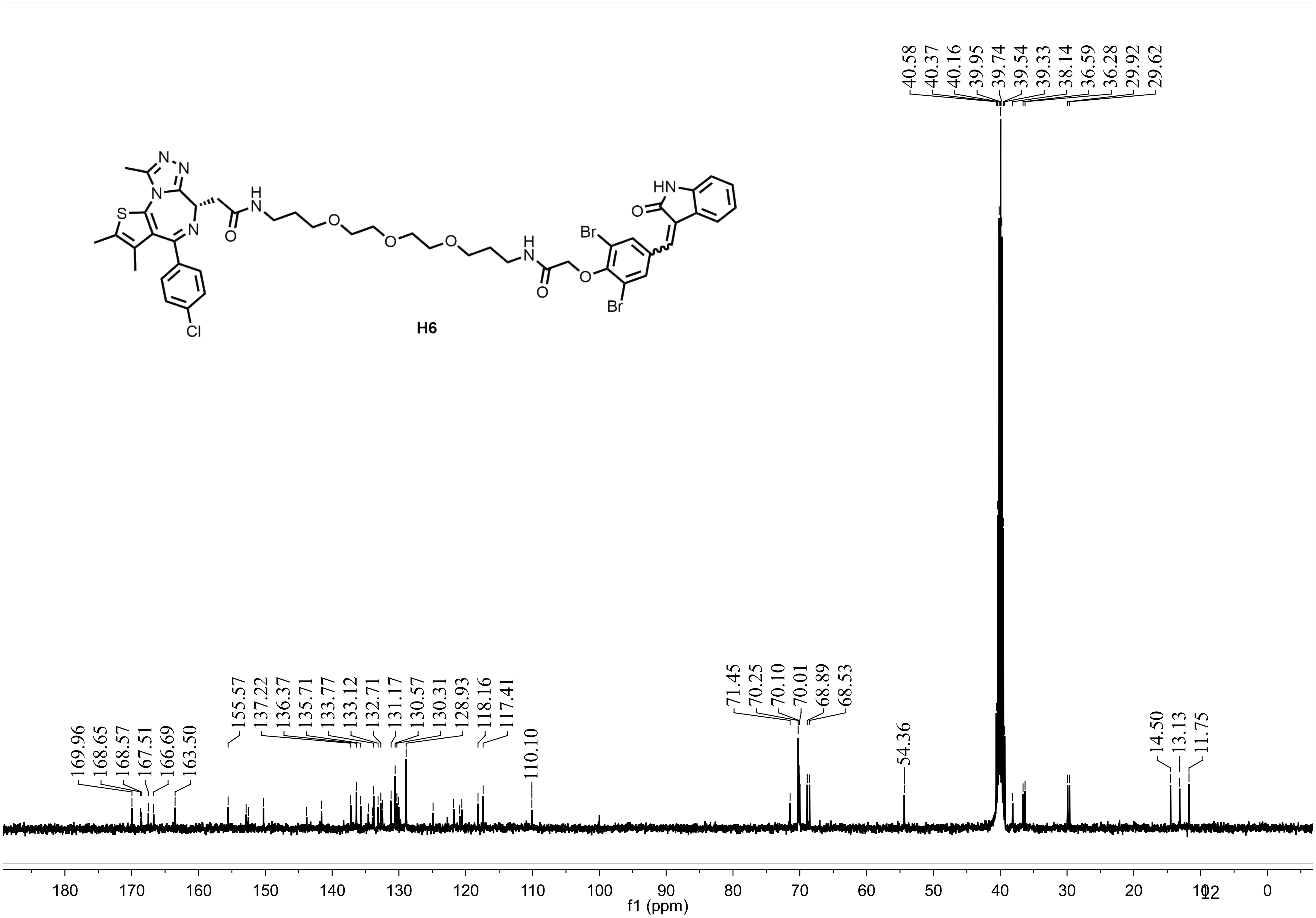
H6
40.58
40.37
40.16
39.95
39.74
39.54
39.33
38.14
36.59
36.28
29.92
29.62
169.96
168.65
168.57
167.51
166.69
163.50
155.57
137.22
136.37
135.71
133.77
133.12
132.71
131.17
130.57
130.31
128.93
118.16
117.41
110.10
71.45
70.25
70.10
70.01
68.89
68.53
54.36
14.50
13.13
11.75
180 170 160 150 140 130 120 110 100 90 80 70 60 50 40 30 20 10 0
f1 (ppm)

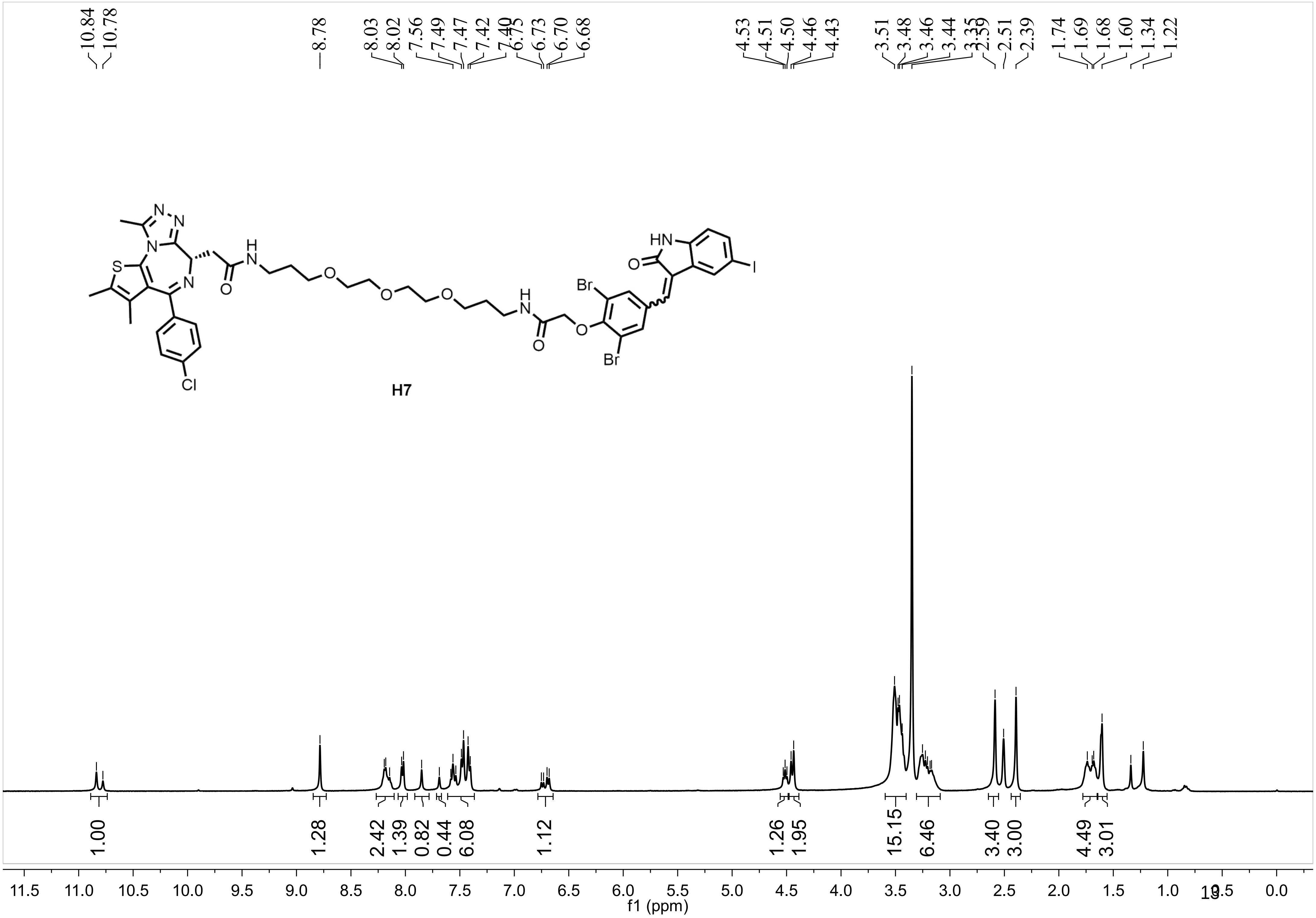

10.84
10.78
8.78
8.03
8.02
7.56
7.49
7.47
7.42
7.40
6.75
6.73
6.70
6.68
4.53
4.51
4.50
4.46
4.43
3.51
3.48
3.46
3.44
3.35
2.59
2.51
2.39
1.74
1.69
1.68
1.60
1.34
1.22
N
N
N
S
N
H
N
O
O
O
O
H
N
O
O
Br
Br
HN
O
I
Cl
H7
1.00
1.28
2.42
1.39
0.82
0.44
6.08
1.12
1.26
1.95
15.15
6.46
3.40
3.00
4.49
3.01
11.5
11.0
10.5
10.0
9.5
9.0
8.5
8.0
7.5
7.0
6.5
6.0
5.5
5.0
4.5
4.0
3.5
3.0
2.5
2.0
1.5
1.0
0.0
f1 (ppm)

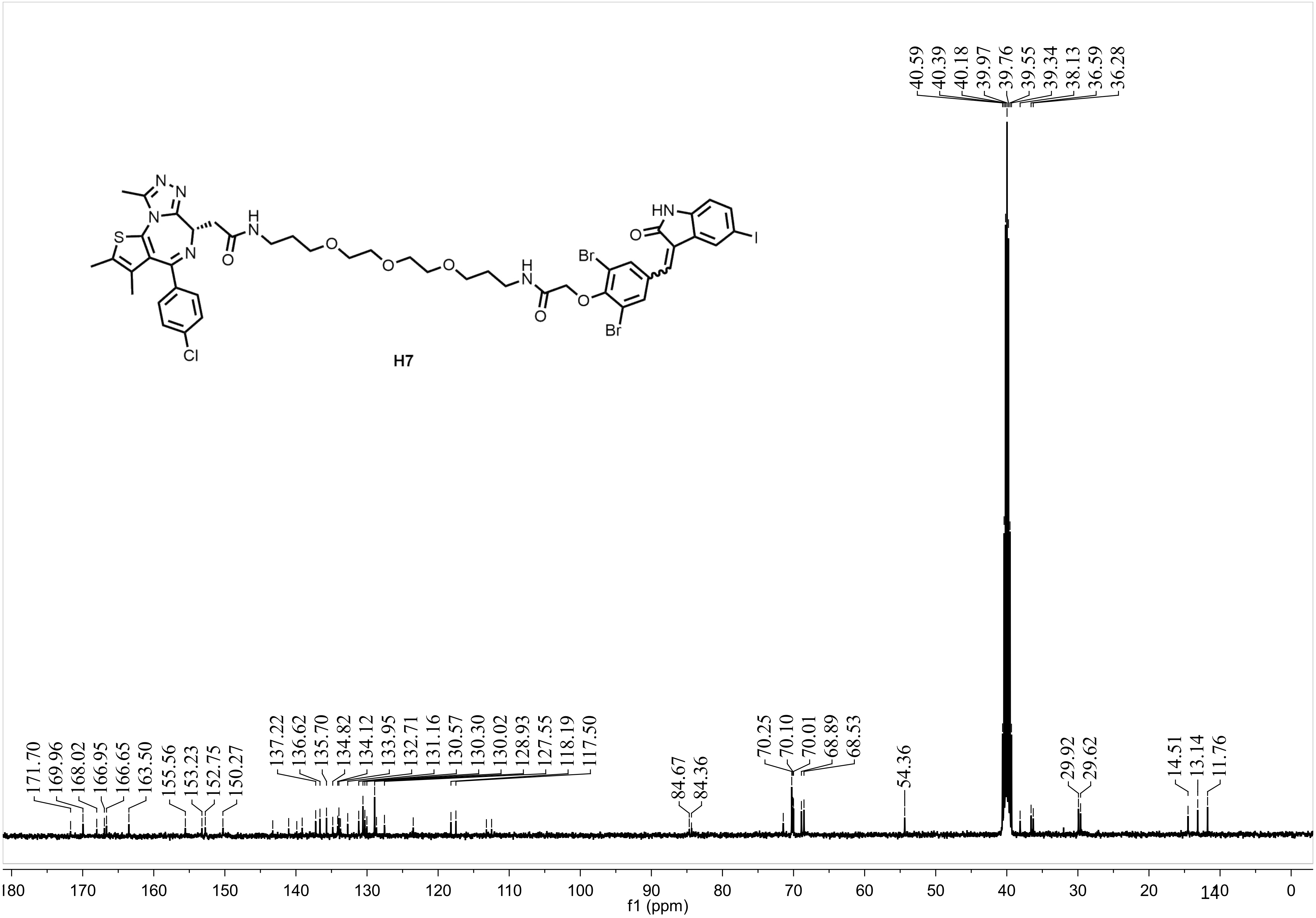
H7
40.59 40.39 40.18 39.97 39.76 39.55 39.34 38.13 36.59 36.28
171.70 169.96 168.02 166.95 166.65 163.50 155.56 153.23 152.75 150.27
137.22 136.62 135.70 134.82 134.12 133.95 132.71 131.16 130.57 130.30 130.02 128.93 127.55 118.19 117.50
84.67 84.36
70.25 70.10 70.01 68.89 68.53
54.36
29.92 29.62
14.51 13.14 11.76
f1 (ppm)

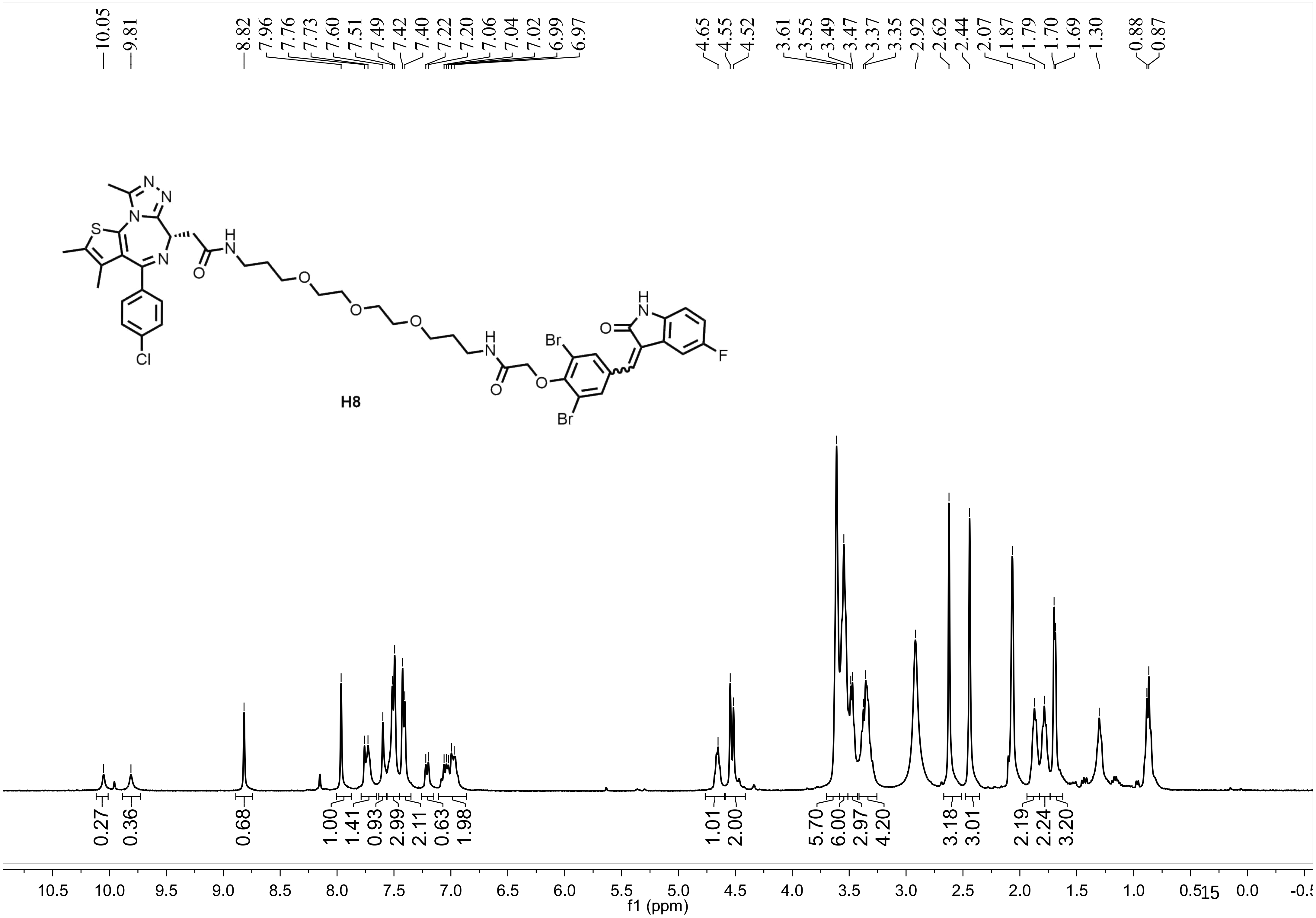
10.05
9.81
8.82
7.96
7.76
7.73
7.60
7.51
7.49
7.42
7.40
7.22
7.20
7.06
7.04
7.02
6.99
6.97
4.65
4.55
4.52
3.61
3.55
3.49
3.47
3.37
3.35
2.92
2.62
2.44
2.07
1.87
1.79
1.70
1.69
1.30
0.88
0.87
H8
0.27
0.36
0.68
1.00
1.41
0.93
2.99
2.11
0.63
1.98
1.01
2.00
5.70
6.00
2.97
4.20
3.18
3.01
2.19
2.24
3.20
10.5
10.0
9.5
9.0
8.5
8.0
7.5
7.0
6.5
6.0
5.5
5.0
4.5
4.0
3.5
3.0
2.5
2.0
1.5
1.0
0.5
0.0
f1 (ppm)

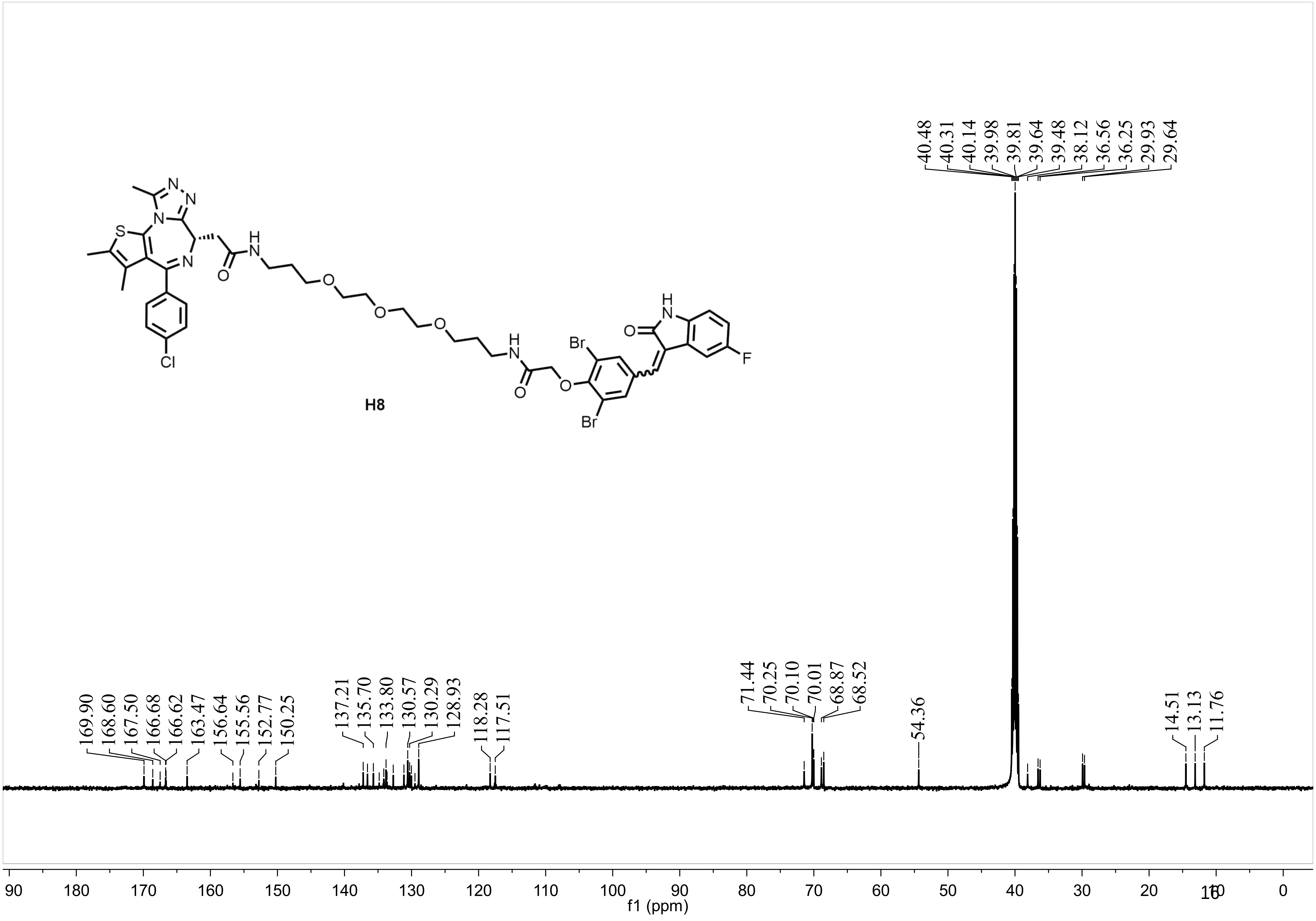

40.48
40.31
40.14
39.98
39.81
39.64
39.48
38.12
36.56
36.25
29.93
29.64
S
N
N
N
N
N
H
O
O
O
O
Cl
H
N
O
Br
Br
O
H
N
O
F
H8
169.90
168.60
167.50
166.68
166.62
163.47
156.64
155.56
152.77
150.25
137.21
135.70
133.80
130.57
130.29
128.93
118.28
117.51
71.44
70.25
70.10
70.01
68.87
68.52
54.36
14.51
13.13
11.76
90
180
170
160
150
140
130
120
110
100
90
80
70
60
50
40
30
20
10
0
f1 (ppm)

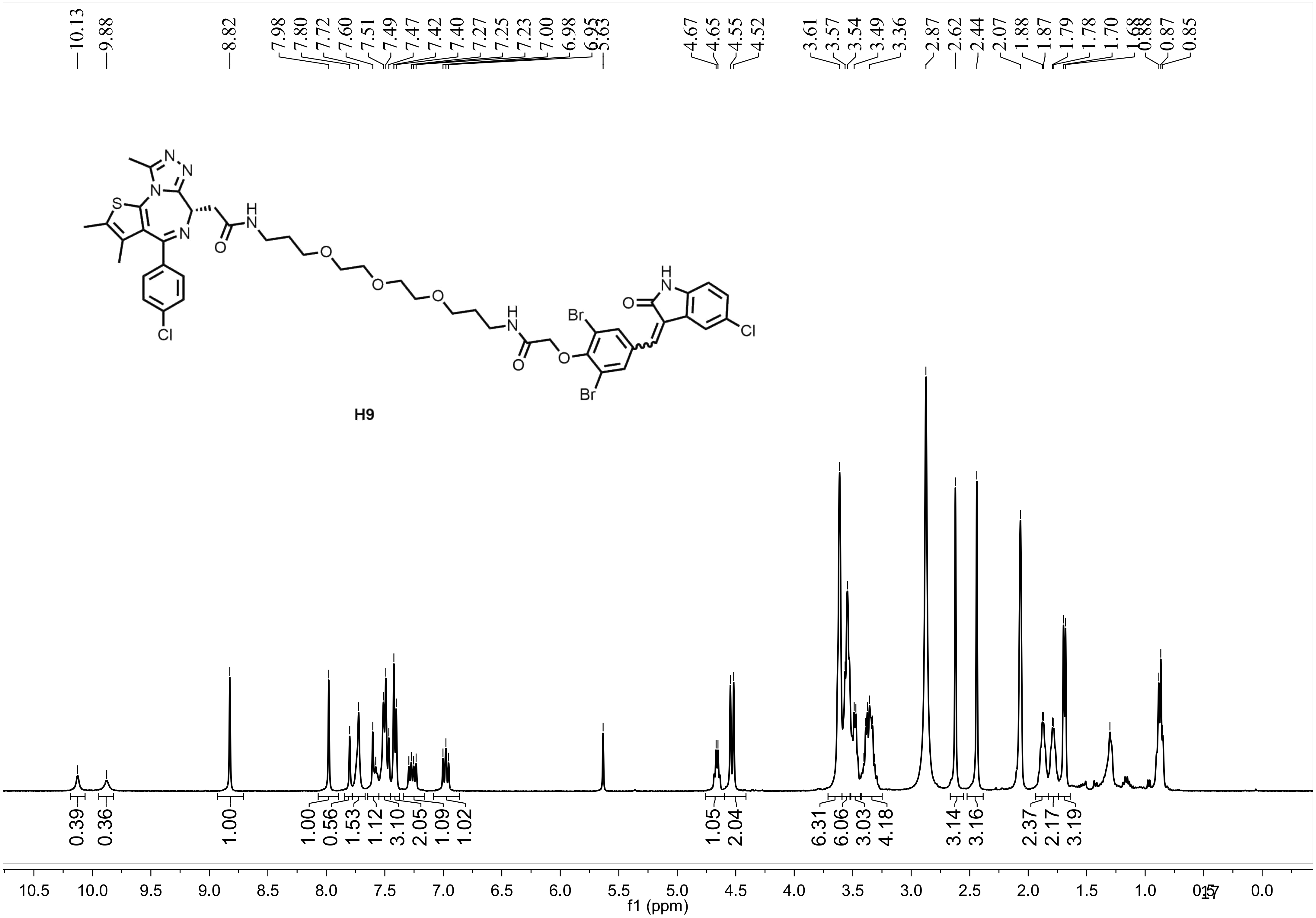
10.13
9.88
8.82
7.98
7.80
7.72
7.60
7.51
7.49
7.47
7.42
7.40
7.27
7.25
7.23
7.00
6.98
6.95
5.63
4.67
4.65
4.55
4.52
3.61
3.57
3.54
3.49
3.36
2.87
2.62
2.44
2.07
1.88
1.87
1.79
1.78
1.70
1.68
0.88
0.87
0.85
H9
0.39
0.36
1.00
1.00
0.56
1.53
1.12
3.10
2.05
1.09
1.02
1.05
2.04
6.31
6.06
3.03
4.18
3.14
3.16
2.37
2.17
3.19
10.5
10.0
9.5
9.0
8.5
8.0
7.5
7.0
6.5
6.0
5.5
5.0
4.5
4.0
3.5
3.0
2.5
2.0
1.5
1.0
0.5
0.0
f1 (ppm)

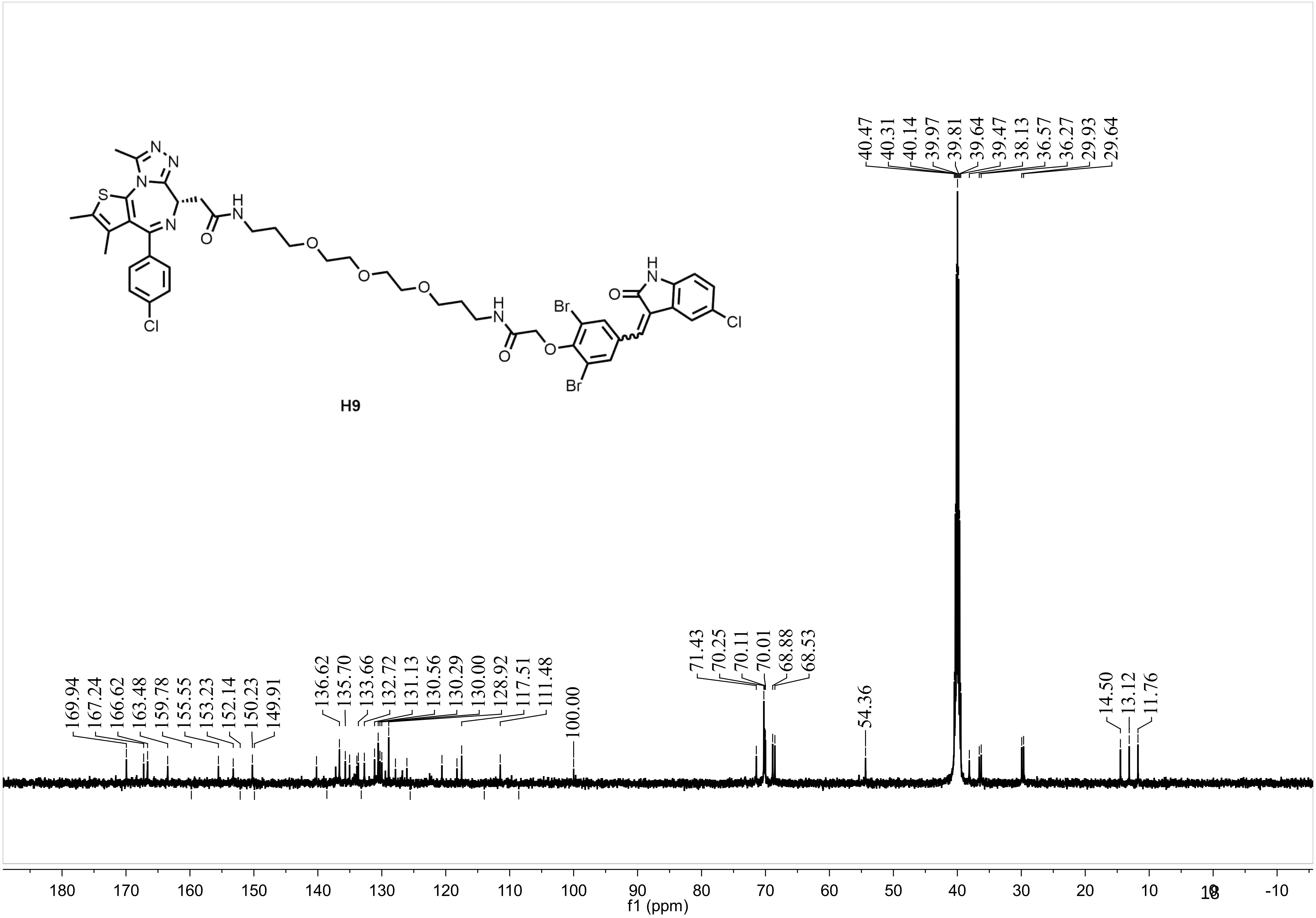
H9
169.94
167.24
166.62
163.48
159.78
155.55
153.23
152.14
150.23
149.91
136.62
135.70
133.66
132.72
131.13
130.56
130.29
130.00
128.92
117.51
111.48
100.00
71.43
70.25
70.11
70.01
68.88
68.53
54.36
40.47
40.31
40.14
39.97
39.81
39.64
39.47
38.13
36.57
36.27
29.93
29.64
14.50
13.12
11.76
180 170 160 150 140 130 120 110 100 90 80 70 60 50 40 30 20 10 0 -10
f1 (ppm)

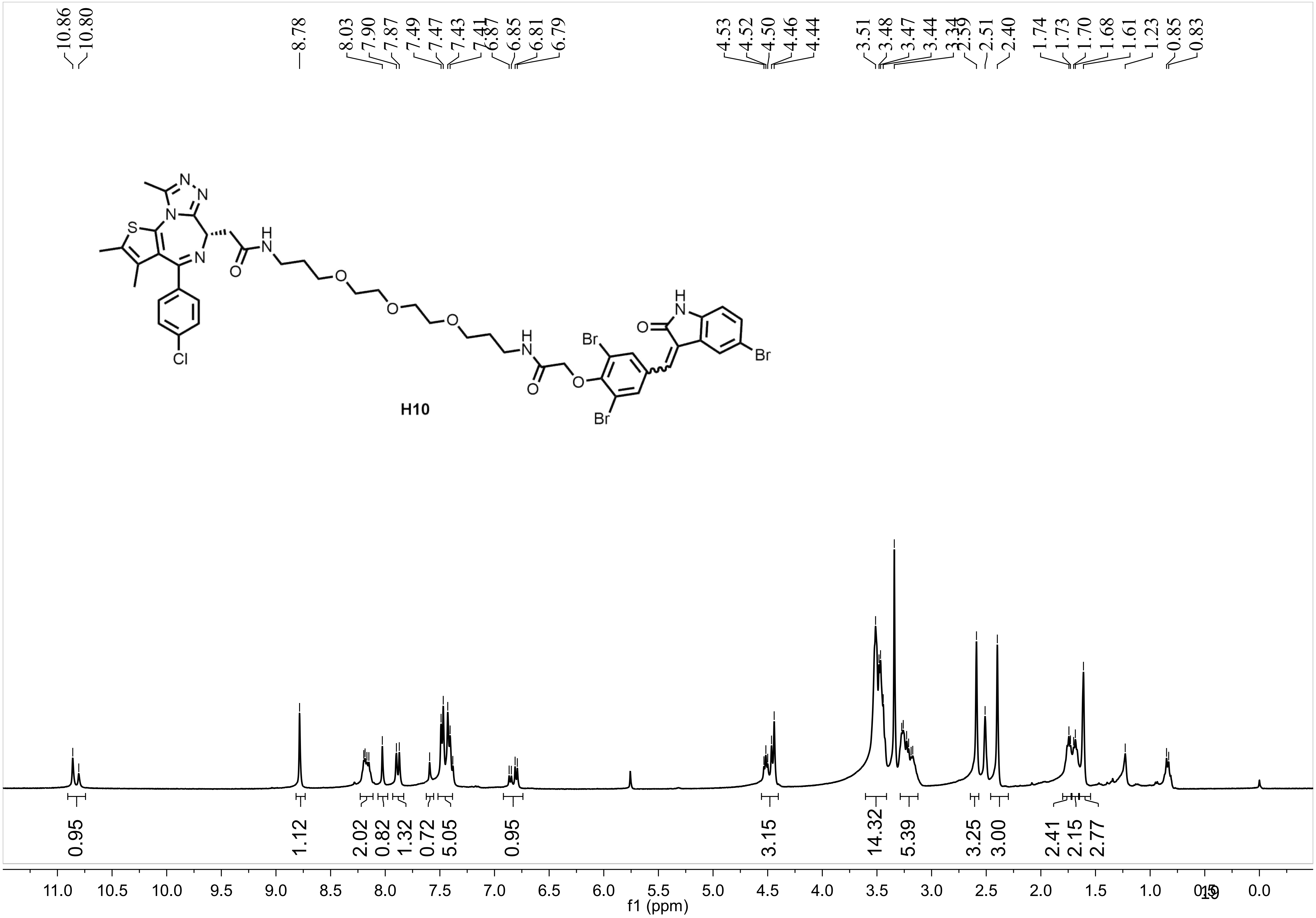

10.86
10.80
8.78
8.03
7.90
7.87
7.49
7.47
7.43
7.41
6.87
6.85
6.81
6.79
4.53
4.52
4.50
4.46
4.44
3.51
3.48
3.47
3.44
3.34
2.59
2.51
2.40
1.74
1.73
1.70
1.68
1.61
1.23
0.85
0.83
H10
Br
Br
Br
Cl
0.95
1.12
2.02
0.82
1.32
0.72
5.05
0.95
3.15
14.32
5.39
3.25
3.00
2.41
2.15
2.77
11.0 10.5 10.0 9.5 9.0 8.5 8.0 7.5 7.0 6.5 6.0 5.5 5.0 4.5 4.0 3.5 3.0 2.5 2.0 1.5 1.0 0.5 0.0
f1 (ppm)

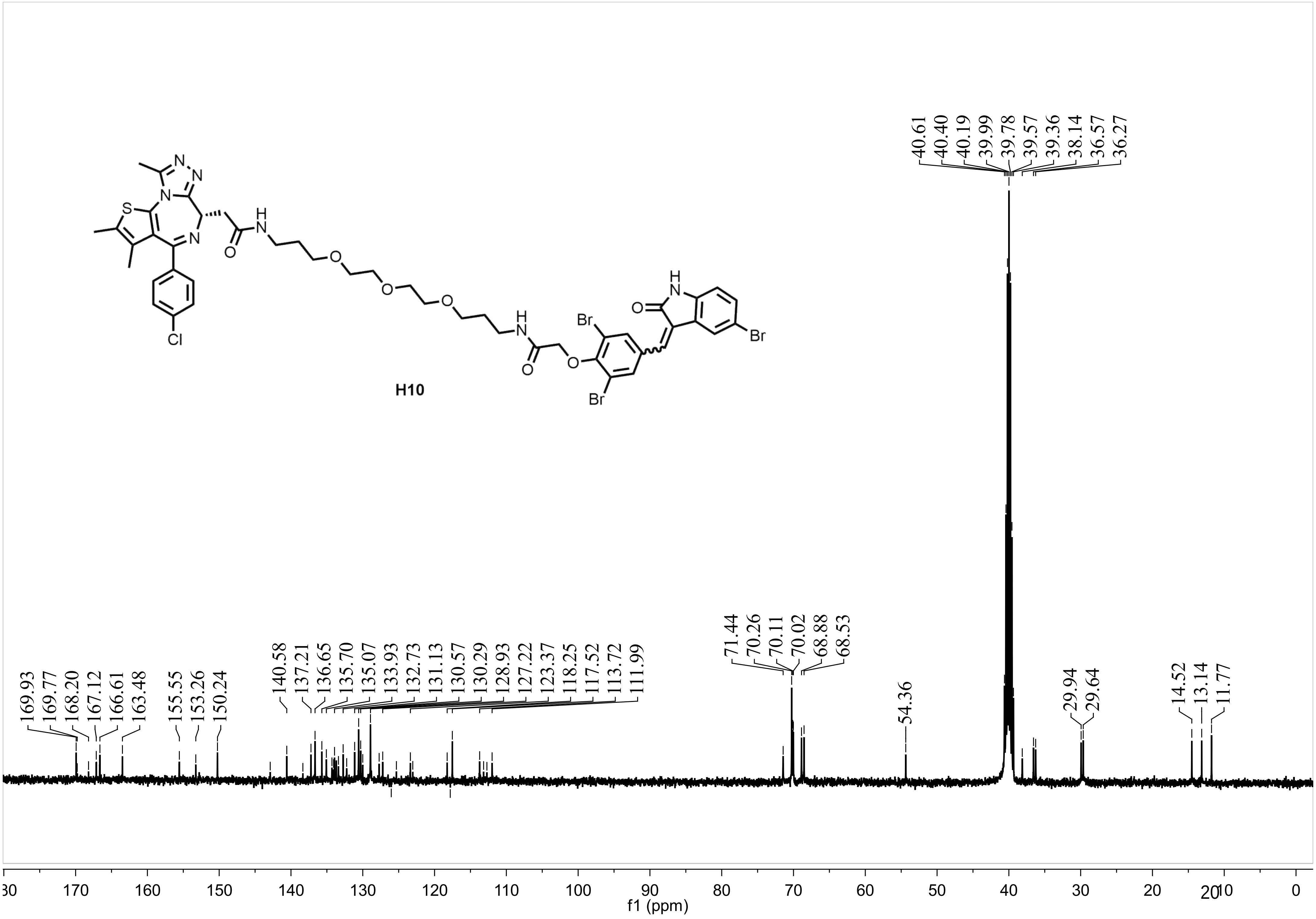
H10
169.93 169.77 168.20 167.12 166.61 163.48 155.55 153.26 150.24
140.58 137.21 136.65 135.70 135.07 133.93 132.73 131.13 130.57 130.29 128.93 127.22 123.37 118.25 117.52 113.72 111.99
71.44 70.26 70.11 70.02 68.88 68.53
54.36
40.61 40.40 40.19 39.99 39.78 39.57 39.36 38.14 36.57 36.27
29.94 29.64
14.52 13.14 11.77
f1 (ppm)

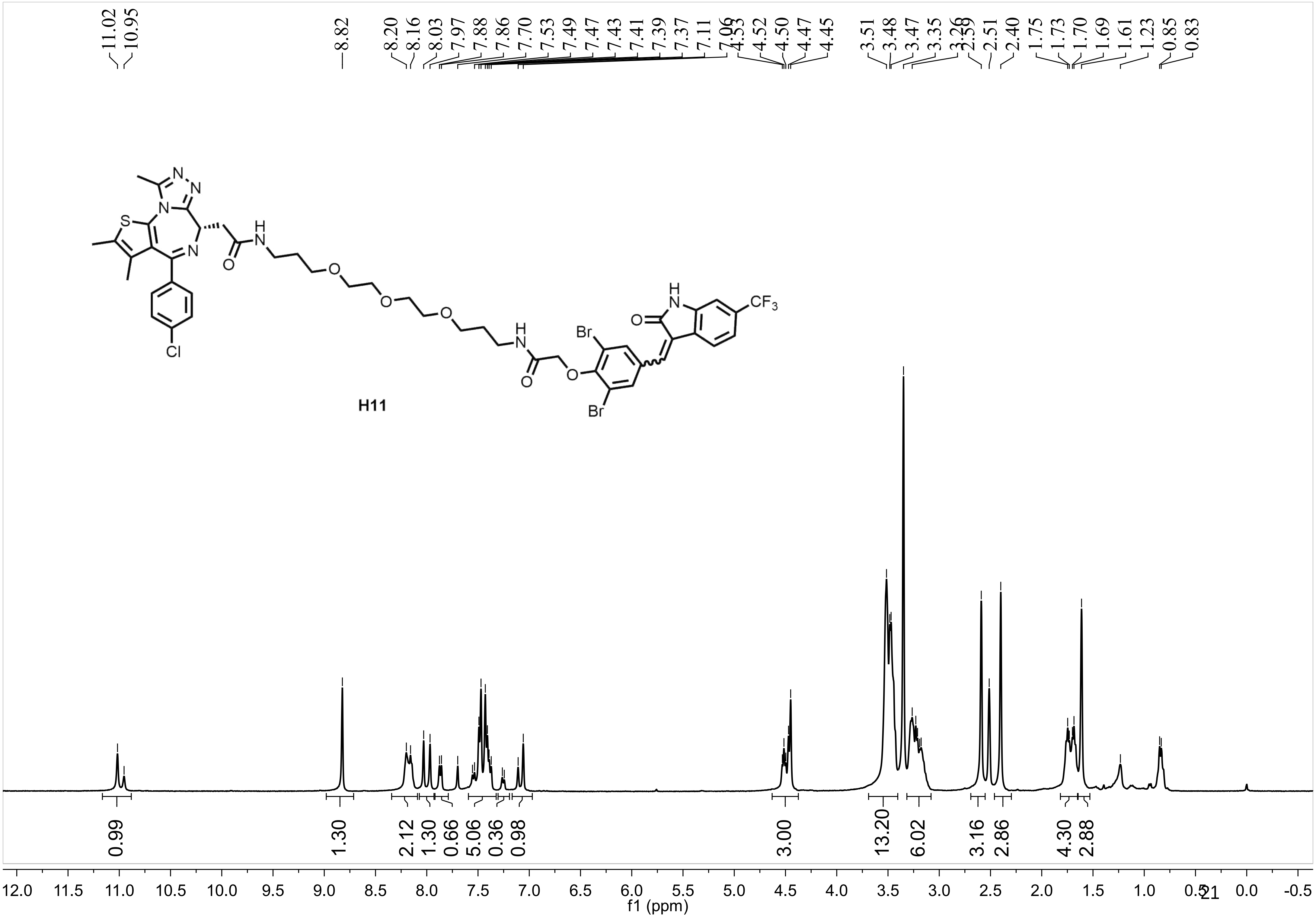
11.02
10.95
8.82
8.20
8.16
8.03
7.97
7.88
7.86
7.70
7.53
7.49
7.47
7.43
7.41
7.39
7.37
7.11
7.06
4.53
4.52
4.50
4.47
4.45
3.51
3.48
3.47
3.35
3.26
2.59
2.51
2.40
1.75
1.73
1.70
1.69
1.61
1.23
0.85
0.83
N
N
N
S
N
H
N
O
O
O
O
H
N
O
Br
O
Br
H
N
O
CF3
Cl
H11
0.99
1.30
2.12
1.30
0.66
5.06
0.36
0.98
3.00
13.20
6.02
3.16
2.86
4.30
2.88
12.0 11.5 11.0 10.5 10.0 9.5 9.0 8.5 8.0 7.5 7.0 6.5 6.0 5.5 5.0 4.5 4.0 3.5 3.0 2.5 2.0 1.5 1.0 0.5 0.0 -0.5
f1 (ppm)

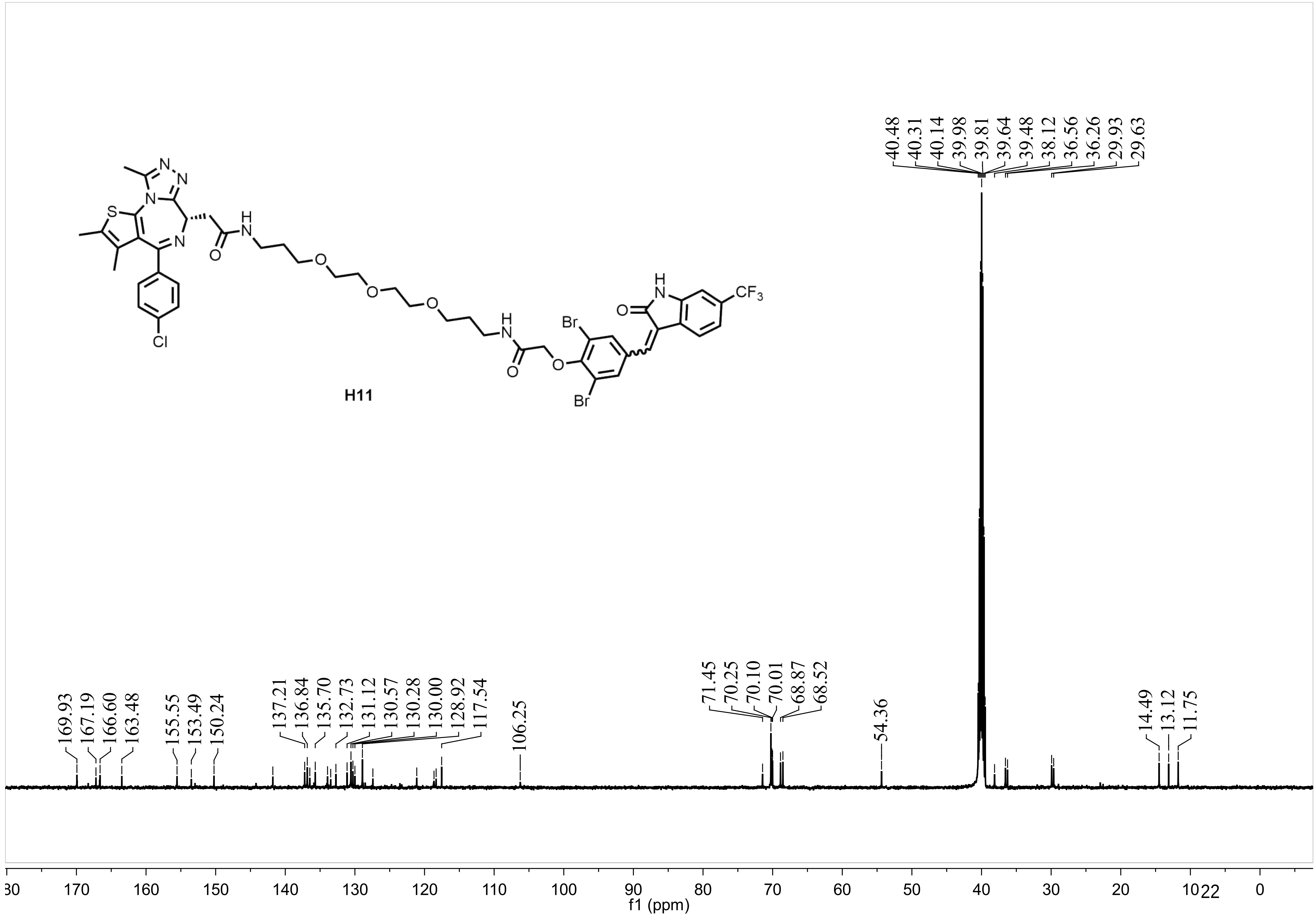

H11
N
N
N
S
N
H
N
O
O
O
O
H
N
O
O
Br
Br
H
N
O
CF3
Cl
40.48
40.31
40.14
39.98
39.81
39.64
39.48
38.12
36.56
36.26
29.93
29.63
169.93
167.19
166.60
163.48
155.55
153.49
150.24
137.21
136.84
135.70
132.73
131.12
130.57
130.28
130.00
128.92
117.54
106.25
71.45
70.25
70.10
70.01
68.87
68.52
54.36
14.49
13.12
11.75
f1 (ppm)

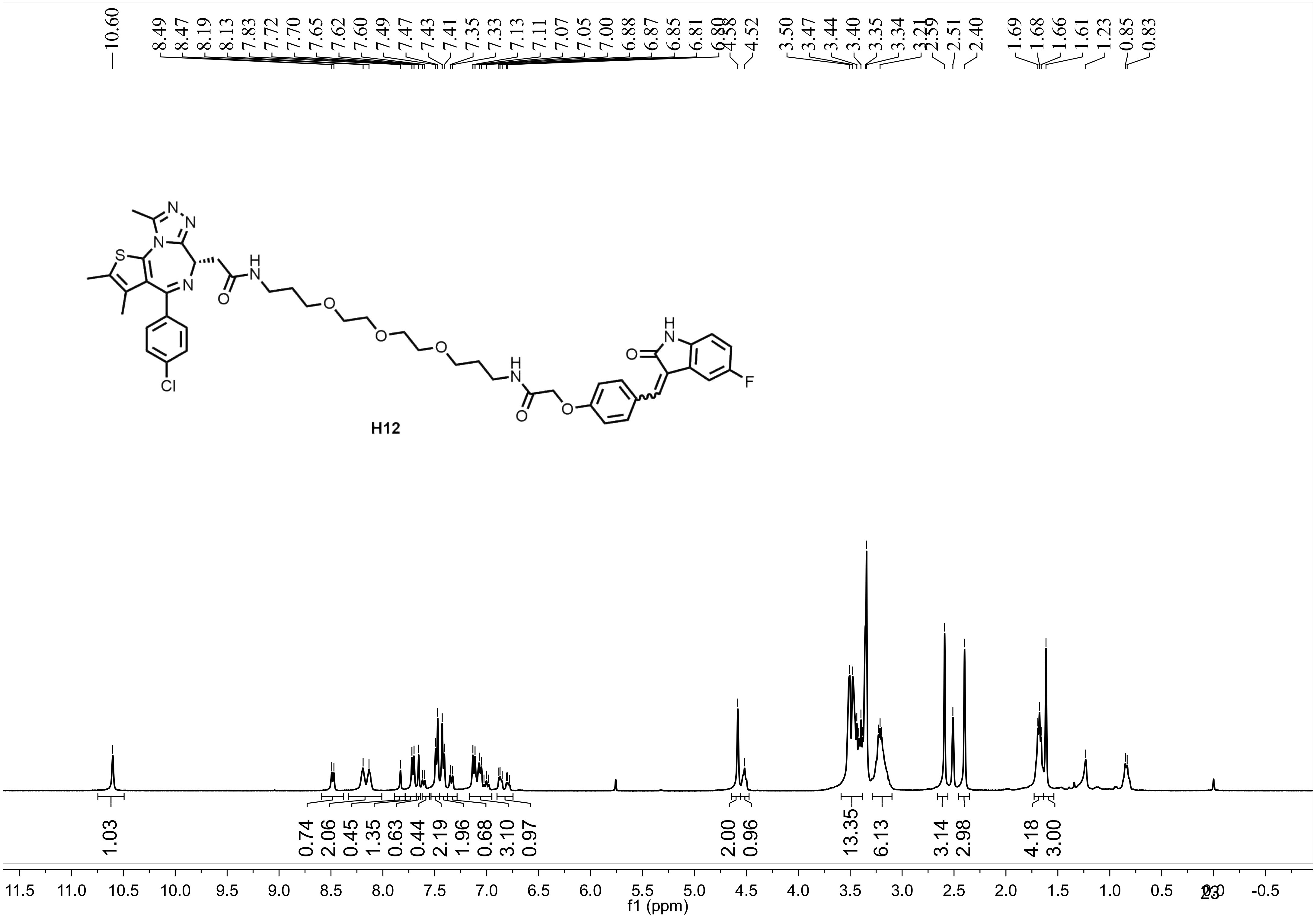
10.60
8.49 8.47 8.19 8.13 7.83 7.72 7.70 7.65 7.62 7.60 7.49 7.47 7.43 7.41 7.35 7.33 7.13 7.11 7.07 7.05 7.00 6.88 6.87 6.85 6.81 6.80 4.58 4.52
3.50 3.47 3.44 3.40 3.35 3.34 3.21 2.59 2.51 2.40
1.69 1.68 1.66 1.61 1.23 0.85 0.83
H12
1.03
0.74 2.06 0.45 1.35 0.63 0.44 2.19 1.96 0.68 3.10 0.97
2.00 0.96
13.35 6.13
3.14 2.98
4.18 3.00
11.5 11.0 10.5 10.0 9.5 9.0 8.5 8.0 7.5 7.0 6.5 6.0 5.5 5.0 4.5 4.0 3.5 3.0 2.5 2.0 1.5 1.0 0.5 0.0 -0.5
f1 (ppm)

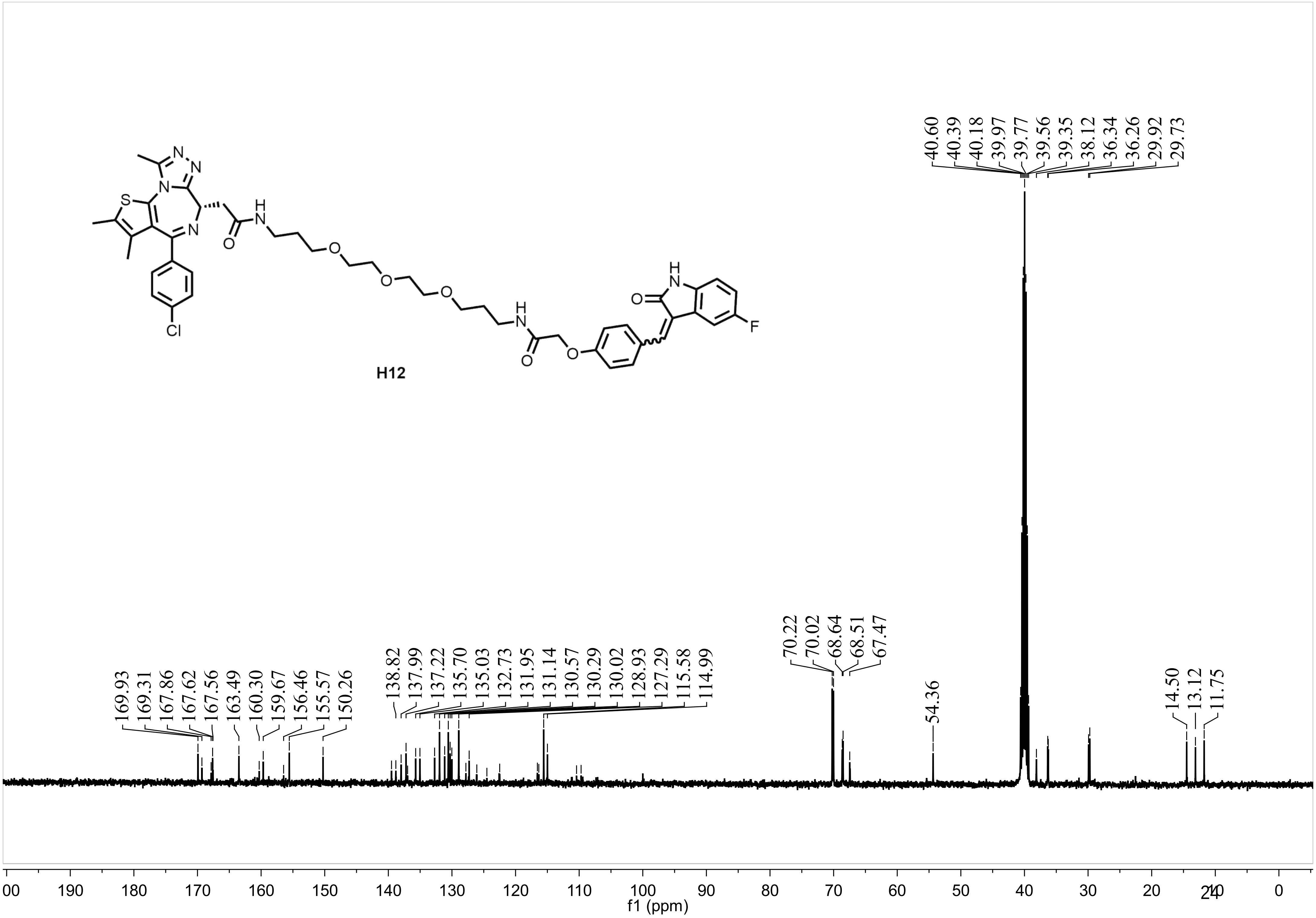

H12
169.93
169.31
167.86
167.62
167.56
163.49
160.30
159.67
156.46
155.57
150.26
138.82
137.99
137.22
135.70
135.03
132.73
131.95
131.14
130.57
130.29
130.02
128.93
127.29
115.58
114.99
70.22
70.02
68.64
68.51
67.47
54.36
40.60
40.39
40.18
39.97
39.77
39.56
39.35
38.12
36.34
36.26
29.92
29.73
14.50
13.12
11.75
f1 (ppm)

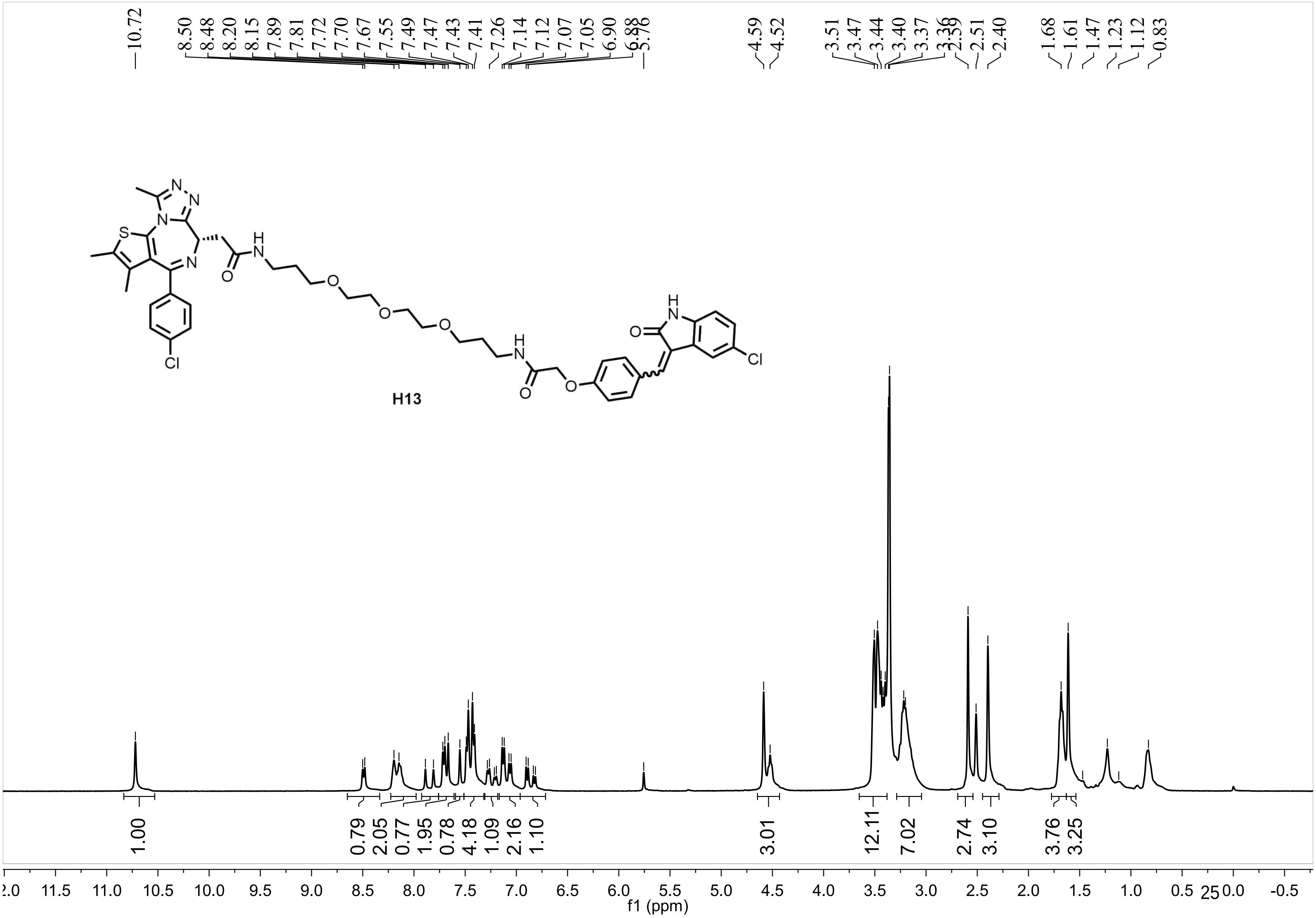
10.72
8.50 8.48 8.20 8.15 7.89 7.81 7.72 7.70 7.67 7.55 7.49 7.47 7.43 7.41 7.26 7.14 7.12 7.07 7.05 6.90 6.88 5.76
4.59 4.52
3.51 3.47 3.44 3.40 3.37 3.36 2.59 2.51 2.40
1.68 1.61 1.47 1.23 1.12 0.83
H13
1.00 0.79 2.05 0.77 1.95 0.78 4.18 1.09 2.16 1.10 3.01 12.11 7.02 2.74 3.10 3.76 3.25
2.0 11.5 11.0 10.5 10.0 9.5 9.0 8.5 8.0 7.5 7.0 6.5 6.0 5.5 5.0 4.5 4.0 3.5 3.0 2.5 2.0 1.5 1.0 0.5 0.0 -0.5
f1 (ppm)

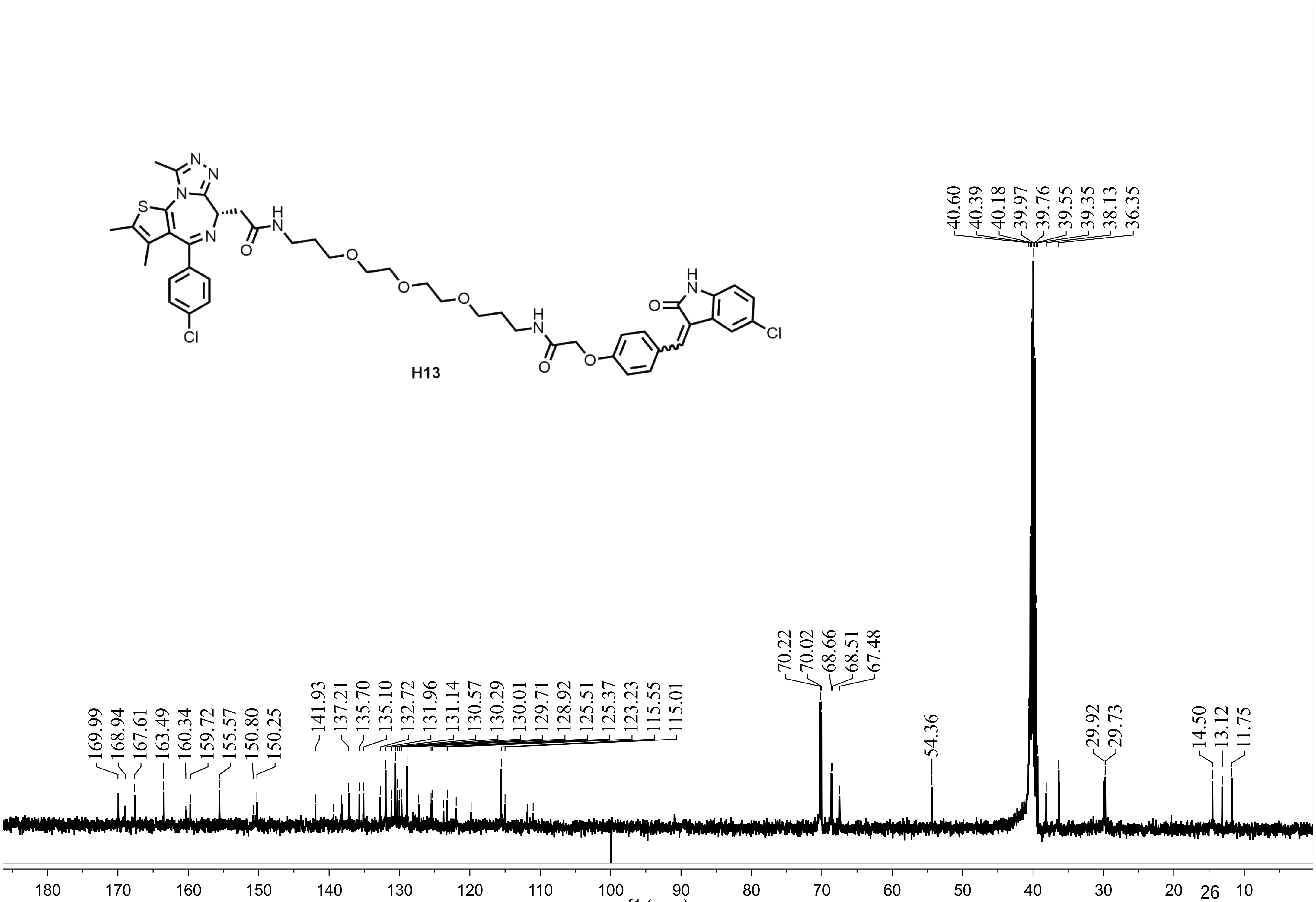
H13
169.99
168.94
167.61
163.49
160.34
159.72
155.57
150.80
150.25
141.93
137.21
135.70
135.10
132.72
131.96
131.14
130.57
130.29
130.01
129.71
128.92
125.51
125.37
123.23
115.55
115.01
70.22
70.02
68.66
68.51
67.48
54.36
40.60
40.39
40.18
39.97
39.76
39.55
39.35
38.13
36.35
29.92
29.73
14.50
13.12
11.75
f1 (ppm)

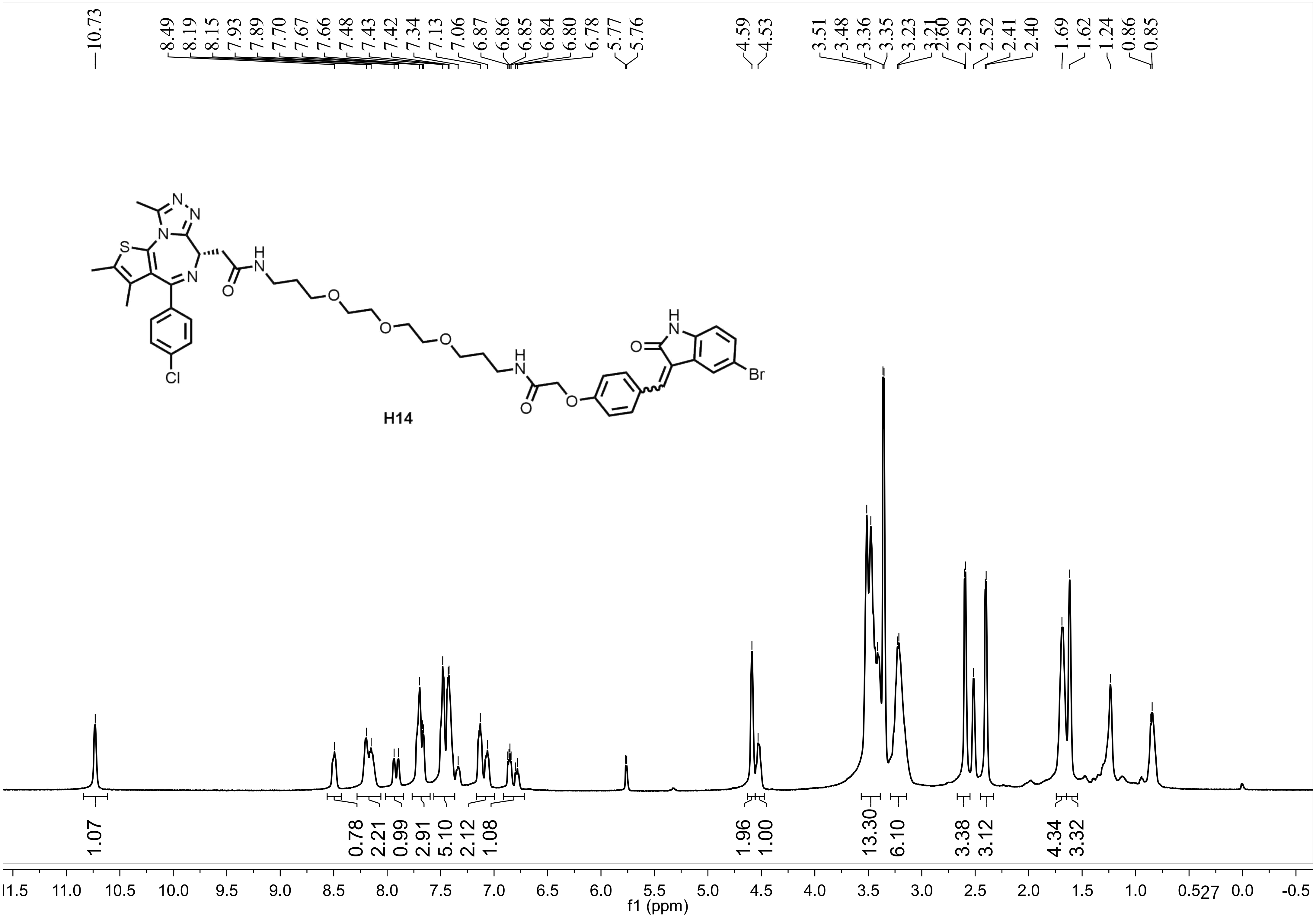
10.73
8.49 8.19 8.15 7.93 7.89 7.70 7.67 7.66 7.48 7.43 7.42 7.34 7.13 7.06 6.87 6.86 6.85 6.84 6.80 6.78 5.77 5.76
4.59 4.53
3.51 3.48 3.36 3.35 3.23 3.21 2.60 2.59 2.52 2.41 2.40 1.69 1.62 1.24 0.86 0.85
S
N
O
H
Cl
Br
H14
1.07
0.78 2.21 0.99 2.91 5.10 2.12 1.08
1.96 1.00
13.30 6.10
3.38 3.12
4.34 3.32
f1 (ppm)

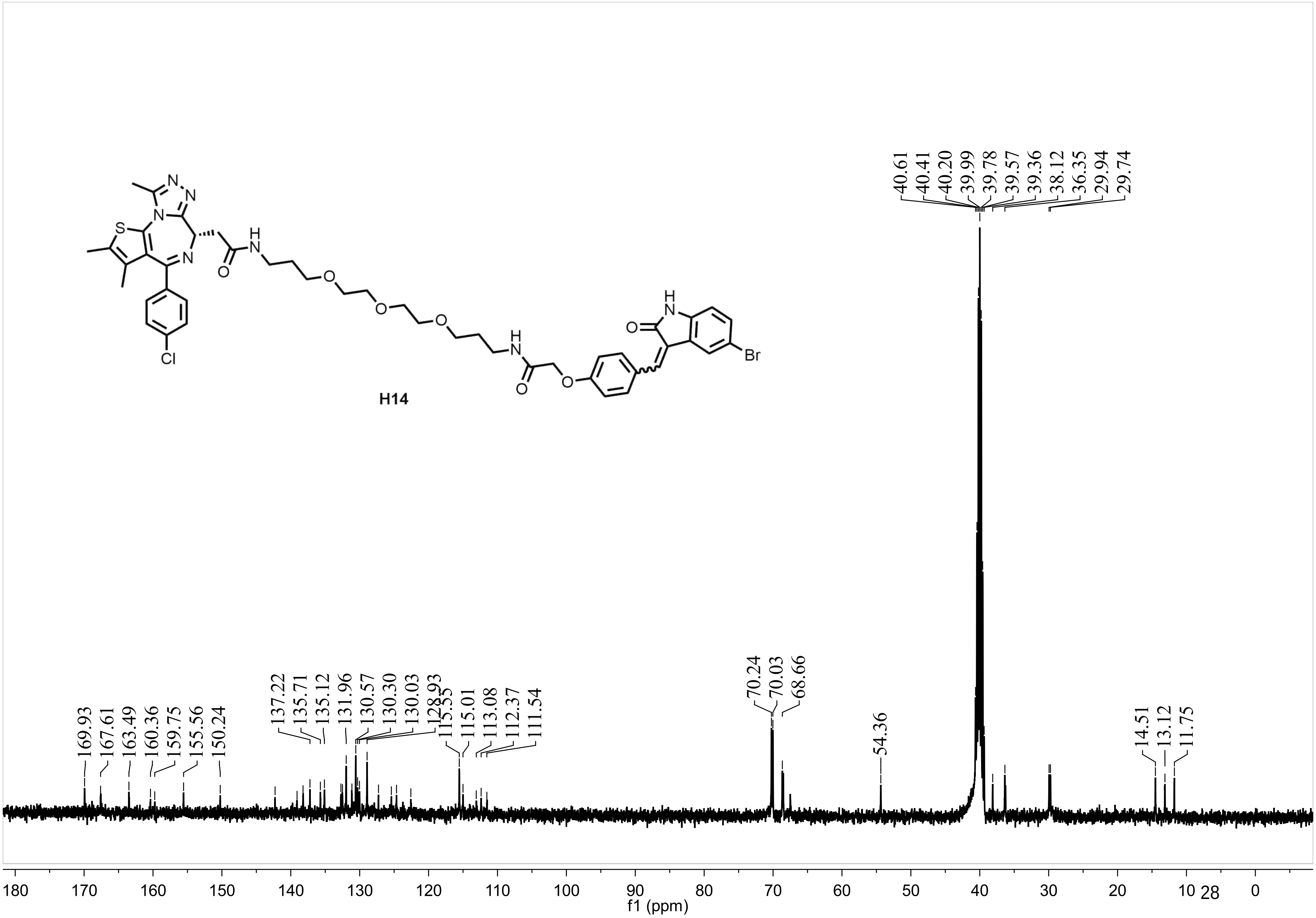
H14
40.61
40.41
40.20
39.99
39.78
39.57
39.36
38.12
36.35
29.94
29.74
169.93
167.61
163.49
160.36
159.75
155.56
150.24
137.22
135.71
135.12
131.96
130.57
130.30
130.03
128.93
115.55
115.01
113.08
112.37
111.54
70.24
70.03
68.66
54.36
14.51
13.12
11.75
180 170 160 150 140 130 120 110 100 90 80 70 60 50 40 30 20 10 0
f1 (ppm)

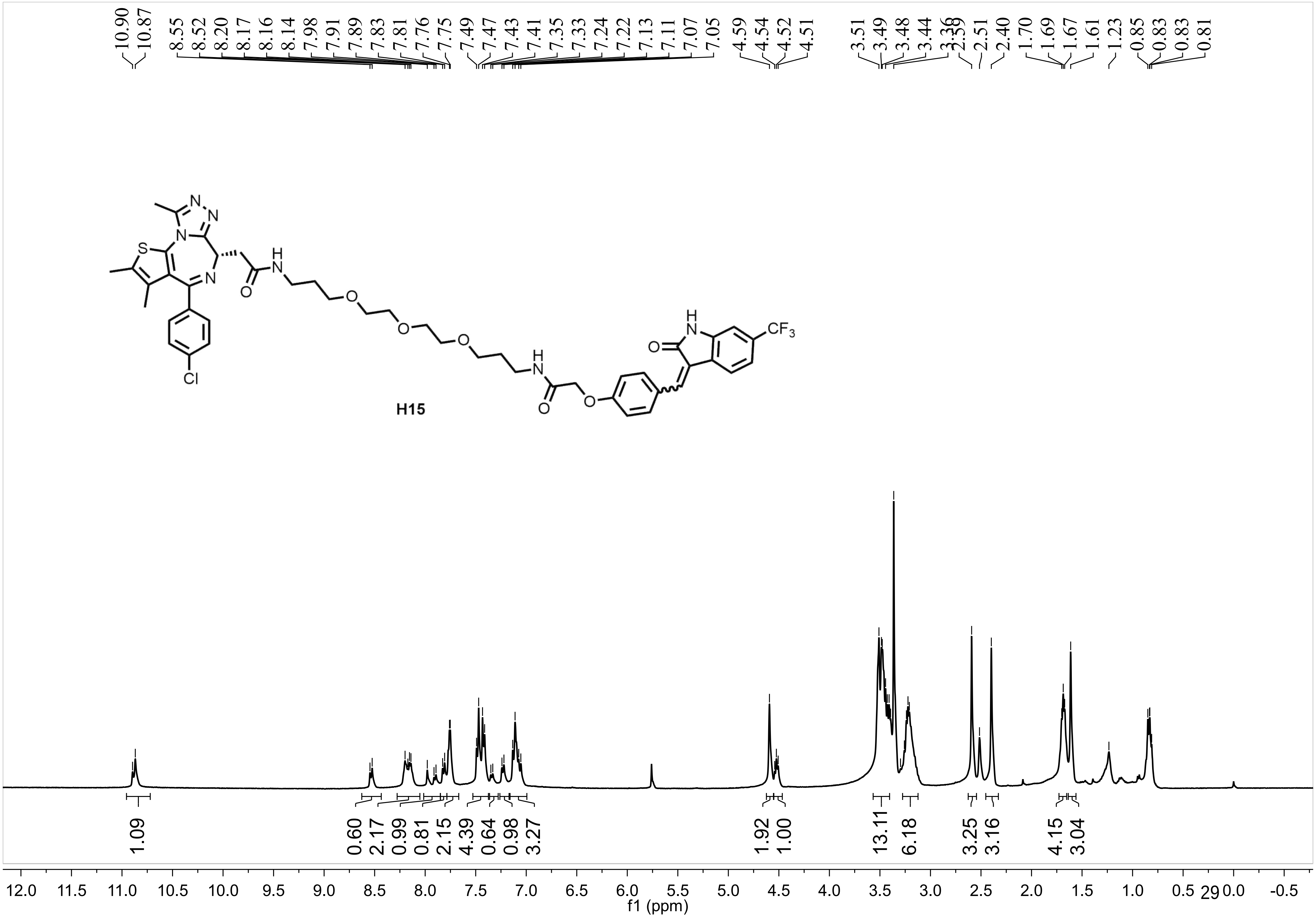
10.90
10.87
8.55
8.52
8.20
8.17
8.16
8.14
7.98
7.91
7.89
7.83
7.81
7.76
7.75
7.49
7.47
7.43
7.41
7.35
7.33
7.24
7.22
7.13
7.11
7.07
7.05
4.59
4.54
4.52
4.51
3.51
3.49
3.48
3.44
3.36
2.59
2.51
2.40
1.70
1.69
1.67
1.61
1.23
0.85
0.83
0.83
0.81
N
N
N
S
N
H
N
O
O
O
O
Cl
H
N
O
O
H
N
O
CF3
H15
1.09
0.60
2.17
0.99
0.81
2.15
4.39
0.64
0.98
3.27
1.92
1.00
13.11
6.18
3.25
3.16
4.15
3.04
12.0
11.5
11.0
10.5
10.0
9.5
9.0
8.5
8.0
7.5
7.0
6.5
6.0
5.5
5.0
4.5
4.0
3.5
3.0
2.5
2.0
1.5
1.0
0.5
0.0
-0.5
f1 (ppm)

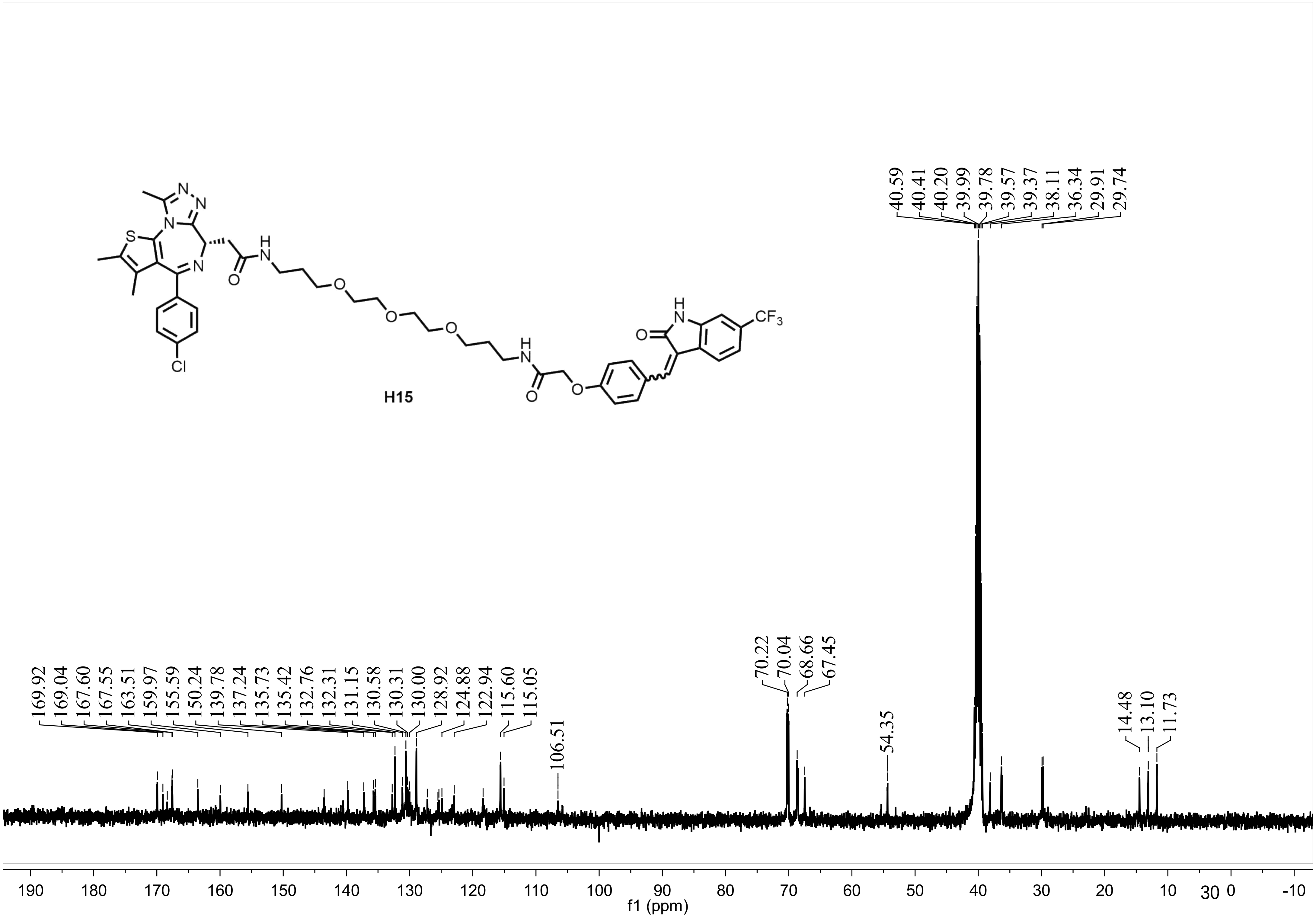
H15
CF3
Cl
40.59
40.41
40.20
39.99
39.78
39.57
39.37
38.11
36.34
29.91
29.74
169.92
169.04
167.60
167.55
163.51
159.97
155.59
150.24
139.78
137.24
135.73
135.42
132.76
132.31
131.15
130.58
130.31
130.00
128.92
124.88
122.94
115.60
115.05
106.51
70.22
70.04
68.66
67.45
54.35
14.48
13.10
11.73
190 180 170 160 150 140 130 120 110 100 90 80 70 60 50 40 30 20 10 0 -10
f1 (ppm)

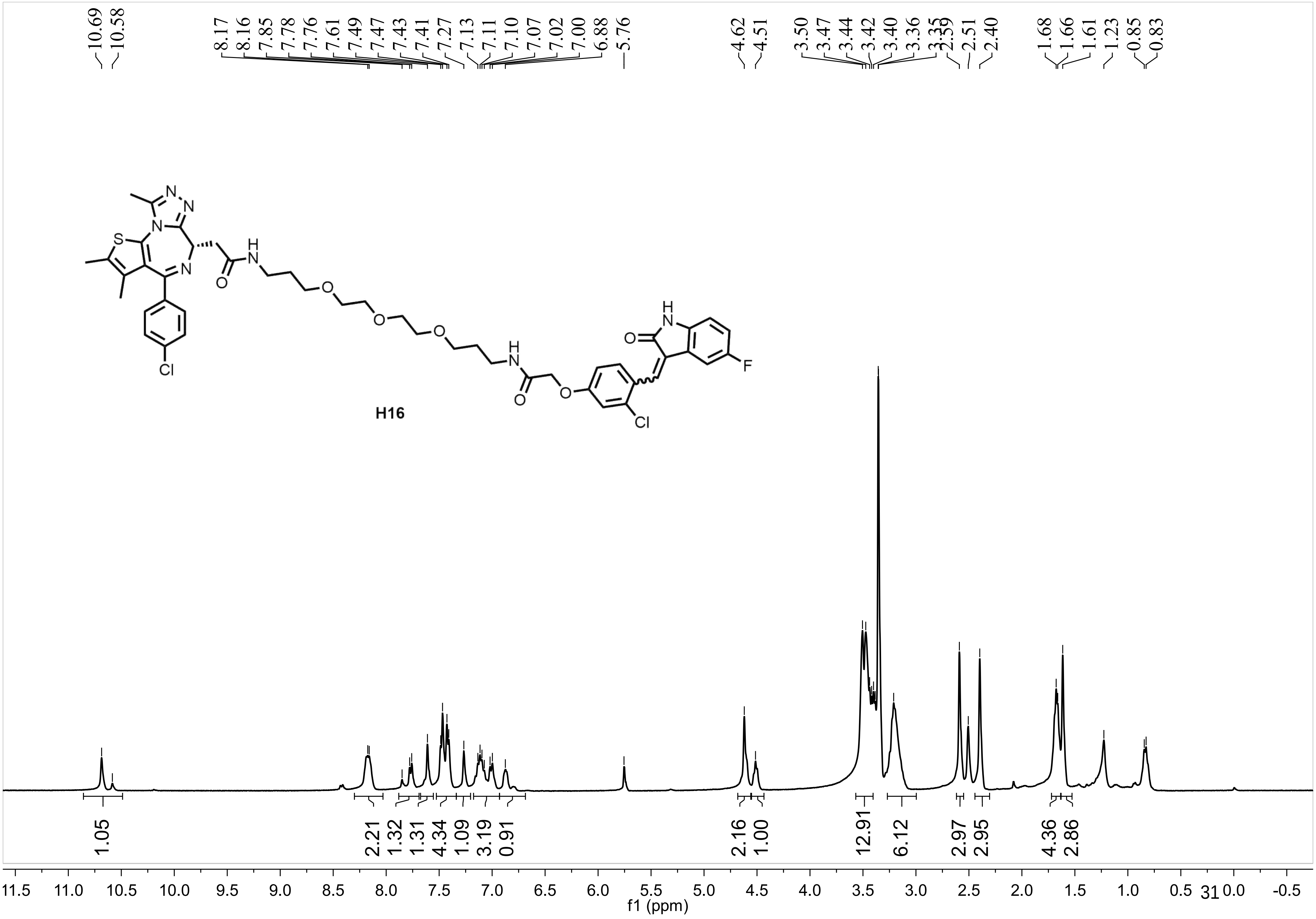
10.69
10.58
8.17
8.16
7.85
7.78
7.76
7.61
7.49
7.47
7.43
7.41
7.27
7.13
7.11
7.10
7.07
7.02
7.00
6.88
5.76
4.62
4.51
3.50
3.47
3.44
3.42
3.40
3.36
3.35
2.59
2.51
2.40
1.68
1.66
1.61
1.23
0.85
0.83
H16
1.05
2.21
1.32
1.31
4.34
1.09
3.19
0.91
2.16
1.00
12.91
6.12
2.97
2.95
4.36
2.86
f1 (ppm)

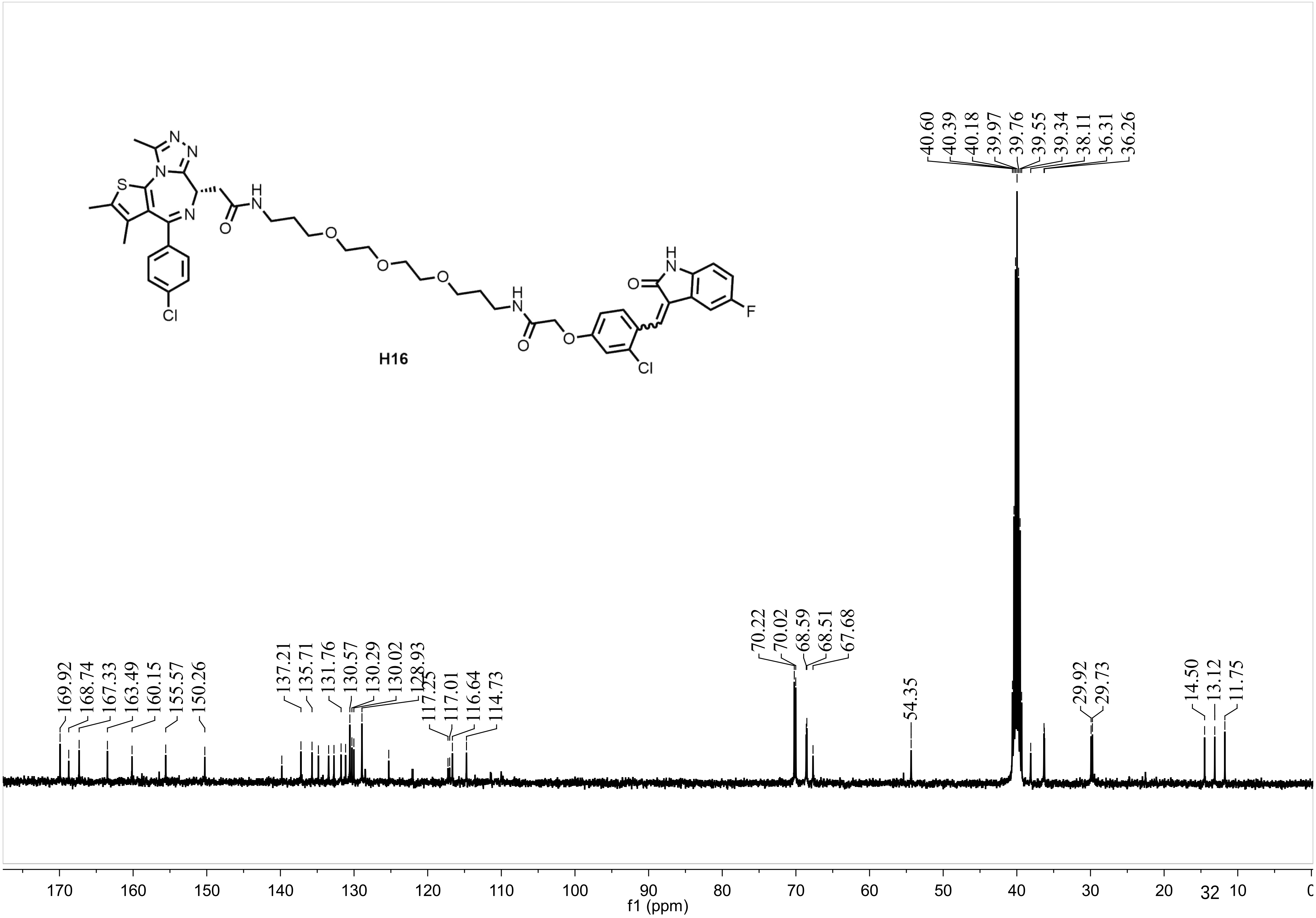

H16
169.92
168.74
167.33
163.49
160.15
155.57
150.26
137.21
135.71
131.76
130.57
130.29
130.02
128.93
117.25
117.01
116.64
114.73
70.22
70.02
68.59
68.51
67.68
54.35
40.60
40.39
40.18
39.97
39.76
39.55
39.34
38.11
36.31
36.26
29.92
29.73
14.50
13.12
11.75
170 160 150 140 130 120 110 100 90 80 70 60 50 40 30 20 10
f1 (ppm)

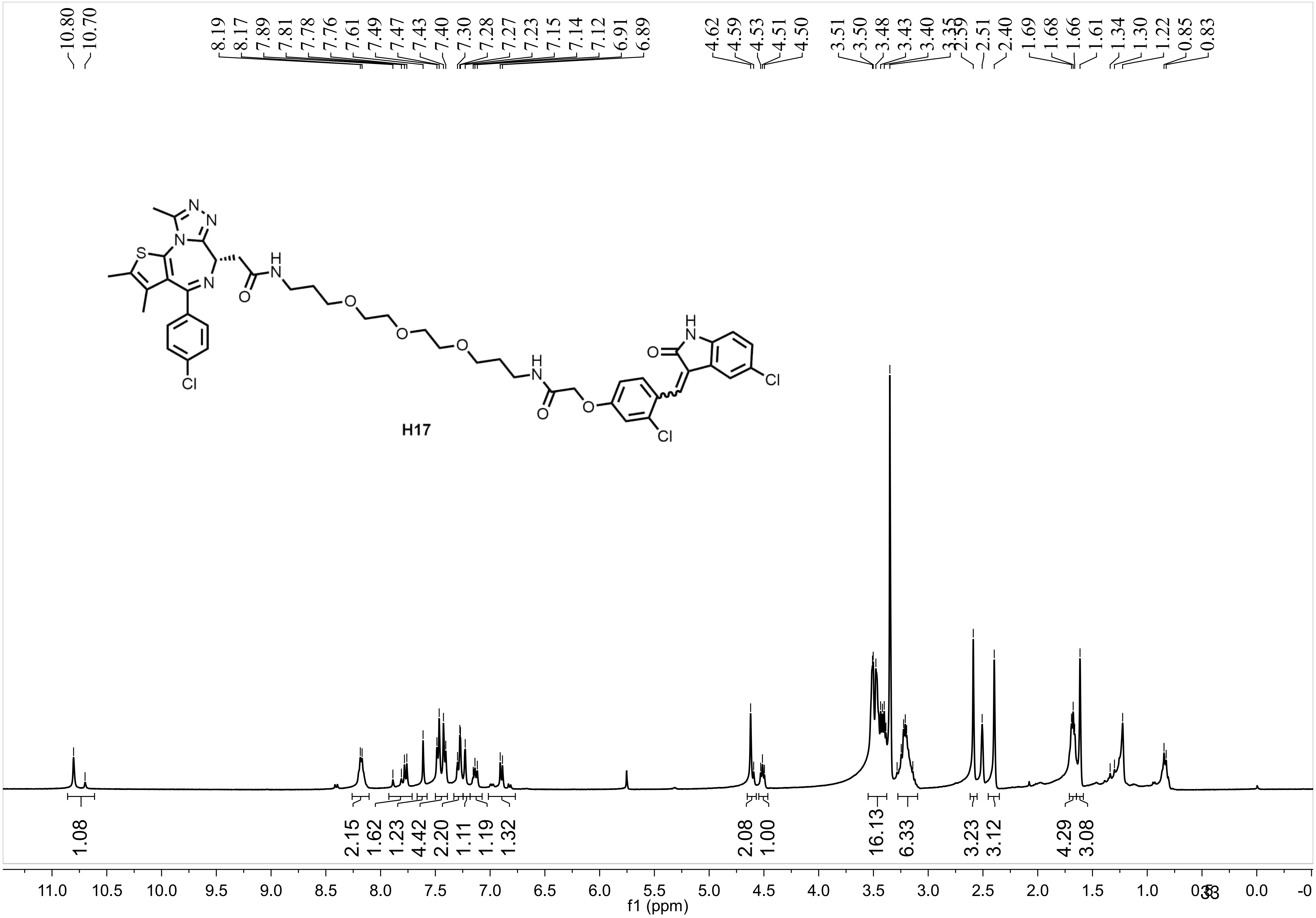
10.80
10.70
8.19
8.17
7.89
7.81
7.78
7.76
7.61
7.49
7.47
7.43
7.40
7.30
7.28
7.27
7.23
7.15
7.14
7.12
6.91
6.89
4.62
4.59
4.53
4.51
4.50
3.51
3.50
3.48
3.43
3.40
3.35
2.59
2.51
2.40
1.69
1.68
1.66
1.61
1.34
1.30
1.22
0.85
0.83
H17
1.08
2.15
1.62
1.23
4.42
2.20
1.11
1.19
1.32
2.08
1.00
16.13
6.33
3.23
3.12
4.29
3.08
f1 (ppm)

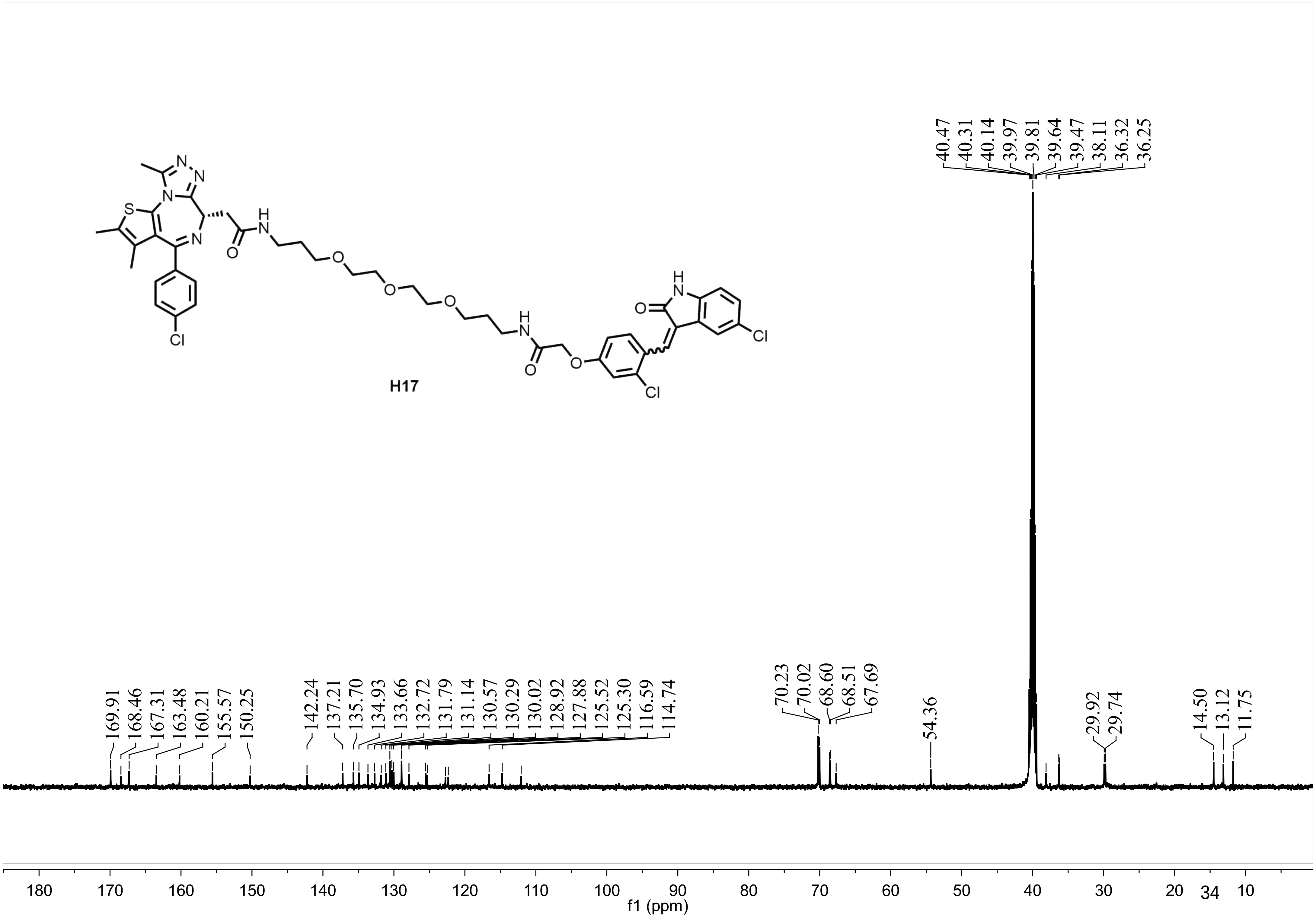
H17
40.47
40.31
40.14
39.97
39.81
39.64
39.47
38.11
36.32
36.25
169.91
168.46
167.31
163.48
160.21
155.57
150.25
142.24
137.21
135.70
134.93
133.66
132.72
131.79
131.14
130.57
130.29
130.02
128.92
127.88
125.52
125.30
116.59
114.74
70.23
70.02
68.60
68.51
67.69
54.36
29.92
29.74
14.50
13.12
11.75
180 170 160 150 140 130 120 110 100 90 80 70 60 50 40 30 20 10
f1 (ppm)

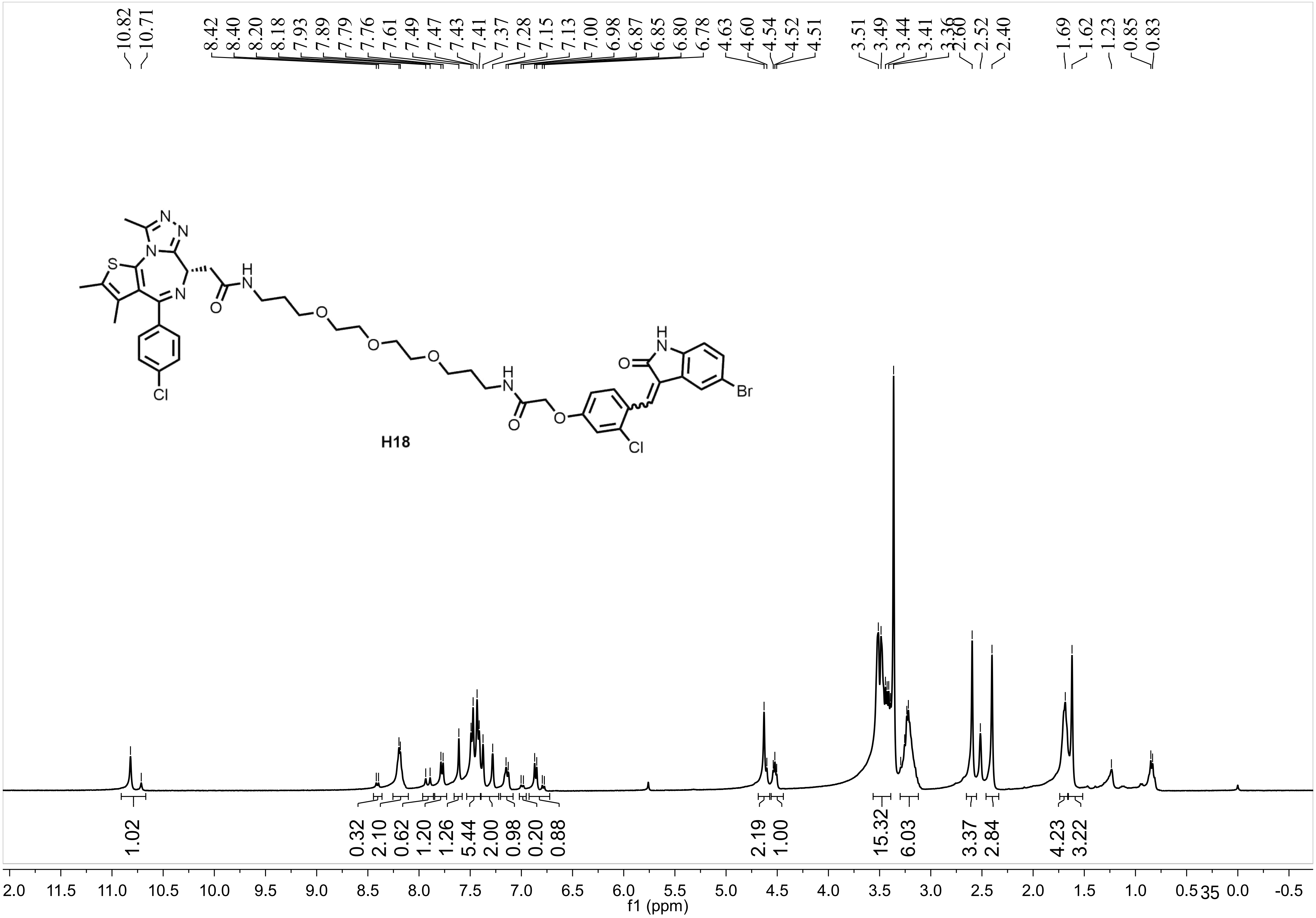
10.82
10.71
8.42
8.40
8.20
8.18
7.93
7.89
7.79
7.76
7.61
7.49
7.47
7.43
7.41
7.37
7.28
7.15
7.13
7.00
6.98
6.87
6.85
6.80
6.78
4.63
4.60
4.54
4.52
4.51
3.51
3.49
3.44
3.41
3.36
2.60
2.52
2.40
1.69
1.62
1.23
0.85
0.83
H18
Br
Cl
1.02
0.32
2.10
0.62
1.20
1.26
5.44
2.00
0.98
0.20
0.88
2.19
1.00
15.32
6.03
3.37
2.84
4.23
3.22
f1 (ppm)

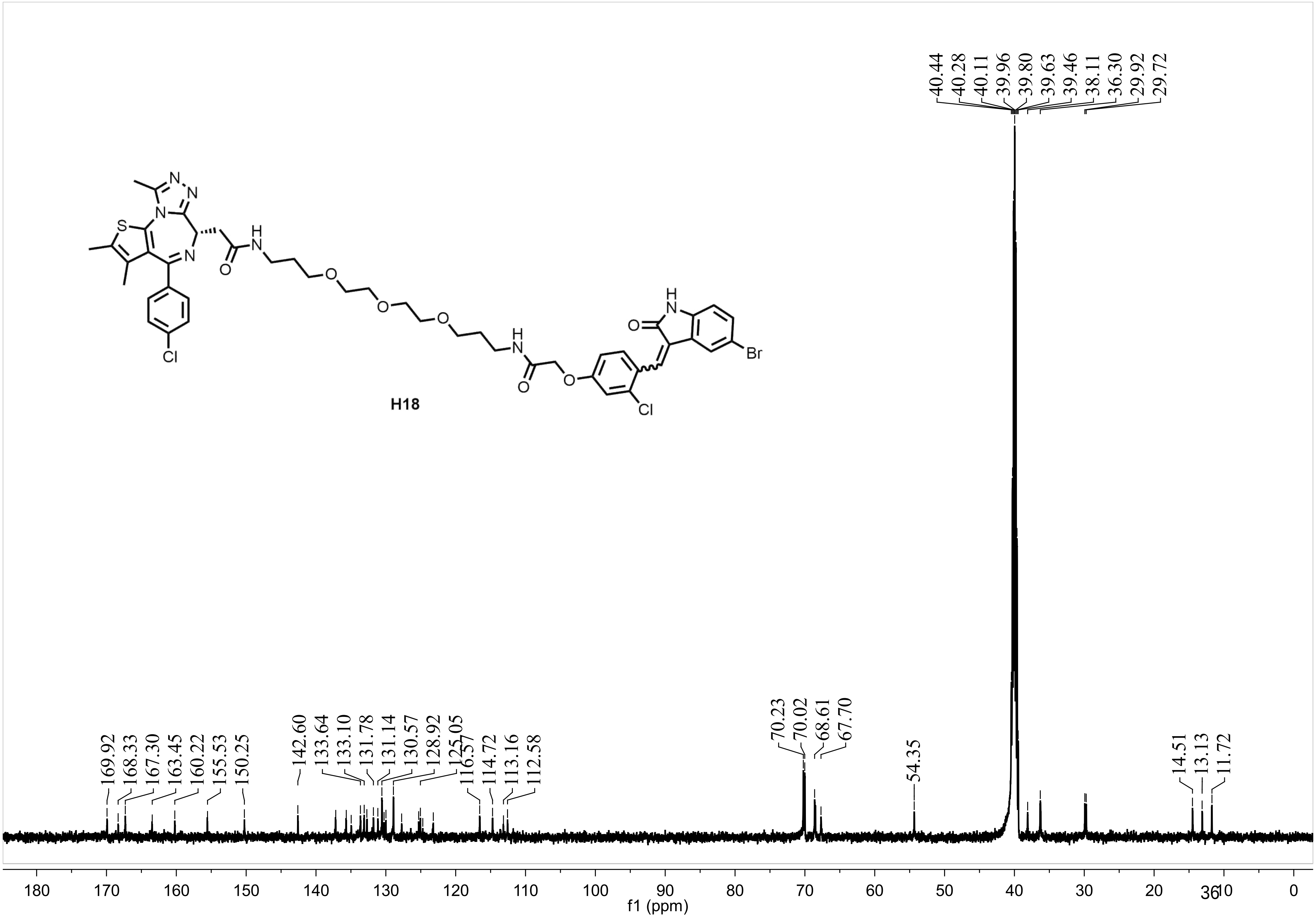
N
N
S
N
N
H
N
O
O
O
O
Cl
H
N
O
O
H
N
O
Br
Cl
H18
40.44
40.28
40.11
39.96
39.80
39.63
39.46
38.11
36.30
29.92
29.72
169.92
168.33
167.30
163.45
160.22
155.53
150.25
142.60
133.64
133.10
131.78
131.14
130.57
128.92
125.05
116.57
114.72
113.16
112.58
70.23
70.02
68.61
67.70
54.35
14.51
13.13
11.72
180 170 160 150 140 130 120 110 100 90 80 70 60 50 40 30 20 10 0
f1 (ppm)

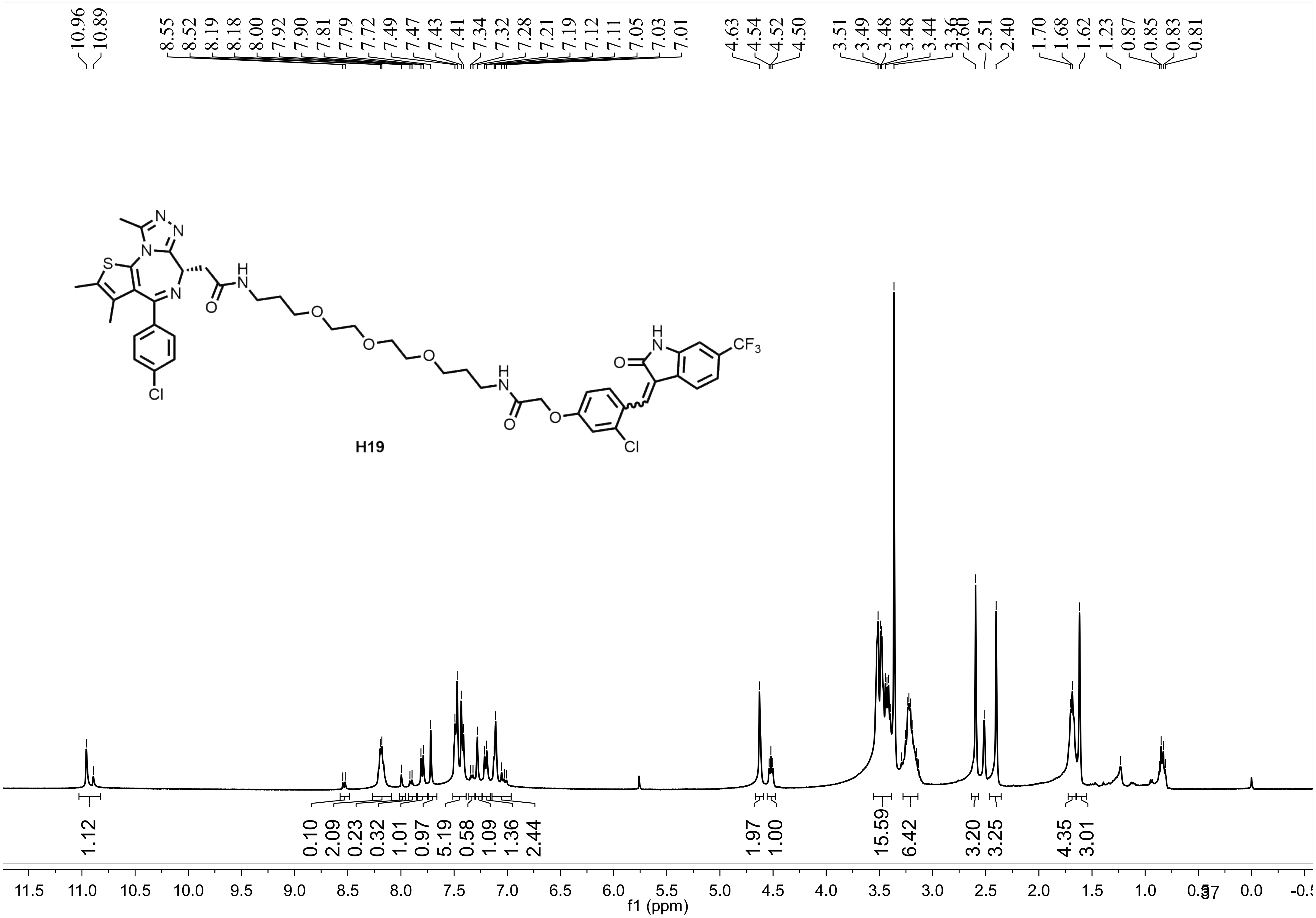
10.96
10.89
8.55
8.52
8.19
8.18
8.00
7.92
7.90
7.81
7.79
7.72
7.49
7.47
7.43
7.41
7.34
7.32
7.28
7.21
7.19
7.12
7.11
7.05
7.03
7.01
4.63
4.54
4.52
4.50
3.51
3.49
3.48
3.48
3.44
3.36
2.60
2.51
2.40
1.70
1.68
1.62
1.23
0.87
0.85
0.83
0.81
H19
1.12
0.10
2.09
0.23
0.32
1.01
0.97
5.19
0.58
1.09
1.36
2.44
1.97
1.00
15.59
6.42
3.20
3.25
4.35
3.01
f1 (ppm)

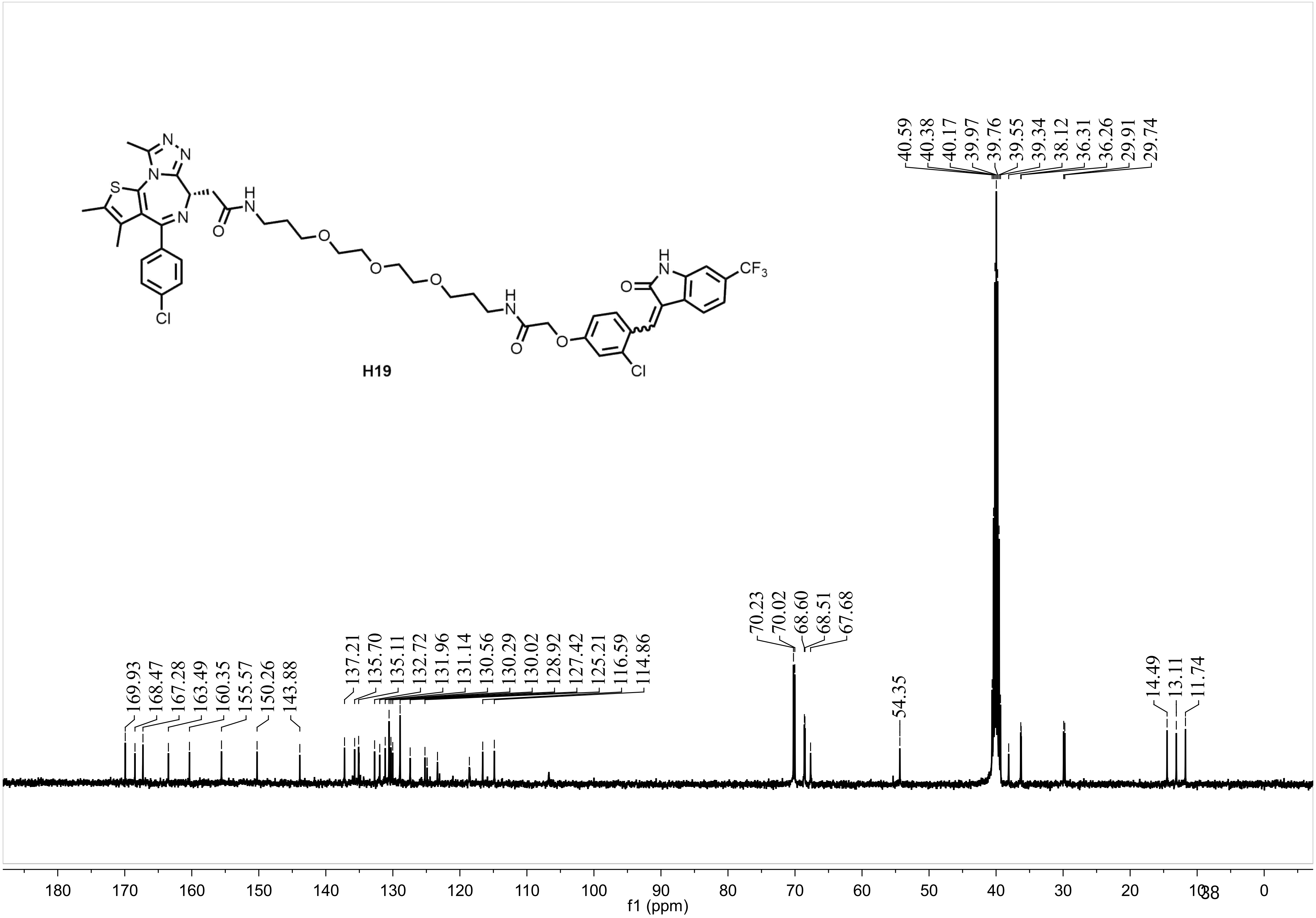
H19
CF3
Cl
169.93
168.47
167.28
163.49
160.35
155.57
150.26
143.88
137.21
135.70
135.11
132.72
131.96
131.14
130.56
130.29
130.02
128.92
127.42
125.21
116.59
114.86
70.23
70.02
68.60
68.51
67.68
54.35
40.59
40.38
40.17
39.97
39.76
39.55
39.34
38.12
36.31
36.26
29.91
29.74
14.49
13.11
11.74
f1 (ppm)

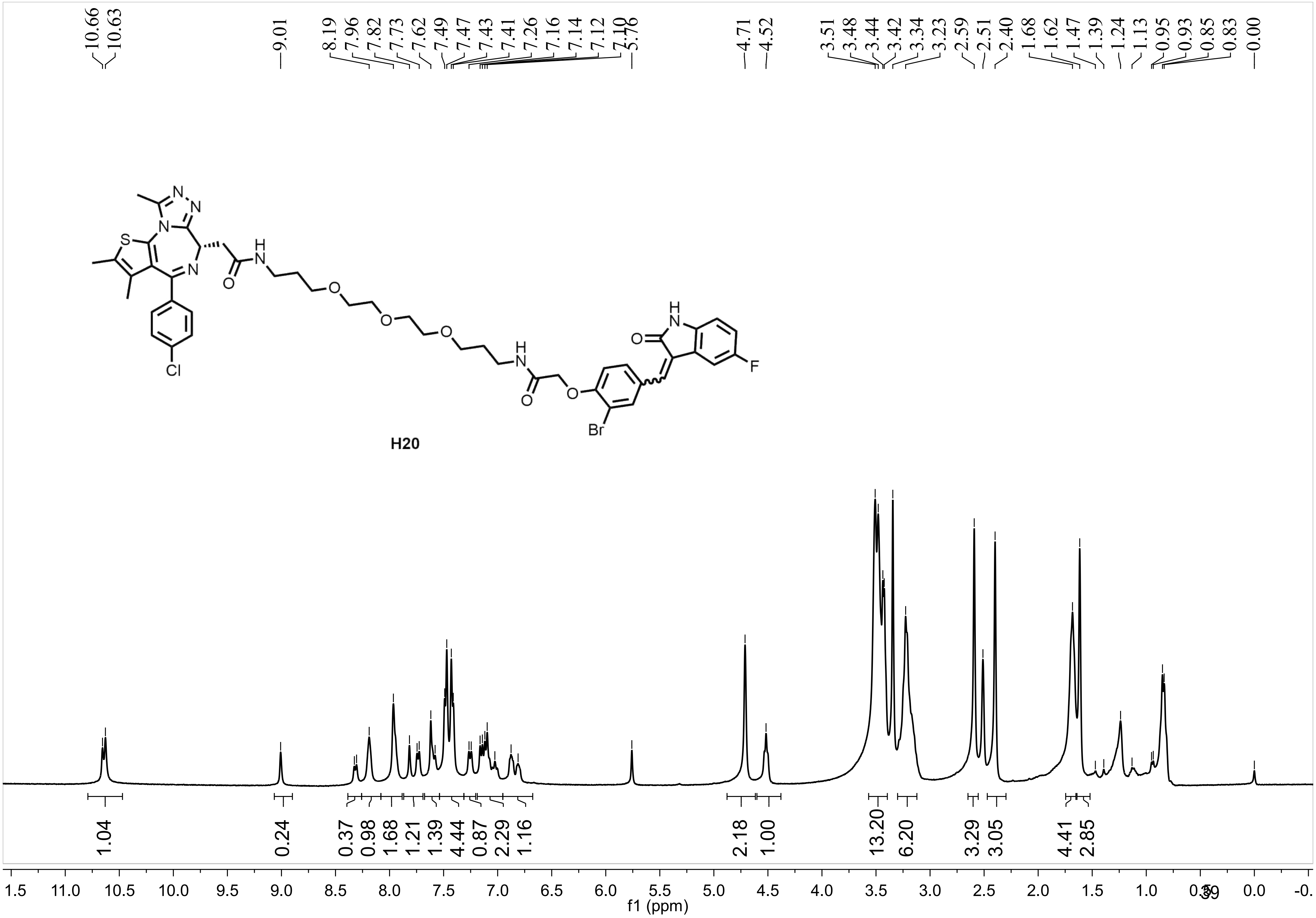
10.66
10.63
9.01
8.19
7.96
7.82
7.73
7.62
7.49
7.47
7.43
7.41
7.26
7.16
7.14
7.12
7.10
5.76
4.71
4.52
3.51
3.48
3.44
3.42
3.34
3.23
2.59
2.51
2.40
1.68
1.62
1.47
1.39
1.24
1.13
0.95
0.93
0.85
0.83
0.00
S
N
N
N
N
H
N
O
O
O
O
H
N
O
O
Cl
Br
H
N
O
F
H20
1.04
0.24
0.37
0.98
1.68
1.21
1.39
4.44
0.87
2.29
1.16
2.18
1.00
13.20
6.20
3.29
3.05
4.41
2.85
f1 (ppm)

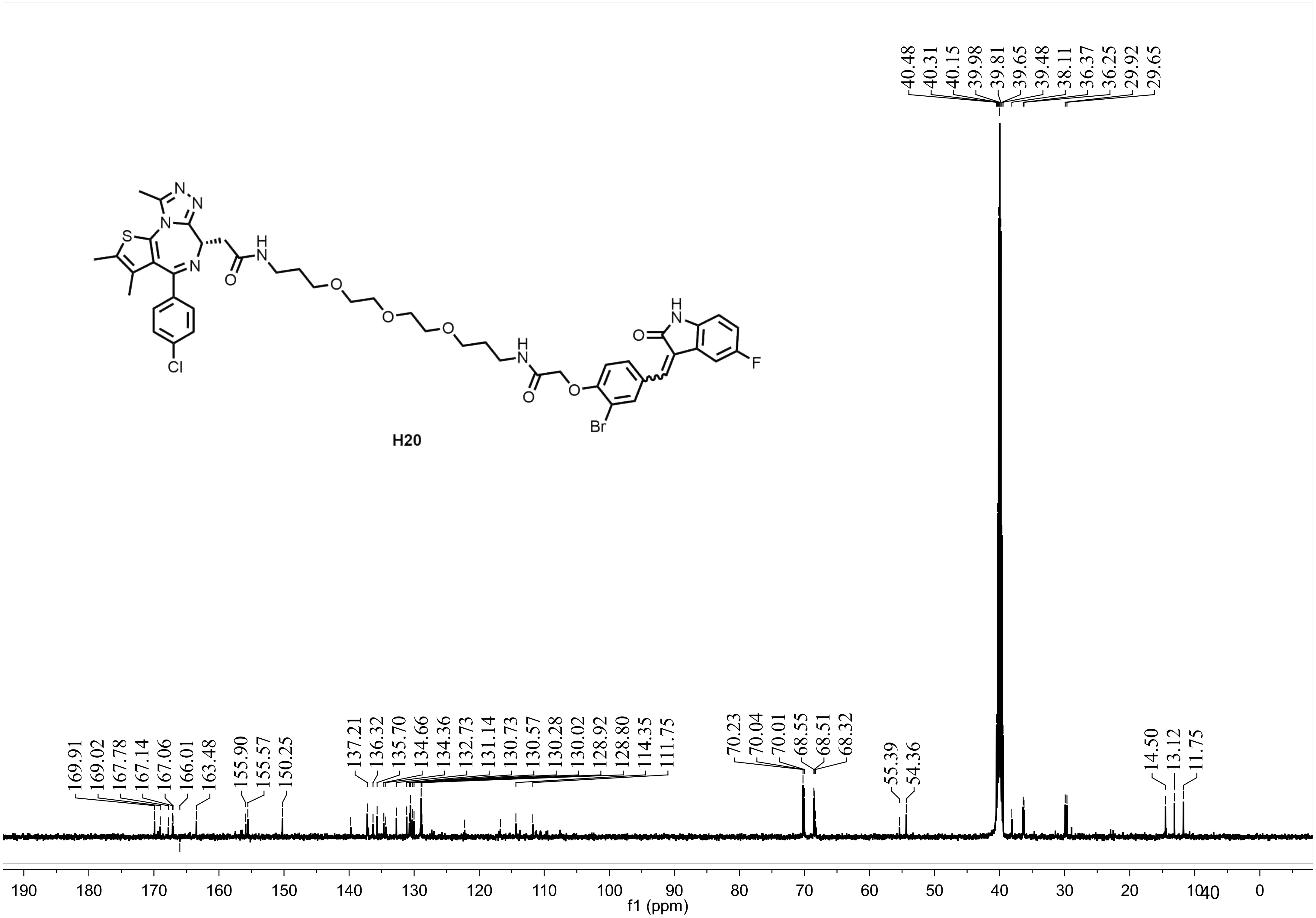
40.48
40.31
40.15
39.98
39.81
39.65
39.48
38.11
36.37
36.25
29.92
29.65
H20
169.91
169.02
167.78
167.14
167.06
166.01
163.48
155.90
155.57
150.25
137.21
136.32
135.70
134.66
134.36
132.73
131.14
130.73
130.57
130.28
130.02
128.92
128.80
114.35
111.75
70.23
70.04
70.01
68.55
68.51
68.32
55.39
54.36
14.50
13.12
11.75
190 180 170 160 150 140 130 120 110 100 90 80 70 60 50 40 30 20 10 0
f1 (ppm)

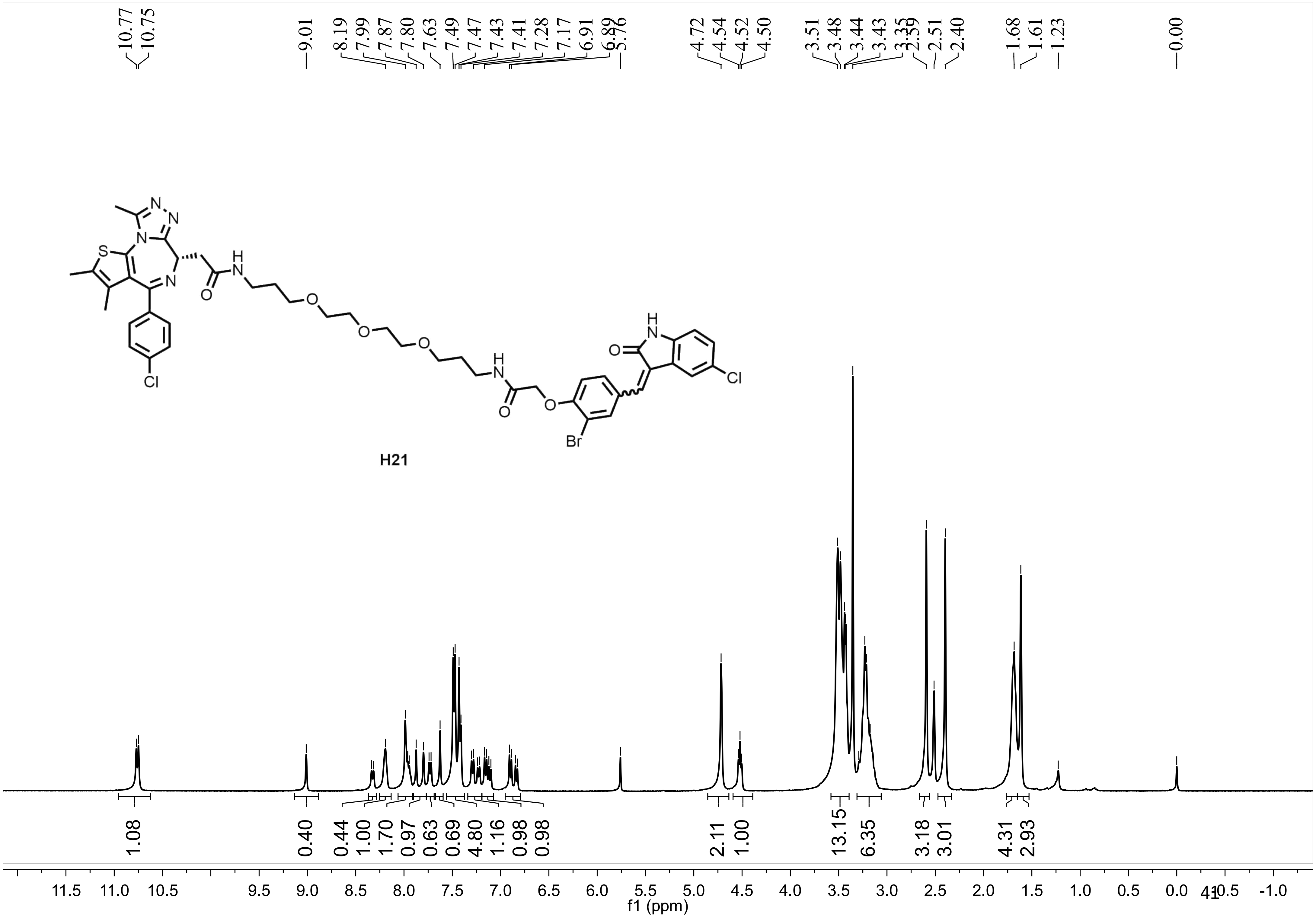
10.77
10.75
9.01
8.19
7.99
7.87
7.80
7.63
7.49
7.47
7.43
7.41
7.28
7.17
6.91
6.89
5.76
4.72
4.54
4.52
4.50
3.51
3.48
3.44
3.43
3.35
2.59
2.51
2.40
1.68
1.61
1.23
0.00
N
N
N
S
N
H
N
O
O
O
O
Cl
H
N
O
Cl
H
N
O
O
Br
H21
1.08
0.40
0.44
1.00
1.70
0.97
0.63
0.69
4.80
1.16
0.98
0.98
2.11
1.00
13.15
6.35
3.18
3.01
4.31
2.93
11.5 11.0 10.5 9.5 9.0 8.5 8.0 7.5 7.0 6.5 6.0 5.5 5.0 4.5 4.0 3.5 3.0 2.5 2.0 1.5 1.0 0.5 0.0 -0.5 -1.0
f1 (ppm)

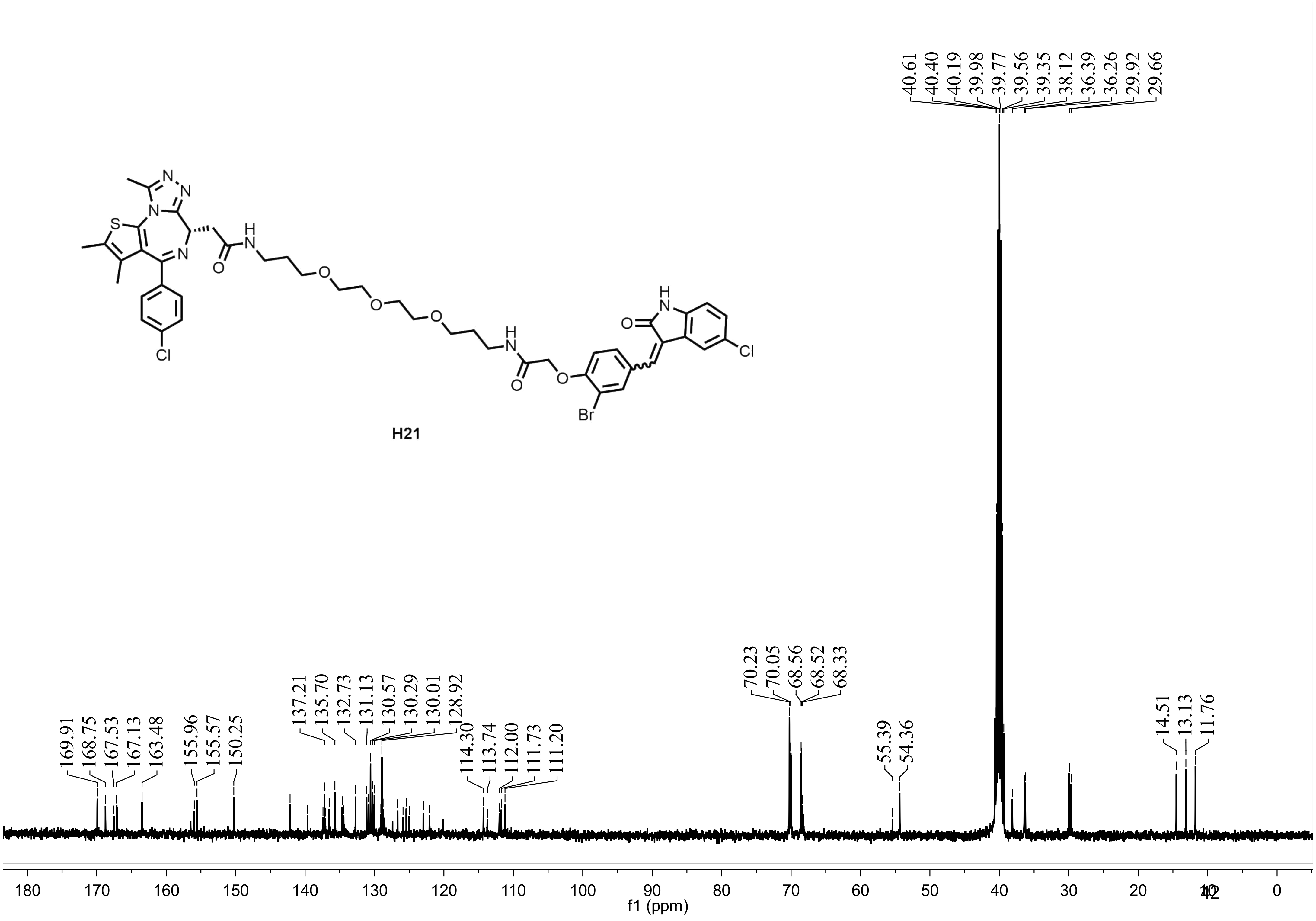
H21
169.91
168.75
167.53
167.13
163.48
155.96
155.57
150.25
137.21
135.70
132.73
131.13
130.57
130.29
130.01
128.92
114.30
113.74
112.00
111.73
111.20
70.23
70.05
68.56
68.52
68.33
55.39
54.36
40.61
40.40
40.19
39.98
39.77
39.56
39.35
38.12
36.39
36.26
29.92
29.66
14.51
13.13
11.76
180 170 160 150 140 130 120 110 100 90 80 70 60 50 40 30 20 10 0
f1 (ppm)

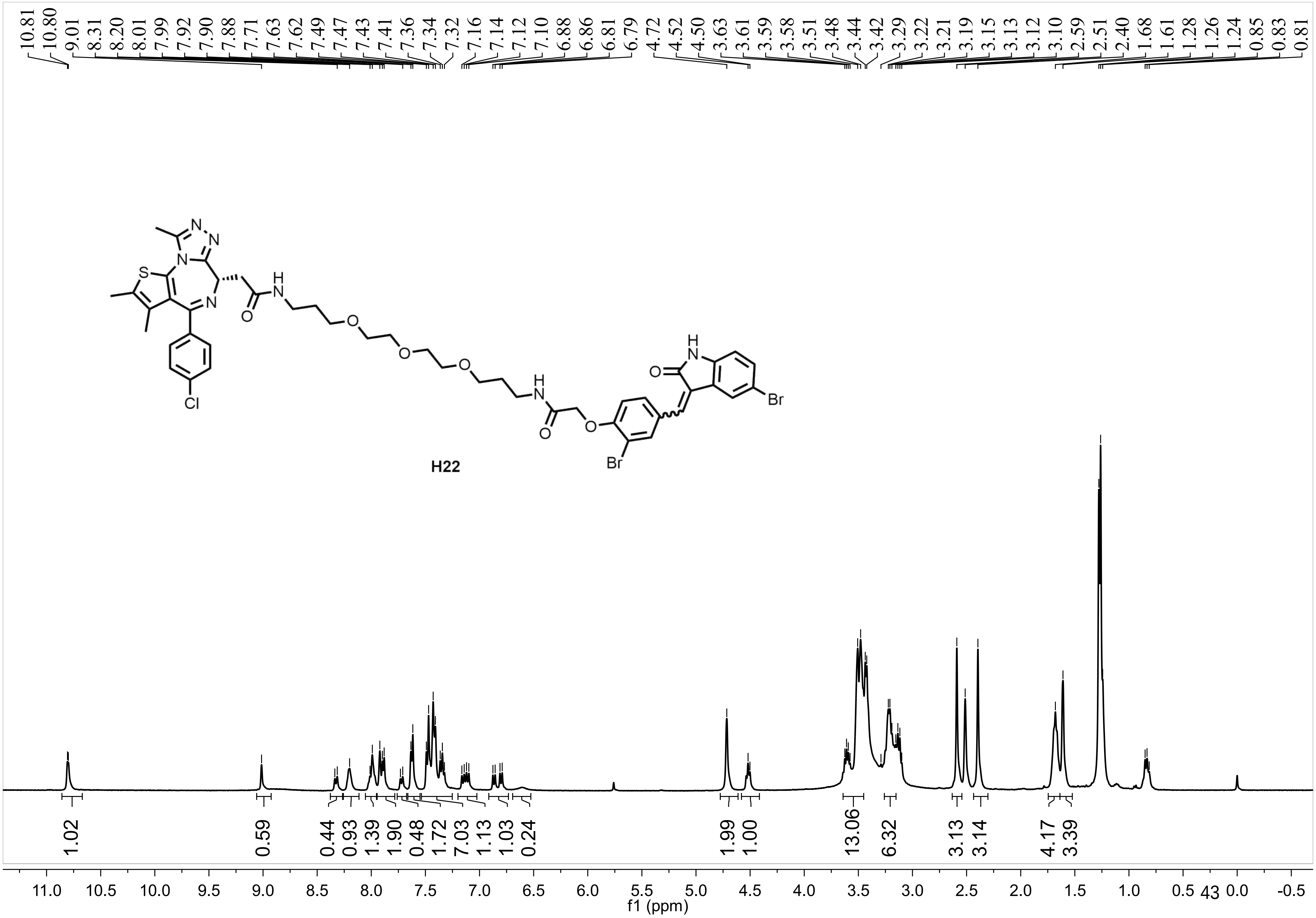
10.81 10.80 9.01 8.31 8.20 8.01 7.99 7.92 7.90 7.88 7.71 7.63 7.62 7.49 7.47 7.43 7.41 7.36 7.34 7.32 7.16 7.14 7.12 7.10 6.88 6.86 6.81 6.79 4.72 4.52 4.50 3.63 3.61 3.59 3.58 3.51 3.48 3.44 3.42 3.29 3.22 3.21 3.19 3.15 3.13 3.12 3.10 2.59 2.51 2.40 1.68 1.61 1.28 1.26 1.24 0.85 0.83 0.81
H22
1.02 0.59 0.44 0.93 1.39 1.90 0.48 1.72 7.03 1.13 1.03 0.24 1.99 1.00 13.06 6.32 3.13 3.14 4.17 3.39
11.0 10.5 10.0 9.5 9.0 8.5 8.0 7.5 7.0 6.5 6.0 5.5 5.0 4.5 4.0 3.5 3.0 2.5 2.0 1.5 1.0 0.5 0.0 -0.5
f1 (ppm)

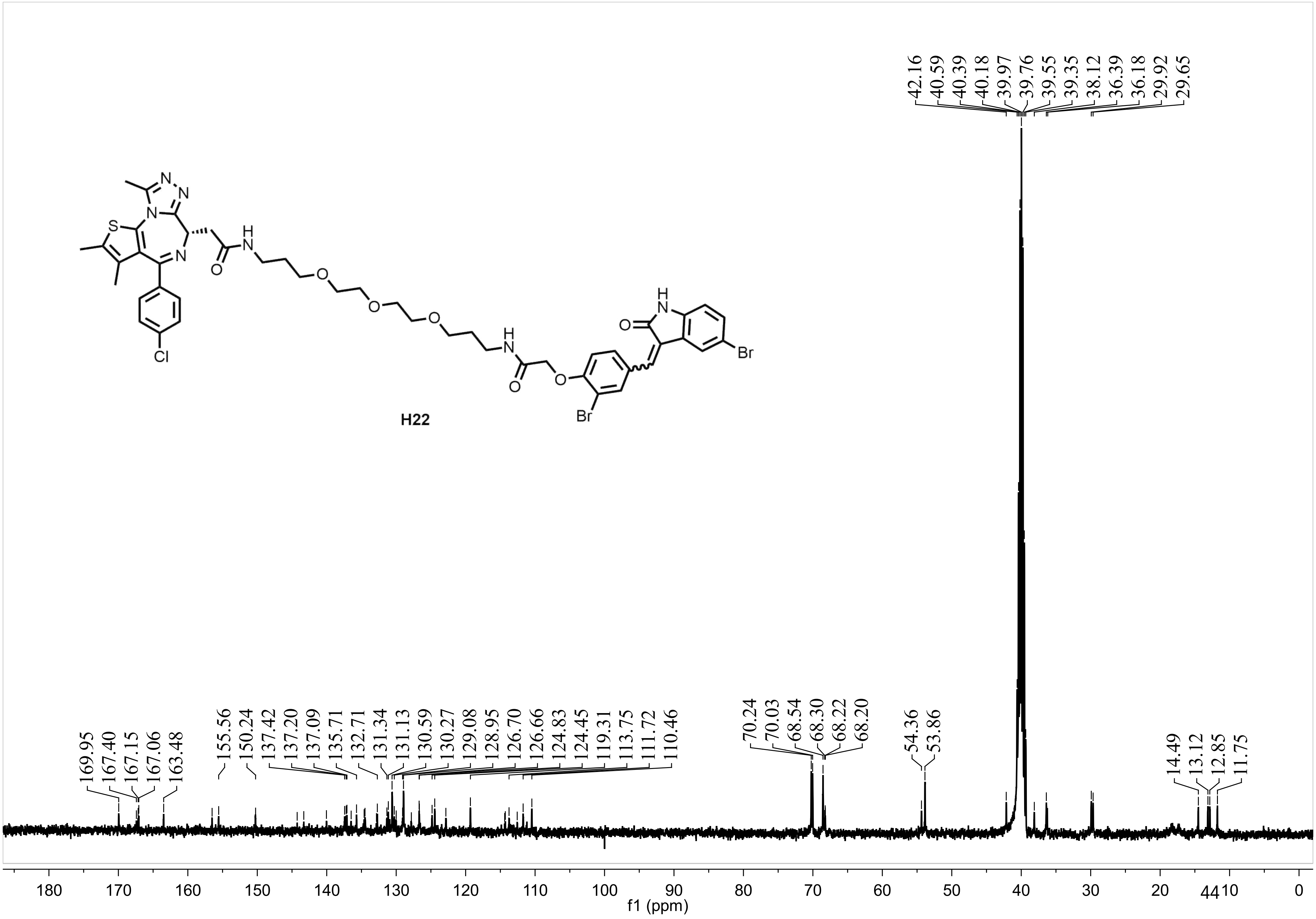

H22
42.16 40.59 40.39 40.18 39.97 39.76 39.55 39.35 38.12 36.39 36.18 29.92 29.65
169.95 167.40 167.15 167.06 163.48
155.56 150.24
137.42 137.20 137.09 135.71 132.71 131.34 131.13 130.59 130.27 129.08 128.95 126.70 126.66 124.83 124.45 119.31 113.75 111.72 110.46
70.24 70.03 68.54 68.30 68.22 68.20
54.36 53.86
14.49 13.12 12.85 11.75
180 170 160 150 140 130 120 110 100 90 80 70 60 50 40 30 20 10 0
f1 (ppm)

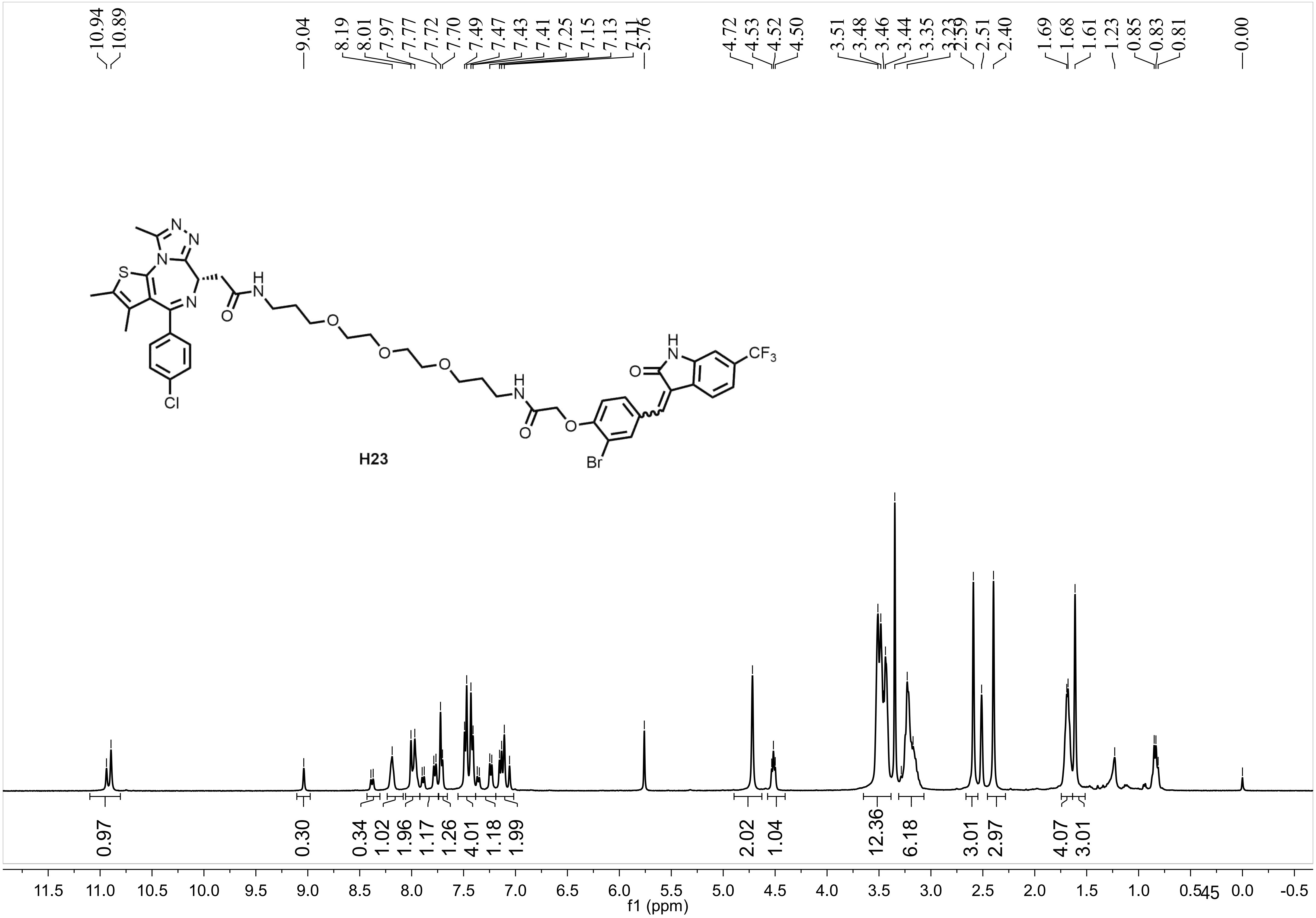
10.94
10.89
9.04
8.19
8.01
7.97
7.77
7.72
7.70
7.49
7.47
7.43
7.41
7.25
7.15
7.13
7.11
5.76
4.72
4.53
4.52
4.50
3.51
3.48
3.46
3.44
3.35
3.23
2.59
2.51
2.40
1.69
1.68
1.61
1.23
0.85
0.83
0.81
0.00
N
N
N
S
N
H
N
O
O
O
O
H
N
O
O
Br
Cl
H
N
O
$CF_3$
0.97
0.30
0.34
1.02
1.96
1.17
1.26
4.01
1.18
1.99
2.02
1.04
12.36
6.18
3.01
2.97
4.07
3.01
11.5 11.0 10.5 10.0 9.5 9.0 8.5 8.0 7.5 7.0 6.5 6.0 5.5 5.0 4.5 4.0 3.5 3.0 2.5 2.0 1.5 1.0 0.5 0.0 -0.5
f1 (ppm)

**H23**

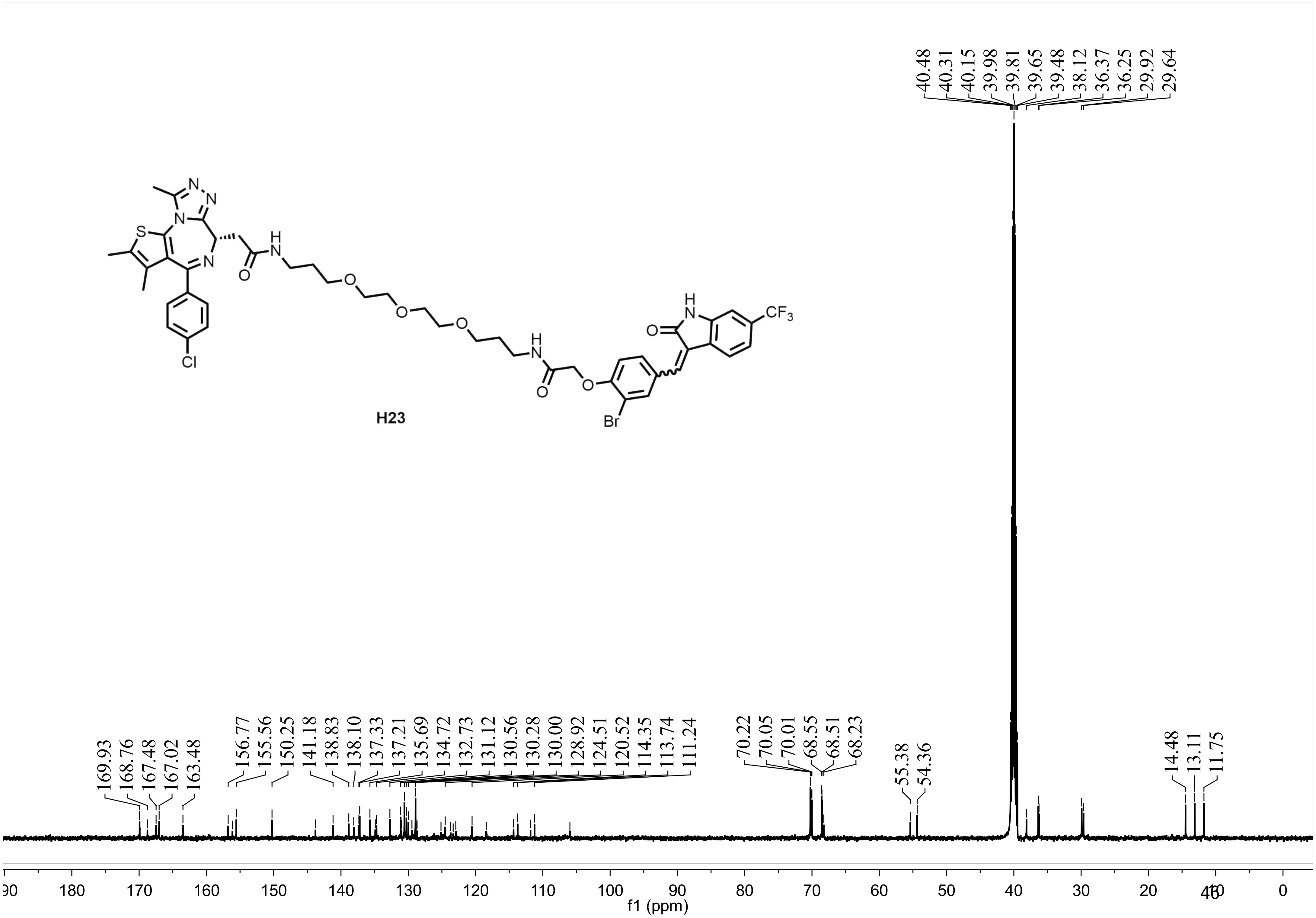
40.48
40.31
40.15
39.98
39.81
39.65
39.48
38.12
36.37
36.25
29.92
29.64
H23
CF3
Cl
Br
169.93
168.76
167.48
167.02
163.48
156.77
155.56
150.25
141.18
138.83
138.10
137.33
137.21
135.69
134.72
132.73
131.12
130.56
130.28
130.00
128.92
124.51
120.52
114.35
113.74
111.24
70.22
70.05
70.01
68.55
68.51
68.23
55.38
54.36
14.48
13.11
11.75
f1 (ppm)

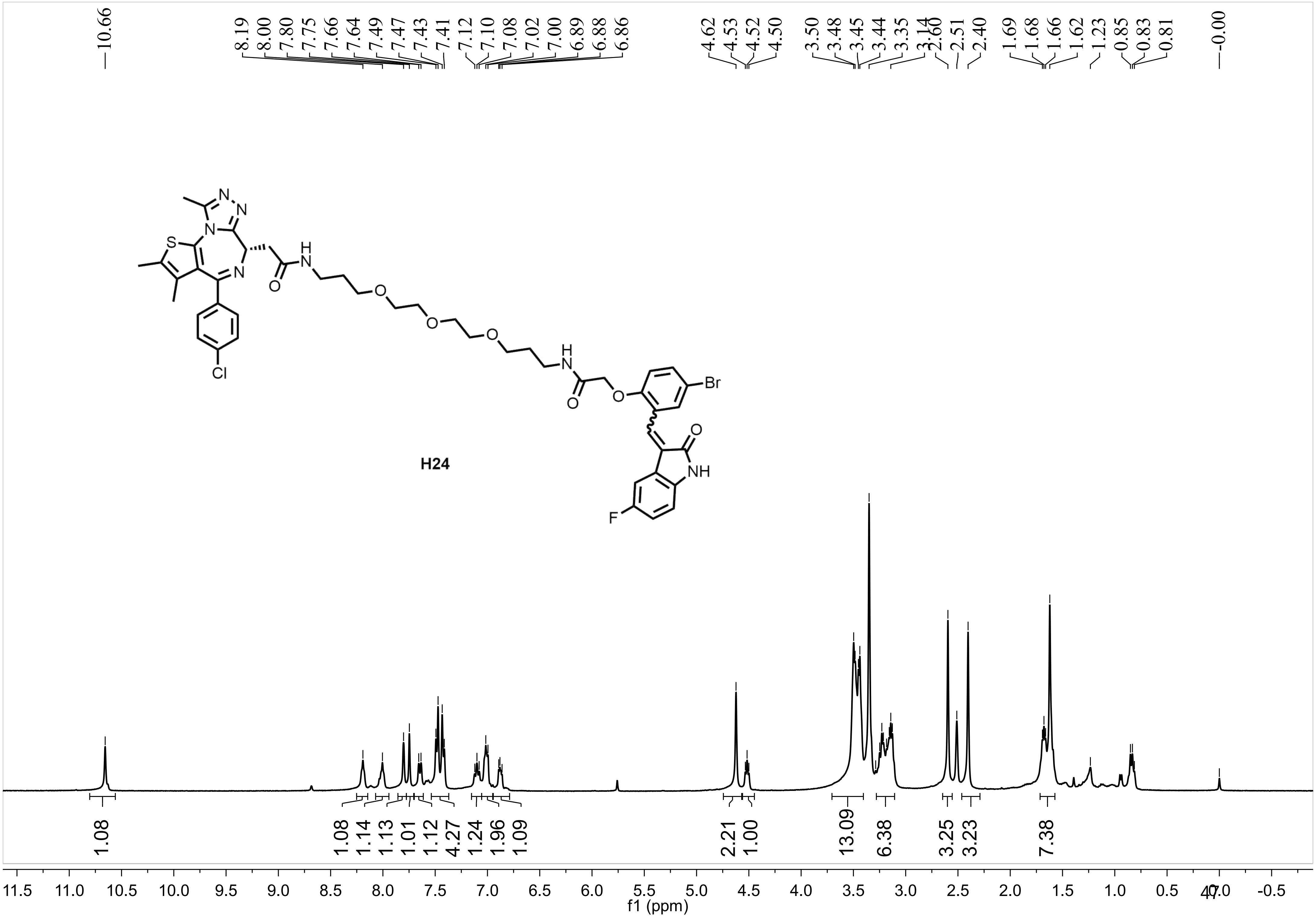
10.66
8.19 8.00 7.80 7.75 7.66 7.64 7.49 7.47 7.43 7.41 7.12 7.10 7.08 7.02 7.00 6.89 6.88 6.86
4.62 4.53 4.52 4.50 3.50 3.48 3.45 3.44 3.35 3.14 2.60 2.51 2.40 1.69 1.68 1.66 1.62 1.23 0.85 0.83 0.81
-0.00
H24
S
N
O
H
Cl
Br
NH
F
1.08
1.08 1.14 1.13 1.01 1.12 4.27 1.24 1.96 1.09
2.21 1.00
13.09
6.38
3.25 3.23
7.38
11.5 11.0 10.5 10.0 9.5 9.0 8.5 8.0 7.5 7.0 6.5 6.0 5.5 5.0 4.5 4.0 3.5 3.0 2.5 2.0 1.5 1.0 0.5 0.0 -0.5
f1 (ppm)

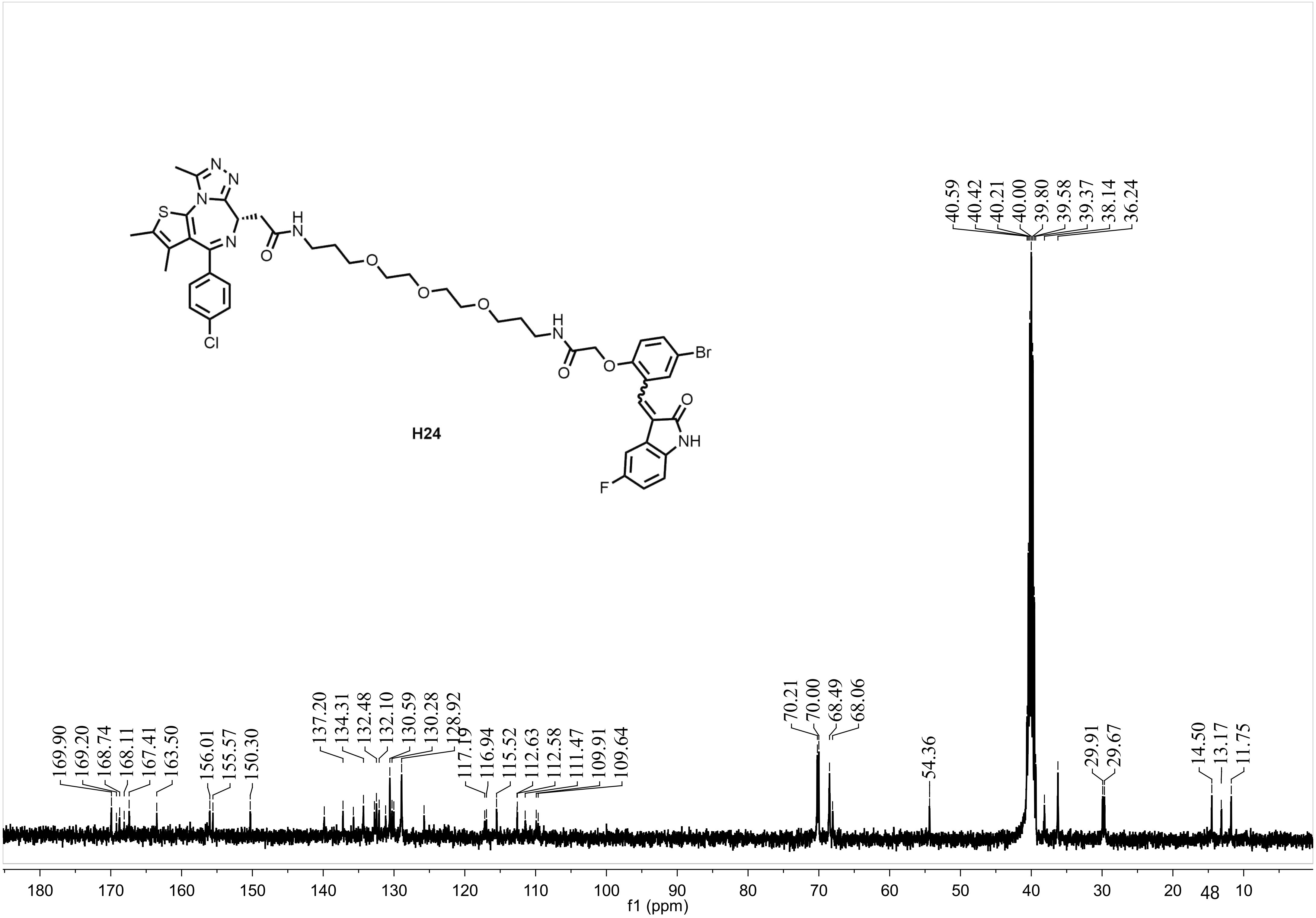
H24
Cl
S
N
O
H
NH
Br
F
40.59
40.42
40.21
40.00
39.80
39.58
39.37
38.14
36.24
169.90
169.20
168.74
168.11
167.41
163.50
156.01
155.57
150.30
137.20
134.31
132.48
132.10
130.59
130.28
128.92
117.19
116.94
115.52
112.63
112.58
111.47
109.91
109.64
70.21
70.00
68.49
68.06
54.36
29.91
29.67
14.50
13.17
11.75
180 170 160 150 140 130 120 110 100 90 80 70 60 50 40 30 20 10
f1 (ppm)

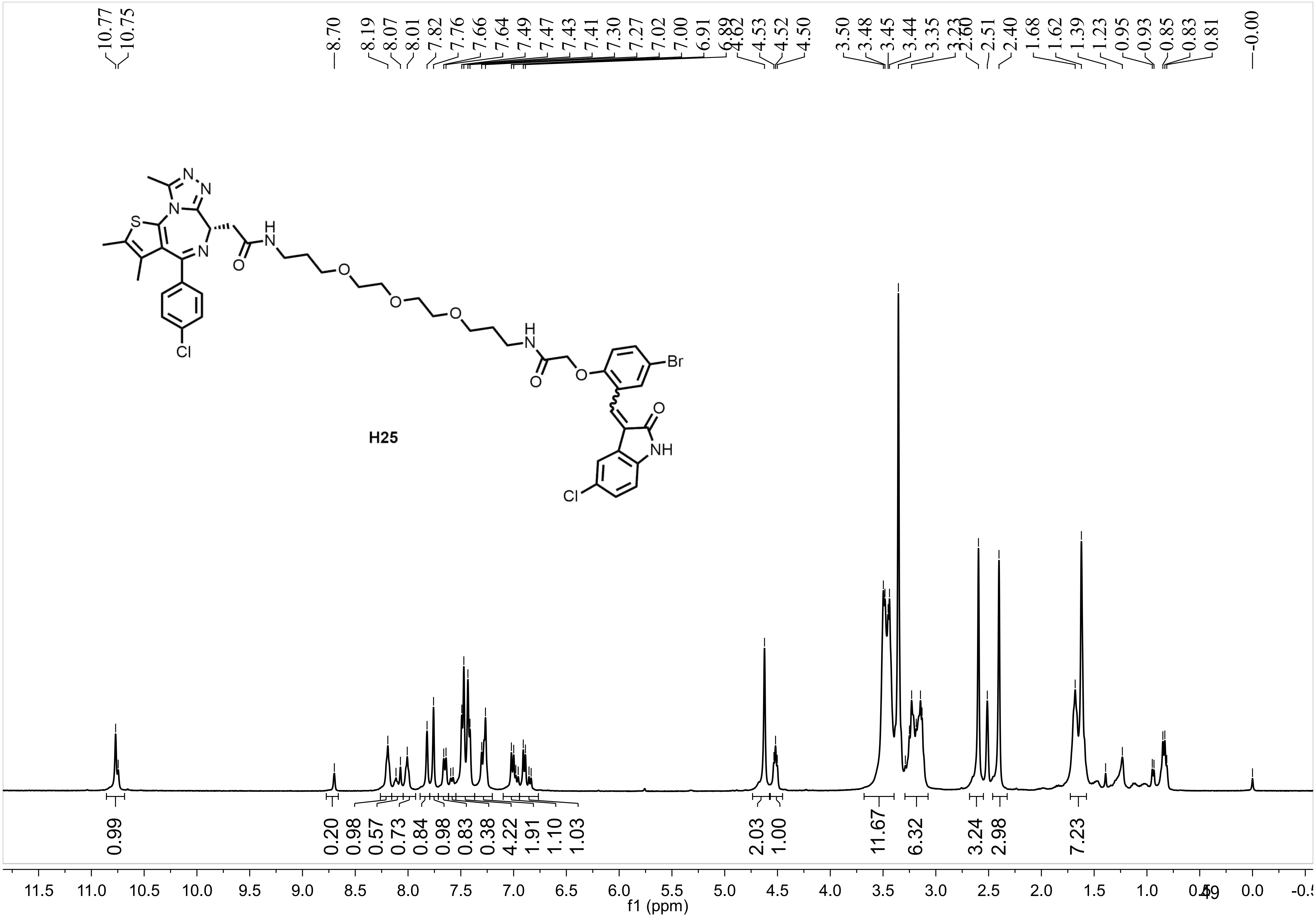
10.77
10.75
8.70
8.19
8.07
8.01
7.82
7.76
7.66
7.64
7.49
7.47
7.43
7.41
7.30
7.27
7.02
7.00
6.91
6.89
4.62
4.53
4.52
4.50
3.50
3.48
3.45
3.44
3.35
3.23
2.60
2.51
2.40
1.68
1.62
1.39
1.23
0.95
0.93
0.85
0.83
0.81
-0.00
H25
0.99
0.20
0.98
0.57
0.73
0.84
0.98
0.83
0.38
4.22
1.91
1.10
1.03
2.03
1.00
11.67
6.32
3.24
2.98
7.23
11.5 11.0 10.5 10.0 9.5 9.0 8.5 8.0 7.5 7.0 6.5 6.0 5.5 5.0 4.5 4.0 3.5 3.0 2.5 2.0 1.5 1.0 0.5 0.0 -0.5
f1 (ppm)

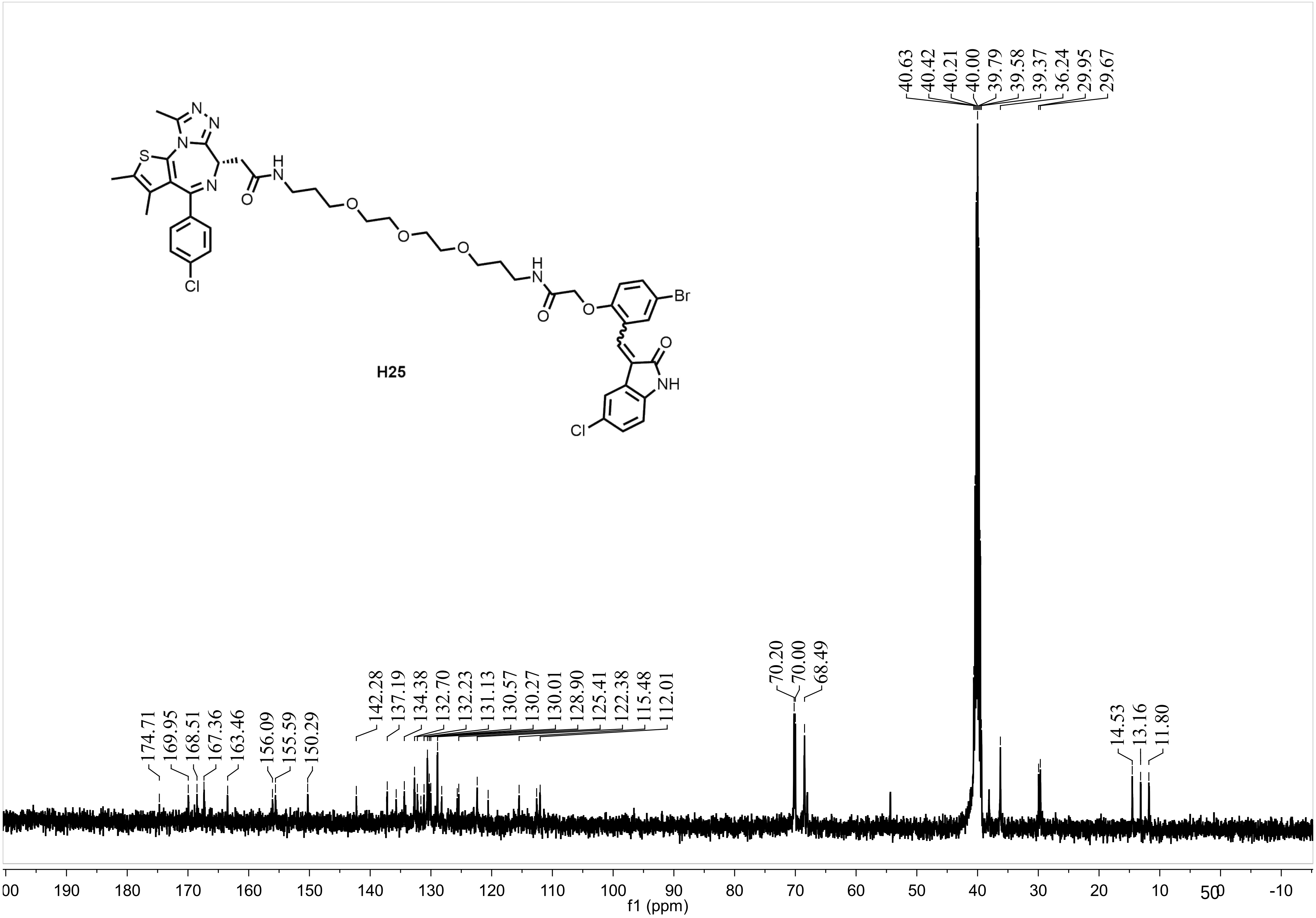
H25
N
N
N
S
N
H
N
O
O
O
O
H
N
O
O
Br
O
NH
Cl
Cl
174.71
169.95
168.51
167.36
163.46
156.09
155.59
150.29
142.28
137.19
134.38
132.70
132.23
131.13
130.57
130.27
130.01
128.90
125.41
122.38
115.48
112.01
70.20
70.00
68.49
40.63
40.42
40.21
40.00
39.79
39.58
39.37
36.24
29.95
29.67
14.53
13.16
11.80
00
190
180
170
160
150
140
130
120
110
100
90
80
70
60
50
40
30
20
10
0
-10
f1 (ppm)

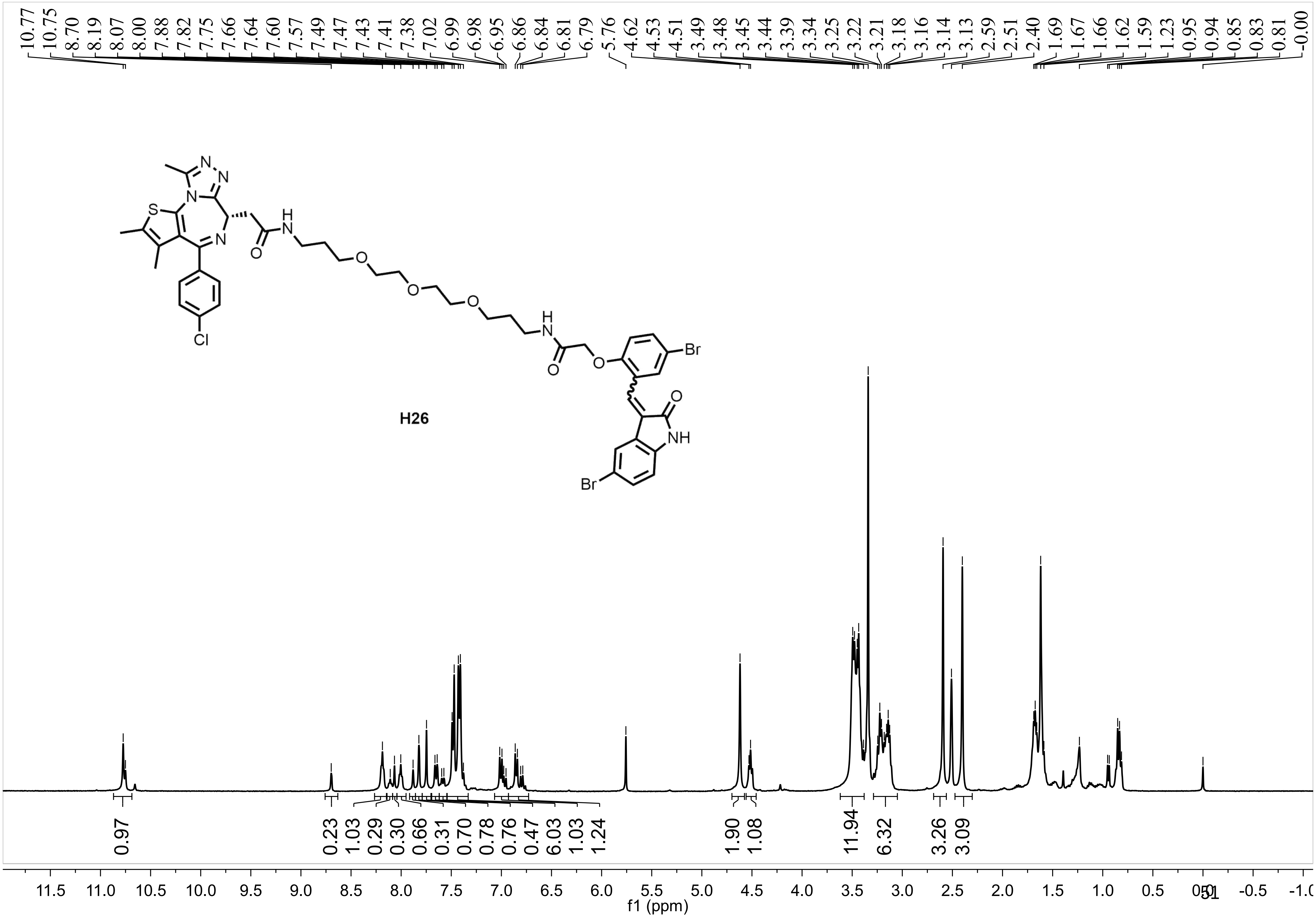
10.77 10.75 8.70 8.19 8.07 8.00 7.88 7.82 7.75 7.66 7.64 7.60 7.57 7.49 7.47 7.43 7.41 7.38 7.02 6.99 6.98 6.95 6.86 6.84 6.81 6.79 5.76 4.62 4.53 4.51 3.49 3.48 3.45 3.44 3.39 3.34 3.25 3.22 3.21 3.18 3.16 3.14 3.13 2.59 2.51 2.40 1.69 1.67 1.66 1.62 1.59 1.23 0.95 0.94 0.85 0.83 0.81 -0.00
H26
0.97 0.23 1.03 0.29 0.30 0.66 0.31 0.70 0.78 0.76 0.47 6.03 1.03 1.24 1.90 1.08 11.94 6.32 3.26 3.09
11.5 11.0 10.5 10.0 9.5 9.0 8.5 8.0 7.5 7.0 6.5 6.0 5.5 5.0 4.5 4.0 3.5 3.0 2.5 2.0 1.5 1.0 0.5 0.0 -0.5 -1.0
f1 (ppm)

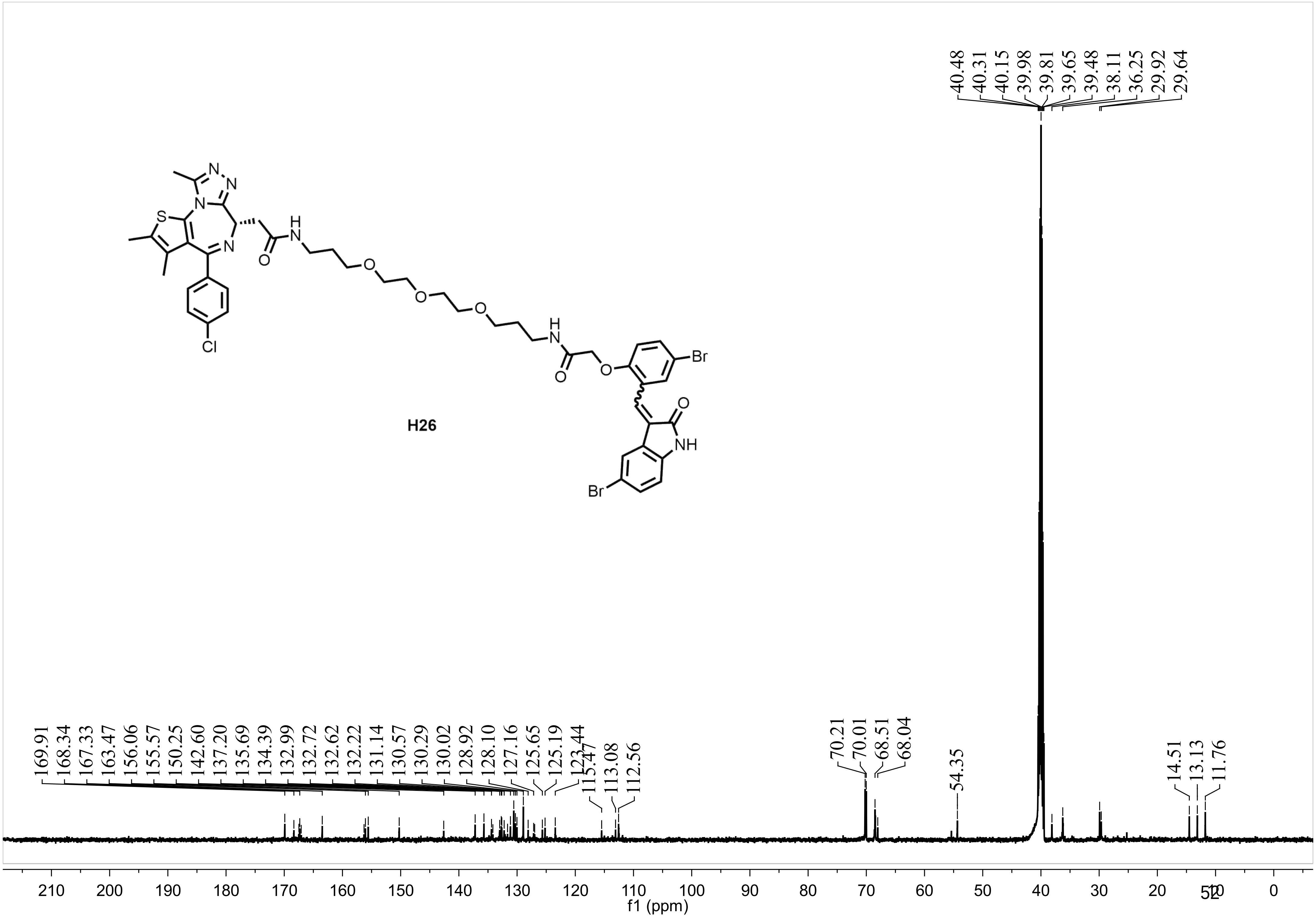
40.48
40.31
40.15
39.98
39.81
39.65
39.48
38.11
36.25
29.92
29.64
N
N
S
N
N
H
N
O
O
O
O
Cl
H
N
O
O
Br
O
NH
Br
H26
169.91
168.34
167.33
163.47
156.06
155.57
150.25
142.60
137.20
135.69
134.39
132.99
132.72
132.62
132.22
131.14
130.57
130.29
130.02
128.92
128.10
127.16
125.65
125.19
123.44
115.47
113.08
112.56
70.21
70.01
68.51
68.04
54.35
14.51
13.13
11.76
210
200
190
180
170
160
150
140
130
120
110
100
90
80
70
60
50
40
30
20
10
0
f1 (ppm)

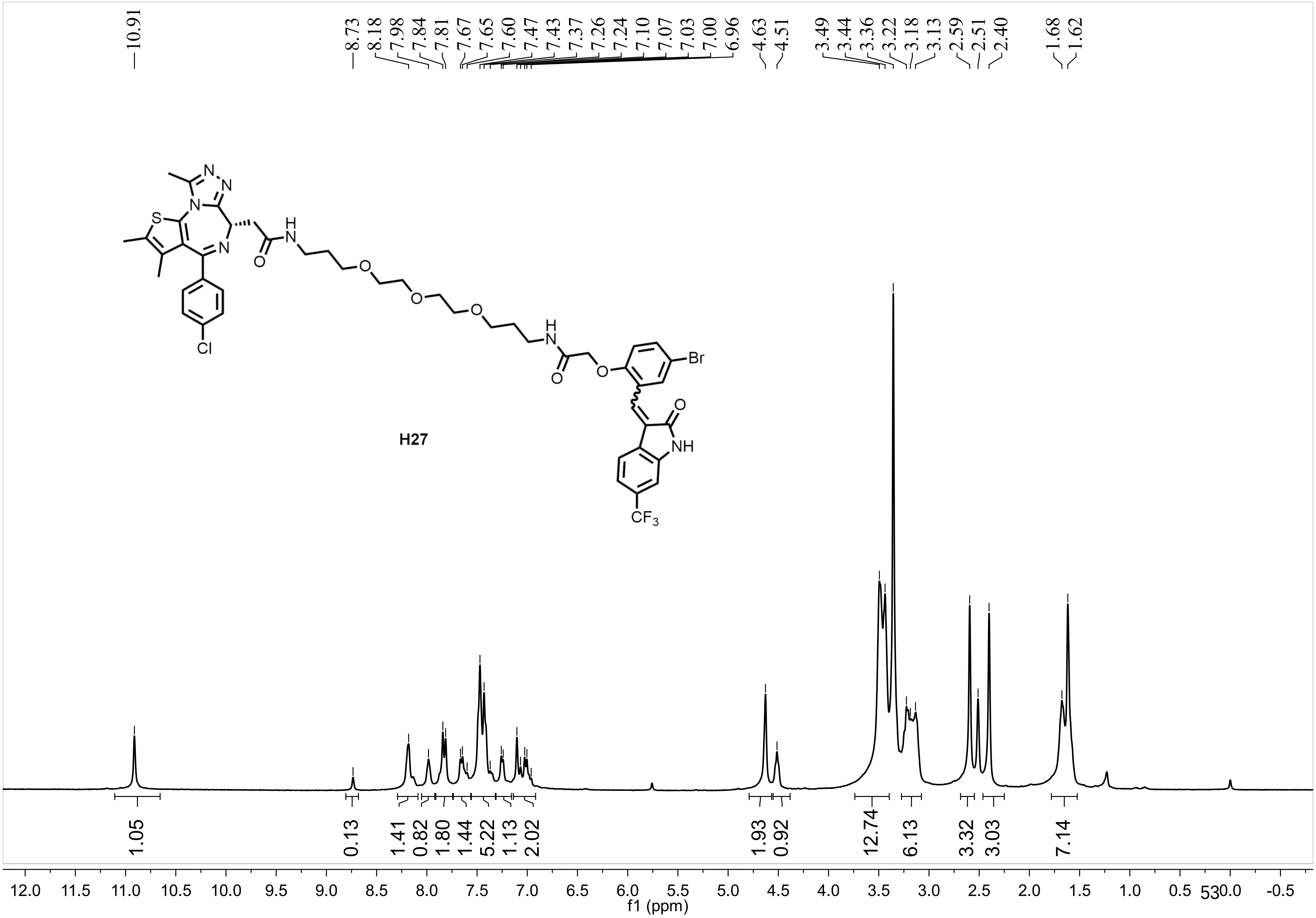
10.91
8.73 8.18 7.98 7.84 7.81 7.67 7.65 7.60 7.47 7.43 7.37 7.26 7.24 7.10 7.07 7.03 7.00 6.96
4.63 4.51
3.49 3.44 3.36 3.22 3.18 3.13 2.59 2.51 2.40
1.68 1.62
H27
N
S
Cl
O
H
Br
NH
$CF_3$
1.05 0.13 1.41 0.82 1.80 1.44 5.22 1.13 2.02 1.93 0.92 12.74 6.13 3.32 3.03 7.14
12.0 11.5 11.0 10.5 10.0 9.5 9.0 8.5 8.0 7.5 7.0 6.5 6.0 5.5 5.0 4.5 4.0 3.5 3.0 2.5 2.0 1.5 1.0 0.5 0.0 -0.5
f1 (ppm)

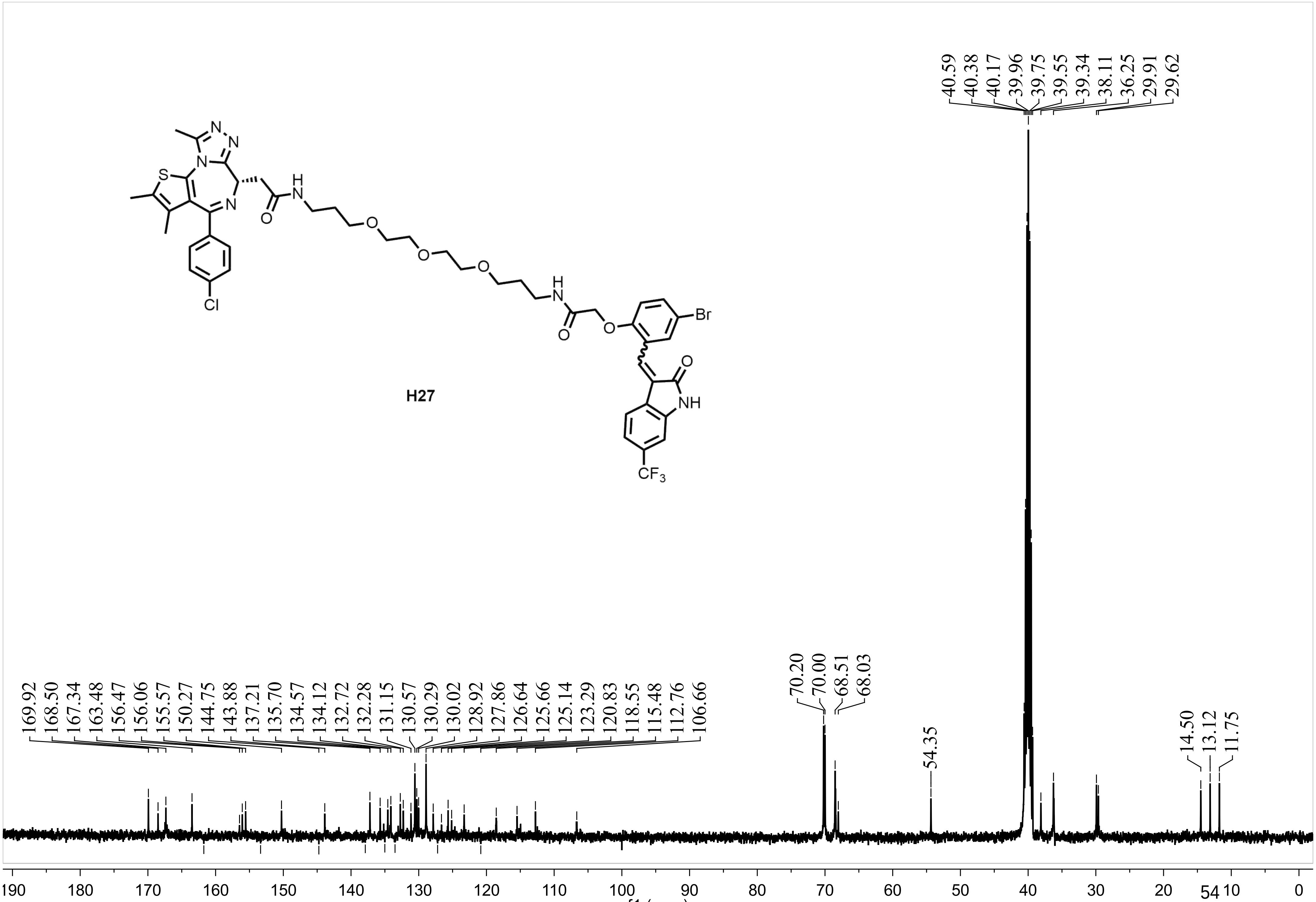
40.59 40.38 40.17 39.96 39.75 39.55 39.34 38.11 36.25 29.91 29.62
H27
Br
O
NH
CF3
Cl
S
169.92 168.50 167.34 163.48 156.47 156.06 155.57 150.27 144.75 143.88 137.21 135.70 134.57 134.12 132.72 132.28 131.15 130.57 130.29 130.02 128.92 127.86 126.64 125.66 125.14 123.29 120.83 118.55 115.48 112.76 106.66
70.20 70.00 68.51 68.03
54.35
14.50 13.12 11.75
190 180 170 160 150 140 130 120 110 100 90 80 70 60 50 40 30 20 10 0
f1 (ppm)

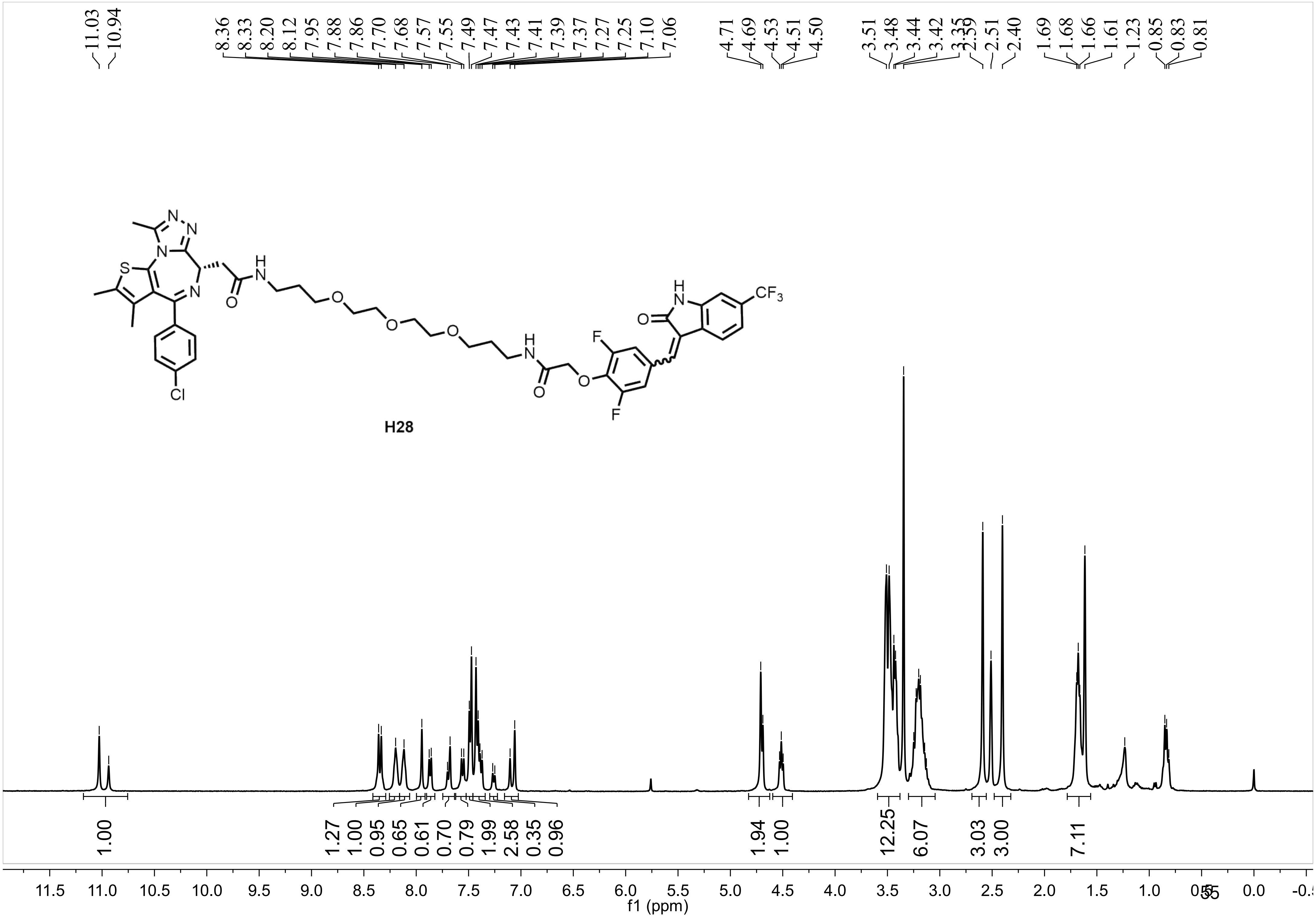
11.03
10.94
8.36
8.33
8.20
8.12
7.95
7.88
7.86
7.70
7.68
7.57
7.55
7.49
7.47
7.43
7.41
7.39
7.37
7.27
7.25
7.10
7.06
4.71
4.69
4.53
4.51
4.50
3.51
3.48
3.44
3.42
3.35
2.59
2.51
2.40
1.69
1.68
1.66
1.61
1.23
0.85
0.83
0.81
H28
1.00
1.27
1.00
0.95
0.65
0.61
0.70
0.79
1.99
2.58
0.35
0.96
1.94
1.00
12.25
6.07
3.03
3.00
7.11
f1 (ppm)

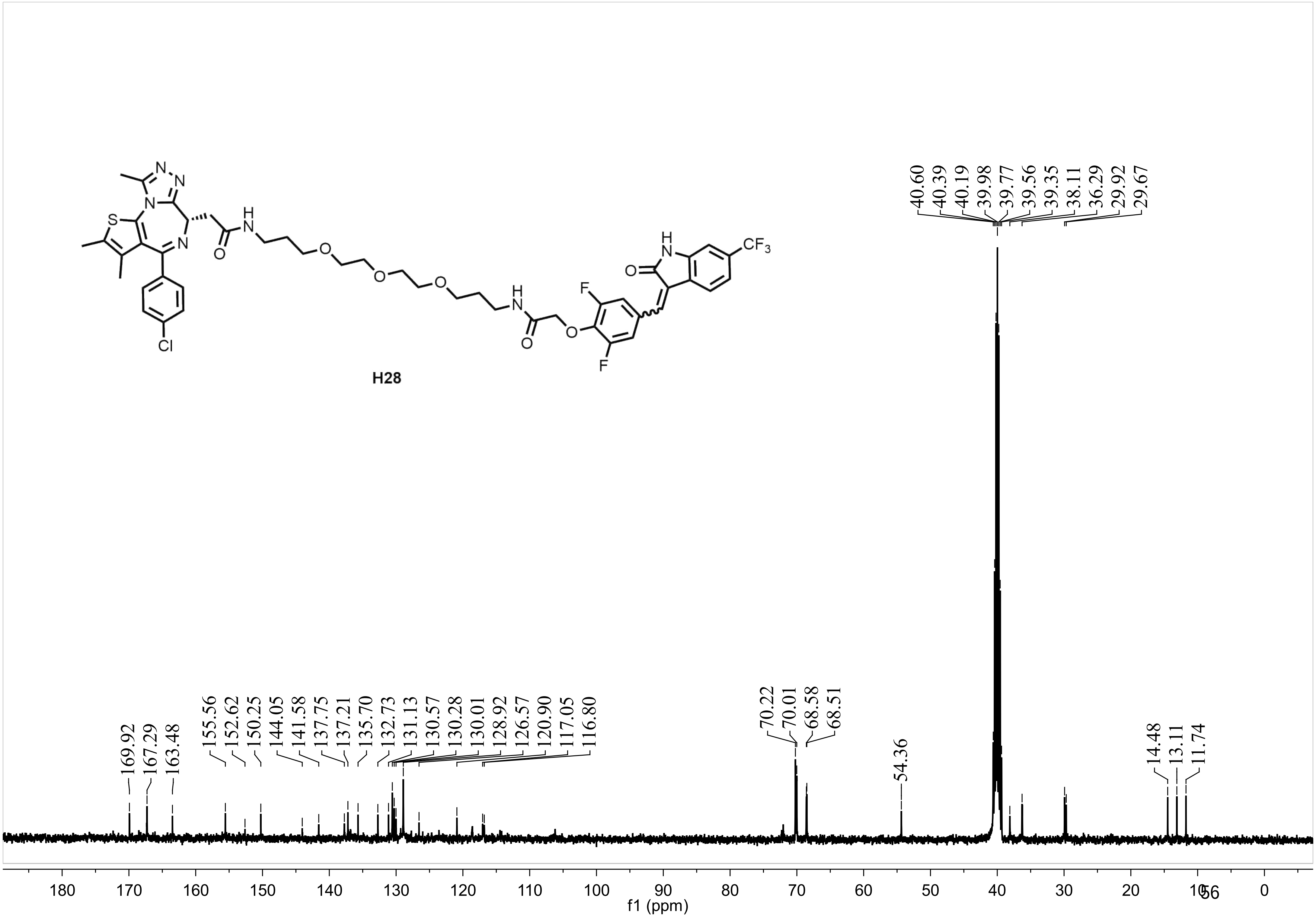
H28
169.92
167.29
163.48
155.56
152.62
150.25
144.05
141.58
137.75
137.21
135.70
132.73
131.13
130.57
130.28
130.01
128.92
126.57
120.90
117.05
116.80
70.22
70.01
68.58
68.51
54.36
40.60
40.39
40.19
39.98
39.77
39.56
39.35
38.11
36.29
29.92
29.67
14.48
13.11
11.74
180 170 160 150 140 130 120 110 100 90 80 70 60 50 40 30 20 10 0
f1 (ppm)

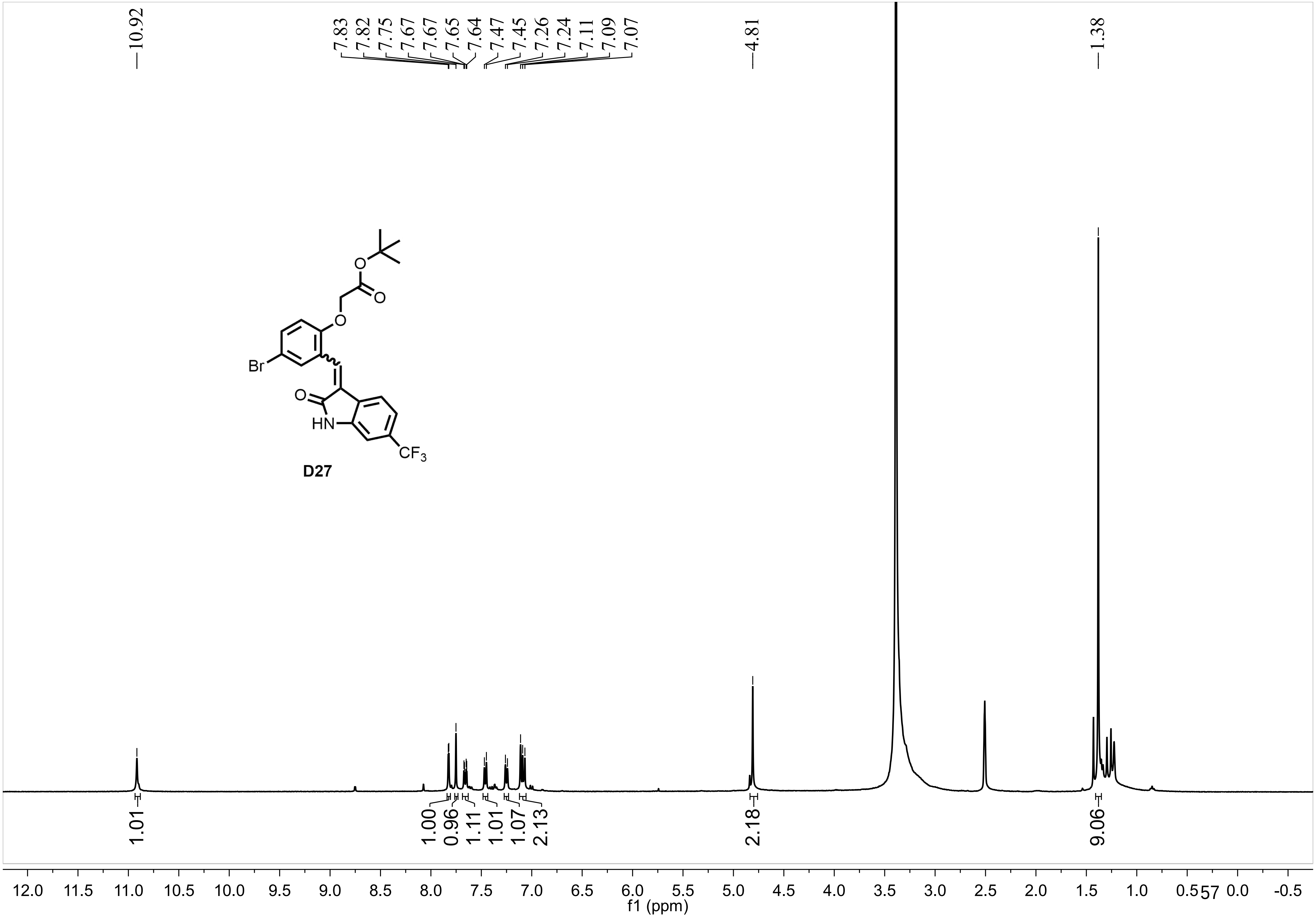
10.92
7.83
7.82
7.75
7.67
7.67
7.65
7.64
7.47
7.45
7.26
7.24
7.11
7.09
7.07
4.81
1.38
Br
O
O
O
O
HN
$CF_3$
D27
1.01
1.00
0.96
1.11
1.01
1.07
2.13
2.18
9.06
12.0 11.5 11.0 10.5 10.0 9.5 9.0 8.5 8.0 7.5 7.0 6.5 6.0 5.5 5.0 4.5 4.0 3.5 3.0 2.5 2.0 1.5 1.0 0.5 0.0 -0.5
f1 (ppm)

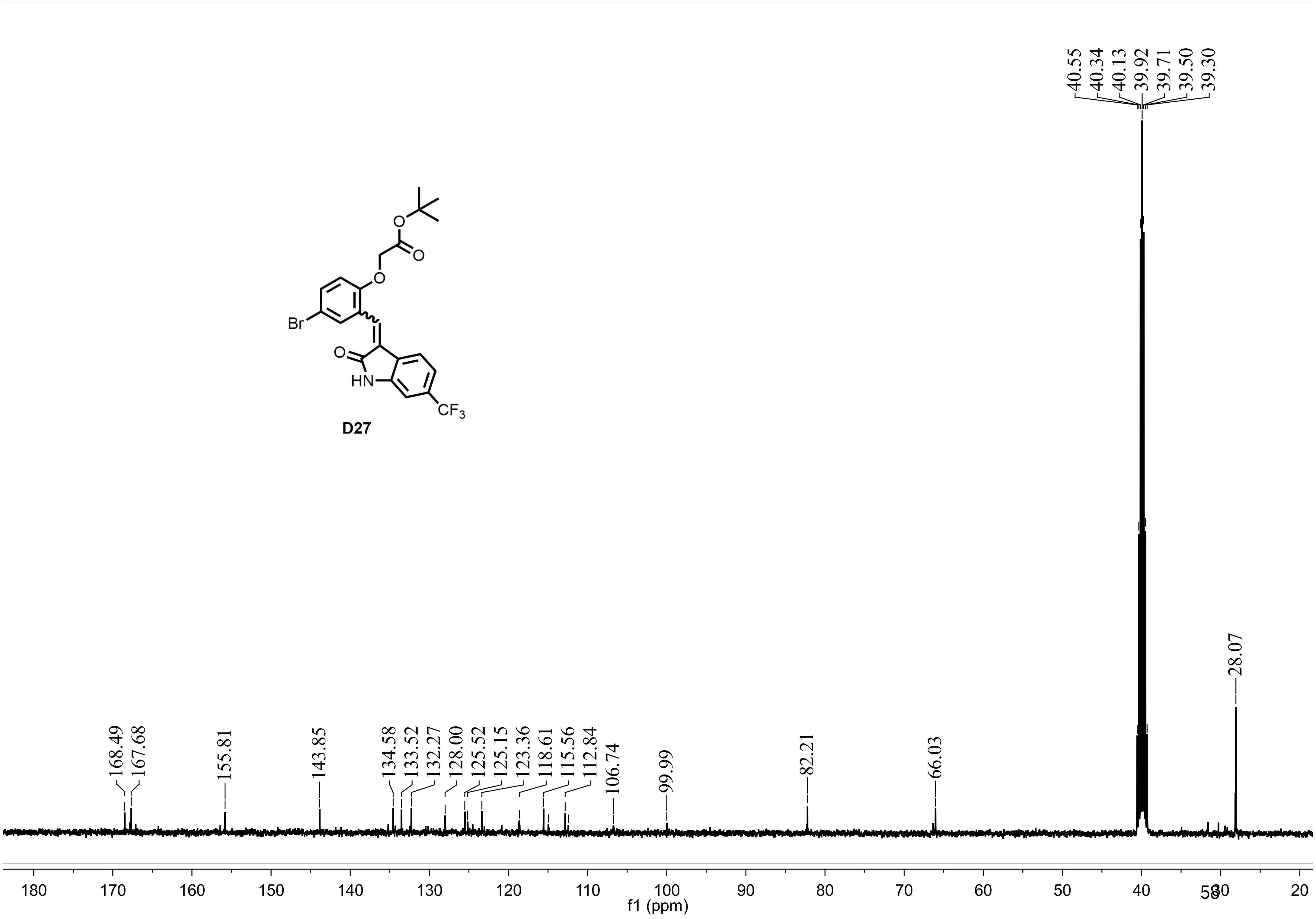
40.55
40.34
40.13
39.92
39.71
39.50
39.30
28.07
168.49
167.68
155.81
143.85
134.58
133.52
132.27
128.00
125.52
125.15
123.36
118.61
115.56
112.84
106.74
99.99
82.21
66.03
O
O
O
O
Br
O
HN
CF3
D27
180
170
160
150
140
130
120
110
100
90
80
70
60
50
40
20
f1 (ppm)
